# High-throughput functional analysis of IncRNA core promoters elucidatesrules governing tissue-specificity

**DOI:** 10.1101/482232

**Authors:** Kaia Mattioli, Pieter-Jan Volders, Chiara Gerhardinger, James C. Lee, Philipp G. Maass, Marta Melé, John L. Rinn

**Affiliations:** Department of Stem Cell and Regenerative Biology, Harvard University, Cambridge, MA, 02138, USA; Department of Biological and Biomedical Sciences, Harvard Medical School, Boston, MA, 02115, USA; Department of Biomolecular Medicine, Ghent University, 9000 Ghent, Belgium; VIB-UGent Center for Medical Biotechnology, VIB, 9000 Ghent, Belgium; Broad Institute of MIT and Harvard, Cambridge, MA, 02142, USA; Department of Medicine, University of Cambridge School of Clinical Medicine, Addenbrooke’s Hospital, Cambridge CB2 0QQ, UK; Genetics and Genome Biology Program, Sickkids Research Institute, Toronto ON, M5G 0A4, Canada; Life Sciences Department, Barcelona Supercomputing Center, Barcelona, Catalonia, 08034, Spain; Department of Pathology, Beth Israel Deaconess Medical Center, Boston, MA, 02115, USA; Department of Biochemistry, University of Colorado, BioFrontiers Institute, Boulder, CO, 80301, USA

## Abstract

Transcription initiates at both coding and non-coding genomic elements, including mRNA and long non-coding RNA (lncRNA) core promoters and enhancer RNAs (eRNAs). However, each class has different expression profiles with lncRNAs and eRNAs being the most tissue-specific. How these complex differences in expression profiles and tissue-specificities are encoded in a single DNA sequence, however, remains unresolved. Here, we address this question using computational approaches and massively parallel reporter assays (MPRA) surveying hundreds of promoters and enhancers. We find that both divergent lncRNA and mRNA core promoters have higher capacities to drive transcription than non-divergent lncRNA and mRNA core promoters, respectively. Conversely, lincRNAs and eRNAs have lower capacities to drive transcription and are more tissue-specific than divergent genes. This higher tissue-specificity is strongly associated with having less complex TF motif profiles at the core promoter. We experimentally validated these findings by testing both engineered single-nucleotide deletions and human single-nucleotide polymorphisms (SNPs) in MPRA. In both cases, we observe that single nucleotides associated with many motifs are important drivers of promoter activity. Thus, we suggest that high TF motif density serves as a robust mechanism to increase promoter activity at the expense of tissue-specificity. Moreover, we find that 22% of common SNPs in core promoter regions have significant regulatory effects. Collectively, our findings show that high TF motif density provides redundancy and increases promoter activity at the expense of tissue specificity, suggesting that specificity of expression may be regulated by simplicity of motif usage.

## Introduction

Transcription factors (TFs) regulate gene expression by binding to DNA regulatory elements at both coding and non-coding genomic elements, including mRNA and long non-coding RNA (lncRNA) promoters and enhancers. Classically, promoters and enhancers have been defined as distinct categories of regulatory elements. However, recent findings suggest that promoters and enhancers share a common regulatory code, as transcription is initiated at both (Core, Waterfall, and Lis 2008; Engreitz et al. 2016). Indeed, at both promoters and enhancers, RNA polymerase II (Pol II) binds to a 50-100bp stretch of DNA termed the “core promoter” and transcribes in both the sense and antisense directions—a phenomenon known as bidirectional transcription (Andersson 2015). Such transcription at promoters typically produces long, stable polyadenylated transcripts in the sense direction and short, unstable, non-polyadenylated transcripts in the antisense direction (Andersson 2015). At enhancers, highly unstable RNAs, named enhancer RNAs (eRNAs), are produced in a bidirectional manner (Forrest et al. 2014).

Although almost all promoters exhibit bidirectional transcription, in some cases, this bidirectional transcription results in two stable transcripts that are arranged in a “head-to-head” orientation (one on the sense strand and one on the antisense strand). These so-called “divergent” transcripts are abundant in the human genome and are evolutionarily conserved and often comprised of two highly expressed individual core promoter sequences (Trinklein et al. 2004). It remains unclear, however, whether their high expression levels are a byproduct of having two promoters in close proximity or whether it is an inherent property of their DNA sequence. Additionally, these divergent transcript pairs can also include lncRNAs, but whether divergent lncRNA promoters have distinct regulatory properties compared to divergent mRNA promoters is also unknown.

Like mRNAs, lncRNAs are transcribed by Pol II, canonically spliced, and polyadenylated. However, lncRNAs also show similarities to enhancers: they have similar chromatin environments (Marques et al. 2013) and they often act as enhancers themselves by activating the transcription of nearby genes (Ørom and Shiekhattar 2013; Rinn and Chang 2012). As a class, lncRNAs are known to be more lowly expressed and more tissue-specific than protein-coding genes (Cabili et al. 2011; Derrien et al. 2012; Molyneaux et al. 2015). Although lncRNAs are less conserved than protein-coding genes, their promoters—and the TF binding sites within their promoters—are remarkably well-conserved (Melé et al. 2017; Ponjavic, Ponting, and Lunter 2007), suggesting that a conserved regulatory logic controls lncRNA transcription. However, the rules that govern lncRNA transcription and that determine their higher tissue-specificity remain unclear. For example, it is unknown whether lncRNAs are more tissue-specific than mRNAs due to differences in their TF binding profiles.

In this work, we address the fundamental question: is there an underlying “code” in lncRNA and mRNA promoter and enhancer sequences that accounts for their established differences in tissue-specificity and abundance? To address this, we used a massively parallel reporter assay (MPRA)—in which thousands of regulatory sequences of interest are assayed in a single experiment (Patwardhan et al. 2012; Melnikov et al. 2012)—to dissect core promoter sequence properties at high resolution and across multiple cell types. MPRAs have previously uncovered important characteristics of promoters and enhancers (Nguyen et al. 2016; Arnold et al. 2017) but to date a systematic analysis of whether intrinsic features of DNA sequence are responsible for differential activity at lncRNA promoters, protein-coding gene promoters, and enhancers has not been performed. Collectively, our data converge on a model of “specificity through simplicity”— lncRNAs and enhancers have less complex TF motif profiles, and this relative simplicity is associated with their low abundance and high tissue-specificity.

### Divergent lncRNA core promoters are strong and ubiquitously expressed

We first defined five biotypes: (1) eRNAs (RNAs emerging from bidirectionally transcribed enhancers that do not overlap protein-coding genes) (2) intergenic lncRNAs (lincRNAs), (3) divergent lncRNAs (lncRNAs that share promoters with either a protein-coding gene or another lncRNA in the antisense direction), (4) mRNAs, and (5) divergent mRNAs (mRNAs that share promoters with either another protein-coding gene or a lncRNA in the antisense direction) (**Figure 1A; see methods**). Note that here the term “divergent” refers to the presence of a stable annotated transcript in the antisense direction, not the potential bidirectionality of the promoter itself. As the TSSs of lncRNAs can be more poorly annotated than the TSSs of mRNAs, which could bias results when comparing them (Lagarde et al. 2017), we carefully selected a set of high-confidence TSSs defined by the FANTOM5 consortium across all biotypes. Specifically, we used the stringent set of enhancer TSSs (for eRNAs) and promoter TSSs (for the remaining biotypes) defined as ‘robust’ in the FANTOM5 project (Andersson et al. 2014; Forrest et al. 2014). For the promoter TSSs, we only considered TSSs that were within 50bp of an annotated gene start site (see methods). In total, our genome-wide set of core promoter regions included 29,807 eRNAs, 4,280 lincRNAs, 1,713 divergent lncRNAs, 14,332 mRNAs, and 4,235 divergent mRNAs. Analysis of Cap Analysis of Gene Expression followed by sequencing (CAGE-seq) data across 550 tissues and cell types (973 samples) for each TSS confirmed that mRNAs were more highly expressed and less tissue-specific than lncRNAs and eRNAs (**Supplemental Fig S1**). Additionally, for both lncRNAs and mRNAs, divergent transcripts were more highly expressed and less tissue-specific than their non-divergent counterparts (**Supplemental Fig S1**; see supplemental methods, genome-wide analysis section).

**Figure 1:**
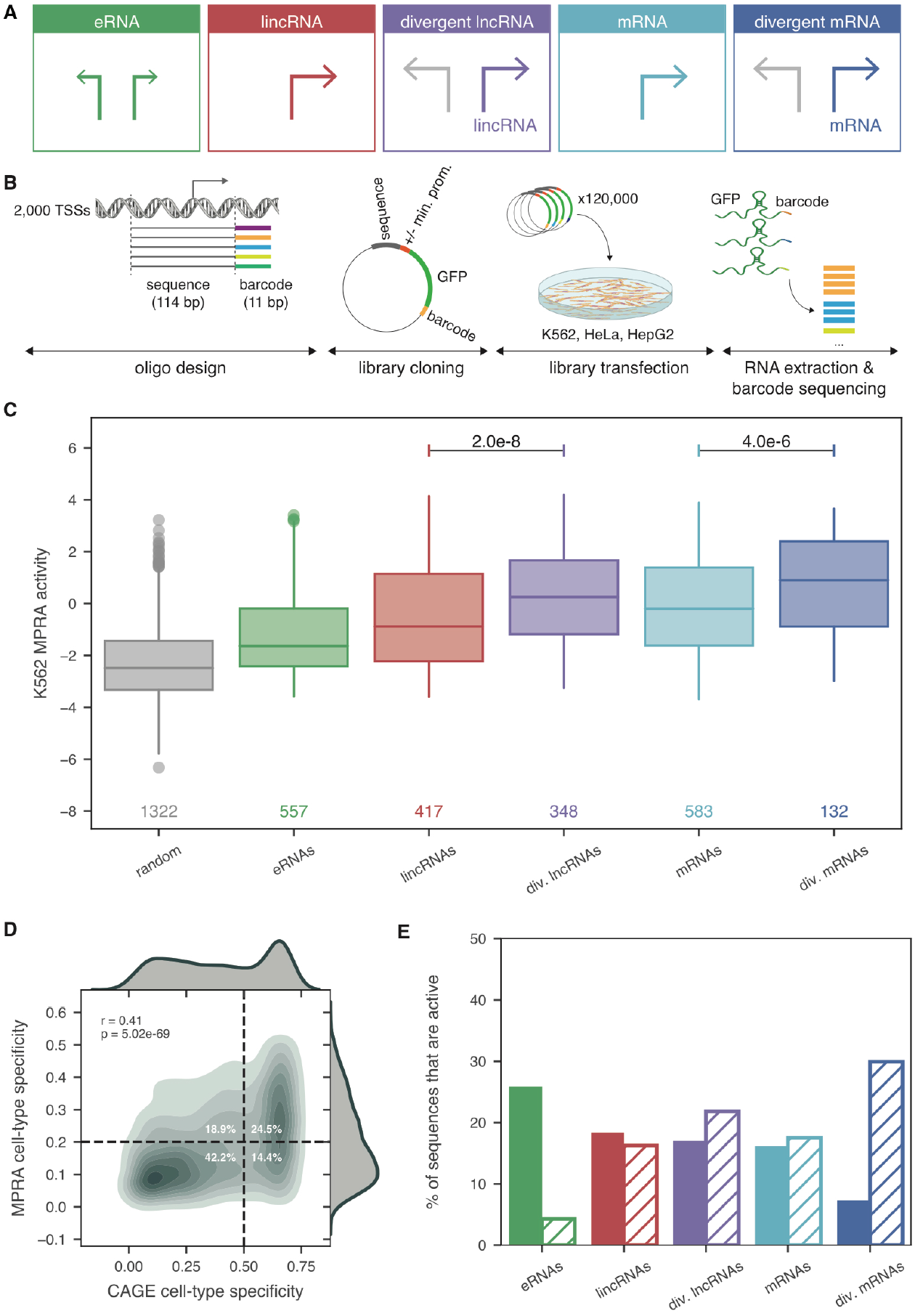
Core promoter sequences of different TSS classes vary in strength and cell-type-specificity. **A.** Overview of TSS classification based on element class (lncRNA, mRNA, or eRNA) and presence or absence of a divergent stable transcript arising from the same promoter region on the antisense strand. **B.** Schematic of MPRA experimental design. “Min. prom.”: minimal promoter. **C.** Comparison of MPRA activities (foldchange between normalized RNA barcodes and input DNA barcodes) of the reference sequences of each TSS class to negative control sequences in K562 cells. Only TSSs which meet the quality criteria of >= 3 barcodes represented each with >= 5 DNA and RNA counts are plotted and *n* values are shown. *P*-values listed are from a two-sided Wilcoxon test. **D.** Correlation between CAGE cell-type-specificity calculated across HeLa, Hepg2, and K562 (x axis) and MPRA cell-type-specificity across the same 3 cell lines (y axis). The upper right and lower left quadrants correspond to sequences that agree with CAGE and MPRA and make up 67% of sequences. Dashed horizontal and vertical line thresholds for specificity were determined from the distribution of specificity values, shown as density plots on the top and to the right of the main plot. Spearman’s rho and p-value are shown. **E.** Percent of sequences that are active in only one cell type (solid bars) or all three cell types (K562, HepG2 and HeLa; hatched bars) within each biotype.

To experimentally test the above computational predictions and dissect the contribution of DNA sequence to the observed expression and tissue-specificity patterns, we designed an MPRA in which we could assess the activity of 2,078 unique TSSs encompassing all 5 biotypes (564 eRNAs, 525 lincRNAs, 353 divergent lncRNAs, 599 mRNAs, and 137 divergent mRNAs) expressed across 3 diverse human cell lines: K562 (chronic myelogenous leukemia), HepG2 (liver carcinoma), and HeLa (cervical adenocarcinoma) (**Figure 1A; Figure 1B; Supplemental Table S1**; see methods). Since most TF motifs and ChIP-seq peaks were enriched near the TSS (**Supplemental Fig S2**), we designed oligonucleotides that spanned the core promoter (from 80 bp upstream to 34 bp downstream of the TSS; see methods). We linked each core promoter to a minimum of 15 unique 11-nucleotide barcodes to ensure redundancy across sequencing measurements (**Supplemental Table S2**). We performed a minimum of 4 replicates and a maximum of 12 replicates per condition. We measured a sequence’s ability to drive transcription—termed “MPRA activity”—by calculating the foldchange between RNA barcode counts and input DNA library barcode counts after normalizing for sequencing depth (see methods). MPRA activity measurements across replicates within a given condition were highly correlated (**Supplemental Fig S3**).

We first validated the MPRA by comparing core promoter activity measurements to negative controls; as expected, core promoters were significantly more active than random sequences in all three cell types (**Figure 1C; Supplemental Fig S4**). In general, MPRA activities correlated well with endogenous CAGE-seq expression (**Supplemental Fig S5**). eRNAs had the lowest activity, followed by lincRNAs, which is consistent with the CAGE-seq results and indicates that lincRNA core promoters are stronger than eRNA core promoters (**Figure 1C; Supplemental Fig S4**). Interestingly, as we saw using CAGE-seq expression, we found that divergent mRNAs were more active than non-divergent mRNAs and that divergent lncRNAs were more active than intergenic lncRNAs (**Figure 1C; Supplemental Fig S4**). This implies that, on average, an individual divergent promoter is stronger than an individual non-divergent promoter. Therefore, the higher CAGE-seq expression levels observed in divergent lncRNAs compared to lincRNAs cannot solely be explained by having two promoters in close proximity. When looking at expression-matched core promoters only, these results were substantially weakened (**Supplemental Fig S6**), indicating that we are capturing innate expression differences between biotypes.

We further tested whether our MPRA could recapitulate endogenous cell-type-specificity patterns. Briefly, we recalculated tissue-specificity values using K562, HepG2, and HeLa CAGE-seq expression data only (termed “cell-type-specificity”) and found that 67% of sequences agreed in CAGE-seq and MPRA cell-type-specificity designations (i.e., were classified as either specific in both or non-specific in both) (**Figure 1D**). Consistently, eRNAs and lincRNAs were more tissue-specific than mRNAs and divergent transcripts (**Figure 1E**). Thus, the DNA sequences of core promoter regions alone drive part of the tissue-specificity pattern that is present endogenously.

We next sought to determine whether differences in expression patterns between biotypes are associated with known core promoter elements. Core promoters are often classified into two types: ubiquitously-expressed promoters—associated with CpG islands and a depletion of TATA box motifs—and tissue-specific promoters—enriched for TATA box and Initiator (Inr) motifs (Medina-Rivera et al. 2018). As expected, we found that more ubiquitously-expressed biotypes had higher CpG content (**Supplemental Fig S7A**). All biotypes had similar numbers of sequences containing Inr motifs (**Supplemental Fig S7B**). Very few sequences (~3%) had canonical TATA box motifs, which are traditionally associated with tissue-specific expression. Intriguingly, while eRNAs and lincRNAs had more TATA boxes than divergent lncRNAs and divergent mRNAs, mRNAs had a relatively high number of TATA boxes and equally high numbers of both TATA boxes and Inr motifs together (**Supplemental Figs S7C, D**). Thus, it would appear that tissue-specificity cannot be explained by core promoter elements alone, as mRNAs—which are more ubiquitously-expressed than eRNAs and lincRNAs—are enriched for more canonical tissue-specific core promoter elements such as the TATA box.

### Fewer overlapping TF motifs in lincRNAs and enhancers contribute to their lowerexpression levels and higher cell-type-specificity

Our earlier results showed that MPRA can partially re-capitulate endogenous patterns of gene expression, including abundance—for which MPRA activity is a proxy—and cell-type-specificity. Therefore, we aimed to further understand what sequence features could be contributing to the lower abundance and higher tissue-specificity of eRNA and lincRNA core promoters. To that end, we focused on two main features: TF motif architecture within a core promoter sequence and the cell-type-specificity of the TFs themselves that are present within a core promoter sequence. To determine core promoter TF motif architecture, we first mapped motifs (corresponding to 519 TFs) within our core promoter sequences using FIMO (Grant, Bailey, and Noble 2011). Since the presence of a computationally-predicted motif does not always indicate physiological binding of the TF (Wasserman and Sandelin 2004), we then intersected these mapped motifs with ChIP-seq peaks corresponding to 771 TFs (218 of which we had motifs for) (Mei et al. 2017) and considered only the motifs that overlap a corresponding ChIP-seq peak (see methods). We divided TF motif architecture into two components: (1) the number of independent motif binding sites in linear sequence space and (2) the number of overlapping motifs, which should be proportional to the number of different TFs that can bind to a specific sequence pattern. As a proxy for the number of independent motif binding sites, we used the number of base pairs covered by at least one motif in a given sequence. As a proxy for the number of overlapping motifs, we used the maximum coverage of motifs per sequence. As a proxy for the cell-type-specificity of the TFs themselves, we calculated the mean cell-type-specificity (across HepG2, HeLa, and K562) of all TF motifs within a given promoter (**Figure 2A**).

**Figure 2:**
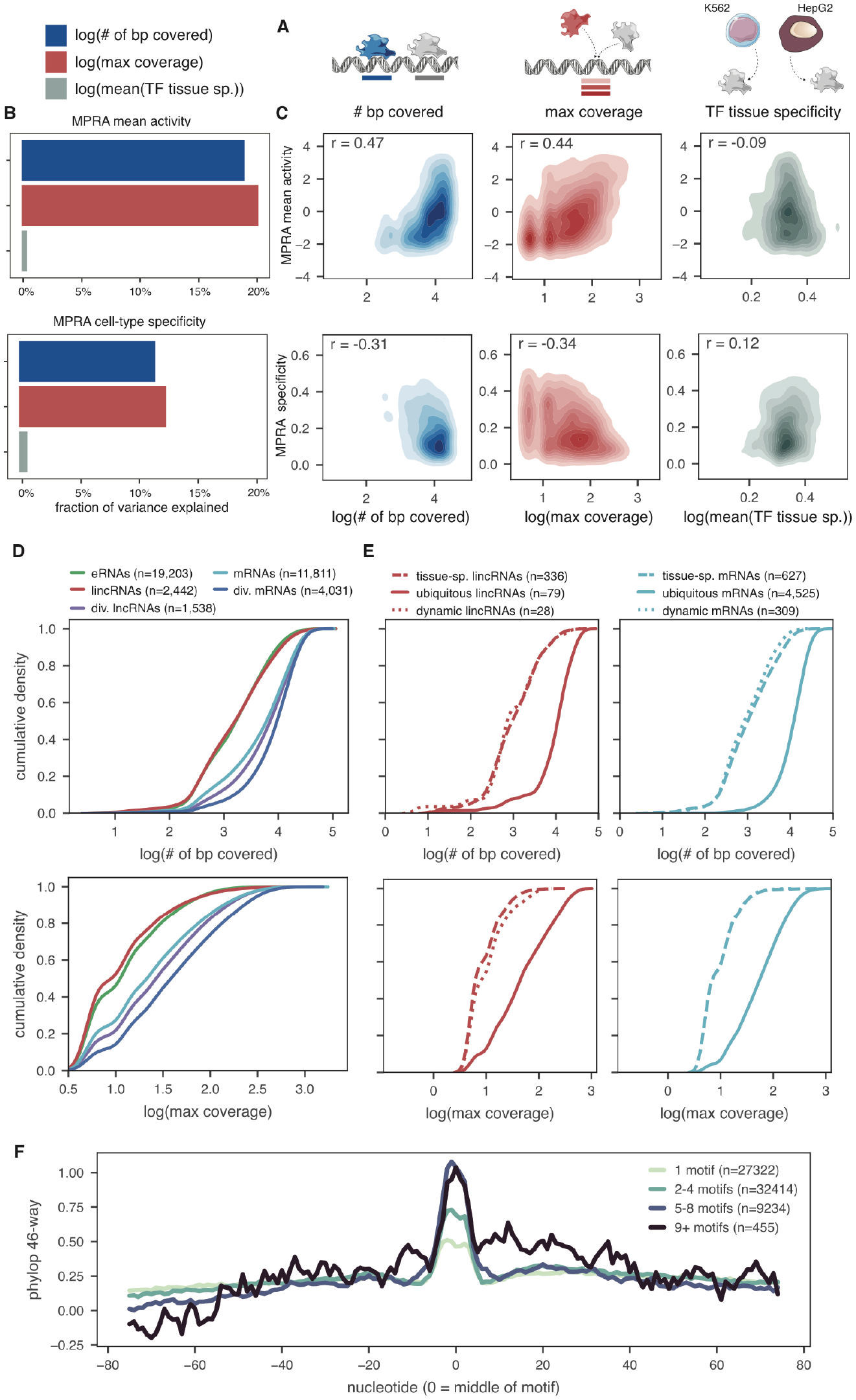
Coverage of TFs within a binding site explains expression levels and cell-type-specificity variability. **A.** Schematic of the three metrics used to model the capacity to drive transcription and cell-type specificity in the MPRA. For each metric, only TF motifs that have been validated by ChIP (i.e., overlap a ChIP peak for the cognate TF) are considered. **B.** Fraction of variance explained by each of the three metrics for either mean MPRA activity (top) or MPRA cell-type-specificity (bottom). **C.** Correlation between the three metrics (x axis) and either the mean MPRA activity (top) or MPRA cell-type-specificity (bottom) across HeLa, HepG2, and K562. Spearman’s rho is shown. **D.** Cumulative density plot of the number of base pairs covered by a motif across all biotypes (top) and the maximum motif coverage across all biotypes (bottom). **E.** Cumulative density plot for number of base pairs covered (top) and maximum motif coverage (bottom) either within lincRNAs (left) or within mRNAs (right), looking only at TSSs that are defined as tissue-specific (“tissue-sp.”), ubiquitous, or dynamically expressed (see text). **F.** Metaplot of the average phyloP 46-way placental mammal conservation score centered on motif regions, broken up by how many individual TF motifs map to the region. In all plots, only sequences with at least 1 validated motif were considered.

To test the relative importance of these three components (number of base pairs covered by a motif, maximum coverage of motifs, and average TF cell-type-specificity), we calculated the proportion of the variation in both MPRA activity and MPRA cell-type-specificity that can be explained by each measurement using a general linear model (see methods). Interestingly, the number of overlapping motifs explains a slightly higher proportion of the variation than the number of base pairs covered by a motif when predicting either mean MPRA activity or MPRA cell-type-specificity (**Figures 2B, C**). Conversely, the cell-type-specificities of the TFs themselves explain relatively little of the variation in MPRA activity and cell-type-specificity (**Figures 2B, C**). We also evaluated how much individual TF motifs contribute to sequence activity (see methods). No single TF motif was able to explain more than 1.5% of the variation (**Supplemental Fig S8B**). Overall, our model suggests that having highly overlapping motifs is substantially predictive of higher transcriptional activity and decreased cell-type-specificity.

Next, we looked at the motif architecture in biotypes that are known to be tissue-specific— such as eRNAs and lincRNAs—compared to biotypes that are known to be ubiquitous—such as mRNAs and divergent genes. We observed that tissue-specific biotypes had both fewer base pairs covered by a motif and fewer overlapping motifs than ubiquitously-expressed biotypes (**Figure 2D**). We then classified individual lincRNA and mRNA TSSs as being either ubiquitously-expressed (>0 CAGE tpm in >90% of samples), tissue-specifically expressed (>0 CAGE tpm in <10% of samples), or dynamically-expressed (a subset of tissue-specific genes, where in at least 1 sample the TSS is expressed at >50 CAGE tpm). Ubiquitously-expressed TSSs within each biotype had both more base pairs covered by a motif and more overlapping motifs than tissue-specific and dynamic TSSs (**Figure 2E**).

Some TF motifs are highly similar to each other, creating potential redundancies in motif databases (Mathelier et al. 2014). To control for this, we used two independent methods to cluster similar motifs. First, we performed unbiased clustering of the 519 motifs using MoSBAT (Lambert et al. 2016), resulting in 223 motif clusters (**Supplemental Fig S9A**). Second, we used a list of 108 non-redundant 8mer motifs generated using protein binding microarrays across 671 TFs (Mariani et al. 2017). We then re-calculated the above metrics (number of base pairs covered by a motif and the maximum motif coverage) after removing each set of redundant motifs. We found that for both metrics, ubiquitously-expressed biotypes had higher maximum coverage values than tissue-specific biotypes (**Supplemental Figs S9B, S9C, S10, S11**). Moreover, DNA regions that harbor many overlapping TF motifs are more conserved than those harboring only one TF motif (**Figure 2F**). Thus, using computationally-mapped TF motifs, endogenous TF binding events via ChIP-seq, and unique TF clusters, we observe that high and ubiquitous expression is correlated with many overlapping motifs.

### Targeted deletions refine functional TF motifs in lncRNA promoters

Our results suggest that overlapping TF motifs that can be bound by many different TFs— potentially in different contexts—are associated with increased expression and decreased tissue-specificity. We thus hypothesized that disruption of highly overlapping motifs should have larger effect sizes than disruption of more specific motifs. To test this, we performed a second MPRA across the core promoters of 21 disease-associated lncRNAs, 5 nearby mRNAs, and 5 nearby eRNAs (**Supplemental Table S3**) and tested the effect of single-nucleotide deletions across each core promoter in HepG2 and K562 cells (**Figure 3A; Supplemental Table S4**). To ensure that we covered all motifs surrounding the TSS, we included 2 tiles for each TSS (from 183 bp upstream to 69 bp upstream and 89 bp upstream to 25 bp downstream of the TSS). Thus, this strategy allows us to assess the contribution of each individual nucleotide to core promoter activity independently in a single experiment (Patwardhan et al. 2009).

**Figure 3.**
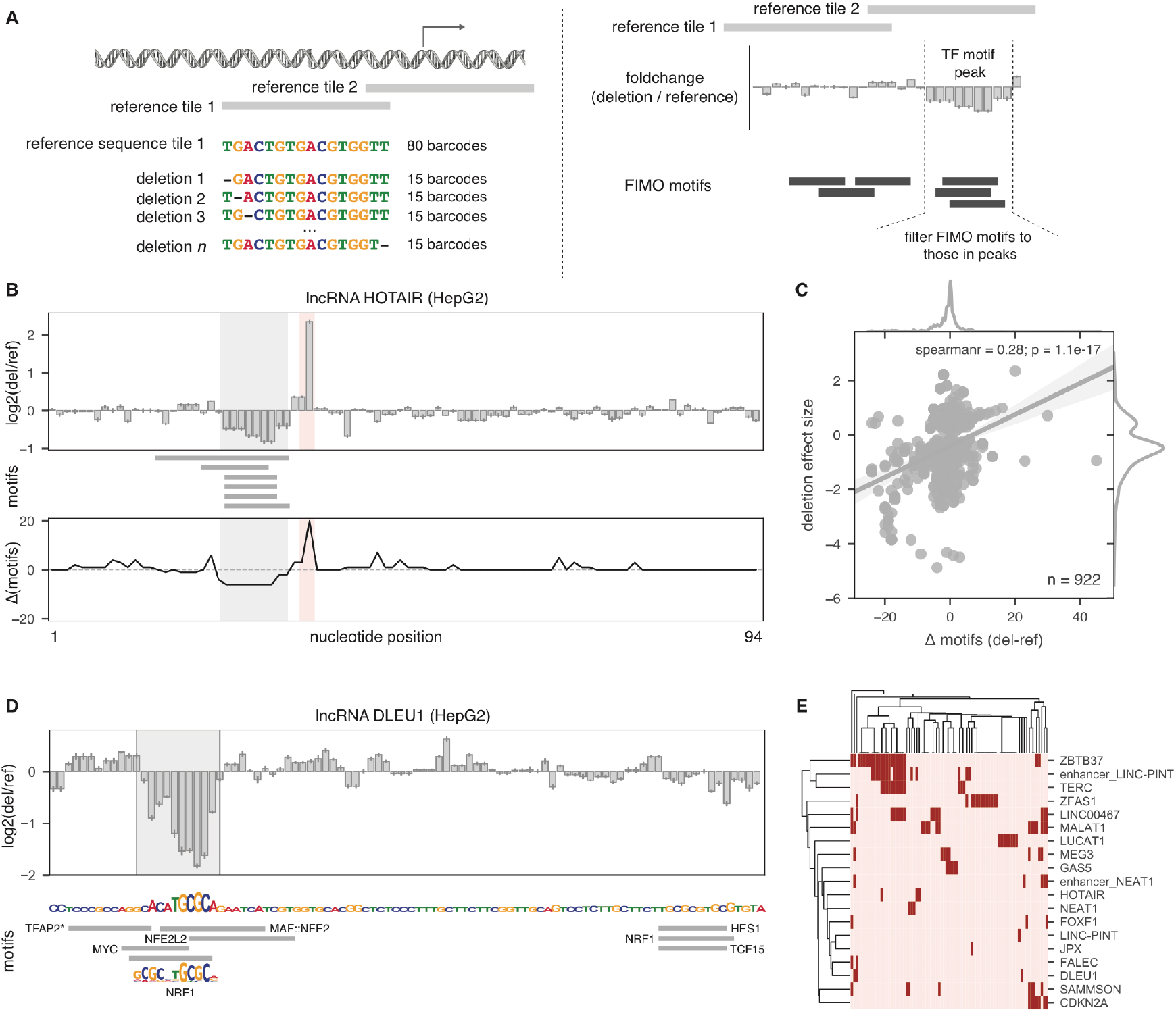
Targeted deletions refine TF motifs within lncRNA promoters. **A.** Schematic of single-nucleotide deletions MPRA design (left) and the output interpretation (right). **B.** MPRA deletion profile for the lncRNA HOTAIR promoter (top), the positions of computationally-mapped motifs in the reference sequence (middle), and the number of motifs predicted to be gained or lost due to the single-nucleotide deletions (bottom). Shaded areas represent the strongest gain (red) or loss (gray) of activity. **C.** Correlation between the number of motifs predicted to be disrupted (x axis) and the effect size of deletions (y axis) for all significant deletions in HepG2. **D.** (top) MPRA deletion profile for the lncRNA DLEU1 promoter. Shaded area is a called peak. (bottom) DLEU1 sequence (plotted with letter heights proportional to loss of activity in the MPRA) and computationally-mapped motifs (gray boxes). The sequence logo for NRF1 is shown. * = TFAP2A, TFAP2B, and TFAP2C all map to the noted gray box. **E.** Heatmap showing all computationally-mapped motifs that overlap deletion peaks in HepG2.

First, we confirmed that test core promoter sequences had significantly more activity than negative control sequences in both cell types (**Supplemental Fig S12**). Next, we calculated the “effect size” of each deletion as a foldchange in MPRA activity relative to the full reference sequence. In order to determine how deletion effect sizes correlate with TF motif profiles, we calculated the number of computationally-mapped motifs that are lost (or gained) in each deletion sequence relative to the full reference sequence (**Figure 3B**). Individual nucleotides that overlap a predicted motif are important in maintaining transcription, as deletion of each nucleotide independently shows a strong loss of activity (**Figure 3B**, gray shaded area). Additionally, we also saw deletions with gain of function effects—for example, deleting a single nucleotide in the lncRNA *HOTAIR* core promoter is predicted to create 20 new TF motifs and causes a strong increase in activity (**Figure 3B**, red shaded area). These observations extended to the remaining core promoters: deletion effect sizes were generally correlated with the number of motifs computationally predicted to be affected by each deletion (**Figure 3C; Supplemental Fig S13A**).

Moreover, single-nucleotide deletions can be used to better identify functional DNA regulatory motifs than computational motif mapping, as the strategy directly tests whether specific nucleotides are required for transcription in a particular cellular context (**Supplemental Fig S13B**). We therefore took advantage of the fact that functional DNA regulatory regions appear as “peaks” in the deletion effect size map and intersected these peaks with computationally-mapped motifs (**Figure 3A**). Of all of the computationally-mapped motifs in these sequences, 41% and 49% were found to be functional in the tested cell line—i.e., overlapped deletion peak regions—in HepG2 and K562, respectively. For example, the lncRNA DLEU1, which is frequently lost in lymphocytic leukemia (Y. Liu et al. 1997), contains 8 predicted TF motifs, but only one motif—NRF1—significantly overlapped the peak found via single-nucleotide deletions (**Figure 3D**). Therefore, we hypothesize that NRF1, which has a known role in the immune system (Suliman et al. 2010), is the primary and direct regulator of DLEU1. Consistent with this, NRF1 also has a corresponding ChIP peak in the DLEU1 promoter. In total, we were able to determine a wide range of functional TF motifs in 15 lncRNAs, 3 eRNAs, and 3 mRNAs (**Figure 3E; Supplemental Fig S14**). These results show the utility of MPRA in combination with single-nucleotide deletions to refine functional TFs.

Finally, we re-examined the idea that sequences that can be bound by many TFs are more broadly expressed. Indeed, we found that sequences that were active in both cell types had more of our detected functional TF motifs than sequences that were active in only one of the tested cell types (p=0.061, one-sided Wilcoxon test; **Supplemental Fig S15**). This again suggests that the more TFs a sequence can bind, the broader its expression pattern.

### Over 20% of genetic variants within core promoters have regulatory effects

We extended our single-nucleotide MPRA studies to examine how human variation (e.g. single nucleotide polymorphisms (SNPs)) affects promoter activity in contrast to engineered deletions. Briefly, we used MPRA to identify regulatory SNPs that could affect a sequence’s ability to drive transcription in our set of 21 disease-associated lncRNA core promoters. The effect sizes of the tested SNPs were highly correlated with the deletion effect sizes (**Supplemental Fig S16A**). More importantly, significant SNPs tended to occur in peaks corresponding to TF motifs (**Supplemental Fig S16B**). In fact, 78% and 90% of significantly regulatory SNPs that decrease expression overlapped deletion-predicted TF peaks in HepG2 and K562, respectively, compared to only 9% and 5% of non-regulatory SNPs. The tumor suppressor lncRNA *MEG3*, for example, harbors 1 regulatory SNP shown to be mutated in breast cancer tumors by two separate studies (Forbes et al. 2017). This SNP lies in a functional TF peak predicted to harbor binding sites for the CREB family of TFs (**Supplemental Fig S16B**). Together, these results show that our MPRA strategy can identify regulatory SNPs that disrupt functional TF motifs.

To gain a wider understanding of how genetic variation affects DNA regulatory elements, we next used MPRA to test all common SNPs annotated in our set of ~2,000 core promoters in HepG2 and K562 (**Figure 4A**). We correctly identified 100% and 71% of positive control variants as significantly regulatory in HepG2 and K562, respectively (**Supplemental Fig S17**). As with the deletion effect sizes, SNP effect sizes also correlated with the number of predicted TF motifs disrupted by the SNP (**Figure 4B; Supplemental Fig S18**), again suggesting that disruption of multiple overlapping motifs is associated with larger expression changes.

**Figure 4:**
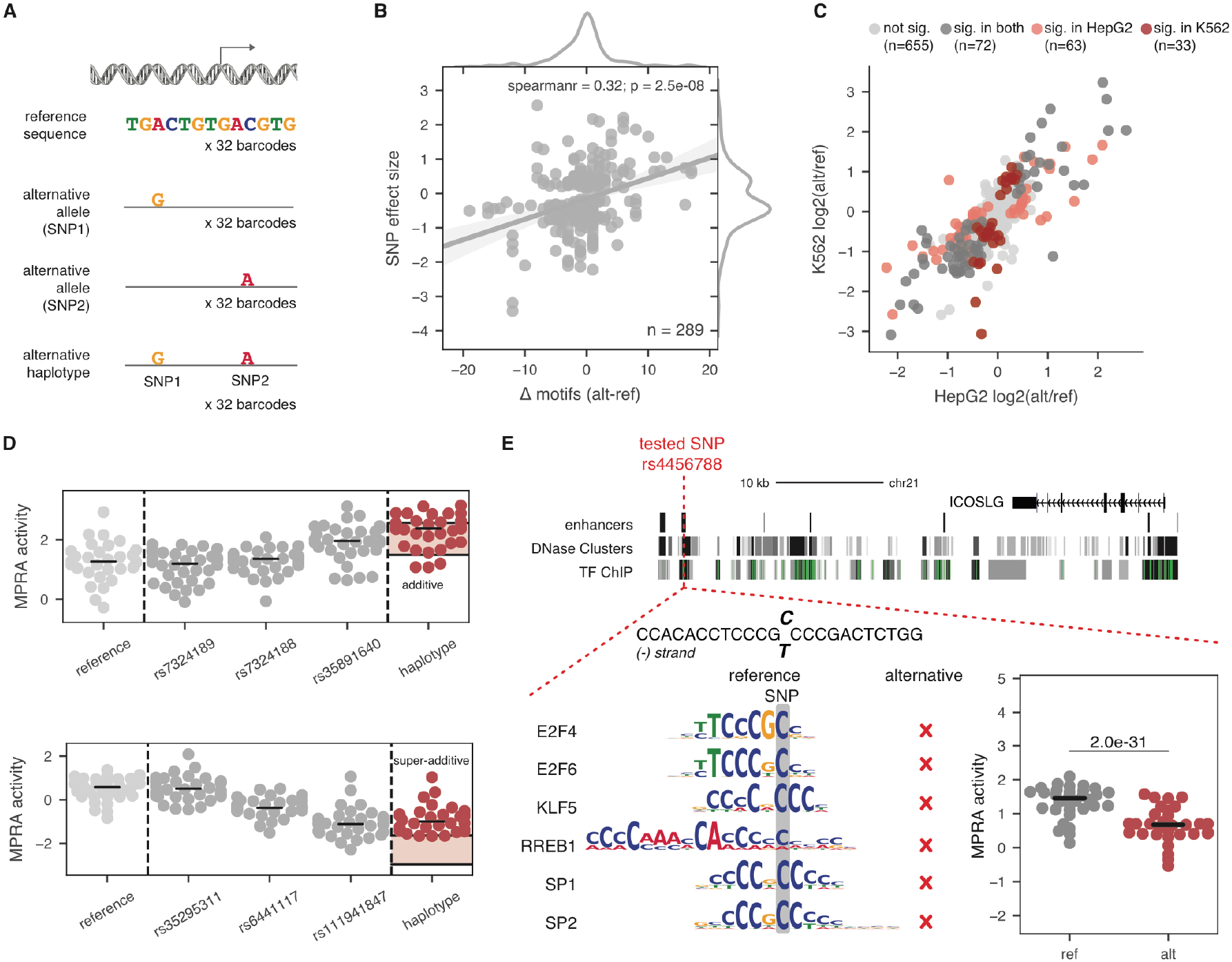
22% of SNPs in promoter and enhancer TSSs have regulatory effects. A. Schematic of SNP and haplotype testing in MPRA. B. Correlation between the number of TF motifs disrupted (x axis) and the SNP effect size (y axis) for all significant SNPs in HepG2. SNP effect size is the mean log2 foldchange in MPRA activity between the alternative and reference alleles. C. Correlation between SNP effect sizes in HepG2 (x axis) and K562 (y axis). D. Examples of two haplotype effects, one additive (top) and one superadditive (bottom). Dots represent barcode activity means across replicates for reference tile (light gray), individual SNP tiles (dark gray), or haplotype tiles (red). Shaded red area in the haplotype column refers to the 90% confidence interval surrounding the expected median additive effect. E. Example of a SNP near *ICOSLG* that disrupts six TF motifs present on the reference allele. The difference in MPRA activity between the reference and alternative alleles in HepG2 is shown. P-value listed is from a two-sided Wilcoxon test.

Overall, we found that as many as 22% of SNPs in the tested TSS regions have an effect on promoter strength (**Supplemental Table S5; Supplemental Fig S19**). We predict that this proportion would increase with a higher number of barcodes (**Supplemental Fig S20A**) and replicates (**Supplemental Fig S20B**). When we looked within each biotype, we found no differences in the number of regulatory SNPs or in the SNP effect sizes (**Supplemental Fig S21**). We found that 55% of regulatory SNPs have an effect in only one of the two cell types (**Figure 4C**).

Due to linkage disequilibrium (LD) in the human genome, multiple individual SNPs tend to be inherited together in haplotypes. However, how individual SNPs interact within a haplotype remains unclear. We therefore sought to determine whether individual SNPs in TSSs tend to interact additively—i.e., the effect of all SNPs together is equal to the sum of their individual effects—or epistatically—i.e., the effect of one SNP masks the effects of the other SNPs. We found that a minority of SNPs acted epistatically as only 16% and 22% of SNPs had a non-additive effect in HepG2 and K562, respectively (**Figure 4D**).

Finally, we sought to identify regulatory SNPs that are in LD with GWAS hits. We identified 96 and 36 such SNPs in HepG2 and K562, respectively (**Supplemental Table S6**). To analyze the putative relationship between the regulatory potential of an MPRA-tested SNP and the GWAS-associated phenotype, we selected SNPs that (i) are regulatory SNPs in both HepG2 and K562 cells (32 total), (ii) disrupt known TF motifs, and (iii) have nearby coding genes that are associated with the GWAS-associated phenotype. We identified three SNPs with significant regulatory effects in both HepG2 and K562 cells that are associated with levels of HDL cholesterol (rs3785098) (Willer et al. 2013), lung cancer (Wang et al. 2008) or schizophrenia (rs3101018) (Goes et al. 2015), and inflammatory bowel disease (IBD) (rs4456788) (J. Z. Liu et al. 2015) respectively (**Figure 4E; Supplemental Fig S22**). For example, the IBD-associated SNP rs4456788 disrupts six TF motifs and shows significantly lower MPRA activity compared to the reference allele (**Figure 4E**). As well as being associated with IBD, this SNP is known to be an eQTL for the protein-coding gene *ICOSLG* (GTEx Consortium 2015) and thus this MPRA result could provide an important clue— and a testable hypothesis—as to the biological pathway that is responsible for this genetic association.

## Discussion

Here, we have characterized the differences between lncRNA, mRNA, and eRNA core promoter sequences by combining computational predictions and experimental testing using high-throughput assays. As many lncRNAs are thought to arise from enhancers (Marques et al. 2013) or bidirectional transcription stemming from protein-coding promoters (Sigova et al. 2013), we sought to determine whether lncRNA promoters are intrinsically different from enhancers and protein-coding promoters. Our findings suggest that the regulation of divergent lncRNAs and intergenic lncRNAs is quite different. Divergent lncRNAs have more TF motifs and consequently have stronger promoters than intergenic lncRNAs. Notably, higher expression levels of divergent lncRNAs compared to lincRNAs cannot solely be explained by having a nearby protein-coding promoter. Rather, we show that both divergent lncRNA and mRNA core promoters are intrinsically stronger than non-divergent lncRNA and mRNA promoters (**Figure 1C**). Conversely, intergenic lncRNA TSSs are similar to enhancer TSSs, both in terms of their TF motif architecture and expression patterns, with both biotypes being highly tissue-specific (**Figure 2D**).

Our results suggest that core promoter sequences play important roles in determining transcript tissue-specificity, as our MPRA results can partially re-capitulate endogenous expression patterns (**Figure 1D, E**). Importantly, using MPRA allows us to thoroughly interrogate the regulatory potential of DNA sequence alone while controlling for other factors such as chromatin differences and effects of post-transcriptional regulation. However, we recognize that this study has some limitations that are intrinsic to using episomal plasmids in MPRA. These limitations are reflected within our own data, in which sequence alone only accounts for ~50% of observed expression profiles when modeled. Nonetheless, our approach has characterized the baseline to which higher order structural and epigenetic information can be added in order to gain a more complete picture of transcriptional regulation.

Our data are consistent with a model where highly abundant genes have complex TF binding profiles, with stretches of promiscuous DNA that can be recognized by many TFs (**Figure 5**). Several lines of evidence point towards overlapping binding sites playing a role in determining abundance and tissue-specificity. First, we see that a model trained on MPRA data finds the number of overlapping motifs to be highly predictive of abundance and anti-correlated to cell-type-specificity (**Figure 2B, C**). Second, we find that tissue-specific biotypes have fewer overlapping motifs than ubiquitously-expressed biotypes (**Figure 2D**). We also find that within one biotype, tissue-specific genes have fewer overlapping motifs than ubiquitously-expressed genes (**Figure 2E**). Third, we show that sequences that are expressed in both HepG2 and K562 have more functional motifs than sequences that are only expressed in one cell type (**Supplemental Fig S15**). Finally, we see that both single-nucleotide deletions and SNPs that are predicted to disrupt more motifs have higher effect sizes (**Figures 3B and 4B**). For example, a single nucleotide deletion in the *HOTAIR* promoter generates 20 new TF motifs and subsequently increases promoter activity by 4-fold (**Figure 3B**).

**Figure 5:**
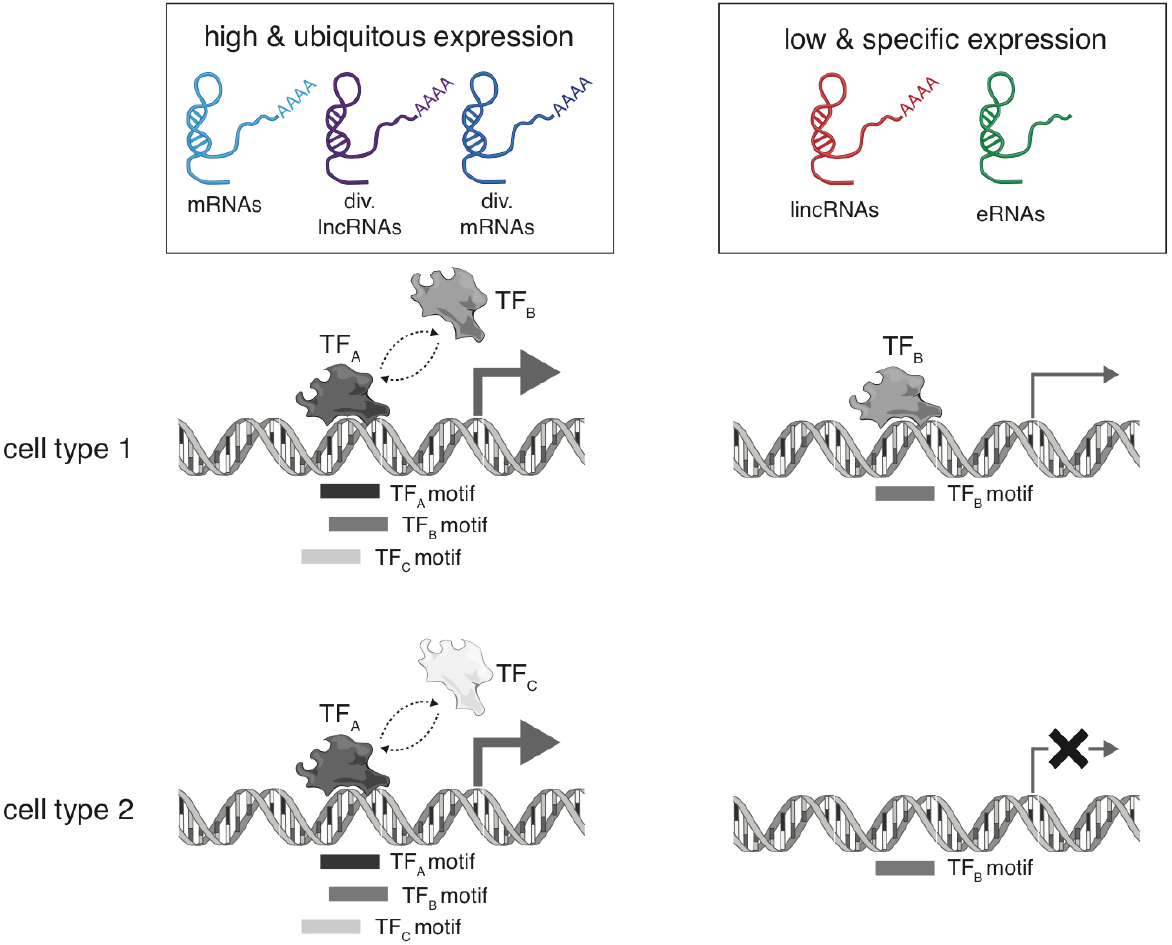
Model: Overlapping TF motifs are associated with high expression and low specificity. Schematic showing biotypes that are highly and ubiquitously expressed (left; big arrow) have more overlapping TF binding sites (gray shaded boxes) and thus more TFs can bind both within a specific cell type or across cell types. Biotypes that are lowly and specifically expressed (right; thin arrow or crossed arrow), have fewer overlapping motifs and thus only a few TFs (one in the example) can bind. TF_A_ Is present in cell type 1 and 2, whereas TF_B_ and TF_C_ are only present in cell type 1 and cell type 2 respectively.

By definition, overlapping binding sites are at the same distance from the TSS; interestingly, where TFs bind in relation to the TSS is important (Tabach et al. 2007). We speculate that this system would allow genes to maintain high and ubiquitous expression levels across different tissues and conditions despite the likely fluctuating expression levels of the TFs. Further, this redundancy could explain why knockdown of certain TFs often does not result in the misregulation of the expected target genes (Cusanovich et al. 2014), as other TFs would be able to bind to the same position. Thus, we propose that promoter specificity may be a function of simplicity in motif usage. Further work remains to be done to experimentally test the extent of the role that these overlapping motif profiles play in regulating expression and specificity.

Our findings also have evolutionary implications. Much attention has been given to the fact that enhancers and lincRNAs have rapid sequence turnover (Hon et al. 2017). Our findings are consistent with this notion. We find that tissue-specific TSSs, such as those of lincRNAs and eRNAs, have less complex motif profiles. Thus, they may be more likely to appear and disappear throughout evolutionary time. In fact, DNA regions with few overlapping motifs are more poorly conserved than DNA regions with many overlapping motifs (**Figure 2F**). Conversely, highly transcribed genes have developed more complex TF binding patterns, which may have evolved to produce stable antisense transcripts because they provide a fitness advantage. Indeed, if we compare human and mouse orthologous genes that have gained a stable antisense transcript in either one of the lineages, they show an overall increase in expression (**Supplemental Fig S23**). Thus, bidirectional transcription may not only allow for *de novo* gene origination but could also be an evolutionary mechanism to increase expression of the gene in the sense direction. This may also help explain why divergent mRNA-lncRNA pairs occur so frequently in the human genome (Sigova et al. 2013). Overall, this study sheds light on the important roles that core promoters play in complicated aspects of gene regulation—including divergent transcription and tissue-specificity—across both coding and non-coding genes.

## Methods

### TSS biotype classification

mRNA and lncRNA TSSs were classified based on Gencode v19 (Harrow et al. 2012) gene annotations. All TSSs from genes annotated as lncRNAs were classified as intergenic lncRNA (lincRNA) TSSs if they did not overlap any annotated protein-coding genes or as divergent lncRNA TSSs if the annotated TSS had an antisense FANTOM5 TSS within 1000bp. Similarly, mRNA TSSs were classified as divergent mRNA TSSs if the annotated TSS had an antisense FANTOM5 TSS within 1000bp. eRNA TSSs were also defined by FANTOM5 (Andersson et al. 2014) and had two TSSs each—a sense TSS and an antisense TSS—due to their inherent definition of being bidirectionally transcribed.

### MPRA TSS selection

To select promoters to include in the MPRA, we used the FANTOM5 robust TSS set (Forrest et al. 2014). These TSSs are expressed robustly in CAGE-seq data (> 10 CAGE reads in one sample and > 10 tpm CAGE expression in at least one sample). Additionally, we only considered FANTOM5 TSSs that were within 50bp of their cognate annotated Gencode v19 transcript TSSs. eRNA TSSs were selected from the enhancer TSS set defined in the same FANTOM5 release (Andersson et al. 2014). Next, we selected TSSs based on their CAGE expression profiles. Specifically, we required the TSSs to either (a) be expressed > 0.5 tpm across all replicates of at least one of the tested cell lines (HeLa, HepG2, or K562) or (b) have an average expression > 0.5 tpm across all FANTOM5 samples (suggesting they had high baseline expression). Finally, we excluded any lncRNA TSSs arising from transcripts with high coding potential (phyloCSF > 0 ((Lin, Jungreis, and Kellis 2011))) or that overlapped a protein-coding gene in the sense direction.

As the MPRA was lncRNA-focused, all lncRNA TSSs (lincRNAs and divergent lncRNA TSS) that filled the criteria above were included for testing in the MPRA. To control for the fact that there were more mRNA and eRNA TSSs than lncRNAs, we selected both expression-matched mRNA and eRNA TSSs (average expression across all FANTOM5 samples matching that of lncRNA TSSs) as well as randomly-selected mRNA and eRNA TSSs for further analysis. We also included all protein-coding TSSs that were in close proximity to the selected lncRNA TSSs (<160bp) in antisense and some of the most highly expressed eRNAs. Additional TSSs were included if they contained at least one SNP in LD with a GWAS hit in their core promoters (see **Supplemental Table S6** for additional details). Overall, we ended up with 2,078 TSSs for testing in MPRA (**Figure 1A; Supplemental Table S1**). More details available in Supplemental Methods (MPRA TSS selection section).

### MPRA pool design

Two 120,000 oligonucleotide (oligo) pools of 170 bp with 11 bp barcodes were designed. The first pool included core promoter sequences across biotypes and common SNPs falling in these regions (see supplemental methods; **Supplemental Table S1; S2**). The second pool included singlenucleotide deletions across the core promoters of 21 lncRNAs, 5 enhancers and 5 mRNAs with two consecutive reference tiles each (see supplemental methods; **Supplemental Table S3; S4**). Random and scrambled sequences were included in both pools as negative controls. More details available in Supplemental Methods (MPRA oligo pool design section).

### MPRA cloning and transfection

Oligo pools were synthesized by Twist Biosciences and then cloned into plasmids to generate a library of constructs where the regulatory sequence is upstream of a reporter gene (here, GFP) that is upstream of a unique barcode (see supplementary methods). Constructs were transfected into live cells and barcode expression was assayed by high-throughput RNA sequencing. A minimum of 4 and a maximum of 12 replicates were performed per condition (cell type and presence/absence of a minimal promoter) adding up to 32 total experiments (**Supplemental Fig S3**). Results are based on the minimal promoter set-up given the high similarity between replicates with and without the minimal promoter and the fact that more replicates were performed for this set-up. More details available in Supplemental Methods (MPRA cloning, transfection, and sequencing section).

### MPRA data analysis

All Python scripts and notebooks used to perform the MPRA analyses are available at https://github.com/kmattioli/2018_lncRNA_promoter_MPRA and provided as Supplemental Materials.

Exact matches to known barcodes and 6 upstream constant nucleotides were mapped after quality-filtering the sequencing reads. Barcodes were filtered to those with >=5 counts (in both DNA and RNA). Barcode activities were calculated as the log-transformed proportion of RNA barcodes to the proportion of DNA barcodes (after normalizing for sequencing depth) and were quantile-normalized across replicates. Element activities were calculated as the median activity value across all cognate barcodes, requiring >=3 barcodes. Significantly active tiles were defined as those with barcode activities that were significantly more activity than random negative control sequences according to a two-sided Wilcoxon test in >=75% of replicates (see supplemental methods). As we had many more replicates in HepG2 than in other cell types and to ensure we had similar power when comparing across cell types (i.e., **Figure 1F**), HepG2 replicates were down-sampled 100 times and sequences were considered significant if they were significant by the rules above in >=75% of samples. More details available in Supplemental Methods (MPRA analysis section).

### Core promoter element analysis

The core promoter was defined as 80 bp upstream to 34 bp downstream of the TSS. CpG content was calculated by counting the number of “CG” dinucleotides in this region. Inr motifs were defined to be matches to the motif BBCABW (B=C/G/T, W=A/T) (Kugel and Goodrich 2017) within 5 bp of the TSS. TATA motifs were defined to be matches to the motifs TATAAA or TATATA within 55 to 15 bp upstream of the TSS. Position weight matrices for TF binding motifs were obtained from the JASPAR database (core, vertebrates, 2016 release) (Mathelier et al. 2014).

### MPRA activity and tissue-specificity predictions

An ANOVA analysis was used to evaluate what properties contribute MPRA activity and specificity. Specificity across the MPRA activity values for HepG2, K562, and HeLa was calculated using the *tau* metric (Kryuchkova-Mostacci and Robinson-Rechavi 2017):

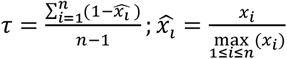

where *x_i_* is the median activity of a TSS in cell type *i* and *n* is the number of cell types. Briefly, *tau* calculates the average difference between the activity of a TSS in a given cell type and the TSS’ maximal expression across all cell types. Thus, “ubiquitous” TSSs will have *tau* values close to zero while “tissue-specific” TSSs will have *tau* values close to one.

To perform the ANOVA analysis, the variance in activity/specificity that is explained by the general sequence features (listed in **Supplemental Fig S6A**) was calculated. The variance explained by each parameter was calculated on its own and the optimal subset of parameters was computed. As the parameters were highly correlated, the optimal subset consisted of only 7 out of 14 parameters yet explained 41% of the total variance (**Supplemental Fig S6A**).

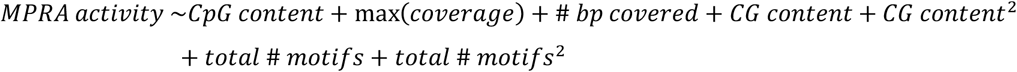

Motifs were then added into the model one by one.

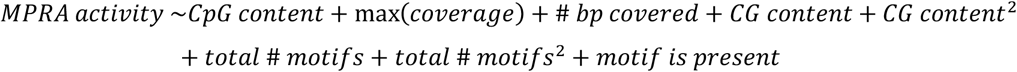

Of the 382 motifs tested, 17 were found to explain a significant fraction of the variance (listed in **Supplemental Fig S6B**). Combining the 7 sequence features and the significant motifs in a model explained a total of 49% of the variance in MPRA activity.

This analysis was performed in R (version 3) using the leaps and tidyverse packages.

### ChIP-seq analysis and TF motif mapping

ChIP-seq files were downloaded from the Cistrome Data Browser (cistrome.org) (Mei et al. 2017) for 771 human TFs (**Supplemental Table S7**)—218 of which overlapped with the set of 519 JASPAR motifs. BedTools (Quinlan and Hall 2010) was used to merge peaks for a given TF and then intersect the merged ChIP peaks with our set of promoters. Since Cistrome peaks were in hg38 and our promoters were in hg19, we first used liftOver (Hinrichs et al. 2006) to convert our promoters to hg38 coordinates. Motifs were mapped in sequences using FIMO (version 4.11.2) (Grant, Bailey, and Noble 2011) with a p-value threshold of 1e-5. Motifs were assigned to ChIP-seq peaks if there was a FIMO motif mapped within 250 bp of the ChIP-seq peak.

### MPRA deletion analysis and functional TF motif mapping

Deletion effect sizes were defined as the log2 foldchange between the mean activity of the deletion sequence across replicates and the mean activity of the reference sequence across replicates, resulting in a value per nucleotide. Peaks were defined as any stretch of >= 5 nucleotides with effect sizes of <= −1.5 * the average standard deviation of the deletion effect sizes in that tile. Mapped motifs were said to be “functional” if >= 1 nucleotide in the motif intersected a peak.

### SNP and haplotype analysis

Regulatory SNPs were defined as those whose barcode activities were significantly different and consistent in direction between reference and alternative tiles using a two-sided Wilcoxon test in >=75% of replicates (see supplemental methods). Again, when comparing between cell types (i.e. **Figure 4C**), HepG2 replicates were down-sampled 100 times as above.

To determine additive haplotypes, first the expected additive haplotype effect size was found by summing the median log2 foldchanges (alternative/reference activities) for each individual SNP in a haplotype. This effect was bootstrapped (n=1000) to determine a 90% confidence interval and a haplotype was considered additive if the actual median log2 foldchange of the haplotype fell within this 90% confidence interval.

The GWAS catalog was downloaded from http://www.ebi.ac.uk/gwas/docs/downloads. Raggr was used to calculate whether any of the MPRA-tested SNPs were in linkage disequilibrium (r^2^<0.6) with any of the GWAS tag SNPs at http://raggr.usc.edu/.

### Data Access

The MPRA sequencing data from this study have been submitted to the NCBI Gene Expression Omnibus (GEO; http://www.ncbi.nlm.nih.gov/geo/) under accession number GSE117594. All scripts required to reproduce this work are available as Supplemental Material as well as on GitHub at https://github.com/kmattioli/2018_lncRNA_promoter_MPRA.

## Supporting information

## Acknowledgements

We thank Catherine Weiner, Abigail Groff, and Julia Rogers for thoughtful comments on the manuscript. We thank Lucas Janson and Kian Hong Kock for helpful discussions throughout the project. M.M. is a Gilead Fellow of the Life Sciences Research Foundation. K.M. is a National Science Foundation Graduate Research Fellow under Grant No. DGE1144152. J.C.L. holds a Wellcome Trust Intermediate Clinical Fellowships (105920/Z/14/Z). J.L.R. is an HHMI faculty scholar. This work was supported by US National Institutes of Health grant P01 GM099117.

## Author Contributions

K.M., M.M., and J.R. designed the project and wrote the manuscript. K.M. and M.M. designed the oligonucleotide libraries. K.M. and M.M. performed all MPRA analyses. P.J.V. performed genome-wide ChIP-seq analysis and the MPRA model. C.G. and P.M. performed the MPRA experiments. J.C.L. contributed to the project design. All authors have read and approved the manuscript for publication.

