## Supplementary material for "High-throughput functional analysis of IncRNA core promoters elucidatesrules governing tissue-specificity"

### Supplemental materials for “High-throughput functional analysis of lncRNA core promoters elucidates rules governing tissue specificity”

Kaia Mattioli, Pieter-Jan Volders, Chiara Gerhardinger, James C. Lee, Philipp G. Maass, Marta Melé*, John L. Rinn*

* contributed equally

#### Supplemental Methods

##### *Genome-wide analyses*

**Selection of TSSs for genome-wide analyses.** All 'robust' FANTOM CAT TSSs (defined as CAGE-seq counts >10 in >=1 sample and CAGE-seq tpm >1 in >=1 sample ) assigned to Gencode v19 genes were selected (Hon et al. 2017) along with all FANTOM5 enhancers meeting the 'robust' criteria (Andersson et al. 2014). When a Gencode gene had more than one TSS assigned by FANTOM, the TSS closest to the annotated Gencode TSS was selected. All 'robust' enhancer TSSs defined by FANTOM5 were selected except those overlapping protein-coding gene loci (Forrest et al. 2014). FANTOM5 CAGE normalized expression values were obtained via the FANTOM portal (Lizio et al. 2015) for both promoters and enhancers.

**CAGE expression and specificity calculations.** FANTOM5 samples corresponding to time courses or fractionated cells were omitted and then grouped by tissue or cell type (**Supplemental Table S8**). Average expression was calculated across these grouped samples. Tissue-specificity was calculated across these grouped samples using the *tau* metric (Kryuchkova-Mostacci and Robinson-Rechavi 2017):

$$\tau = \frac{\sum_{i=1}^{n}(1-\widehat{x_i})}{n-1}; \widehat{x_i} = \frac{x_i}{\max\limits_{1\le i\le n}(x_i)}$$

where $x_i$ is the median expression of a TSS in tissue *i* and *n* is the number of tissues. Briefly, *tau* calculates the average difference between the expression of a TSS in a given sample and the maximal expression of that TSS across all samples. Thus, "ubiquitous" TSSs will have *tau* values close to zero while "tissue-specific" TSSs will have *tau* values close to one. We then defined three TSS expression profiles: (1) "ubiquitous": TSSs expressed (CAGE tpm > 0) in > 90% of all grouped CAGE samples), (2) "dynamic": TSSs expressed (CAGE tpm > 0) in < 10% of all grouped CAGE samples, with very high expression (CAGE tpm > 50) in at least 1 sample, and (3) "tissue-specific": TSSs expressed (CAGE tpm > 0) in < 10% of all grouped CAGE samples, with no samples expressed at CAGE tpm > 50. All other TSSs were considered "moderate".

##### *MPRA TSS/gene selection and oligonucleotide pool design*

We designed two 120,000 oligonucleotide (oligo) pools of 170 bp. All oligos contained universal primers, two restriction enzyme sites, an 11 bp barcode, and the regulatory sequences of interest as follows (Melnikov et al. 2012):

**5'**---**universal primer 1** (16 bp)---**variable sequence** (144 bp)---**XbaI** (6bp)---**KpnI** (6bp)---**barcode** (11 bp)---**universal primer 2** (17 bp)---**3'**

**Universal primer 1:** ACTGGCCGCTTCACTG
**Universal primer 2:** AGATCGGAAGAGCGTCG

In the first pool, reference sequences correspond to core promoter sequences (-80 to +34 bp from the TSS) from different biotypes (see below; **Table S2**). In the second pool, we selected 21 lncRNAs, 5 enhancers and 5 mRNAs to perform single nucleotide deletions within their TSSs (see below; **Table S4**).

###### Pool 1

**Selection of TSSs.** We used TSSs (mRNAs, lncRNAs and enhancers) from the FANTOM5 consortium (http://fantom.gsc.riken.jp) (Forrest et al. 2014). For mRNA and lncRNA TSSs, we only considered TSSs that are within 50bp of an annotated Gencode v19 transcript TSS. For enhancers, which have two TSSs (one on the minus and one on the plus strand), we selected the TSSs that are either expressed at higher than 0.5 TPMs across all three replicates of at least one of the tested cell lines (K562, HeLa, HepG2) or have an average expression of > 0.5 TPM across all FANTOM5 samples (suggesting that they will have a high baseline expression). If there was more than 1 TSS assigned to a single lncRNA or mRNA transcript, we selected the one with the highest expression. All TSSs that belong to lncRNA transcripts with phyloCSF scores < 0 (Lin, Jungreis, and Kellis 2011) and do not overlap a protein-coding gene in the sense direction were selected for further analysis. We also included 14 lncRNAs TSSs that have a GWAS hit in LD with a SNP in their core promoters. We then included expression-matched and randomly selected mRNAs (180 each) and eRNA TSSs (100 enhancers; 200 eTSSs each). We also added all mRNAs ($r2>0.70$) and all eRNAs ($r2>0.6$) that have at least one SNP in LD with a GWAS hit. Finally, we included all protein-coding TSSs that share their promoter with the selected lncRNAs (<160bp). Additionally, we selected some of the 75 most expressed eRNAs. We selected all SNPs present in 1000 Genomes (Gibbs et al. 2015) at a frequency > 0.01 that fell in our core promoter regions.

**Oligo pool design.** In this pool, all test sequences correspond to core promoter sequences (-80 to +34 bp from the TSS) from different biotypes selected as explained above (**Table S1**). For 25% of the sequences, we also included their reverse complements. Specifically, we did it for 1) the 100 most expressed lncRNA TSSs and 2) their expression-matched most highly expressed eRNA (50 eRNAs, 100 TSSs) and mRNA TSSs (100 mRNAs) and 3) all divergent promoters and enhancers for which their antisense TSS was closer than 160bp and further than 70bp. We selected all SNPs as described above, and when there was more than one SNP present in a core promoter, we tested each SNP individually as well as the haplotype with all the alternative variants together (**Figure 4A**). We assigned 15 barcodes for tiles with no SNPs and 32 barcodes for tiles with SNPs (**Table S2**). As positive controls, we used the main HBB and the A100S1 promoters (in both sense and antisense directions) from a previous publication with a similar set-up (Patwardhan et al. 2009). We also added the substitutions with the strongest effects to use as positive controls for the genetic variants (2 depleting substitutions for each promoter and also one activating substitution for HBB). For enhancers, we used the AldoB enhancer (Patwardhan et al. 2012) as a positive control. We added the three substitutions with the strongest effects (two depleting one activating). We assigned 80 barcodes to control tiles. We also included 100 scrambled sequences (chosen randomly and each permuted 10 times) and 1,000 random sequences to serve as negative controls.

<u>**Pool 2**</u>

**Selection of TSSs.** We selected 21 TSSs from lncRNAs using the following criteria: (1) high average expression and high expression across at least one of the cell lines of interest (HeLa, K562, HepG2), (2) known function (Quek et al. 2015), and (3) related to specific diseases (mostly cancer) (Huarte 2015). We also included 5 protein-coding promoters that are divergent with 5 of the lncRNAs as well as 5 nearby enhancers (**Table S4**).

**Oligo pool design.** We designed two consecutive tiles to cover the TSS region (tile 1: -183 to -69 bp from the TSS, tile 2: -89 to +25 bp from the TSS). For 4 of the TSSs (8 tiles), we included their reverse complements. We performed single-nucleotide deletions in the inner 94 bp of each tile (so, excluding the flanking 10 bp on each side). We also selected all SNPs present in 1000 Genomes (including low frequency variants) that fall in our tiles, filtering out insertions and deletions. We used 80 barcodes for the reference tiles and 26 barcodes per deletion tile or alternative SNP tiles (**Table S5**). For each reference sequence, we included 40 scrambled permutations to serve as negative controls as well as 2,000 random sequences.

#### *MPRA cloning, transfection, and sequencing*

**ePCR amplification of oligo pools.** The synthesized oligo pool was amplified by emulsion-PCR (ePCR, Micellula DNA Emulsion & Purification Kit, Chimerx), according to the manufacturers' instructions. ePCR primers harboring Sfi I restriction sites were designed to add for subsequent cloning purposes (5' primer: GCTAAGGGCCTAACTGGCCGCTTCACTG; 3' primer: GTTTAAGGCCTCCGAGGCCGACGCTCTTC). To determine the oligos representation of the ePCR-amplified oligopool (based on the unique 3' barcode of each oligo), 1 ng of the amplified oligo pool was used as input for library preparation (see below) and sequenced on a MiSeq (SR, Illumina).

**Cloning.** The ePCR-amplified oligopools were digested with SfiI, and inserted in the multiple cloning region of an empty vector. In the first cloning step, the ligation reaction (100 ng backbone + 4x molar excess of oligopool) was transformed into 20 x DH5$\alpha$ tubes (ThermoScientific), spread out on 50 ampicillin LB plates, and incubated overnight at 37°C. After scraping all bacteria of each LB plate in 5 ml LB and pooling all colonies, plasmids were purified with the endotoxin-free Qiagen Plasmid Plus Maxi kit (Qiagen). The oligo representation was determined by MiSeq-sequencing. Next, the cloned oligopool was sequentially cut with Kpn I and Xba I and ligated as described above with and without a minimal promoter (5'-AGAGGGTATATAATGGAAGCTCGACTTCCAG-3') and an ORF for GFP. Illumina sequencing determined barcode representation of the oligopools. Finally, to remove plasmids without inserted oligos, the oligopools were digested with Kpn I and the plasmid fractions containing oligos were size-selected by agarose gel-electrophoresis, re-ligated and cloned as described above.

**Library preparation.** Cloned oligopools (50 ng) were amplified with PfU HS DNA polymerase (Agilent) and 5 $\mu$l of 2 $\mu$M index primer (cloning step 1, universal 3' primer: AATGATACGGCGACCACCGAGATCTACACTCTTTCCCTACACGACGCTCTTCCGATCT; Index 1 5' primer: caagcagaagacggcatacgagat<u>CGTGAT</u>gtgactggagttcagacgtgtgctcttccgatctACTGGCCGCTTCACTG; cloning step 2 & 3 and cDNA libraries, universal 3' primer: AATGATACGGCGACCACCGAGATCTACACTCTTTCCCTACACGACGCTCTTCCGATCT; Index 1 5' primer: caagcagaagacggcatacgagat<u>CGTGAT</u>gtgactggagttcagacgtgtgctcttccgatctCGCCGCGTGGAGGAGGA, underlined nucleotides = index; PCR setting: 95°C 2 min, 95°C 30 sec, 55°C 30 sec, 72°C 15 sec [x 18-24 cycles], 72°C 10 min, 4°C hold). After 18 cycles of amplification, molarity was checked on a BioAnalyzer. Insufficient amplifications were run for 2-6 additional cycles (total of 24 cycles). Amplified libraries were then cleaned three times with AMPure beads with the following ratios according to manufacturer's instruction: 0.6x, 1.6x, 1.0x.

**Transient transfections.** K562 (chronic myelogenous leukemia lymphoblasts), HepG2 (liver carcinoma epithelial cells) and HeLa (cervical carcinoma epithelial cells) cells were ordered from ATCC and PCR-tested for mycoplasma contamination (LookOut, SigmaAldrich). Media were prepared according to ATCC recommendations and tissue culture conditions were standard (5 % $CO_2$, 37°C). K562 cells were transiently transfected for 48 hours with a ratio of 1:1 with Xtreme Gene HP (Roche), HeLa and HepG2 both with 3:1 Xtreme Gene HP according to the manufacturer protocol. 4x10e6 K562 cells in 6 cm dish, 1.6x10e6 HeLa or HepG2 cells in 10 cm dish were seeded 12-16 hours prior to the transfection. Cells were counted before the transfection and per 1 x 10e6 cells, 5 $\mu$g oligopool were transfected. Media was not changed and transfection efficiency was microscopically determined by GFP expression 48 hours post transfection.

**RNA extraction and cDNA library preparation.** RNA from transiently transfected K562, HeLa, and HepG2 was precipitated by phenol-chloroform extraction according to standard protocols. DNase treatment (Worthington) was followed by cDNA synthesis with SuperScript III First-Strand Synthesis System (Invitrogen). cDNA was subject to library amplification as listed above (primer: cloning step 2 & 3 and cDNA libraries), and libraries were cleaned up with AMPure beads (0.6x, 1.6x, 1.0x).

**Sequencing & quality control.** We sequenced RNA illumina polyA+ single end either 50 bp or 100 bp. We used cutadapt (Martin 2011) to remove adapters and trim bases with a Phred score lower than 20. We required each filtered read to exactly match one of our pre-designed 11-nucleotide barcodes as well as the neighboring upstream 10 constant nucleotides (TCTAGAATTA).

#### *MPRA analysis*

All scripts used to do the MPRA analysis are available at https://github.com/kmattioli/2018__lncRNA_promoter_MPRA.

**Normalization and activity calculation.** We only used barcodes that had at least 5 reads present (in both DNA and RNA). We then normalized the read counts for sequencing depth within each replicate. To calculate activity per barcode, we calculated the proportion of RNA barcodes to the proportion of DNA barcodes, log-transformed the proportion, and quantile normalized the activities across replicates. To get an activity level per element (i.e., each unique sequence, all of which had multiple barcodes associated with them), we selected the median activity value across all of its cognate barcodes, requiring a minimum of 3 barcodes to be present.

**Defining significantly active tiles.** To define significantly active tiles, we compared the barcode activities of a sequence of interest to barcode activities corresponding to the random negative control sequences using a two-sided Wilcoxon test. We did this per replicate, and then combined the resulting *p*-values across replicates using Stouffer's method (Stouffer et al. 1949). We corrected the *p*-values for multiple testing using the Bonferroni method. We determined a sequence to be significantly active if its corrected combined *p*-value was < 0.05 and if the mean log2 foldchange between reference and negative controls was >0.5 in >75% of replicates. Since we performed far more replicates in HepG2 than in either K562 or HeLa, to ensure we had similar power when comparing across cell types (i.e., **Figure 1F**), we down-sampled the HepG2 replicates 100 times, and considered the sequences as significant if they were significant by the rules above in >75% of samples.

**Deletion analysis.** To determine the effect size of deletions, we found the log2 foldchange between the mean activity of the deletion sequence across replicates (median across all barcodes) and the mean activity of the reference sequence across replicates (median across all barcodes), resulting in a value per nucleotide. To find significant deletions, we compared these two barcode distributions (mean across replicates) using a two-sided Wilcoxon test, and then combined and corrected the *p*-values as mentioned above. We determined deletions to be significant if the combined corrected *p*-value was < 0.05 and if the mean log2 foldchange between deletion and reference tiles was in the same direction in > 75% of replicates.

**SNP analysis.** We first defined significant SNPs within each replicate by comparing the barcode activities of the reference allele to the barcode activities of the alternative allele using a two-sided Wilcoxon test. We then combined and corrected the *p*-values as above. We determined a SNP to be significant if its corrected *p*-value was < 0.05, and if the direction of the effect between the reference and alternative allele was consistent in > 75% of replicates. Again, when comparing between cell types (i.e. **Figure 4C**), we down-sampled HepG2 replicates 100 times as above.

**Haplotype analysis.** To determine whether SNPs were interacting additively, we first found the expected haplotype effect size by summing the median log2 foldchanges (alternative/reference activities) for each individual SNP in a haplotype. Then, we bootstrapped this effect to determine a 90% confidence interval by re-sampling both the reference values and each of the alternative values 1000 times and repeating the process. We considered a haplotype as additive if the actual median log2 foldchange of the haplotype (all alternative alleles/reference activities) fell within this 90% confidence interval. If the actual median foldchange fell below the interval, we considered it “sub-additive”, and if it fell above the interval, we considered it “super-additive”.

#### Supplemental Tables

More detailed column descriptions provided in each Supplemental Table file.

**Supplemental Table S1: Properties associated with TSSs included in TSS MPRA.** Table is a txt file.

**Supplemental Table S2: Barcode and biotype information included in TSS MPRA.** All reference sequences correspond to core promoter sequences, defined as -80 to +34 bp from the TSS. Only common SNPs (frequency > 0.01) and their corresponding haplotypes are included in this pool. *Note that the barcodes for scrambled tiles actually correspond to the number of permutations (10), but each permutation itself only has 1 barcode.

| sequence type | # unique sequences | # barcodes | total # oligos |
|---|---|---|---|
| Reference (sense) | 1229 | 15 | 18,435 |
| Reference (antisense) | 544 | 15 | 8,160 |
| Reference that has a corresponding SNP | 993 | 32 | 31,776 |
| SNP (individual) | 1404 | 32 | 44,928 |
| SNP (haplotype) | 361 | 32 | 11,552 |
| Positive control | 32 | 80 | 2,560 |
| Random | 1589 | 1 | 1,589 |
| Scrambled | 100 | 10* | 1,000 |
| | | | ***120,000*** |

**Supplemental Table S3: Properties associated with genes included in deletion MPRA.** Table is in an excel file.

**Supplemental Table S4: Barcode and biotype information included in deletion MPRA.** Note that here, reference sequences include both core promoter tiles, defined as -89 to +25 bp from the TSS, and an upstream tile, defined as -183 to -69 bp from the TSS. Both rare and common SNPs are included in this pool. Note that the barcodes for scrambled tiles actually correspond to the number of permutations (40), but each permutation itself only has 1 barcode.

| sequence type | # unique sequences | # barcodes | total # oligos |
|---|---|---|---|
| Reference (sense) | 54 | 80 | 4320 |
| Reference (antisense) | 8 | 80 | 640 |
| Deletion | 3964 | 26 | 103,064 |
| SNP (individual) | 173 | 26 | 4498 |
| Positive control | 30 | 80 | 2400 |
| Random | 2000 | 1 | 2000 |
| Scrambled | 54 | 40* | 2160 |
| | | | ***119,082*** |

**Supplemental Table S5: SNP results.** Table is a txt file.

**Supplemental Table S6: GWAS-associated SNP information.** Table is a txt file.

**Supplemental Table S7: ChIP datasets used in analyses.** Table is a txt file.

**Supplemental Table S8: FANTOM5 CAGE sample groupings used in analyses.** Table is a txt file.

#### Supplemental Figures

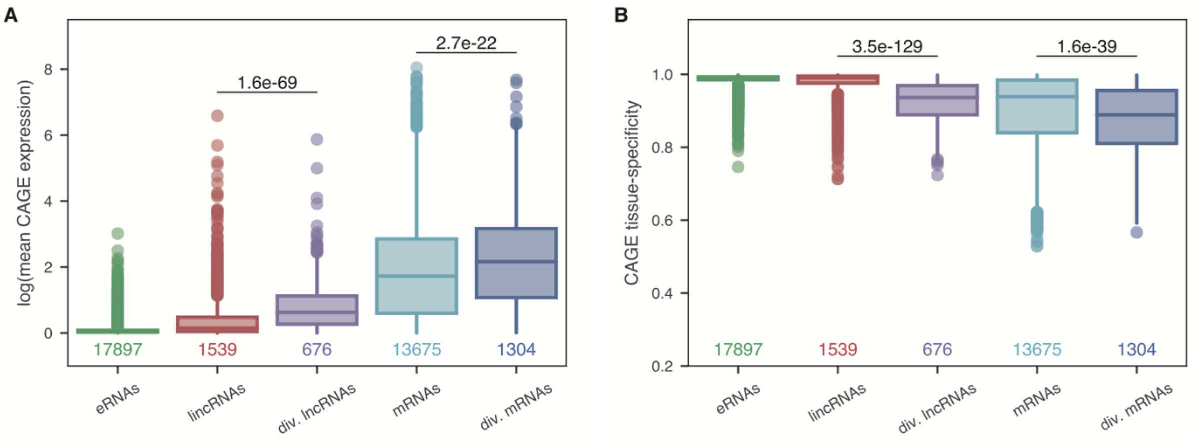

**Supplemental Fig S1: CAGE expression and tissue specificity across biotypes. A.** Mean CAGE expression per biotype plotted in log space. **B.** CAGE tissue specificities per biotype, where 0 is ubiquitous and 1 is tissue-specific. Only "robust" TSSs as defined by FANTOM5 are included (>= 10 CAGE reads in at least 1 sample and >= 10 tpm CAGE expression in at least 1 sample), with *n* TSSs plotted below each distribution. *P*-values are from a two-sided Wilcoxon test.

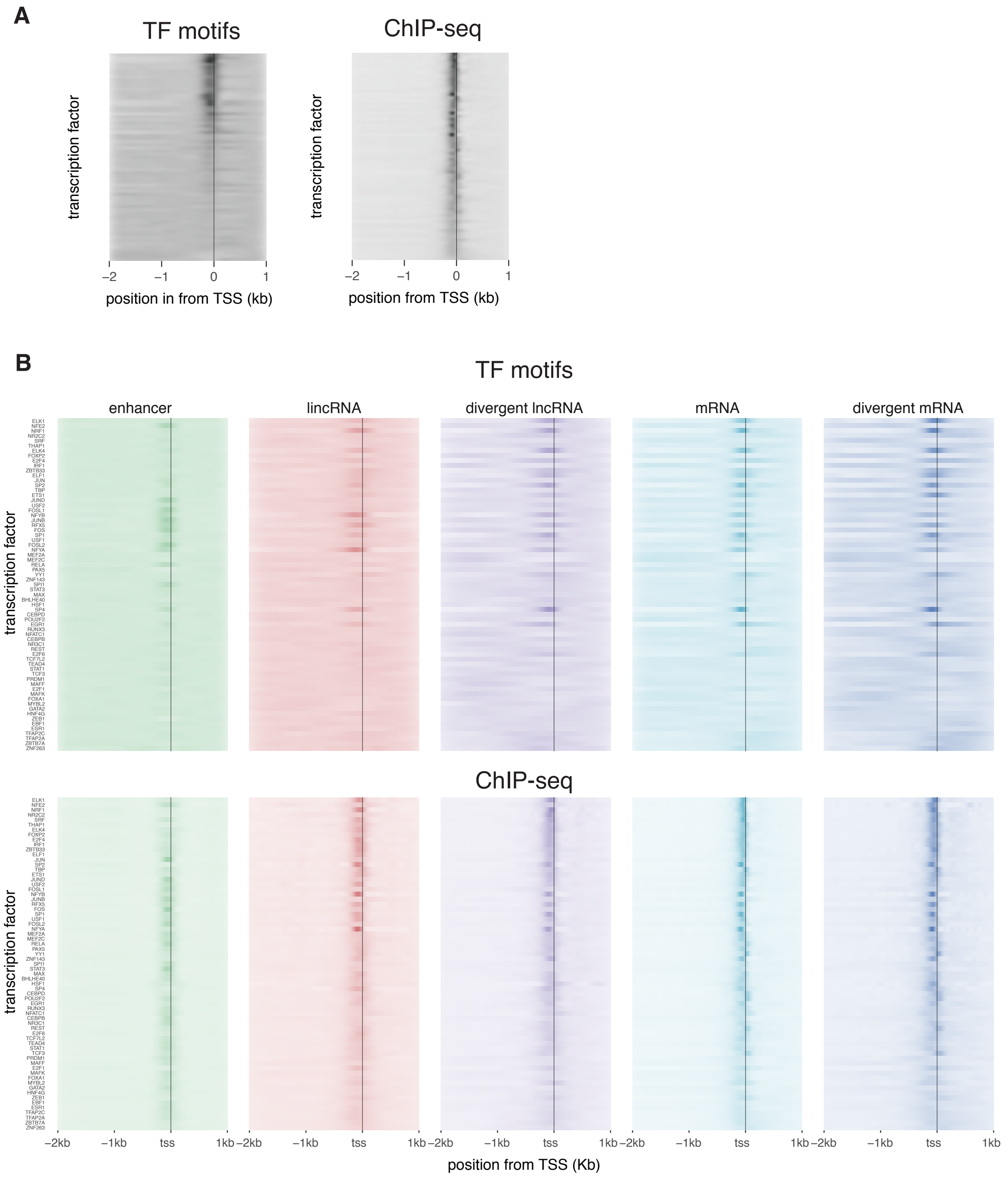

**Supplemental Fig S2: TF motifs and ChIP sequencing peaks are enriched in a narrow region around the TSS. A.** Positioning of TF motifs and ChIP peaks for 67 transcription factors across ~76,000 TSSs. TFs (y axis) are sorted based on the relative enrichment in the core promoter, where TFs that are higher on the y-axis are more enriched. B. TF motifs and ChIP peaks for 67 transcription factors across ~76,000 TSSs divided by biotype.

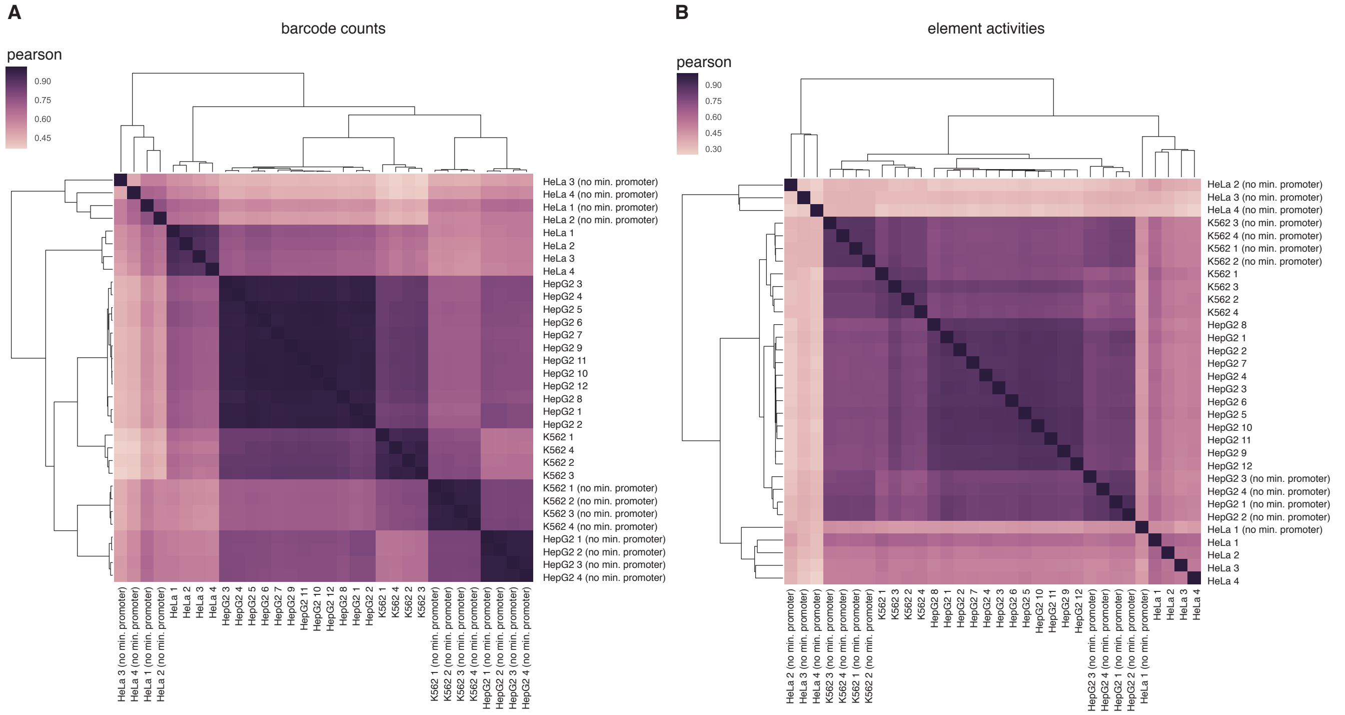

**Supplemental Fig S3: Correlation across replicates**. Clustering was performed for all replicates, including those with and without a minimal promoter. Hierarchical clustering is based on Euclidean distance with average linkage. **A.** Pearson correlation of counts *per barcode* across replicates. We only included barcodes with >5 counts in at least 1 replicate. **B.** Pearson correlation of activity values *per element* across replicates. We only included elements with >=3 barcodes represented at with counts >5 in at least 1 replicate. Activity values per element were calculated by collapsing all barcodes that map to the same unique tested sequence or element and dividing RNA by DNA normalized counts (see methods). Negative controls were excluded from the plot since each only had 1 barcode by design. In both cases, replicates cluster per cell type whereas presence or absence of a minimal CMV promoter plays a secondary role.

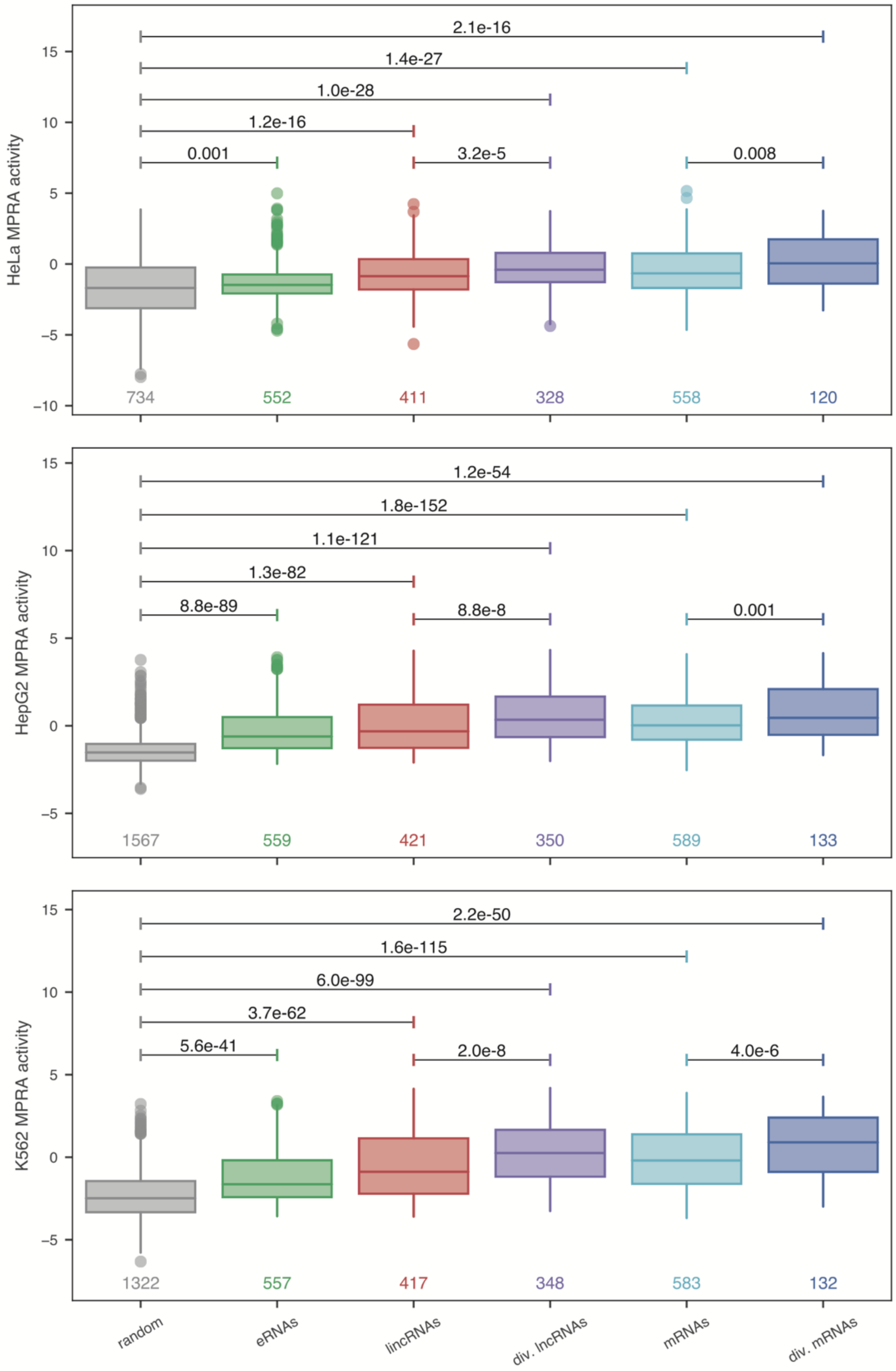

**Supplemental Fig S4: Test sequences are significantly more active than negative controls and vary across TSS classes.** Activity of core promoter sequences across TSS classes compared to random sequences in HeLa, HepG2, and K562. Activity is plotted as the mean across replicates of the median barcode activity values per element. *P*-values listed are from a two-sided Wilcoxon test.

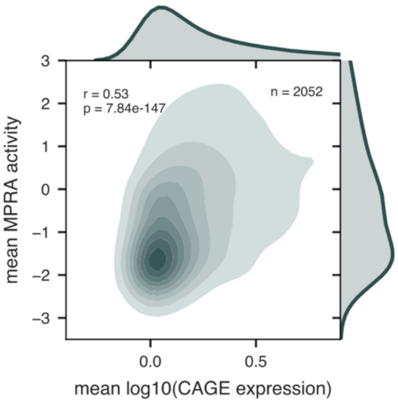

**Supplemental Fig S5: TSS CAGE-seq expression and MPRA activity are correlated.** Mean CAGE expression was calculated across HepG2, HeLa, and K562 samples. Mean MPRA activity was calculated across HepG2, HeLa, and K562. Spearman's rho and *p*-value are shown.

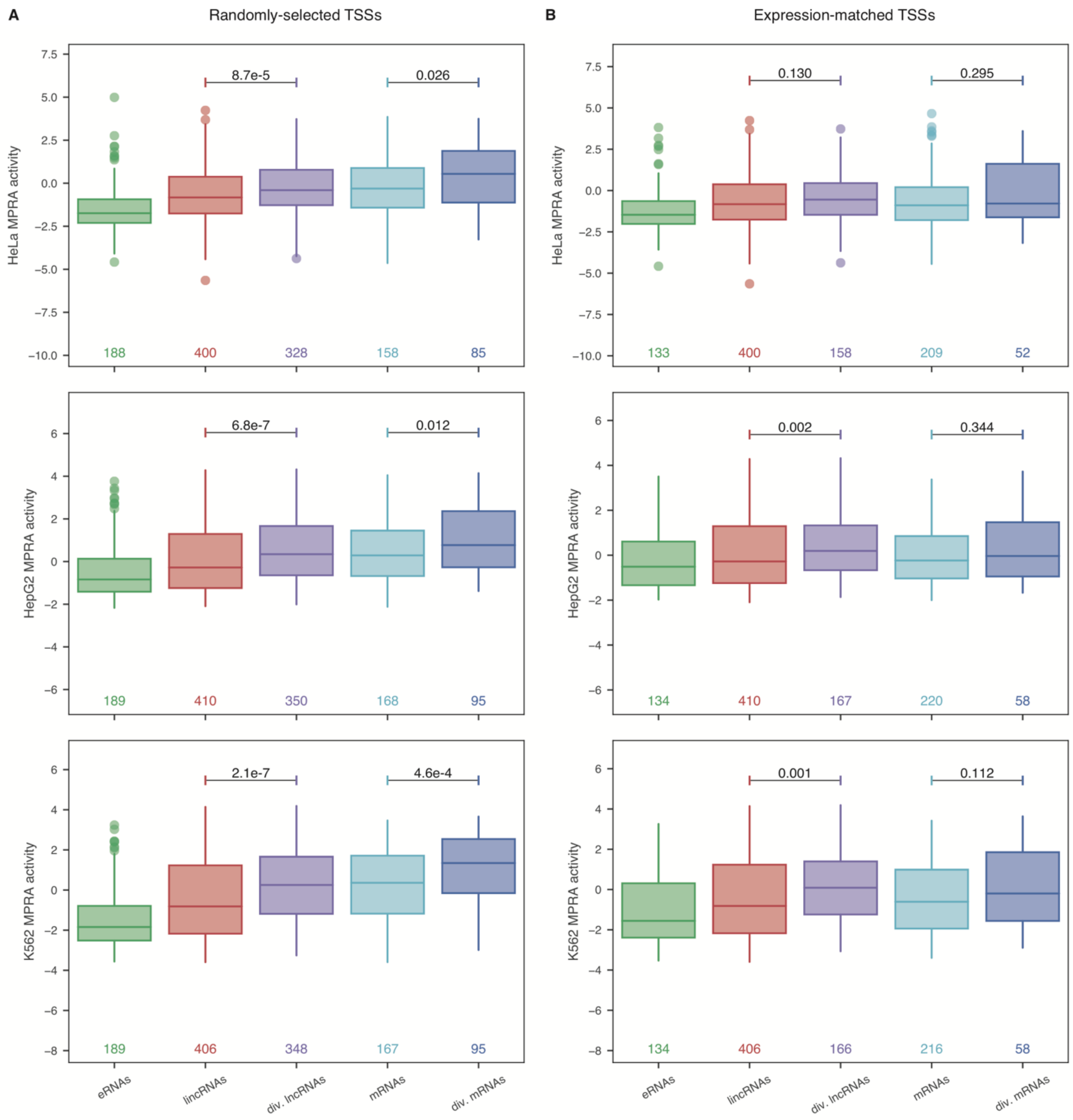

**Supplemental Fig S6: MPRA activity patterns in randomly-selected vs. expression-matched TSSs.** **A.** MPRA activities across TSS classes in randomly-selected TSSs that were included in the original MPRA design. **B.** MPRA activities across TSS classes in expression-matched TSSs. Expression matching was done by matching the average CAGE-seq expression across all samples of eRNAs, mRNAs, and divergent TSSs to the average CAGE-seq expression of lncRNAs using a k-nearest neighbors approach. *P*-values listed are from a two-sided Wilcoxon test.

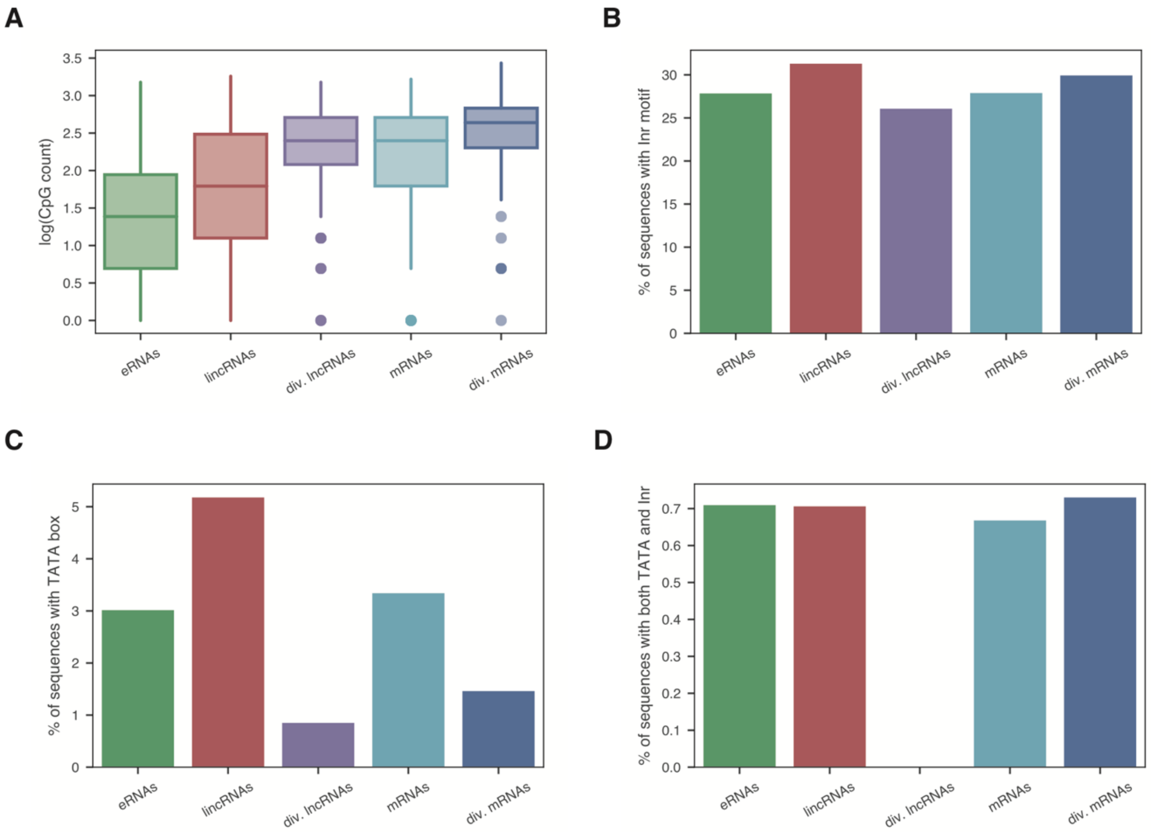

**Supplemental Fig S7: Distribution of core promoter elements across biotypes. A.** CpG count per biotype, where CpG count is the number of "CG" dinucleotides in a 114bp sequence (-80 to +34 from TSS). **B.** Percent of sequences containing an Inr motif, where the Inr motif is defined as BBCABW (B = C/G/T, W = A/T) within 5 nucleotides of the TSS. **C.** Percent of sequences containing a TATA box, where the TATA box is defined as either TATAAA or TATATA within -55 to -15 bp from the TSS. **D.** Percent of sequences with both an Inr motif and TATA box, as defined in B and C above.

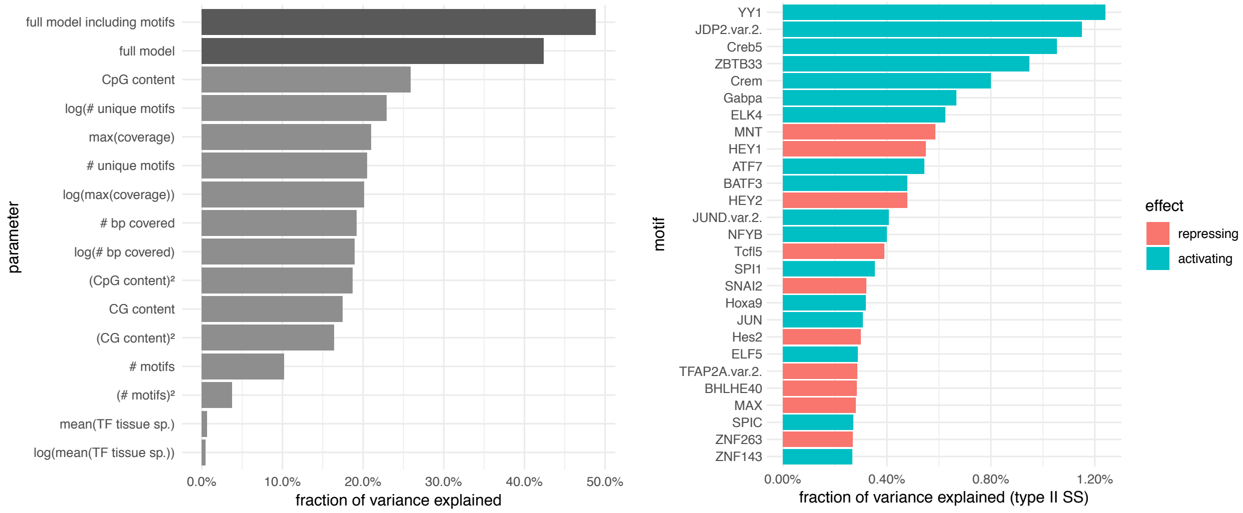

**Supplemental Fig S8: ANOVA analysis of the measured MPRA activity. A.** Fraction of the variance explained by each parameter in the model on its own (light gray) or by all parameters combined together (dark gray) either including or excluding motifs from panel B. **B.** Fraction of the variance explained by each of the 17 TF motifs that significantly contribute to the model. Addition of the 17 TF motifs in (panel B) significantly explains additional variance when including the light gray parameters in panel A (compare the first two dark gray bars in panel A). Both activating and repressing motifs are represented.

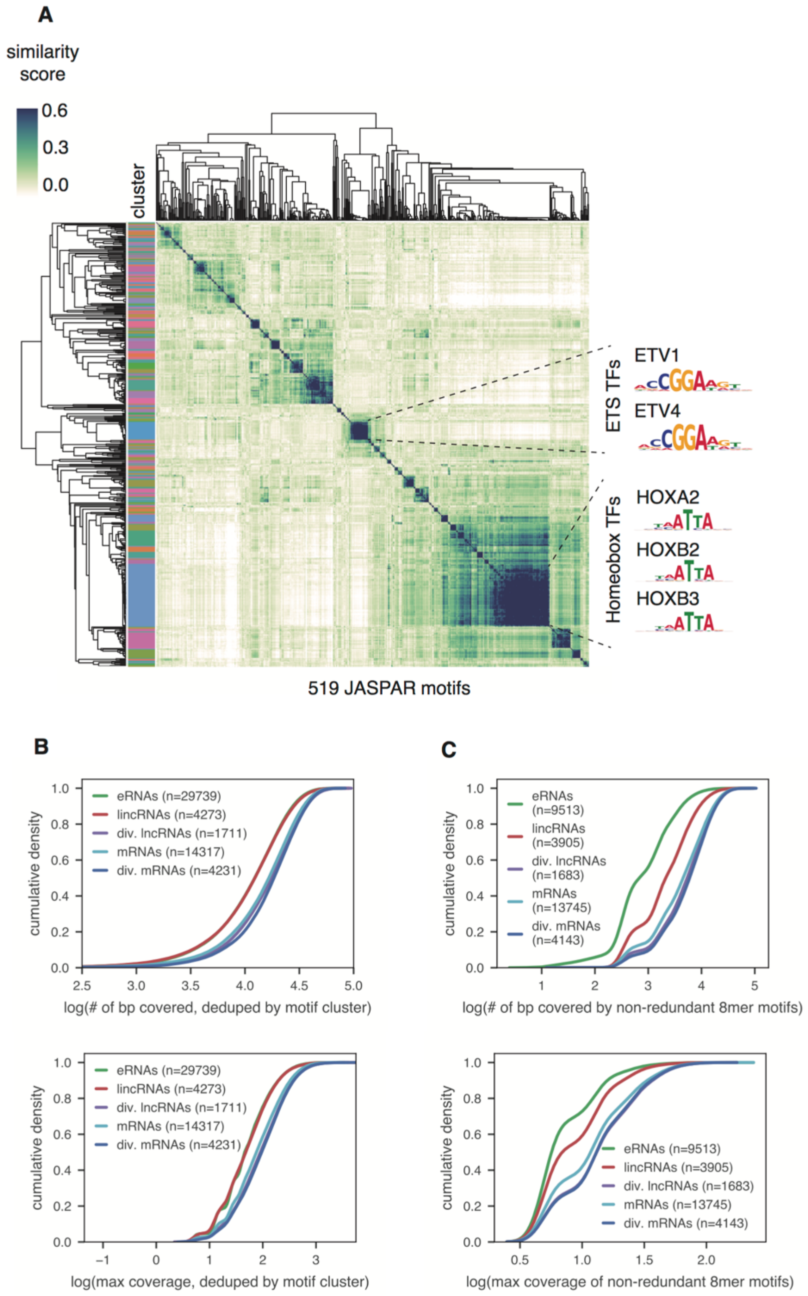

**Supplemental Fig S9: High motif coverage in ubiquitous biotypes is not due to motif redundancy.** **A.** Heatmap shows the MoSBAT similarity scores between all pairwise comparisons of TF motif position weight matrices. To cluster similar motifs, we used average hierarchical clustering using correlation as the pairwise distance measurement. We assigned motifs to clusters using a distance cut-off of 0.1. This clustering methodology successfully clusters the largest similar family of motifs (Homeobox TFs) as well as other similar families, including the ETS TFs. **B.** Cumulative density plot of the number of base pairs covered by a motif (top) and maximum motif coverage (bottom), after de-duping by motif cluster, across all core promoters. **C.** Cumulative density plot of the number of base pairs covered by a motif (top) and maximum motif coverage (bottom), using only a set of published non-redundant 8mer motifs (Mariani et al. 2017), across all core promoters. In B and C, only sequences with at least 1 validated motif were considered.

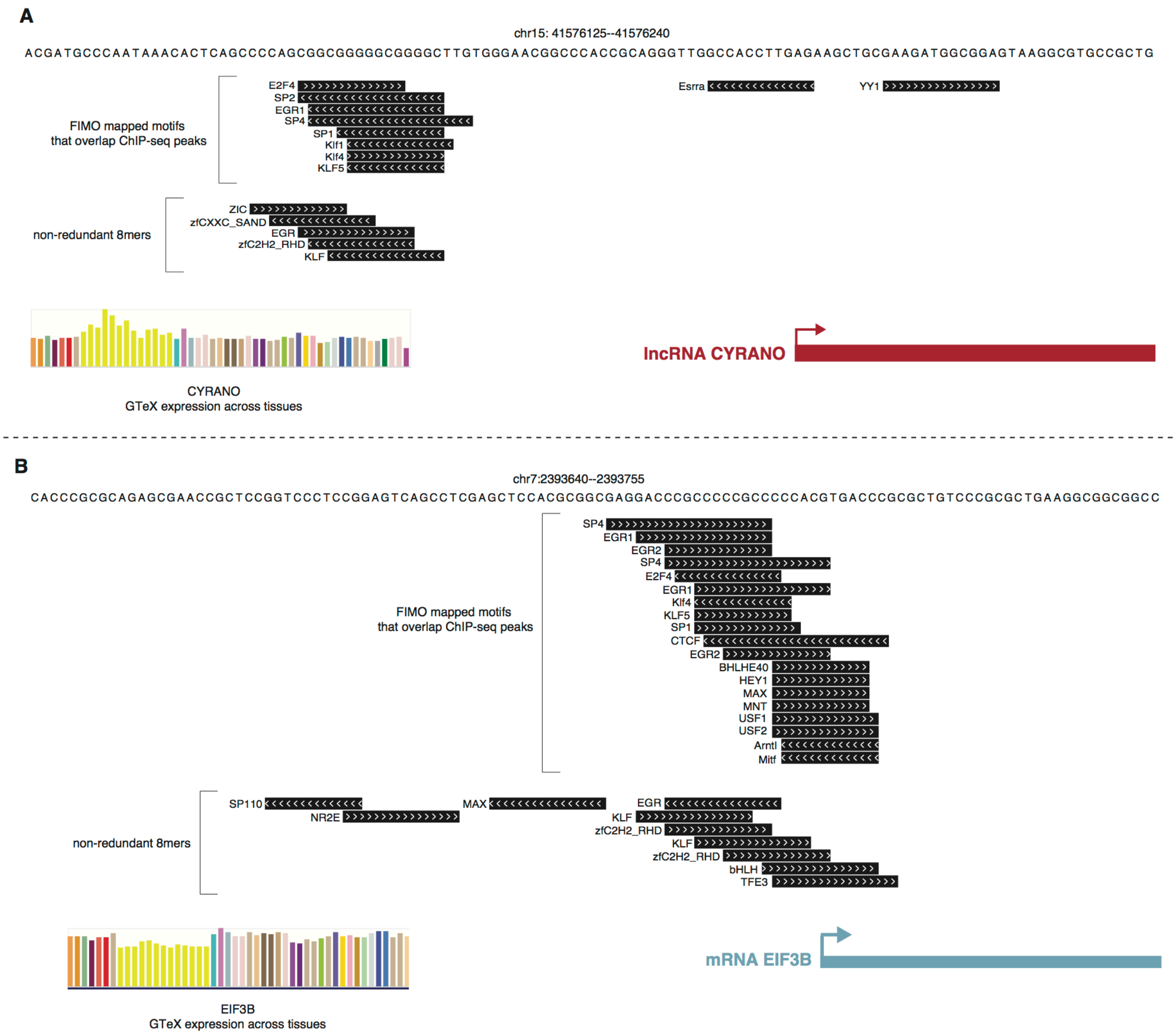

**Supplemental Fig S10: Examples of motif profiles of a ubiquitously-expressed lncRNA (A) and mRNA (B).** UCSC Genome Browser (http://genome.ucsc.edu) screenshots (Kent et al. 2002) of all full FIMO-mapped motifs in the 114bp core promoter regions that intersect with ChIP peaks in Cistrome (Mei et al. 2017) are shown, as well as all non-redundant 8mers (Mariani et al. 2017). The GTEx expression across tissues for the gene of interest is also shown, where each bar color represents a different human tissue. Coordinates listed are in the hg19 assembly.

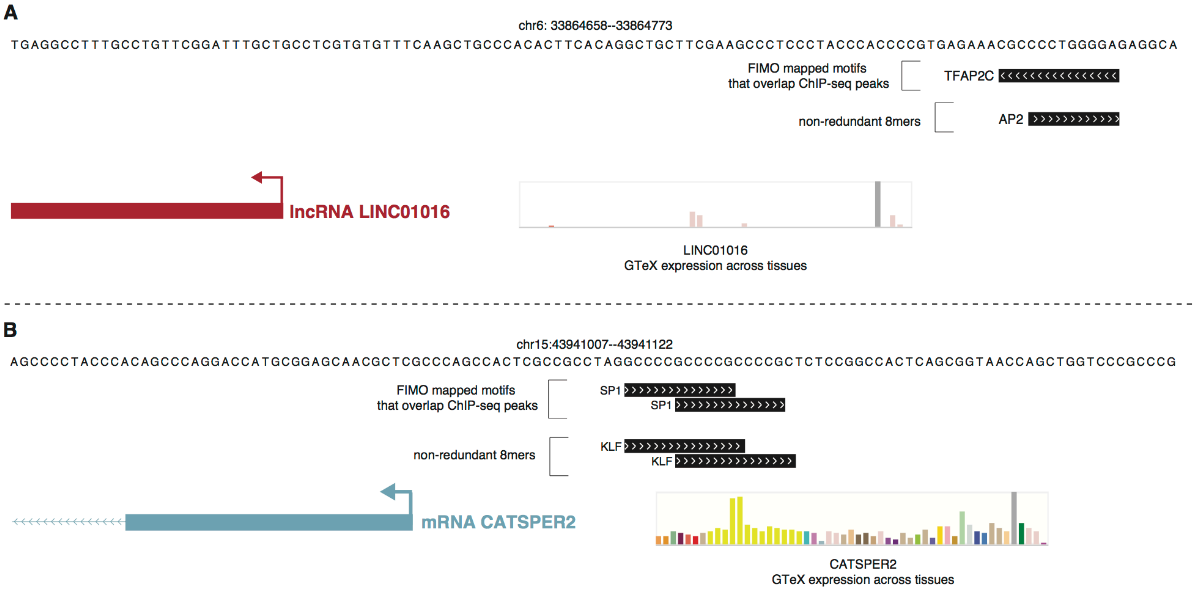

**Supplemental Fig S11: Examples of motif profiles of a tissue-specifically-expressed lncRNA (A) and mRNA (B).** UCSC Genome Browser (http://genome.ucsc.edu) screenshots (Kent et al. 2002) of all full FIMO-mapped motifs in the 114bp core promoter regions that intersect with ChIP peaks in Cistrome (Mei et al. 2017) are shown, as well as all non-redundant 8mers (Mariani et al. 2017). The GTEx expression across tissues for the gene of interest is also shown, where each bar color represents a different human tissue. Coordinates listed are in the hg19 assembly.

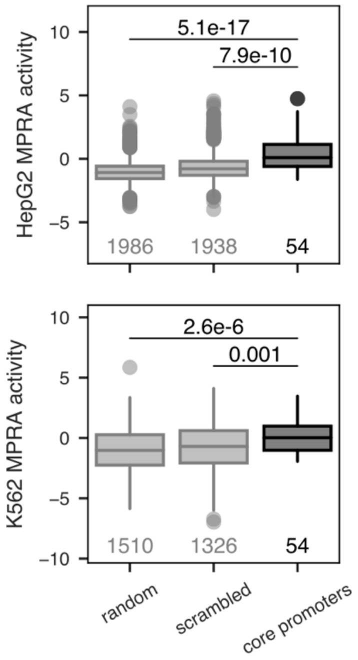

**Supplemental Fig S12: Core promoter sequences are significantly more active than negative controls in deletion MPRA.** Activity of reference sequences compared to random and scrambled sequences in HepG2 (top) and K562 (bottom). Activity is plotted as the mean across replicates of the median barcode activity values per element. *P*-values listed are from a two-sided Wilcoxon test.

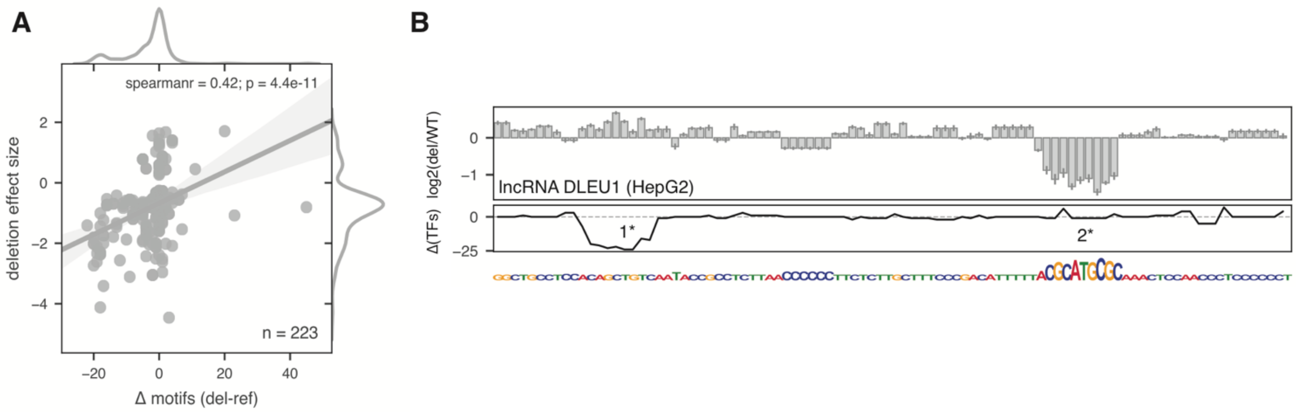

**Supplemental Fig S13: Deletion effect sizes are correlated with the number of motifs predicted to be gained or lost. A.** Correlation between the number of TF motifs predicted to be disrupted by a deletion (x axis) and the effect size of that deletion (y axis, defined as the log2 foldchange between the deletion tile activity and the reference tile activity) for all significant deletions in K562. Spearman's rho and corresponding *p*-value are shown. **B.** Example of a tile (lncRNA DLEU1) that contains both a motif prediction false positive (1*) and false negative (2*). Top panel shows the deletion effect sizes across the sequence, middle panel shows the number of motifs predicted to be disrupted by a given deletion, and the bottom panel shows the actual sequence of the reference tile, with letter heights plotted proportionally to the loss of activity they cause. In the false positive (1*), computational motif mapping predicts a high number of motifs to be disrupted by single nucleotide deletions (middle panel), but the MPRA deletion effect sizes at those positions are minimal (top panel). This is true even when the predicted TF is expressed in the cell line (of the 14 unique TFs with CAGE data that are predicted to bind in region 1*, 6 are expressed in HepG2). In the false negative (2*), computational motif mapping does not predict many motifs to be disrupted (middle panel), but the MPRA deletion effect sizes are very high (top panel).

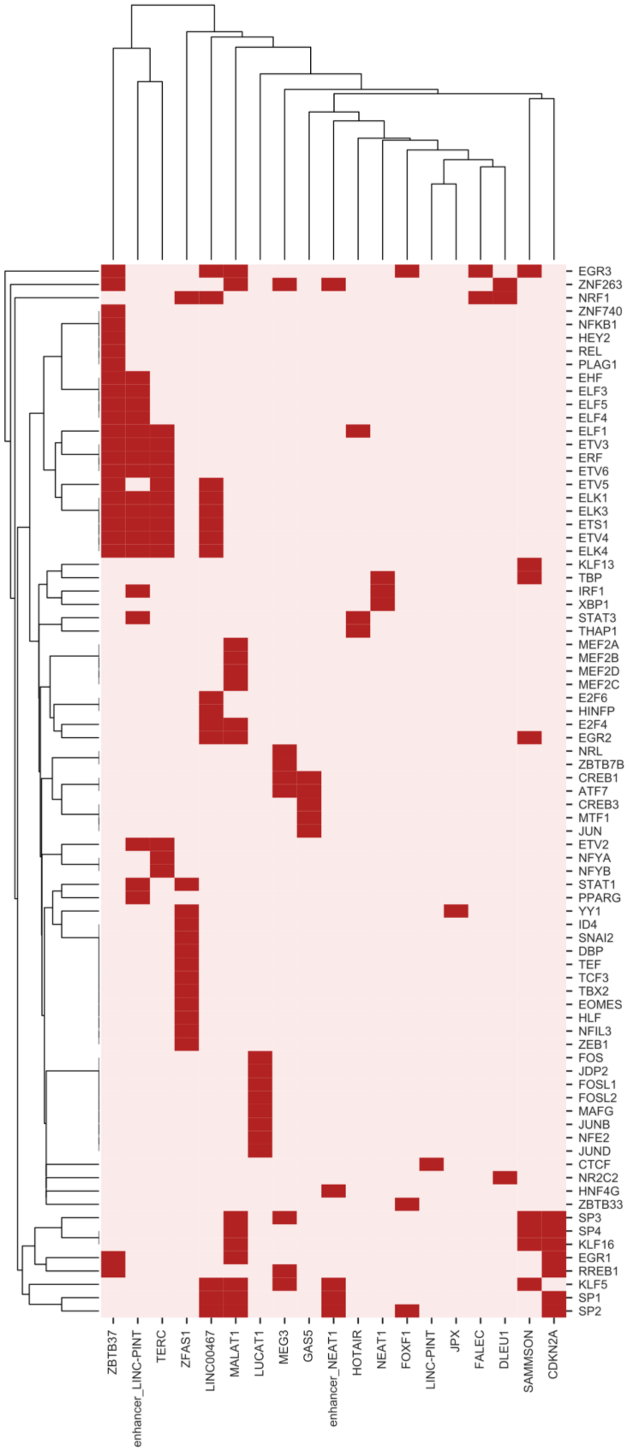

**Supplemental Fig S14: LncRNAs are regulated by a variety of TFs.** Functional TF motifs found across lncRNA core promoter sequences analyzed using targeted deletions. We define functional motifs as those computationally-mapped TF motifs that overlap deletion peaks. Peaks were defined as any stretch of >= 5 nucleotides with effect sizes of <= -1.5 * the average standard deviation of the deletion effect sizes in that tile. Mapped motifs were said to be "functional" if >= 1 nucleotide in the motif intersected a peak. Dark red means a functional motif is present in that core promoter sequence, light red means it is absent.

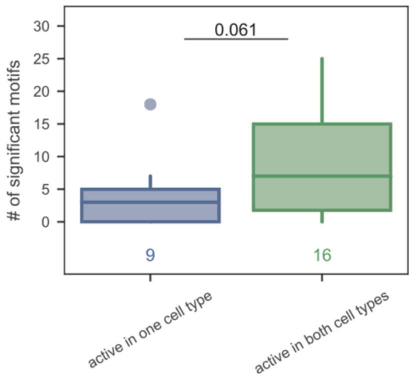

**Supplemental Fig S15: The number of functional TF motifs correlates with cell-type specificity.** Number of functional TF motifs (i.e., computationally-mapped TFs that overlap deletion peaks) in reference tiles that are either active in one cell type only or both cell types. *P*-value listed is from one-sided Wilcoxon test.

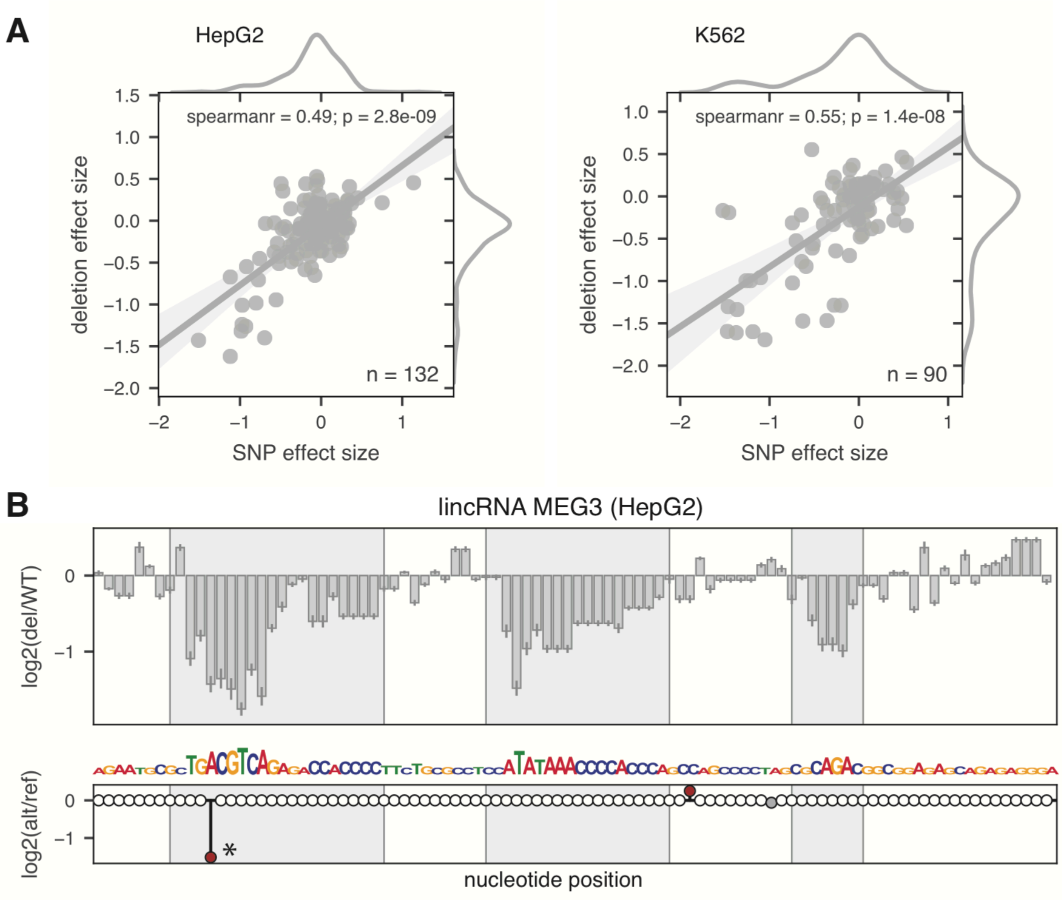

**Supplemental Fig S16: Deletion and SNP effect sizes are correlated. A.** Correlation between deletion effect size (y axis) and SNP effect size (x axis) for all SNPs in HepG2 (left) and K562 (right). Spearman's rho and corresponding *p*-value are shown. **B.** Deletion and SNP effect sizes across the lncRNA *MEG3* promoter in HepG2. Top: deletion effect sizes across nucleotides. Middle: SNP effect sizes across nucleotides. Colored circles indicate tested SNPs: gray circles indicate SNPs that are not called significant; red circles indicate SNPs that are called as significantly down regulatory. Shadowed gray boxes are TF motif peaks called from the single-nucleotide deletion. The starred SNP is a cancer-associated SNP.

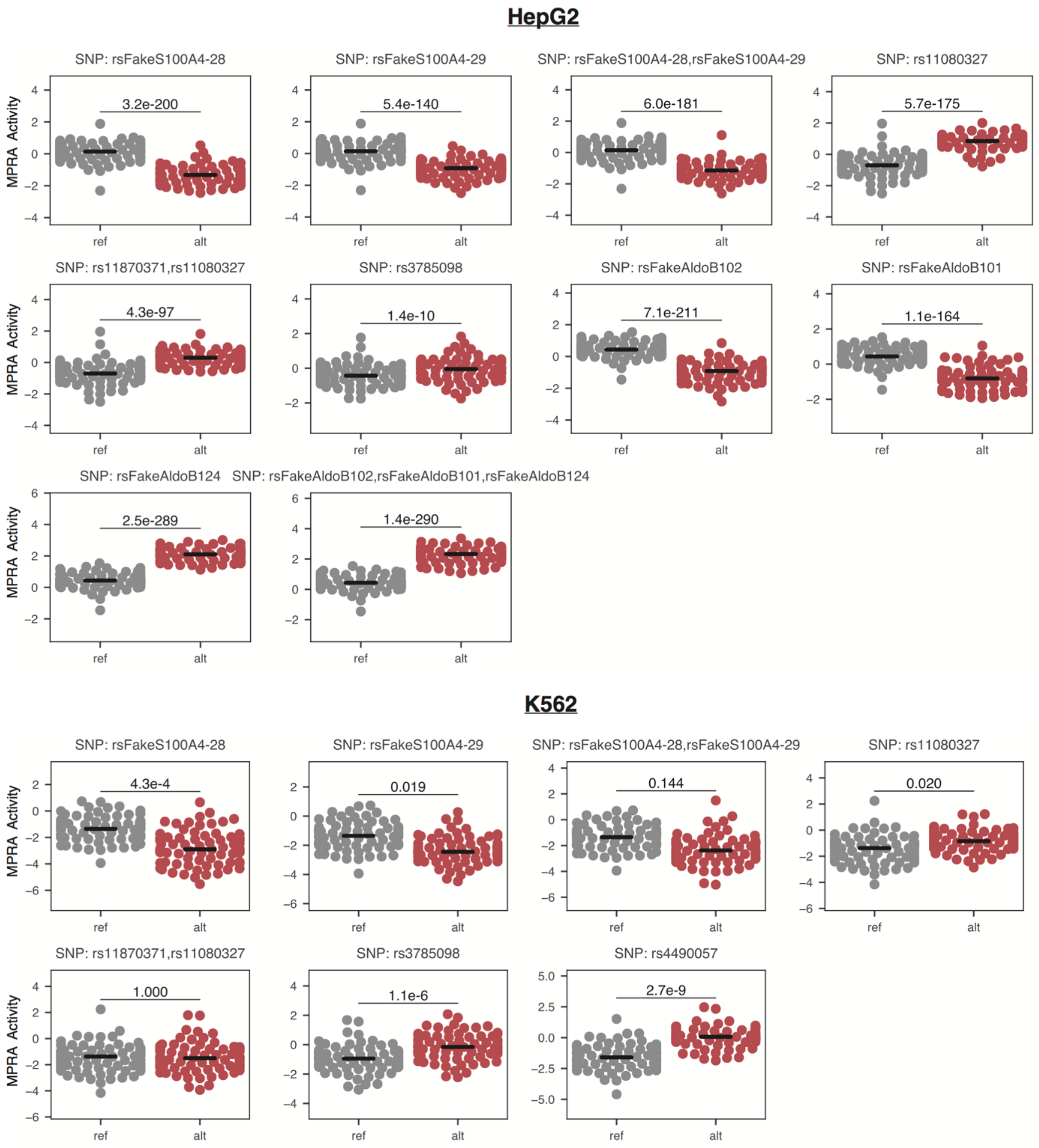

**Supplemental Fig S17: Positive control SNPs are significantly regulatory.** Each plot is an individual SNP, with the reference allele (ref) plotted in gray and the alternative allele (alt) plotted in red. Each dot represents the mean MPRA activity of a barcode across replicates; controls had 80 barcodes. Only SNPs where either the reference tile or the alternative tile (or both) are significantly active (see methods) in the given cell line were analyzed; since more tiles are significantly active in HepG2 than in K562, more controls are plotted in HepG2. *P*-values listed are from adjusted two-sided Wilcoxon tests. *P*-values were all adjusted using the Bonferroni correction. In HepG2, 10/10 controls are significant, and in K562, 5/7 controls are significant at an alpha of 0.05.

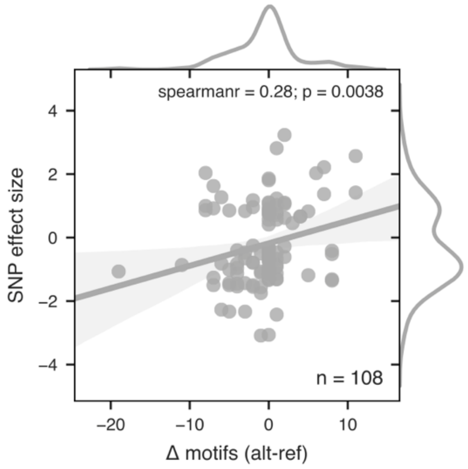

**Supplemental Fig S18: SNP effect sizes correlate with the number of motifs predicted to be disrupted**. Correlation between the number of motifs predicted to be disrupted by the SNP (x axis) and the effect size of the SNP (y axis), defined as the log2 foldchange in activity between the alternative and reference tiles, in K562. Spearman's rho and corresponding *p*-value are shown.

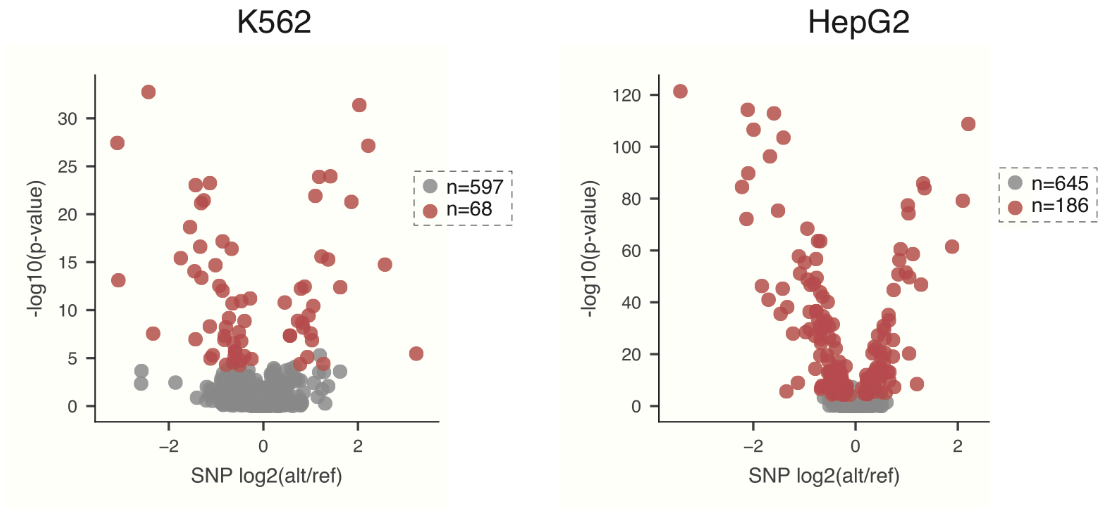

**Supplemental Fig S19: As many as 22% of SNPs in core promoters are regulatory.** Volcano plots for SNPs analyzed in K562 and HepG2 showing the SNP effect size (x axis) and the negative log10 *p*-value (y axis). Only SNPs where either the reference tile or the alternative tile (or both) are significantly active are analyzed (see methods); since more tiles are active in HepG2, more SNPs were analyzed. Red circles correspond to significant SNPs (combined adjusted *p*-value < 0.05). Note that the unadjusted *p*-values are plotted for better visualization but coloring of circles uses the adjusted *p*-values. 10.5% of SNPs are significant in K562; 21.8% of SNPs are significant in HepG2.

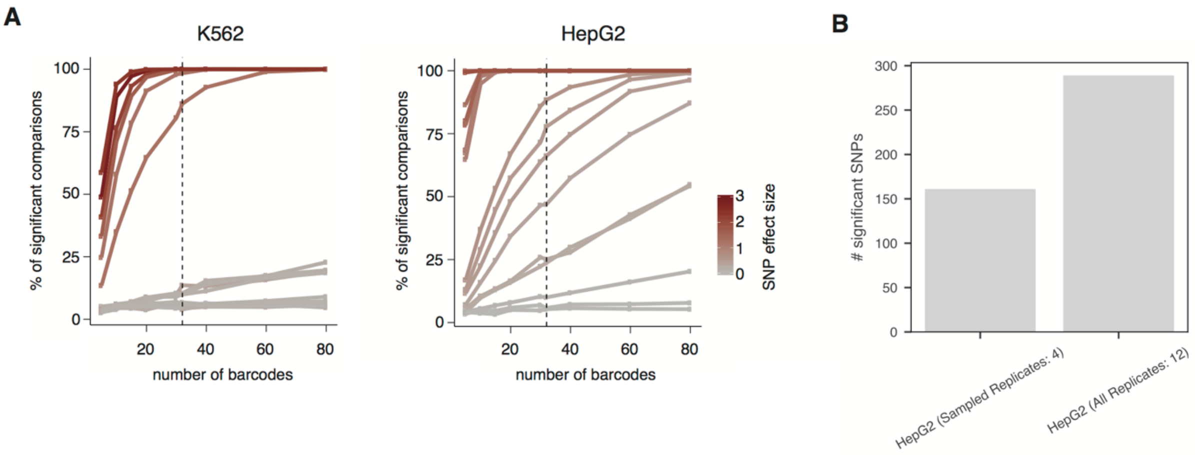

**Supplemental Fig S20: Power to detect regulatory SNPs increases with barcodes and replicates.** **A.** Relationship between the number of barcodes considered (x axis) and the percentage of times we call a SNP regulatory (adjusted *p*-value < 0.05 by two-sided Wilcoxon test) (y axis) in K562 and HepG2. This analysis was done by down-sampling the barcodes of positive control SNPs, which had 80 barcodes. SNPs with different effect sizes are colored accordingly. Straight vertical line is set at 32 barcodes as this is the number of barcodes used to test the majority of SNPs. **B.** The number of significant SNPs (adjusted *p*-value < 0.05 by two-sided Wilcoxon test) found when down-sampling HepG2 replicates to 4 vs using all 12 HepG2 replicates.

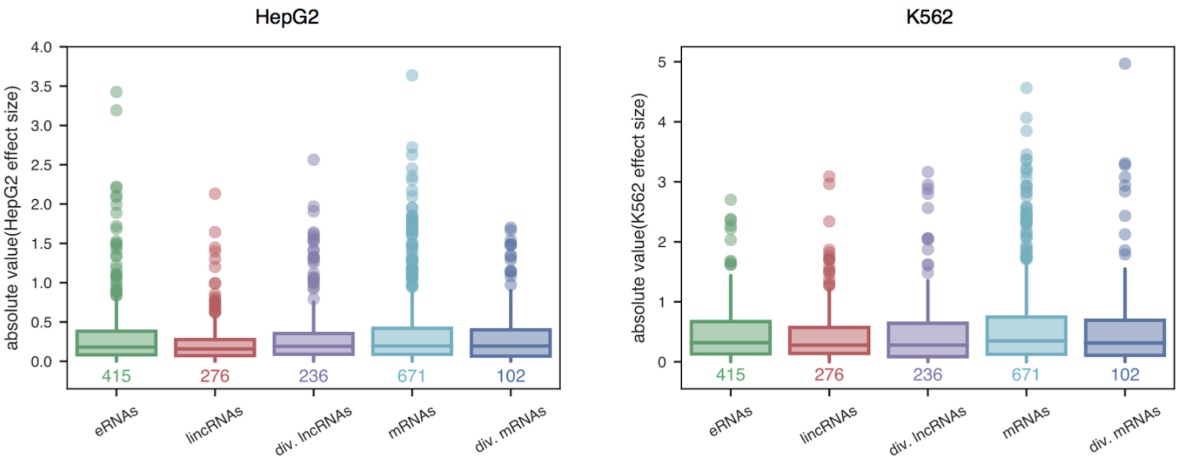

**Supplemental Fig S21: No differences between SNP effect sizes across biotypes.** SNP effect sizes, plotted as the absolute value of the log2 foldchange between alternative (alt) and reference (ref) tiles, per biotype in HepG2 and K562.

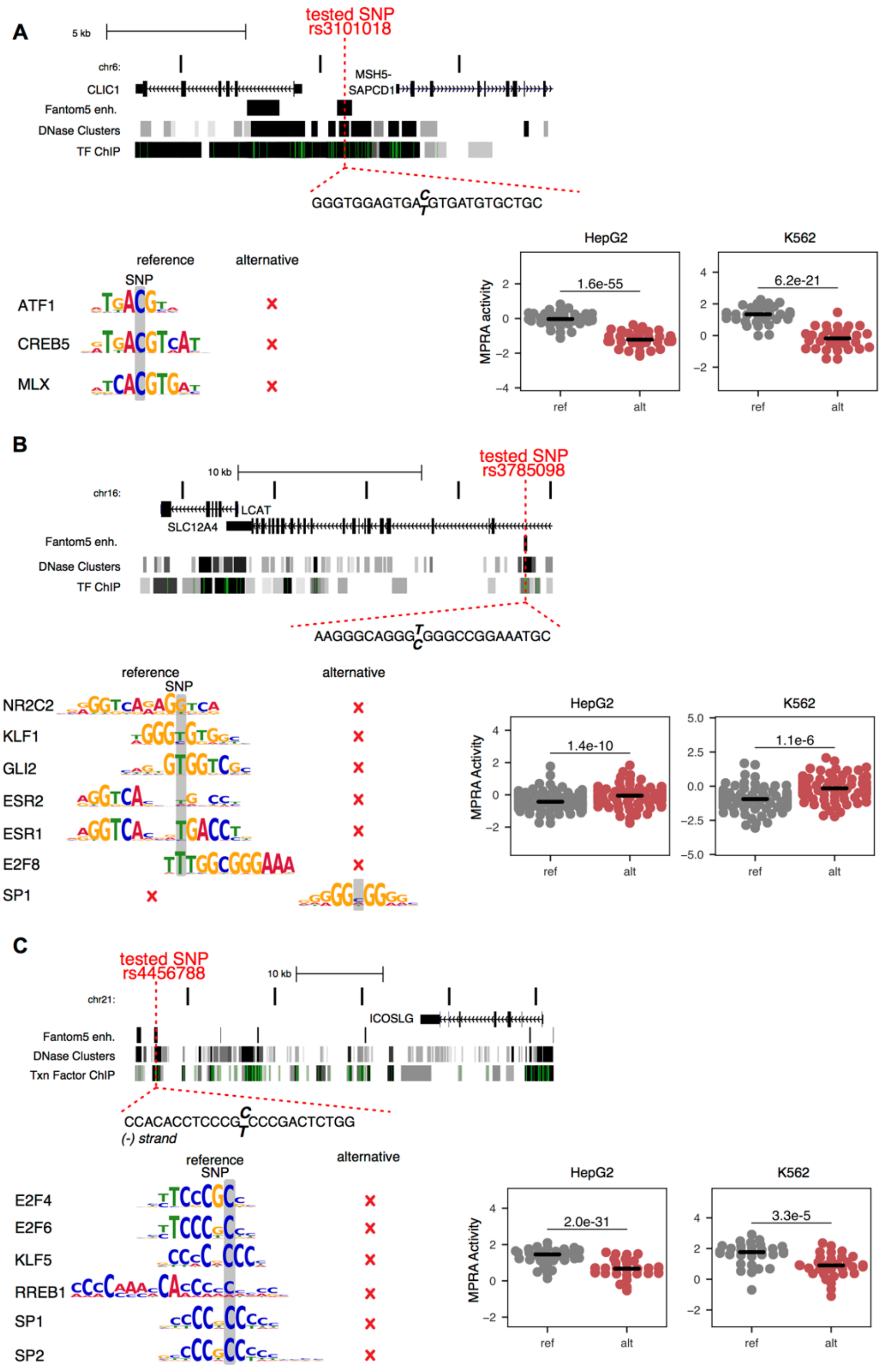

**Supplemental Fig S22: MPRA detects disease-associated regulatory SNPs.** Shown are 3 SNPs that are in LD with GWAS hits (see methods for selection criteria): lung cancer/schizophrenia (**A.**), high HDL cholesterol (**B.**), and inflammatory bowel disease (**C.**). In each panel, the SNP is highlighted in red, and the nearby genes are shown, as are known FANTOM enhancers, DNase clusters, and TF ChIP peaks. The sequence surrounding each SNP is shown in the inset between the red dotted lines, with the reference allele bolded above the alternative allele. Motifs that are predicted to be disrupted by the SNP are shown. Finally, MPRA activities for the reference and alternative alleles in both HepG2 and K562 are shown. *P*-values listed are from a two-sided Wilcoxon test.

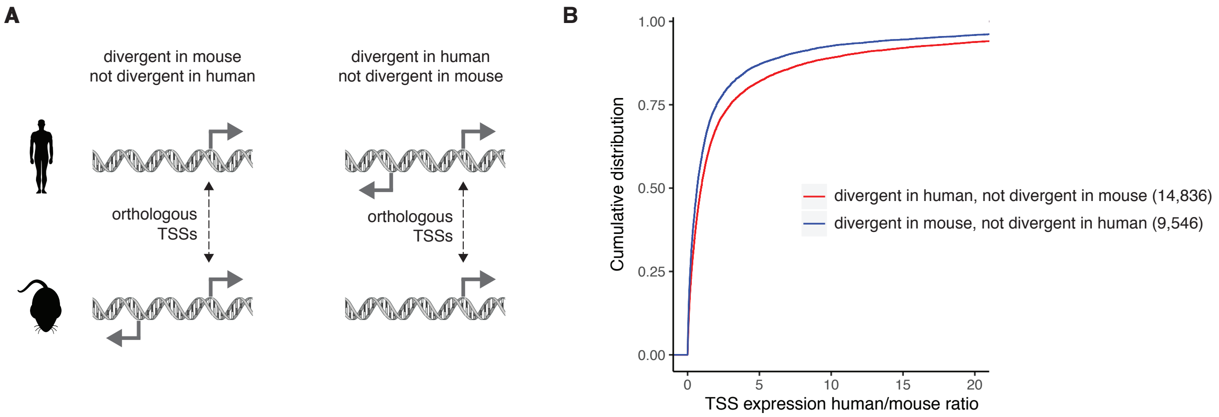

**Supplemental Fig S23: Evolutionary occurrence of an antisense transcript is associated with increased expression of the sense gene. A.** Schematic of analysis performed. We mapped 1:1 orthologous TSSs between human and mouse and found those where an antisense transcript exists (i.e., "divergent" genes) in at least one species (see methods). **B.** Cumulative density plot of the sense TSS expression ratio between human and mouse for the two classifications. Higher TSS expression ratios indicate a sense gene that is more highly expressed in human compared to mouse.

Quek, Xiu Cheng, Daniel W Thomson, Jesper L V Maag, Nenad Bartonicek, Bethany Signal, Michael B Clark, Brian S Gloss, and Marcel E Dinger. 2015. “LncRNAdb v2.0: Expanding the Reference Database for Functional Long Noncoding RNAs.” *Nucleic Acids Research* 43 (Database issue):

D168-73. https://doi.org/10.1093/nar/gku988.
Stouffer, SA, EA Suchman, LC DeVinney, and SA Star. 1949. “The American Soldier: Adjustment during Army Life.(Studies in Social Psychology in World War II), Vol. 1.” http://psycnet.apa.org/record/1950-00790-000.
