## Supplementary material for "High-throughput functional analysis of IncRNA core promoters elucidatesrules governing tissue-specificity": 00__calculate_cage_exp_and_ts.py

```
# coding: utf-8

# # 00__calculate_cage_exp_and_ts
# # calculating CAGE expression and tissue-specificity from FANTOM5 files
# 
# in this notebook, i parse FANTOM5 files and calculate CAGE expression & tissue specificities. i use the groups outlined in Supplemental Table S7 to group samples by sample type and omit ambiguous samples, and then calculate average expression across these groups. i also calculate tissue specificity across these groups and assign TSSs to be either "ubiquitous" (expressed in >90% of groups), "dynamic" (expressed in <90% of groups, but expressed at >50tpm in at least 1 group), or "tissue-specific" (expressed in <90% of groups, and never above 50tpm in any group).
# 
# i also get a list of 'robust' enhancers using the same criteria defined by fantom5 for the 'robust' TSSs: A ‘robust’ threshold, for which a peak must include a CTSS with more than 10 read counts and 1 TPM (tags per million) at least one sample, was employed to define a stringent subset of the CAGE peaks (from [here](https://www.nature.com/articles/sdata2017112))
# 
# ------
# 
# figures in this notebook:
# - **Fig S1**: average CAGE-seq expression and CAGE-seq tissue-specificity across TSS classes

# In[1]:


import warnings
warnings.filterwarnings('ignore')

import pandas as pd
import numpy as np
import matplotlib as mpl
import matplotlib.pyplot as plt
import seaborn as sns
import sys
import time

from os import walk
from scipy.stats import spearmanr

# import utils
sys.path.append("../../utils")
from plotting_utils import *
from misc_utils import *
from norm_utils import *

get_ipython().magic('matplotlib inline')


# In[2]:


sns.set(**PAPER_PRESET)
fontsize = PAPER_FONTSIZE


# ## variables

# In[3]:


all_tss_f = "../../data/00__index/0__all_tss/All.TSS.114bp.bed"


# In[4]:


tss_cage_exp_f = "../../misc/01__cage/TSS.CAGE_grouped_exp.tissue_sp.txt.gz"
enh_cage_exp_f = "../../misc/01__cage/Enh.CAGE_grouped_exp.tissue_sp.txt.gz"


# In[5]:


robust_enh_f = "../../misc/01__cage/human_robust_enhancers.txt"


# ## 1. import data

# In[6]:


all_tss = pd.read_table(all_tss_f, sep="\t", header=None)
all_tss.columns = ["chr", "start", "end", "name", "score", "strand"]
all_tss.head()


# In[7]:


tss_cage_exp = pd.read_table(tss_cage_exp_f, sep="\t")
tss_cage_exp.head()


# In[8]:


tss_cage_exp["cage_id"] = tss_cage_exp["00Annotation"]
tss_cage_exp.drop(["00Annotation", "short_description"], axis=1, inplace=True)


# In[9]:


enh_cage_exp = pd.read_table(enh_cage_exp_f, sep="\t")
enh_cage_exp.head()


# In[10]:


enh_cage_exp["cage_id"] = enh_cage_exp["Id"]
enh_cage_exp.drop(["Id"], axis=1, inplace=True)


# In[11]:


robust_enh = pd.read_table(robust_enh_f, sep="\t", skiprows=1, header=None)
robust_enh.columns = ["cage_id"]
robust_enh.head()


# In[12]:


# limit enhancers to robust only!
enh_cage_exp = enh_cage_exp[enh_cage_exp["cage_id"].isin(robust_enh["cage_id"])]
len(enh_cage_exp)


# In[13]:


all_cage_exp = tss_cage_exp.append(enh_cage_exp)
all_cage_exp.sample(5)


# ## 2. parse CAGE IDs

# In[14]:


def get_cage_id(row):
    if "Enhancer" in row["name"]:
        return row["name"].split("__")[1]
    else:
        return row["name"].split("__")[2]


# In[15]:


all_tss["cage_id"] = all_tss.apply(get_cage_id, axis=1)
all_tss.sample(5)


# ## 3. get av exp & t.s. across all samples

# In[16]:


samples = [x for x in all_cage_exp.columns if "Group_" in x]
all_cage_exp["av_exp"] = all_cage_exp[samples].mean(axis=1)
all_cage_exp.sample(5)


# ## 4. merge

# In[17]:


all_tss = all_tss.merge(all_cage_exp[["cage_id", "av_exp", "tissue_sp_all"]], on="cage_id")
all_tss.sample(5)


# ## 5. define promtype2

# In[18]:


all_tss["PromType2"] = all_tss["name"].str.split("__", expand=True)[0]
all_tss.sample(5)


# In[19]:


all_tss.PromType2.unique()


# ## 5. plot

# In[20]:


all_tss["log_av_exp"] = np.log(all_tss["av_exp"]+1)


# In[21]:


fig = plt.figure(figsize=(3.5, 2.5))
ax = sns.boxplot(data=all_tss, x="PromType2", y="log_av_exp", 
                 flierprops = dict(marker='o', markersize=5), order=TSS_CLASS_ORDER, palette=TSS_CLASS_PALETTE)
ax.set_xticklabels(["eRNAs", "lincRNAs", "div. lncRNAs", "mRNAs", "div. mRNAs"], rotation=30)
mimic_r_boxplot(ax)


# calc p-vals b/w divergent and non
enhs = np.asarray(all_tss[all_tss["PromType2"] == "Enhancer"]["log_av_exp"])
lincs = np.asarray(all_tss[all_tss["PromType2"] == "intergenic"]["log_av_exp"])
div_lncs = np.asarray(all_tss[all_tss["PromType2"] == "div_lnc"]["log_av_exp"])
pcs = np.asarray(all_tss[all_tss["PromType2"] == "protein_coding"]["log_av_exp"])
div_pcs = np.asarray(all_tss[all_tss["PromType2"] == "div_pc"]["log_av_exp"])

enhs = enhs[~np.isnan(enhs)]
lincs = lincs[~np.isnan(lincs)]
div_lncs = div_lncs[~np.isnan(div_lncs)]
pcs = pcs[~np.isnan(pcs)]
div_pcs = div_pcs[~np.isnan(div_pcs)]

lnc_u, lnc_pval = stats.mannwhitneyu(lincs, div_lncs, alternative="two-sided", use_continuity=False)
pc_u, pc_pval = stats.mannwhitneyu(pcs, div_pcs, alternative="two-sided", use_continuity=False)

annotate_pval(ax, 1, 2, 7.3, 0, 0, lnc_pval, fontsize, False, None, None)
annotate_pval(ax, 3, 4, 8.5, 0, 0, pc_pval, fontsize, False, None, None)

plt.ylim((-1, 9.5))
plt.xlabel("")
plt.ylabel("log(mean CAGE expression)")

x_ax_0, y_ax = axis_data_coords_sys_transform(ax, 0, 0, inverse=True)
x_ax_1, y_ax = axis_data_coords_sys_transform(ax, 1, 0, inverse=True)
x_ax_diff = x_ax_1 - x_ax_0

ax.annotate(str(len(enhs)), xy=(x_ax_0, 0.02), xycoords="axes fraction", xytext=(0, 0), 
            textcoords="offset pixels", ha='center', va='bottom', 
            color=TSS_CLASS_PALETTE["Enhancer"], size=fontsize)
ax.annotate(str(len(lincs)), xy=(x_ax_0+(x_ax_diff*1), 0.02), xycoords="axes fraction", xytext=(0, 0), 
            textcoords="offset pixels", ha='center', va='bottom', 
            color=TSS_CLASS_PALETTE["intergenic"], size=fontsize)
ax.annotate(str(len(div_lncs)), xy=(x_ax_0+(x_ax_diff*2), 0.02), xycoords="axes fraction", xytext=(0, 0), 
            textcoords="offset pixels", ha='center', va='bottom', 
            color=TSS_CLASS_PALETTE["div_lnc"], size=fontsize)
ax.annotate(str(len(pcs)), xy=(x_ax_0+(x_ax_diff*3), 0.02), xycoords="axes fraction", xytext=(0, 0), 
            textcoords="offset pixels", ha='center', va='bottom', 
            color=TSS_CLASS_PALETTE["protein_coding"], size=fontsize)
ax.annotate(str(len(div_pcs)), xy=(x_ax_0+(x_ax_diff*4), 0.02), xycoords="axes fraction", xytext=(0, 0), 
            textcoords="offset pixels", ha='center', va='bottom', 
            color=TSS_CLASS_PALETTE["div_pc"], size=fontsize)
    

fig.savefig("Fig_S1A.pdf", dpi="figure", bbox_inches="tight")


# In[22]:


print("lnc pval: %s" % lnc_pval)
print("pc pval: %s" % pc_pval)


# In[23]:


fig = plt.figure(figsize=(3.5, 2.5))
ax = sns.boxplot(data=all_tss, x="PromType2", y="tissue_sp_all", 
                 flierprops = dict(marker='o', markersize=5), order=TSS_CLASS_ORDER, palette=TSS_CLASS_PALETTE)
ax.set_xticklabels(["eRNAs", "lincRNAs", "div. lncRNAs", "mRNAs", "div. mRNAs"], rotation=30)
mimic_r_boxplot(ax)


# calc p-vals b/w divergent and non
enhs = np.asarray(all_tss[all_tss["PromType2"] == "Enhancer"]["tissue_sp_all"])
lincs = np.asarray(all_tss[all_tss["PromType2"] == "intergenic"]["tissue_sp_all"])
div_lncs = np.asarray(all_tss[all_tss["PromType2"] == "div_lnc"]["tissue_sp_all"])
pcs = np.asarray(all_tss[all_tss["PromType2"] == "protein_coding"]["tissue_sp_all"])
div_pcs = np.asarray(all_tss[all_tss["PromType2"] == "div_pc"]["tissue_sp_all"])

enhs = enhs[~np.isnan(enhs)]
lincs = lincs[~np.isnan(lincs)]
div_lncs = div_lncs[~np.isnan(div_lncs)]
pcs = pcs[~np.isnan(pcs)]
div_pcs = div_pcs[~np.isnan(div_pcs)]

lnc_u, lnc_pval = stats.mannwhitneyu(lincs, div_lncs, alternative="two-sided", use_continuity=False)
pc_u, pc_pval = stats.mannwhitneyu(pcs, div_pcs, alternative="two-sided", use_continuity=False)
    
annotate_pval(ax, 1, 2, 1.05, 0, 0, lnc_pval, fontsize, False, None, None)
annotate_pval(ax, 3, 4, 1.05, 0, 0, pc_pval, fontsize, False, None, None)

x_ax_0, y_ax = axis_data_coords_sys_transform(ax, 0, 0, inverse=True)
x_ax_1, y_ax = axis_data_coords_sys_transform(ax, 1, 0, inverse=True)
x_ax_diff = x_ax_1 - x_ax_0

ax.annotate(str(len(enhs)), xy=(x_ax_0, 0.02), xycoords="axes fraction", xytext=(0, 0), 
            textcoords="offset pixels", ha='center', va='bottom', 
            color=TSS_CLASS_PALETTE["Enhancer"], size=fontsize)
ax.annotate(str(len(lincs)), xy=(x_ax_0+(x_ax_diff*1), 0.02), xycoords="axes fraction", xytext=(0, 0), 
            textcoords="offset pixels", ha='center', va='bottom', 
            color=TSS_CLASS_PALETTE["intergenic"], size=fontsize)
ax.annotate(str(len(div_lncs)), xy=(x_ax_0+(x_ax_diff*2), 0.02), xycoords="axes fraction", xytext=(0, 0), 
            textcoords="offset pixels", ha='center', va='bottom', 
            color=TSS_CLASS_PALETTE["div_lnc"], size=fontsize)
ax.annotate(str(len(pcs)), xy=(x_ax_0+(x_ax_diff*3), 0.02), xycoords="axes fraction", xytext=(0, 0), 
            textcoords="offset pixels", ha='center', va='bottom', 
            color=TSS_CLASS_PALETTE["protein_coding"], size=fontsize)
ax.annotate(str(len(div_pcs)), xy=(x_ax_0+(x_ax_diff*4), 0.02), xycoords="axes fraction", xytext=(0, 0), 
            textcoords="offset pixels", ha='center', va='bottom', 
            color=TSS_CLASS_PALETTE["div_pc"], size=fontsize)

plt.ylim((0.2, 1.15))
plt.xlabel("")
plt.ylabel("CAGE tissue-specificity")
fig.savefig("Fig_S1B.pdf", dpi="figure", bbox_inches="tight")


# In[24]:


print("lnc pval: %s" % lnc_pval)
print("pc pval: %s" % pc_pval)


# ## define promoters as "ubiquitous" v "tissue-specific"
# ubiquitous = on in >50% of samples
# dynamic = on in < 50% of samples, in at least 1 sample on at > 10
# tissue-sp = on in < 50% of samples

# In[25]:


all_cage_exp["n_expr"] = all_cage_exp[samples].astype(bool).sum(axis=1)
all_cage_exp.head()


# In[26]:


len(all_cage_exp)


# In[27]:


def expr_type(row, samples, thresh):
    if row.n_expr > 0.9*len(samples):
        return "ubiquitous"
    elif row.n_expr < 0.1*len(samples):
        exprs = list(row[samples])
        over_thresh = any(i >= thresh for i in exprs)
        if over_thresh > 0:
            return "dynamic"
        else:
            return "tissue-specific"
    else:
        return "moderate"

all_cage_exp["tss_type"] = all_cage_exp.apply(expr_type, axis=1, samples=samples, thresh=50)
all_cage_exp.sample(10)


# In[28]:


all_cage_exp.tss_type.value_counts()


# ## write file

# In[29]:


final = all_cage_exp[["cage_id", "av_exp", "tissue_sp_all",  "tissue_sp_3", "n_expr", "tss_type"]]
final.sample(5)


# In[30]:


final[final["tss_type"] == "dynamic"].sample(5)


# In[31]:


final.to_csv("../../misc/01__cage/All_TSS_and_enh.CAGE_grouped_exp.tissue_sp.txt", sep="\t", index=False)


# In[32]:


len(final)


# In[ ]:
```
