## Supplementary material for "High-throughput functional analysis of IncRNA core promoters elucidatesrules governing tissue-specificity": 01__preprocess.py

```
# coding: utf-8

# # 01__preprocess
# # turn files of counts into large dataframe of counts
# in this notebook, I load in the barcode counts that we got from the .fastq files and put everything in one nice dataframe. additionally, I filter the barcodes to only include those that have >= 5 reads in a given sample, and then filter them again to only include those that correspond to TSSs (elements) for which we have at least 3 barcodes represented.
# 
# - throughout these notebooks, "pool1" refers to the TSS pool, while "pool2" refers to the deletion pool
# - pool1 was performed in HepG2, HeLa, and K562; pool2 was performed in HepG2 and K562
# 
# ------
# 
# figures in this notebook:
# - **Fig S3A**: heatmap of barcode count correlation between replicates

# In[1]:


import warnings
warnings.filterwarnings('ignore')

import pandas as pd
import matplotlib.pyplot as plt
import numpy as np
import re
import seaborn as sns
import sys

# import utils
sys.path.append("../../utils")
from plotting_utils import *

get_ipython().magic('matplotlib inline')


# In[2]:


sns.set(**PAPER_PRESET)
fontsize = PAPER_FONTSIZE


# ## functions

# In[3]:


def import_dna(counts_dir, dna_f):
    """
    function that will import the DNA counts for a given experiment
    ----
    counts_dir: string defining where the counts files are stored
    dna_f: list of strings defining where the DNA counts files are
    
    returns a dataframe with barcodes + counts
    """
    dna_dfs = []
    for i in range(len(dna_f)):
        f = dna_f[i]
        cols = ["barcode", "dna_%s" % (i+1)]
        tmp = pd.read_table("%s/%s" % (counts_dir, f), sep="\t", header=None)
        tmp.columns = cols
        dna_dfs.append(tmp)
    if len(dna_dfs) > 1:
        dna = reduce(lambda x, y: pd.merge(x, y, on = "barcode"), dna_dfs)
    else:
        dna = dna_dfs[0]
    return dna


# In[4]:


def import_rna(counts_dir, rna_f, dna):
    """
    function that will import the RNA counts for a given experiment
    ----
    counts_dir: string defining where the counts files are stored
    rna_f: list of strings defining where the RNA counts files are
    dna: dataframe w/ DNA counts
    
    returns a dataframe with barcodes + counts, incl. RNA
    """
    data = dna.copy()
    data_cols = list(dna.columns)
    for f in rna_f:
        rep = re.findall(r'\d+', f.split("__")[3])[0]
        tmp = pd.read_table("%s/%s" % (counts_dir, f), sep="\t", header=None)
        tmp.columns = ["barcode", "rna_%s" % rep]
        data_cols.append("rna_%s" % rep)
        data = data.merge(tmp, on="barcode", how="outer")
    return data, data_cols


# ## variables

# In[5]:


counts_dir = "../../data/01__counts"


# In[6]:


barcode_dna_read_threshold = 5
barcode_rna_read_threshold = 5
n_barcodes_per_elem_threshold = 3


# ### DNA files

# In[7]:


pool1_pMPRA1_dna_f = ["POOL1__pMPRA1__COUNTS.txt"]
pool1_pNoCMVMPRA1_dna_f = ["POOL1__pNoCMVMPRA1__COUNTS.txt"]
pool2_pMPRA1_dna_f = ["POOL2__pMPRA1__COUNTS.txt"]


# ### RNA files

# In[8]:


pool1_pMPRA1_HeLa_rna_f = ["POOL1__pMPRA1__HeLa__rep1__COUNTS.txt",
                           "POOL1__pMPRA1__HeLa__rep2__COUNTS.txt",
                           "POOL1__pMPRA1__HeLa__rep3__COUNTS.txt", 
                           "POOL1__pMPRA1__HeLa__rep4__COUNTS.txt"]
pool1_pMPRA1_HeLa_out_f = "POOL1__pMPRA1__HeLa__all_counts.txt"


# In[9]:


pool1_pMPRA1_HepG2_rna_f = ["POOL1__pMPRA1__HepG2__rep3__COUNTS.txt",
                            "POOL1__pMPRA1__HepG2__rep4__COUNTS.txt",
                            "POOL1__pMPRA1__HepG2__rep5__COUNTS.txt", 
                            "POOL1__pMPRA1__HepG2__rep6__COUNTS.txt",
                            "POOL1__pMPRA1__HepG2__rep7__COUNTS.txt",
                            "POOL1__pMPRA1__HepG2__rep8__COUNTS.txt",
                            "POOL1__pMPRA1__HepG2__rep9__COUNTS.txt", 
                            "POOL1__pMPRA1__HepG2__rep10__COUNTS.txt",
                            "POOL1__pMPRA1__HepG2__rep11__COUNTS.txt",
                            "POOL1__pMPRA1__HepG2__rep12__COUNTS.txt",
                            "POOL1__pMPRA1__HepG2__rep13__COUNTS.txt", 
                            "POOL1__pMPRA1__HepG2__rep14__COUNTS.txt"]
pool1_pMPRA1_HepG2_out_f = "POOL1__pMPRA1__HepG2__all_counts.txt"


# In[10]:


pool1_pMPRA1_K562_rna_f = ["POOL1__pMPRA1__K562__rep1__COUNTS.txt",
                           "POOL1__pMPRA1__K562__rep2__COUNTS.txt",
                           "POOL1__pMPRA1__K562__rep3__COUNTS.txt", 
                           "POOL1__pMPRA1__K562__rep4__COUNTS.txt"]
pool1_pMPRA1_K562_out_f = "POOL1__pMPRA1__K562__all_counts.txt"


# In[11]:


pool1_pNoCMVMPRA1_HeLa_rna_f = ["POOL1__pNoCMVMPRA1__HeLa__rep1__COUNTS.txt",
                                "POOL1__pNoCMVMPRA1__HeLa__rep2__COUNTS.txt",
                                "POOL1__pNoCMVMPRA1__HeLa__rep3__COUNTS.txt", 
                                "POOL1__pNoCMVMPRA1__HeLa__rep4__COUNTS.txt"]
pool1_pNoCMVMPRA1_HeLa_out_f = "POOL1__pNoCMVMPRA1__HeLa__all_counts.txt"


# In[12]:


pool1_pNoCMVMPRA1_HepG2_rna_f = ["POOL1__pNoCMVMPRA1__HepG2__rep3__COUNTS.txt",
                                 "POOL1__pNoCMVMPRA1__HepG2__rep4__COUNTS.txt",
                                 "POOL1__pNoCMVMPRA1__HepG2__rep5__COUNTS.txt", 
                                 "POOL1__pNoCMVMPRA1__HepG2__rep6__COUNTS.txt"]
pool1_pNoCMVMPRA1_HepG2_out_f = "POOL1__pNoCMVMPRA1__HepG2__all_counts.txt"


# In[13]:


pool1_pNoCMVMPRA1_K562_rna_f = ["POOL1__pNoCMVMPRA1__K562__rep1__COUNTS.txt",
                                "POOL1__pNoCMVMPRA1__K562__rep2__COUNTS.txt",
                                "POOL1__pNoCMVMPRA1__K562__rep3__COUNTS.txt", 
                                "POOL1__pNoCMVMPRA1__K562__rep4__COUNTS.txt"]
pool1_pNoCMVMPRA1_K562_out_f = "POOL1__pNoCMVMPRA1__K562__all_counts.txt"


# In[14]:


pool2_pMPRA1_HepG2_rna_f = ["POOL2__pMPRA1__HepG2__rep3__COUNTS.txt",
                            "POOL2__pMPRA1__HepG2__rep4__COUNTS.txt",
                            "POOL2__pMPRA1__HepG2__rep5__COUNTS.txt", 
                            "POOL2__pMPRA1__HepG2__rep6__COUNTS.txt",
                            "POOL2__pMPRA1__HepG2__rep7__COUNTS.txt",
                            "POOL2__pMPRA1__HepG2__rep8__COUNTS.txt",
                            "POOL2__pMPRA1__HepG2__rep9__COUNTS.txt", 
                            "POOL2__pMPRA1__HepG2__rep10__COUNTS.txt"]
pool2_pMPRA1_HepG2_out_f = "POOL2__pMPRA1__HepG2__all_counts.txt"


# In[15]:


pool2_pMPRA1_K562_rna_f = ["POOL2__pMPRA1__K562__rep1__COUNTS.txt",
                           "POOL2__pMPRA1__K562__rep2__COUNTS.txt",
                           "POOL2__pMPRA1__K562__rep3__COUNTS.txt", 
                           "POOL2__pMPRA1__K562__rep4__COUNTS.txt"]
pool2_pMPRA1_K562_out_f = "POOL2__pMPRA1__K562__all_counts.txt"


# ### Index Files

# In[16]:


pool1_index_f = "../../data/00__index/tss_oligo_pool.index.txt"
pool2_index_f = "../../data/00__index/dels_oligo_pool.index.txt"


# ## 1. import indexes

# In[17]:


pool1_index = pd.read_table(pool1_index_f, sep="\t")
pool2_index = pd.read_table(pool2_index_f, sep="\t")
pool1_index.head()


# ## 2. import dna

# In[18]:


pool1_pMPRA1_dna = import_dna(counts_dir, pool1_pMPRA1_dna_f)
pool1_pMPRA1_dna.head()


# In[19]:


pool1_pNoCMVMPRA1_dna = import_dna(counts_dir, pool1_pNoCMVMPRA1_dna_f)
pool1_pNoCMVMPRA1_dna.head()


# In[20]:


pool2_pMPRA1_dna = import_dna(counts_dir, pool2_pMPRA1_dna_f)
pool2_pMPRA1_dna.head()


# ## 3. import rna

# In[21]:


pool1_pMPRA1_HeLa_data, pool1_pMPRA1_HeLa_cols = import_rna(counts_dir, pool1_pMPRA1_HeLa_rna_f, pool1_pMPRA1_dna)
pool1_pMPRA1_HeLa_data.head()


# In[22]:


pool1_pMPRA1_HepG2_data, pool1_pMPRA1_HepG2_cols = import_rna(counts_dir, pool1_pMPRA1_HepG2_rna_f, pool1_pMPRA1_dna)
pool1_pMPRA1_HepG2_data.head()


# In[23]:


pool1_pMPRA1_K562_data, pool1_pMPRA1_K562_cols = import_rna(counts_dir, pool1_pMPRA1_K562_rna_f, pool1_pMPRA1_dna)
pool1_pMPRA1_K562_data.head()


# In[24]:


pool1_pNoCMVMPRA1_HeLa_data, pool1_pNoCMVMPRA1_HeLa_cols = import_rna(counts_dir, pool1_pNoCMVMPRA1_HeLa_rna_f, pool1_pNoCMVMPRA1_dna)
pool1_pNoCMVMPRA1_HeLa_data.head()


# In[25]:


pool1_pNoCMVMPRA1_HepG2_data, pool1_pNoCMVMPRA1_HepG2_cols = import_rna(counts_dir, pool1_pNoCMVMPRA1_HepG2_rna_f, pool1_pNoCMVMPRA1_dna)
pool1_pNoCMVMPRA1_HepG2_data.head()


# In[26]:


pool1_pNoCMVMPRA1_K562_data, pool1_pNoCMVMPRA1_K562_cols = import_rna(counts_dir, pool1_pNoCMVMPRA1_K562_rna_f, pool1_pNoCMVMPRA1_dna)
pool1_pNoCMVMPRA1_K562_data.head()


# In[27]:


pool2_pMPRA1_HepG2_data, pool2_pMPRA1_HepG2_cols = import_rna(counts_dir, pool2_pMPRA1_HepG2_rna_f, pool2_pMPRA1_dna)
pool2_pMPRA1_HepG2_data.head()


# In[28]:


pool2_pMPRA1_K562_data, pool2_pMPRA1_K562_cols = import_rna(counts_dir, pool2_pMPRA1_K562_rna_f, pool2_pMPRA1_dna)
pool2_pMPRA1_K562_data.head()


# ## 4. filter barcodes

# In[29]:


pool1_pMPRA1_HeLa_data = pool1_pMPRA1_HeLa_data.fillna(0)
pool1_pMPRA1_HepG2_data = pool1_pMPRA1_HepG2_data.fillna(0)
pool1_pMPRA1_K562_data = pool1_pMPRA1_K562_data.fillna(0)
pool1_pNoCMVMPRA1_HeLa_data = pool1_pNoCMVMPRA1_HeLa_data.fillna(0)
pool1_pNoCMVMPRA1_HepG2_data = pool1_pNoCMVMPRA1_HepG2_data.fillna(0)
pool1_pNoCMVMPRA1_K562_data = pool1_pNoCMVMPRA1_K562_data.fillna(0)
pool2_pMPRA1_HepG2_data = pool2_pMPRA1_HepG2_data.fillna(0)
pool2_pMPRA1_K562_data = pool2_pMPRA1_K562_data.fillna(0)


# In[30]:


pool1_pMPRA1_HeLa_data.head()


# In[31]:


pool1_pMPRA1_HeLa_data_filt = pool1_pMPRA1_HeLa_data[pool1_pMPRA1_HeLa_data["dna_1"] >= barcode_dna_read_threshold]
pool1_pMPRA1_HepG2_data_filt = pool1_pMPRA1_HepG2_data[pool1_pMPRA1_HepG2_data["dna_1"] >= barcode_dna_read_threshold]
pool1_pMPRA1_K562_data_filt = pool1_pMPRA1_K562_data[pool1_pMPRA1_K562_data["dna_1"] >= barcode_dna_read_threshold]
pool1_pNoCMVMPRA1_HeLa_data_filt = pool1_pNoCMVMPRA1_HeLa_data[pool1_pNoCMVMPRA1_HeLa_data["dna_1"] >= barcode_dna_read_threshold]
pool1_pNoCMVMPRA1_HepG2_data_filt = pool1_pNoCMVMPRA1_HepG2_data[pool1_pNoCMVMPRA1_HepG2_data["dna_1"] >= barcode_dna_read_threshold]
pool1_pNoCMVMPRA1_K562_data_filt = pool1_pNoCMVMPRA1_K562_data[pool1_pNoCMVMPRA1_K562_data["dna_1"] >= barcode_dna_read_threshold]
pool2_pMPRA1_HepG2_data_filt = pool2_pMPRA1_HepG2_data[pool2_pMPRA1_HepG2_data["dna_1"] >= barcode_dna_read_threshold]
pool2_pMPRA1_K562_data_filt = pool2_pMPRA1_K562_data[pool2_pMPRA1_K562_data["dna_1"] >= barcode_dna_read_threshold]


# In[32]:


pool1_pMPRA1_HeLa_reps = [x for x in pool1_pMPRA1_HeLa_data_filt.columns if "rna_" in x]
pool1_pMPRA1_HepG2_reps = [x for x in pool1_pMPRA1_HepG2_data_filt.columns if "rna_" in x]
pool1_pMPRA1_K562_reps = [x for x in pool1_pMPRA1_K562_data_filt.columns if "rna_" in x]
pool1_pNoCMVMPRA1_HeLa_reps = [x for x in pool1_pNoCMVMPRA1_HeLa_data_filt.columns if "rna_" in x]
pool1_pNoCMVMPRA1_HepG2_reps = [x for x in pool1_pNoCMVMPRA1_HepG2_data_filt.columns if "rna_" in x]
pool1_pNoCMVMPRA1_K562_reps = [x for x in pool1_pNoCMVMPRA1_K562_data_filt.columns if "rna_" in x]
pool2_pMPRA1_HepG2_reps = [x for x in pool2_pMPRA1_HepG2_data_filt.columns if "rna_" in x]
pool2_pMPRA1_K562_reps = [x for x in pool2_pMPRA1_K562_data_filt.columns if "rna_" in x]


# In[33]:


pool1_pMPRA1_HeLa_data_filt[pool1_pMPRA1_HeLa_reps] = pool1_pMPRA1_HeLa_data_filt[pool1_pMPRA1_HeLa_data_filt > barcode_rna_read_threshold][pool1_pMPRA1_HeLa_reps]
pool1_pMPRA1_HepG2_data_filt[pool1_pMPRA1_HepG2_reps] = pool1_pMPRA1_HepG2_data_filt[pool1_pMPRA1_HepG2_data_filt > barcode_rna_read_threshold][pool1_pMPRA1_HepG2_reps]
pool1_pMPRA1_K562_data_filt[pool1_pMPRA1_K562_reps] = pool1_pMPRA1_K562_data_filt[pool1_pMPRA1_K562_data_filt > barcode_rna_read_threshold][pool1_pMPRA1_K562_reps]
pool1_pNoCMVMPRA1_HeLa_data_filt[pool1_pNoCMVMPRA1_HeLa_reps] = pool1_pNoCMVMPRA1_HeLa_data_filt[pool1_pNoCMVMPRA1_HeLa_data_filt > barcode_rna_read_threshold][pool1_pNoCMVMPRA1_HeLa_reps]
pool1_pNoCMVMPRA1_HepG2_data_filt[pool1_pNoCMVMPRA1_HepG2_reps] = pool1_pNoCMVMPRA1_HepG2_data_filt[pool1_pNoCMVMPRA1_HepG2_data_filt > barcode_rna_read_threshold][pool1_pNoCMVMPRA1_HepG2_reps]
pool1_pNoCMVMPRA1_K562_data_filt[pool1_pNoCMVMPRA1_K562_reps] = pool1_pNoCMVMPRA1_K562_data_filt[pool1_pNoCMVMPRA1_K562_data_filt > barcode_rna_read_threshold][pool1_pNoCMVMPRA1_K562_reps]
pool2_pMPRA1_HepG2_data_filt[pool2_pMPRA1_HepG2_reps] = pool2_pMPRA1_HepG2_data_filt[pool2_pMPRA1_HepG2_data_filt > barcode_rna_read_threshold][pool2_pMPRA1_HepG2_reps]
pool2_pMPRA1_K562_data_filt[pool2_pMPRA1_K562_reps] = pool2_pMPRA1_K562_data_filt[pool2_pMPRA1_K562_data_filt > barcode_rna_read_threshold][pool2_pMPRA1_K562_reps]
pool1_pMPRA1_HeLa_data_filt.head()


# In[34]:


all_names = ["pool1_pMPRA1_HeLa", "pool1_pMPRA1_HepG2", "pool1_pMPRA1_K562", "pool1_pNoCMVMPRA1_HeLa",
             "pool1_pNoCMVMPRA1_HepG2", "pool1_pNoCMVMPRA1_K562", "pool2_pMPRA1_HepG2", "pool2_pMPRA1_K562"]

all_dfs = [pool1_pMPRA1_HeLa_data_filt, pool1_pMPRA1_HepG2_data_filt, pool1_pMPRA1_K562_data_filt,
           pool1_pNoCMVMPRA1_HeLa_data_filt, pool1_pNoCMVMPRA1_HepG2_data_filt, pool1_pNoCMVMPRA1_K562_data_filt,
           pool2_pMPRA1_HepG2_data_filt, pool2_pMPRA1_K562_data_filt]

all_cols = [pool1_pMPRA1_HeLa_cols, pool1_pMPRA1_HepG2_cols, pool1_pMPRA1_K562_cols,
            pool1_pNoCMVMPRA1_HeLa_cols, pool1_pNoCMVMPRA1_HepG2_cols, pool1_pNoCMVMPRA1_K562_cols,
            pool2_pMPRA1_HepG2_cols, pool2_pMPRA1_K562_cols]

print("FILTERING RESULTS:")
for n, df, cs in zip(all_names, all_dfs, all_cols):
    if "pool1" in n:
        index_len = len(pool1_index)
    elif "pool2" in n:
        index_len = len(pool2_index)
        
    dna_barc_len = len(df)
    dna_barc_perc = (float(dna_barc_len)/index_len)*100
    
    print("%s: from %s barcodes to %s at DNA level (%s%%)" % (n, index_len, dna_barc_len, dna_barc_perc))
    
    reps = [x for x in cs if "rna_" in x]
    
    for r in reps:
        rep = r.split("_")[1]
        
        rna_barc_len = sum(~pd.isnull(df[r]))
        rna_barc_perc = (float(rna_barc_len)/index_len)*100
        
        print("\trep %s: %s barcodes at RNA level (%s%%)" % (rep, rna_barc_len, rna_barc_perc))
    print("")


# ## 5. join with index

# In[35]:


pool1_pMPRA1_HeLa_data_filt = pool1_pMPRA1_HeLa_data_filt.merge(pool1_index, on="barcode", how="inner")
pool1_pMPRA1_HepG2_data_filt = pool1_pMPRA1_HepG2_data_filt.merge(pool1_index, on="barcode", how="inner")
pool1_pMPRA1_K562_data_filt = pool1_pMPRA1_K562_data_filt.merge(pool1_index, on="barcode", how="inner")


# In[36]:


pool1_pNoCMVMPRA1_HeLa_data_filt = pool1_pNoCMVMPRA1_HeLa_data_filt.merge(pool1_index, on="barcode", how="inner")
pool1_pNoCMVMPRA1_HepG2_data_filt = pool1_pNoCMVMPRA1_HepG2_data_filt.merge(pool1_index, on="barcode", how="inner")
pool1_pNoCMVMPRA1_K562_data_filt = pool1_pNoCMVMPRA1_K562_data_filt.merge(pool1_index, on="barcode", how="inner")


# In[37]:


pool2_pMPRA1_HepG2_data_filt = pool2_pMPRA1_HepG2_data_filt.merge(pool2_index, on="barcode", how="inner")
pool2_pMPRA1_K562_data_filt = pool2_pMPRA1_K562_data_filt.merge(pool2_index, on="barcode", how="inner")


# ## 6. filter elements (remove ones with < 3 barcodes represented at dna level)

# In[38]:


pool1_pMPRA1_HeLa_barcodes_per_elem = pool1_pMPRA1_HeLa_data_filt.groupby(["unique_id", "oligo_type"])["barcode"].agg("count").reset_index()
pool1_pMPRA1_HeLa_barcodes_per_elem_neg = pool1_pMPRA1_HeLa_barcodes_per_elem[pool1_pMPRA1_HeLa_barcodes_per_elem["oligo_type"].isin(["RANDOM", "SCRAMBLED"])]
pool1_pMPRA1_HeLa_barcodes_per_elem_no_neg = pool1_pMPRA1_HeLa_barcodes_per_elem[~pool1_pMPRA1_HeLa_barcodes_per_elem["oligo_type"].isin(["RANDOM", "SCRAMBLED"])]

pool1_pMPRA1_HeLa_barcodes_per_elem_no_neg_filt = pool1_pMPRA1_HeLa_barcodes_per_elem_no_neg[pool1_pMPRA1_HeLa_barcodes_per_elem_no_neg["barcode"] >= n_barcodes_per_elem_threshold]
pool1_pMPRA1_HeLa_total_elems_rep = len(pool1_pMPRA1_HeLa_barcodes_per_elem_no_neg)
pool1_pMPRA1_HeLa_total_elems_filt_rep = len(pool1_pMPRA1_HeLa_barcodes_per_elem_no_neg_filt)


# In[39]:


pool1_pMPRA1_HepG2_barcodes_per_elem = pool1_pMPRA1_HepG2_data_filt.groupby(["unique_id", "oligo_type"])["barcode"].agg("count").reset_index()
pool1_pMPRA1_HepG2_barcodes_per_elem_neg = pool1_pMPRA1_HepG2_barcodes_per_elem[pool1_pMPRA1_HepG2_barcodes_per_elem["oligo_type"].isin(["RANDOM", "SCRAMBLED"])]
pool1_pMPRA1_HepG2_barcodes_per_elem_no_neg = pool1_pMPRA1_HepG2_barcodes_per_elem[~pool1_pMPRA1_HepG2_barcodes_per_elem["oligo_type"].isin(["RANDOM", "SCRAMBLED"])]

pool1_pMPRA1_HepG2_barcodes_per_elem_no_neg_filt = pool1_pMPRA1_HepG2_barcodes_per_elem_no_neg[pool1_pMPRA1_HepG2_barcodes_per_elem_no_neg["barcode"] >= n_barcodes_per_elem_threshold]
pool1_pMPRA1_HepG2_total_elems_rep = len(pool1_pMPRA1_HepG2_barcodes_per_elem_no_neg)
pool1_pMPRA1_HepG2_total_elems_filt_rep = len(pool1_pMPRA1_HepG2_barcodes_per_elem_no_neg_filt)


# In[40]:


pool1_pMPRA1_K562_barcodes_per_elem = pool1_pMPRA1_K562_data_filt.groupby(["unique_id", "oligo_type"])["barcode"].agg("count").reset_index()
pool1_pMPRA1_K562_barcodes_per_elem_neg = pool1_pMPRA1_K562_barcodes_per_elem[pool1_pMPRA1_K562_barcodes_per_elem["oligo_type"].isin(["RANDOM", "SCRAMBLED"])]
pool1_pMPRA1_K562_barcodes_per_elem_no_neg = pool1_pMPRA1_K562_barcodes_per_elem[~pool1_pMPRA1_K562_barcodes_per_elem["oligo_type"].isin(["RANDOM", "SCRAMBLED"])]

pool1_pMPRA1_K562_barcodes_per_elem_no_neg_filt = pool1_pMPRA1_K562_barcodes_per_elem_no_neg[pool1_pMPRA1_K562_barcodes_per_elem_no_neg["barcode"] >= n_barcodes_per_elem_threshold]
pool1_pMPRA1_K562_total_elems_rep = len(pool1_pMPRA1_K562_barcodes_per_elem_no_neg)
pool1_pMPRA1_K562_total_elems_filt_rep = len(pool1_pMPRA1_K562_barcodes_per_elem_no_neg_filt)


# In[41]:


pool1_pNoCMVMPRA1_HeLa_barcodes_per_elem = pool1_pNoCMVMPRA1_HeLa_data_filt.groupby(["unique_id", "oligo_type"])["barcode"].agg("count").reset_index()
pool1_pNoCMVMPRA1_HeLa_barcodes_per_elem_neg = pool1_pNoCMVMPRA1_HeLa_barcodes_per_elem[pool1_pNoCMVMPRA1_HeLa_barcodes_per_elem["oligo_type"].isin(["RANDOM", "SCRAMBLED"])]
pool1_pNoCMVMPRA1_HeLa_barcodes_per_elem_no_neg = pool1_pNoCMVMPRA1_HeLa_barcodes_per_elem[~pool1_pNoCMVMPRA1_HeLa_barcodes_per_elem["oligo_type"].isin(["RANDOM", "SCRAMBLED"])]

pool1_pNoCMVMPRA1_HeLa_barcodes_per_elem_no_neg_filt = pool1_pNoCMVMPRA1_HeLa_barcodes_per_elem_no_neg[pool1_pNoCMVMPRA1_HeLa_barcodes_per_elem_no_neg["barcode"] >= n_barcodes_per_elem_threshold]
pool1_pNoCMVMPRA1_HeLa_total_elems_rep = len(pool1_pNoCMVMPRA1_HeLa_barcodes_per_elem_no_neg)
pool1_pNoCMVMPRA1_HeLa_total_elems_filt_rep = len(pool1_pNoCMVMPRA1_HeLa_barcodes_per_elem_no_neg_filt)


# In[42]:


pool1_pNoCMVMPRA1_HepG2_barcodes_per_elem = pool1_pNoCMVMPRA1_HepG2_data_filt.groupby(["unique_id", "oligo_type"])["barcode"].agg("count").reset_index()
pool1_pNoCMVMPRA1_HepG2_barcodes_per_elem_neg = pool1_pNoCMVMPRA1_HepG2_barcodes_per_elem[pool1_pNoCMVMPRA1_HepG2_barcodes_per_elem["oligo_type"].isin(["RANDOM", "SCRAMBLED"])]
pool1_pNoCMVMPRA1_HepG2_barcodes_per_elem_no_neg = pool1_pNoCMVMPRA1_HepG2_barcodes_per_elem[~pool1_pNoCMVMPRA1_HepG2_barcodes_per_elem["oligo_type"].isin(["RANDOM", "SCRAMBLED"])]

pool1_pNoCMVMPRA1_HepG2_barcodes_per_elem_no_neg_filt = pool1_pNoCMVMPRA1_HepG2_barcodes_per_elem_no_neg[pool1_pNoCMVMPRA1_HepG2_barcodes_per_elem_no_neg["barcode"] >= n_barcodes_per_elem_threshold]
pool1_pNoCMVMPRA1_HepG2_total_elems_rep = len(pool1_pNoCMVMPRA1_HepG2_barcodes_per_elem_no_neg)
pool1_pNoCMVMPRA1_HepG2_total_elems_filt_rep = len(pool1_pNoCMVMPRA1_HepG2_barcodes_per_elem_no_neg_filt)


# In[43]:


pool1_pNoCMVMPRA1_K562_barcodes_per_elem = pool1_pNoCMVMPRA1_K562_data_filt.groupby(["unique_id", "oligo_type"])["barcode"].agg("count").reset_index()
pool1_pNoCMVMPRA1_K562_barcodes_per_elem_neg = pool1_pNoCMVMPRA1_K562_barcodes_per_elem[pool1_pNoCMVMPRA1_K562_barcodes_per_elem["oligo_type"].isin(["RANDOM", "SCRAMBLED"])]
pool1_pNoCMVMPRA1_K562_barcodes_per_elem_no_neg = pool1_pNoCMVMPRA1_K562_barcodes_per_elem[~pool1_pNoCMVMPRA1_K562_barcodes_per_elem["oligo_type"].isin(["RANDOM", "SCRAMBLED"])]

pool1_pNoCMVMPRA1_K562_barcodes_per_elem_no_neg_filt = pool1_pNoCMVMPRA1_K562_barcodes_per_elem_no_neg[pool1_pNoCMVMPRA1_K562_barcodes_per_elem_no_neg["barcode"] >= n_barcodes_per_elem_threshold]
pool1_pNoCMVMPRA1_K562_total_elems_rep = len(pool1_pNoCMVMPRA1_K562_barcodes_per_elem_no_neg)
pool1_pNoCMVMPRA1_K562_total_elems_filt_rep = len(pool1_pNoCMVMPRA1_K562_barcodes_per_elem_no_neg_filt)


# In[44]:


pool2_pMPRA1_HepG2_barcodes_per_elem = pool2_pMPRA1_HepG2_data_filt.groupby(["unique_id", "oligo_type"])["barcode"].agg("count").reset_index()
pool2_pMPRA1_HepG2_barcodes_per_elem_neg = pool2_pMPRA1_HepG2_barcodes_per_elem[pool2_pMPRA1_HepG2_barcodes_per_elem["oligo_type"].isin(["RANDOM", "SCRAMBLED"])]
pool2_pMPRA1_HepG2_barcodes_per_elem_no_neg = pool2_pMPRA1_HepG2_barcodes_per_elem[~pool2_pMPRA1_HepG2_barcodes_per_elem["oligo_type"].isin(["RANDOM", "SCRAMBLED"])]

pool2_pMPRA1_HepG2_barcodes_per_elem_no_neg_filt = pool2_pMPRA1_HepG2_barcodes_per_elem_no_neg[pool2_pMPRA1_HepG2_barcodes_per_elem_no_neg["barcode"] >= n_barcodes_per_elem_threshold]
pool2_pMPRA1_HepG2_total_elems_rep = len(pool2_pMPRA1_HepG2_barcodes_per_elem_no_neg)
pool2_pMPRA1_HepG2_total_elems_filt_rep = len(pool2_pMPRA1_HepG2_barcodes_per_elem_no_neg_filt)


# In[45]:


pool2_pMPRA1_K562_barcodes_per_elem = pool2_pMPRA1_K562_data_filt.groupby(["unique_id", "oligo_type"])["barcode"].agg("count").reset_index()
pool2_pMPRA1_K562_barcodes_per_elem_neg = pool2_pMPRA1_K562_barcodes_per_elem[pool2_pMPRA1_K562_barcodes_per_elem["oligo_type"].isin(["RANDOM", "SCRAMBLED"])]
pool2_pMPRA1_K562_barcodes_per_elem_no_neg = pool2_pMPRA1_K562_barcodes_per_elem[~pool2_pMPRA1_K562_barcodes_per_elem["oligo_type"].isin(["RANDOM", "SCRAMBLED"])]

pool2_pMPRA1_K562_barcodes_per_elem_no_neg_filt = pool2_pMPRA1_K562_barcodes_per_elem_no_neg[pool2_pMPRA1_K562_barcodes_per_elem_no_neg["barcode"] >= n_barcodes_per_elem_threshold]
pool2_pMPRA1_K562_total_elems_rep = len(pool2_pMPRA1_K562_barcodes_per_elem_no_neg)
pool2_pMPRA1_K562_total_elems_filt_rep = len(pool2_pMPRA1_K562_barcodes_per_elem_no_neg_filt)


# In[46]:


print("ELEMENT FILTERING RESULTS:")
print("Pool1 pMPRA1 HeLa: filtered %s elements to %s represented at >= %s barcodes (%s%%)" % (pool1_pMPRA1_HeLa_total_elems_rep, pool1_pMPRA1_HeLa_total_elems_filt_rep,
                                                                                              n_barcodes_per_elem_threshold,
                                                                                              float(pool1_pMPRA1_HeLa_total_elems_filt_rep)/pool1_pMPRA1_HeLa_total_elems_rep*100))
print("Pool1 pMPRA1 HepG2: filtered %s elements to %s represented at >= %s barcodes (%s%%)" % (pool1_pMPRA1_HepG2_total_elems_rep, pool1_pMPRA1_HepG2_total_elems_filt_rep,
                                                                                               n_barcodes_per_elem_threshold,
                                                                                               float(pool1_pMPRA1_HepG2_total_elems_filt_rep)/pool1_pMPRA1_HepG2_total_elems_rep*100))
print("Pool1 pMPRA1 K562: filtered %s elements to %s represented at >= %s barcodes (%s%%)" % (pool1_pMPRA1_K562_total_elems_rep, pool1_pMPRA1_K562_total_elems_filt_rep,
                                                                                              n_barcodes_per_elem_threshold,
                                                                                              float(pool1_pMPRA1_K562_total_elems_filt_rep)/pool1_pMPRA1_K562_total_elems_rep*100))

print("")
print("Pool1 pNoCMVMPRA1 HeLa: filtered %s elements to %s represented at >= %s barcodes (%s%%)" % (pool1_pNoCMVMPRA1_HeLa_total_elems_rep, pool1_pNoCMVMPRA1_HeLa_total_elems_filt_rep,
                                                                                                   n_barcodes_per_elem_threshold,
                                                                                                   float(pool1_pNoCMVMPRA1_HeLa_total_elems_filt_rep)/pool1_pNoCMVMPRA1_HeLa_total_elems_rep*100))
print("Pool1 pNoCMVMPRA1 HepG2: filtered %s elements to %s represented at >= %s barcodes (%s%%)" % (pool1_pNoCMVMPRA1_HepG2_total_elems_rep, pool1_pNoCMVMPRA1_HepG2_total_elems_filt_rep,
                                                                                                    n_barcodes_per_elem_threshold,
                                                                                                    float(pool1_pNoCMVMPRA1_HepG2_total_elems_filt_rep)/pool1_pNoCMVMPRA1_HepG2_total_elems_rep*100))
print("Pool1 pNoCMVMPRA1 K562: filtered %s elements to %s represented at >= %s barcodes (%s%%)" % (pool1_pNoCMVMPRA1_K562_total_elems_rep, pool1_pNoCMVMPRA1_K562_total_elems_filt_rep,
                                                                                                   n_barcodes_per_elem_threshold,
                                                                                                   float(pool1_pNoCMVMPRA1_K562_total_elems_filt_rep)/pool1_pNoCMVMPRA1_K562_total_elems_rep*100))

print("")
print("Pool2 pMPRA1 HepG2: filtered %s elements to %s represented at >= %s barcodes (%s%%)" % (pool2_pMPRA1_HepG2_total_elems_rep, pool2_pMPRA1_HepG2_total_elems_filt_rep,
                                                                                               n_barcodes_per_elem_threshold,
                                                                                               float(pool2_pMPRA1_HepG2_total_elems_filt_rep)/pool2_pMPRA1_HepG2_total_elems_rep*100))
print("Pool2 pMPRA1 K562: filtered %s elements to %s represented at >= %s barcodes (%s%%)" % (pool2_pMPRA1_K562_total_elems_rep, pool2_pMPRA1_K562_total_elems_filt_rep,
                                                                                              n_barcodes_per_elem_threshold,
                                                                                              float(pool2_pMPRA1_K562_total_elems_filt_rep)/pool2_pMPRA1_K562_total_elems_rep*100))


# In[47]:


pool1_pMPRA1_good_elems = list(pool1_pMPRA1_HeLa_barcodes_per_elem_no_neg_filt["unique_id"]) + list(pool1_pMPRA1_HeLa_barcodes_per_elem_neg["unique_id"])
pool1_pNoCMVMPRA1_good_elems = list(pool1_pNoCMVMPRA1_HeLa_barcodes_per_elem_no_neg_filt["unique_id"]) + list(pool1_pNoCMVMPRA1_HeLa_barcodes_per_elem_neg["unique_id"])
pool2_pMPRA1_good_elems = list(pool2_pMPRA1_HepG2_barcodes_per_elem_no_neg_filt["unique_id"]) + list(pool2_pMPRA1_HepG2_barcodes_per_elem_neg["unique_id"])


# In[48]:


pool1_pMPRA1_HeLa_data_filt = pool1_pMPRA1_HeLa_data_filt[pool1_pMPRA1_HeLa_data_filt["unique_id"].isin(pool1_pMPRA1_good_elems)]
pool1_pMPRA1_HepG2_data_filt = pool1_pMPRA1_HepG2_data_filt[pool1_pMPRA1_HepG2_data_filt["unique_id"].isin(pool1_pMPRA1_good_elems)]
pool1_pMPRA1_K562_data_filt = pool1_pMPRA1_K562_data_filt[pool1_pMPRA1_K562_data_filt["unique_id"].isin(pool1_pMPRA1_good_elems)]

pool1_pNoCMVMPRA1_HeLa_data_filt = pool1_pNoCMVMPRA1_HeLa_data_filt[pool1_pNoCMVMPRA1_HeLa_data_filt["unique_id"].isin(pool1_pNoCMVMPRA1_good_elems)]
pool1_pNoCMVMPRA1_HepG2_data_filt = pool1_pNoCMVMPRA1_HepG2_data_filt[pool1_pNoCMVMPRA1_HepG2_data_filt["unique_id"].isin(pool1_pNoCMVMPRA1_good_elems)]
pool1_pNoCMVMPRA1_K562_data_filt = pool1_pNoCMVMPRA1_K562_data_filt[pool1_pNoCMVMPRA1_K562_data_filt["unique_id"].isin(pool1_pNoCMVMPRA1_good_elems)]

pool2_pMPRA1_HepG2_data_filt = pool2_pMPRA1_HepG2_data_filt[pool2_pMPRA1_HepG2_data_filt["unique_id"].isin(pool2_pMPRA1_good_elems)]
pool2_pMPRA1_K562_data_filt = pool2_pMPRA1_K562_data_filt[pool2_pMPRA1_K562_data_filt["unique_id"].isin(pool2_pMPRA1_good_elems)]


# ## 5. write final file

# In[49]:


pool1_pMPRA1_HeLa_counts = pool1_pMPRA1_HeLa_data_filt[pool1_pMPRA1_HeLa_cols]
pool1_pMPRA1_HepG2_counts = pool1_pMPRA1_HepG2_data_filt[pool1_pMPRA1_HepG2_cols]
pool1_pMPRA1_K562_counts = pool1_pMPRA1_K562_data_filt[pool1_pMPRA1_K562_cols]

pool1_pNoCMVMPRA1_HeLa_counts = pool1_pNoCMVMPRA1_HeLa_data_filt[pool1_pNoCMVMPRA1_HeLa_cols]
pool1_pNoCMVMPRA1_HepG2_counts = pool1_pNoCMVMPRA1_HepG2_data_filt[pool1_pNoCMVMPRA1_HepG2_cols]
pool1_pNoCMVMPRA1_K562_counts = pool1_pNoCMVMPRA1_K562_data_filt[pool1_pNoCMVMPRA1_K562_cols]

pool2_pMPRA1_HepG2_counts = pool2_pMPRA1_HepG2_data_filt[pool2_pMPRA1_HepG2_cols]
pool2_pMPRA1_K562_counts = pool2_pMPRA1_K562_data_filt[pool2_pMPRA1_K562_cols]

pool2_pMPRA1_HepG2_counts.head()


# In[50]:


pool1_pMPRA1_HeLa_counts.to_csv("%s/%s" % (counts_dir, pool1_pMPRA1_HeLa_out_f), sep="\t", header=True, index=False)
pool1_pMPRA1_HepG2_counts.to_csv("%s/%s" % (counts_dir, pool1_pMPRA1_HepG2_out_f), sep="\t", header=True, index=False)
pool1_pMPRA1_K562_counts.to_csv("%s/%s" % (counts_dir, pool1_pMPRA1_K562_out_f), sep="\t", header=True, index=False)

pool1_pNoCMVMPRA1_HeLa_counts.to_csv("%s/%s" % (counts_dir, pool1_pNoCMVMPRA1_HeLa_out_f), sep="\t", header=True, index=False)
pool1_pNoCMVMPRA1_HepG2_counts.to_csv("%s/%s" % (counts_dir, pool1_pNoCMVMPRA1_HepG2_out_f), sep="\t", header=True, index=False)
pool1_pNoCMVMPRA1_K562_counts.to_csv("%s/%s" % (counts_dir, pool1_pNoCMVMPRA1_K562_out_f), sep="\t", header=True, index=False)

pool2_pMPRA1_HepG2_counts.to_csv("%s/%s" % (counts_dir, pool2_pMPRA1_HepG2_out_f), sep="\t", header=True, index=False)
pool2_pMPRA1_K562_counts.to_csv("%s/%s" % (counts_dir, pool2_pMPRA1_K562_out_f), sep="\t", header=True, index=False)


# ## 6. heatmap comparing barcode counts

# In[51]:


pool1_pMPRA1_HeLa_cols = ["barcode"]
pool1_pMPRA1_HepG2_cols = ["barcode"]
pool1_pMPRA1_K562_cols = ["barcode"]
pool1_pNoCMVMPRA1_HeLa_cols = ["barcode"]
pool1_pNoCMVMPRA1_HepG2_cols = ["barcode"]
pool1_pNoCMVMPRA1_K562_cols = ["barcode"]


# In[52]:


pool1_pMPRA1_HeLa_cols.extend(["HeLa_%s" % x for x in pool1_pMPRA1_HeLa_reps])
pool1_pMPRA1_HepG2_cols.extend(["HepG2_%s" % x for x in pool1_pMPRA1_HepG2_reps])
pool1_pMPRA1_K562_cols.extend(["K562_%s" % x for x in pool1_pMPRA1_K562_reps])
pool1_pNoCMVMPRA1_HeLa_cols.extend(["HeLa_%s (no min. prom.)" % x for x in pool1_pNoCMVMPRA1_HeLa_reps])
pool1_pNoCMVMPRA1_HepG2_cols.extend(["HepG2_%s (no min. prom.)" % x for x in pool1_pNoCMVMPRA1_HepG2_reps])
pool1_pNoCMVMPRA1_K562_cols.extend(["K562_%s (no min. prom.)" % x for x in pool1_pNoCMVMPRA1_K562_reps])
pool1_pNoCMVMPRA1_K562_cols


# In[53]:


pool1_pMPRA1_HeLa_counts.drop("dna_1", axis=1, inplace=True)
pool1_pMPRA1_HepG2_counts.drop("dna_1", axis=1, inplace=True)
pool1_pMPRA1_K562_counts.drop("dna_1", axis=1, inplace=True)
pool1_pNoCMVMPRA1_HeLa_counts.drop("dna_1", axis=1, inplace=True)
pool1_pNoCMVMPRA1_HepG2_counts.drop("dna_1", axis=1, inplace=True)
pool1_pNoCMVMPRA1_K562_counts.drop("dna_1", axis=1, inplace=True)


# In[54]:


pool1_pMPRA1_HeLa_counts.columns = pool1_pMPRA1_HeLa_cols
pool1_pMPRA1_HepG2_counts.columns = pool1_pMPRA1_HepG2_cols
pool1_pMPRA1_K562_counts.columns = pool1_pMPRA1_K562_cols
pool1_pNoCMVMPRA1_HeLa_counts.columns = pool1_pNoCMVMPRA1_HeLa_cols
pool1_pNoCMVMPRA1_HepG2_counts.columns = pool1_pNoCMVMPRA1_HepG2_cols
pool1_pNoCMVMPRA1_K562_counts.columns = pool1_pNoCMVMPRA1_K562_cols


# In[55]:


all_pool1 = pool1_pMPRA1_HeLa_counts.merge(pool1_pMPRA1_HepG2_counts, on="barcode").merge(pool1_pMPRA1_K562_counts, on = "barcode").merge(pool1_pNoCMVMPRA1_HeLa_counts, on="barcode").merge(pool1_pNoCMVMPRA1_HepG2_counts, on="barcode").merge(pool1_pNoCMVMPRA1_K562_counts, on="barcode")
all_pool1_corr = all_pool1.corr(method="pearson")


# In[56]:


cmap = sns.cubehelix_palette(as_cmap=True)
cg = sns.clustermap(all_pool1_corr, figsize=(7.2,7.2), cmap=cmap, annot=False)
_ = plt.setp(cg.ax_heatmap.yaxis.get_majorticklabels(), rotation=0)
plt.suptitle("pearson correlation of replicates\nall barcodes at counts > 5")
cg.savefig("Fig_S3A.pdf", dpi="figure", transparent=True, bbox_inches="tight")
```
