## Supplementary material for "High-throughput functional analysis of IncRNA core promoters elucidatesrules governing tissue-specificity": 02__normalize.py

```
# coding: utf-8

# # 02__normalize
# # normalizing data (log2, median center, quantile norm)
# 
# in this notebook, i take all the filtered RNA and DNA barcode counts and calculate activities both per barcode and per element. the steps are as follows:
# 1. add a pseudocount of 1 to every barcode
# 2. normalize each replicate for sequencing depth by converting counts to counts per million (CPMs)
# 3. calculate activity as RNA/DNA
# 4. transform activities to log2
# 5. median center each replicate
# 6. quantile-normalize the replicates within a given condition
# 
# to calculate activities per element, i take the median activity value across corresponding barcodes. 
# 
# ------
# 
# figures in this notebook:
# - **Fig S3B**: heatmap of element activity correlation between replicates

# In[1]:


import warnings
warnings.filterwarnings('ignore')

import pandas as pd
import numpy as np
import matplotlib as mpl
import matplotlib.pyplot as plt
import seaborn as sns
import sys

# import utils
sys.path.append("../../utils")
from plotting_utils import *
from misc_utils import *
from norm_utils import *

get_ipython().magic('matplotlib inline')


# In[2]:


sns.set(**PAPER_PRESET)
fontsize = PAPER_FONTSIZE


# ## variables

# In[3]:


counts_dir = "../../data/01__counts"
out_dir = "../../data/02__activs"
get_ipython().system('mkdir -p $out_dir')


# In[4]:


pool1_pMPRA1_HeLa_barc_counts_f = "POOL1__pMPRA1__HeLa__all_counts.txt"
pool1_pMPRA1_HepG2_barc_counts_f = "POOL1__pMPRA1__HepG2__all_counts.txt"
pool1_pMPRA1_K562_barc_counts_f = "POOL1__pMPRA1__K562__all_counts.txt"

pool1_pNoCMVMPRA1_HeLa_barc_counts_f = "POOL1__pNoCMVMPRA1__HeLa__all_counts.txt"
pool1_pNoCMVMPRA1_HepG2_barc_counts_f = "POOL1__pNoCMVMPRA1__HepG2__all_counts.txt"
pool1_pNoCMVMPRA1_K562_barc_counts_f = "POOL1__pNoCMVMPRA1__K562__all_counts.txt"

pool2_pMPRA1_HepG2_barc_counts_f = "POOL2__pMPRA1__HepG2__all_counts.txt"
pool2_pMPRA1_K562_barc_counts_f = "POOL2__pMPRA1__K562__all_counts.txt"


# In[5]:


pool1_index_f = "../../data/00__index/tss_oligo_pool.index.txt"
pool2_index_f = "../../data/00__index/dels_oligo_pool.index.txt"


# In[6]:


pool1_pMPRA1_HeLa_activ_elem_f = "%s/POOL1__pMPRA1__HeLa__activities_per_element.txt" % out_dir
pool1_pMPRA1_HeLa_activ_barc_f = "%s/POOL1__pMPRA1__HeLa__activities_per_barcode.txt" % out_dir

pool1_pMPRA1_HepG2_activ_elem_f = "%s/POOL1__pMPRA1__HepG2__activities_per_element.txt" % out_dir
pool1_pMPRA1_HepG2_activ_barc_f = "%s/POOL1__pMPRA1__HepG2__activities_per_barcode.txt" % out_dir

pool1_pMPRA1_K562_activ_elem_f = "%s/POOL1__pMPRA1__K562__activities_per_element.txt" % out_dir
pool1_pMPRA1_K562_activ_barc_f = "%s/POOL1__pMPRA1__K562__activities_per_barcode.txt" % out_dir

pool1_pNoCMVMPRA1_HeLa_activ_elem_f = "%s/POOL1__pNoCMVMPRA1__HeLa__activities_per_element.txt" % out_dir
pool1_pNoCMVMPRA1_HeLa_activ_barc_f = "%s/POOL1__pNoCMVMPRA1__HeLa__activities_per_barcode.txt" % out_dir

pool1_pNoCMVMPRA1_HepG2_activ_elem_f = "%s/POOL1__pNoCMVMPRA1__HepG2__activities_per_element.txt" % out_dir
pool1_pNoCMVMPRA1_HepG2_activ_barc_f = "%s/POOL1__pNoCMVMPRA1__HepG2__activities_per_barcode.txt" % out_dir

pool1_pNoCMVMPRA1_K562_activ_elem_f = "%s/POOL1__pNoCMVMPRA1__K562__activities_per_element.txt" % out_dir
pool1_pNoCMVMPRA1_K562_activ_barc_f = "%s/POOL1__pNoCMVMPRA1__K562__activities_per_barcode.txt" % out_dir

pool2_pMPRA1_HepG2_activ_elem_f = "%s/POOL2__pMPRA1__HepG2__activities_per_element.txt" % out_dir
pool2_pMPRA1_HepG2_activ_barc_f = "%s/POOL2__pMPRA1__HepG2__activities_per_barcode.txt" % out_dir

pool2_pMPRA1_K562_activ_elem_f = "%s/POOL2__pMPRA1__K562__activities_per_element.txt" % out_dir
pool2_pMPRA1_K562_activ_barc_f = "%s/POOL2__pMPRA1__K562__activities_per_barcode.txt" % out_dir


# ## 1. import data

# In[7]:


pool1_pMPRA1_HeLa_barc_counts = pd.read_table("%s/%s" % (counts_dir, pool1_pMPRA1_HeLa_barc_counts_f), sep="\t")
pool1_pMPRA1_HepG2_barc_counts = pd.read_table("%s/%s" % (counts_dir, pool1_pMPRA1_HepG2_barc_counts_f), sep="\t")
pool1_pMPRA1_K562_barc_counts = pd.read_table("%s/%s" % (counts_dir, pool1_pMPRA1_K562_barc_counts_f), sep="\t")

pool1_pNoCMVMPRA1_HeLa_barc_counts = pd.read_table("%s/%s" % (counts_dir, pool1_pNoCMVMPRA1_HeLa_barc_counts_f), sep="\t")
pool1_pNoCMVMPRA1_HepG2_barc_counts = pd.read_table("%s/%s" % (counts_dir, pool1_pNoCMVMPRA1_HepG2_barc_counts_f), sep="\t")
pool1_pNoCMVMPRA1_K562_barc_counts = pd.read_table("%s/%s" % (counts_dir, pool1_pNoCMVMPRA1_K562_barc_counts_f), sep="\t")

pool2_pMPRA1_HepG2_barc_counts = pd.read_table("%s/%s" % (counts_dir, pool2_pMPRA1_HepG2_barc_counts_f), sep="\t")
pool2_pMPRA1_K562_barc_counts = pd.read_table("%s/%s" % (counts_dir, pool2_pMPRA1_K562_barc_counts_f), sep="\t")

pool1_pMPRA1_HeLa_barc_counts.head()


# In[8]:


print(len(pool1_pMPRA1_HeLa_barc_counts))
print(len(pool1_pMPRA1_HepG2_barc_counts))
print(len(pool1_pMPRA1_K562_barc_counts))

print(len(pool1_pNoCMVMPRA1_HeLa_barc_counts))
print(len(pool1_pNoCMVMPRA1_HepG2_barc_counts))
print(len(pool1_pNoCMVMPRA1_K562_barc_counts))

print(len(pool2_pMPRA1_HepG2_barc_counts))
print(len(pool2_pMPRA1_K562_barc_counts))


# In[9]:


pool1_index = pd.read_table(pool1_index_f, sep="\t")
pool2_index = pd.read_table(pool2_index_f, sep="\t")


# In[10]:


pool1_index_elem = pool1_index[["element", "oligo_type", "unique_id", "dupe_info", "SNP"]]
pool2_index_elem = pool2_index[["element", "oligo_type", "unique_id", "dupe_info", "SNP"]]

pool1_index_elem = pool1_index_elem.drop_duplicates()
pool2_index_elem = pool2_index_elem.drop_duplicates()


# ## 2. normalize

# In[11]:


pool1_pMPRA1_HeLa_barc_pseudo = pseudocount(pool1_pMPRA1_HeLa_barc_counts)
pool1_pMPRA1_HepG2_barc_pseudo = pseudocount(pool1_pMPRA1_HepG2_barc_counts)
pool1_pMPRA1_K562_barc_pseudo = pseudocount(pool1_pMPRA1_K562_barc_counts)

pool1_pNoCMVMPRA1_HeLa_barc_pseudo = pseudocount(pool1_pNoCMVMPRA1_HeLa_barc_counts)
pool1_pNoCMVMPRA1_HepG2_barc_pseudo = pseudocount(pool1_pNoCMVMPRA1_HepG2_barc_counts)
pool1_pNoCMVMPRA1_K562_barc_pseudo = pseudocount(pool1_pNoCMVMPRA1_K562_barc_counts)

pool2_pMPRA1_HepG2_barc_pseudo = pseudocount(pool2_pMPRA1_HepG2_barc_counts)
pool2_pMPRA1_K562_barc_pseudo = pseudocount(pool2_pMPRA1_K562_barc_counts)

pool1_pMPRA1_HeLa_barc_pseudo.head()


# In[12]:


pool1_pMPRA1_HeLa_barc_cpm = to_cpm(pool1_pMPRA1_HeLa_barc_pseudo)
pool1_pMPRA1_HepG2_barc_cpm = to_cpm(pool1_pMPRA1_HepG2_barc_pseudo)
pool1_pMPRA1_K562_barc_cpm = to_cpm(pool1_pMPRA1_K562_barc_pseudo)

pool1_pNoCMVMPRA1_HeLa_barc_cpm = to_cpm(pool1_pNoCMVMPRA1_HeLa_barc_pseudo)
pool1_pNoCMVMPRA1_HepG2_barc_cpm = to_cpm(pool1_pNoCMVMPRA1_HepG2_barc_pseudo)
pool1_pNoCMVMPRA1_K562_barc_cpm = to_cpm(pool1_pNoCMVMPRA1_K562_barc_pseudo)

pool2_pMPRA1_HepG2_barc_cpm = to_cpm(pool2_pMPRA1_HepG2_barc_pseudo)
pool2_pMPRA1_K562_barc_cpm = to_cpm(pool2_pMPRA1_K562_barc_pseudo)

pool1_pMPRA1_HeLa_barc_cpm.head()


# In[13]:


pool1_pMPRA1_HeLa_barc_activ = to_activ(pool1_pMPRA1_HeLa_barc_cpm)
pool1_pMPRA1_HepG2_barc_activ = to_activ(pool1_pMPRA1_HepG2_barc_cpm)
pool1_pMPRA1_K562_barc_activ = to_activ(pool1_pMPRA1_K562_barc_cpm)

pool1_pNoCMVMPRA1_HeLa_barc_activ = to_activ(pool1_pNoCMVMPRA1_HeLa_barc_cpm)
pool1_pNoCMVMPRA1_HepG2_barc_activ = to_activ(pool1_pNoCMVMPRA1_HepG2_barc_cpm)
pool1_pNoCMVMPRA1_K562_barc_activ = to_activ(pool1_pNoCMVMPRA1_K562_barc_cpm)

pool2_pMPRA1_HepG2_barc_activ = to_activ(pool2_pMPRA1_HepG2_barc_cpm)
pool2_pMPRA1_K562_barc_activ = to_activ(pool2_pMPRA1_K562_barc_cpm)

pool1_pMPRA1_HeLa_barc_activ.head()


# In[14]:


pool1_pMPRA1_HeLa_barc_log2 = to_log2(pool1_pMPRA1_HeLa_barc_activ)
pool1_pMPRA1_HepG2_barc_log2 = to_log2(pool1_pMPRA1_HepG2_barc_activ)
pool1_pMPRA1_K562_barc_log2 = to_log2(pool1_pMPRA1_K562_barc_activ)

pool1_pNoCMVMPRA1_HeLa_barc_log2 = to_log2(pool1_pNoCMVMPRA1_HeLa_barc_activ)
pool1_pNoCMVMPRA1_HepG2_barc_log2 = to_log2(pool1_pNoCMVMPRA1_HepG2_barc_activ)
pool1_pNoCMVMPRA1_K562_barc_log2 = to_log2(pool1_pNoCMVMPRA1_K562_barc_activ)

pool2_pMPRA1_HepG2_barc_log2 = to_log2(pool2_pMPRA1_HepG2_barc_activ)
pool2_pMPRA1_K562_barc_log2 = to_log2(pool2_pMPRA1_K562_barc_activ)

pool1_pMPRA1_HeLa_barc_log2.head()


# In[15]:


pool1_pMPRA1_HeLa_barc_norm = median_norm(pool1_pMPRA1_HeLa_barc_log2)
pool1_pMPRA1_HepG2_barc_norm = median_norm(pool1_pMPRA1_HepG2_barc_log2)
pool1_pMPRA1_K562_barc_norm = median_norm(pool1_pMPRA1_K562_barc_log2)

pool1_pNoCMVMPRA1_HeLa_barc_norm = median_norm(pool1_pNoCMVMPRA1_HeLa_barc_log2)
pool1_pNoCMVMPRA1_HepG2_barc_norm = median_norm(pool1_pNoCMVMPRA1_HepG2_barc_log2)
pool1_pNoCMVMPRA1_K562_barc_norm = median_norm(pool1_pNoCMVMPRA1_K562_barc_log2)

pool2_pMPRA1_HepG2_barc_norm = median_norm(pool2_pMPRA1_HepG2_barc_log2)
pool2_pMPRA1_K562_barc_norm = median_norm(pool2_pMPRA1_K562_barc_log2)

pool1_pMPRA1_HeLa_barc_norm.head()


# In[16]:


pool1_pMPRA1_HeLa_barc_quant = quantile_norm(pool1_pMPRA1_HeLa_barc_norm)
pool1_pMPRA1_HepG2_barc_quant = quantile_norm(pool1_pMPRA1_HepG2_barc_norm)
pool1_pMPRA1_K562_barc_quant = quantile_norm(pool1_pMPRA1_K562_barc_norm)

pool1_pNoCMVMPRA1_HeLa_barc_quant = quantile_norm(pool1_pNoCMVMPRA1_HeLa_barc_norm)
pool1_pNoCMVMPRA1_HepG2_barc_quant = quantile_norm(pool1_pNoCMVMPRA1_HepG2_barc_norm)
pool1_pNoCMVMPRA1_K562_barc_quant = quantile_norm(pool1_pNoCMVMPRA1_K562_barc_norm)

pool2_pMPRA1_HepG2_barc_quant = quantile_norm(pool2_pMPRA1_HepG2_barc_norm)
pool2_pMPRA1_K562_barc_quant = quantile_norm(pool2_pMPRA1_K562_barc_norm)

pool1_pMPRA1_HeLa_barc_quant.head()


# In[17]:


print(len(pool1_pMPRA1_HeLa_barc_quant))
print(len(pool1_pMPRA1_HepG2_barc_quant))
print(len(pool1_pMPRA1_K562_barc_quant))

print(len(pool1_pNoCMVMPRA1_HeLa_barc_quant))
print(len(pool1_pNoCMVMPRA1_HepG2_barc_quant))
print(len(pool1_pNoCMVMPRA1_K562_barc_quant))

print(len(pool2_pMPRA1_HepG2_barc_quant))
print(len(pool2_pMPRA1_K562_barc_quant))


# ## 3. join with index

# In[18]:


pool1_pMPRA1_HeLa_barc_quant = pool1_index.merge(pool1_pMPRA1_HeLa_barc_quant, left_on="barcode", right_on="barcode", how="left")
pool1_pMPRA1_HepG2_barc_quant = pool1_index.merge(pool1_pMPRA1_HepG2_barc_quant, left_on="barcode", right_on="barcode", how="left")
pool1_pMPRA1_K562_barc_quant = pool1_index.merge(pool1_pMPRA1_K562_barc_quant, left_on="barcode", right_on="barcode", how="left")

pool1_pNoCMVMPRA1_HeLa_barc_quant = pool1_index.merge(pool1_pNoCMVMPRA1_HeLa_barc_quant, left_on="barcode", right_on="barcode", how="left")
pool1_pNoCMVMPRA1_HepG2_barc_quant = pool1_index.merge(pool1_pNoCMVMPRA1_HepG2_barc_quant, left_on="barcode", right_on="barcode", how="left")
pool1_pNoCMVMPRA1_K562_barc_quant = pool1_index.merge(pool1_pNoCMVMPRA1_K562_barc_quant, left_on="barcode", right_on="barcode", how="left")

pool2_pMPRA1_HepG2_barc_quant = pool2_index.merge(pool2_pMPRA1_HepG2_barc_quant, left_on="barcode", right_on="barcode", how="left")
pool2_pMPRA1_K562_barc_quant = pool2_index.merge(pool2_pMPRA1_K562_barc_quant, left_on="barcode", right_on="barcode", how="left")

pool1_pMPRA1_HeLa_barc_quant.head()


# In[19]:


print(len(pool1_pMPRA1_HeLa_barc_quant))
print(len(pool1_pMPRA1_HepG2_barc_quant))
print(len(pool1_pMPRA1_K562_barc_quant))

print(len(pool1_pNoCMVMPRA1_HeLa_barc_quant))
print(len(pool1_pNoCMVMPRA1_HepG2_barc_quant))
print(len(pool1_pNoCMVMPRA1_K562_barc_quant))

print(len(pool2_pMPRA1_HepG2_barc_quant))
print(len(pool2_pMPRA1_K562_barc_quant))


# ## 4. collapse per element

# In[20]:


pool1_pMPRA1_HeLa_reps = [x for x in pool1_pMPRA1_HeLa_barc_quant.columns if "rna" in x]
pool1_pMPRA1_HepG2_reps = [x for x in pool1_pMPRA1_HepG2_barc_quant.columns if "rna" in x]
pool1_pMPRA1_K562_reps = [x for x in pool1_pMPRA1_K562_barc_quant.columns if "rna" in x]

pool1_pNoCMVMPRA1_HeLa_reps = [x for x in pool1_pNoCMVMPRA1_HeLa_barc_quant.columns if "rna" in x]
pool1_pNoCMVMPRA1_HepG2_reps = [x for x in pool1_pNoCMVMPRA1_HepG2_barc_quant.columns if "rna" in x]
pool1_pNoCMVMPRA1_K562_reps = [x for x in pool1_pNoCMVMPRA1_K562_barc_quant.columns if "rna" in x]

pool2_pMPRA1_HepG2_reps = [x for x in pool2_pMPRA1_HepG2_barc_quant.columns if "rna" in x]
pool2_pMPRA1_K562_reps = [x for x in pool2_pMPRA1_K562_barc_quant.columns if "rna" in x]


# In[21]:


pool1_pMPRA1_HeLa_barc_quant_grp = pool1_pMPRA1_HeLa_barc_quant.groupby(["element", "oligo_type", "unique_id"])[pool1_pMPRA1_HeLa_reps].agg("median").reset_index()
pool1_pMPRA1_HepG2_barc_quant_grp = pool1_pMPRA1_HepG2_barc_quant.groupby(["element", "oligo_type", "unique_id"])[pool1_pMPRA1_HepG2_reps].agg("median").reset_index()
pool1_pMPRA1_K562_barc_quant_grp = pool1_pMPRA1_K562_barc_quant.groupby(["element", "oligo_type", "unique_id"])[pool1_pMPRA1_K562_reps].agg("median").reset_index()

pool1_pNoCMVMPRA1_HeLa_barc_quant_grp = pool1_pNoCMVMPRA1_HeLa_barc_quant.groupby(["element", "oligo_type", "unique_id"])[pool1_pNoCMVMPRA1_HeLa_reps].agg("median").reset_index()
pool1_pNoCMVMPRA1_HepG2_barc_quant_grp = pool1_pNoCMVMPRA1_HepG2_barc_quant.groupby(["element", "oligo_type", "unique_id"])[pool1_pNoCMVMPRA1_HepG2_reps].agg("median").reset_index()
pool1_pNoCMVMPRA1_K562_barc_quant_grp = pool1_pNoCMVMPRA1_K562_barc_quant.groupby(["element", "oligo_type", "unique_id"])[pool1_pNoCMVMPRA1_K562_reps].agg("median").reset_index()

pool2_pMPRA1_HepG2_barc_quant_grp = pool2_pMPRA1_HepG2_barc_quant.groupby(["element", "oligo_type", "unique_id"])[pool2_pMPRA1_HepG2_reps].agg("median").reset_index()
pool2_pMPRA1_K562_barc_quant_grp = pool2_pMPRA1_K562_barc_quant.groupby(["element", "oligo_type", "unique_id"])[pool2_pMPRA1_K562_reps].agg("median").reset_index()


# ## 5. write files

# ### activities per element, column per replicate

# In[22]:


pool1_pMPRA1_HeLa_activs_per_elem = pool1_pMPRA1_HeLa_barc_quant.groupby(["unique_id", "element"])[pool1_pMPRA1_HeLa_reps].agg("median").reset_index()
pool1_pMPRA1_HepG2_activs_per_elem = pool1_pMPRA1_HepG2_barc_quant.groupby(["unique_id", "element"])[pool1_pMPRA1_HepG2_reps].agg("median").reset_index()
pool1_pMPRA1_K562_activs_per_elem = pool1_pMPRA1_K562_barc_quant.groupby(["unique_id", "element"])[pool1_pMPRA1_K562_reps].agg("median").reset_index()

pool1_pNoCMVMPRA1_HeLa_activs_per_elem = pool1_pNoCMVMPRA1_HeLa_barc_quant.groupby(["unique_id", "element"])[pool1_pNoCMVMPRA1_HeLa_reps].agg("median").reset_index()
pool1_pNoCMVMPRA1_HepG2_activs_per_elem = pool1_pNoCMVMPRA1_HepG2_barc_quant.groupby(["unique_id", "element"])[pool1_pNoCMVMPRA1_HepG2_reps].agg("median").reset_index()
pool1_pNoCMVMPRA1_K562_activs_per_elem = pool1_pNoCMVMPRA1_K562_barc_quant.groupby(["unique_id", "element"])[pool1_pNoCMVMPRA1_K562_reps].agg("median").reset_index()

pool2_pMPRA1_HepG2_activs_per_elem = pool2_pMPRA1_HepG2_barc_quant.groupby(["unique_id", "element"])[pool2_pMPRA1_HepG2_reps].agg("median").reset_index()
pool2_pMPRA1_K562_activs_per_elem = pool2_pMPRA1_K562_barc_quant.groupby(["unique_id", "element"])[pool2_pMPRA1_K562_reps].agg("median").reset_index()


# In[23]:


pool1_pMPRA1_HeLa_activs_per_elem.to_csv(pool1_pMPRA1_HeLa_activ_elem_f, sep="\t", header=True, index=False)
pool1_pMPRA1_HepG2_activs_per_elem.to_csv(pool1_pMPRA1_HepG2_activ_elem_f, sep="\t", header=True, index=False)
pool1_pMPRA1_K562_activs_per_elem.to_csv(pool1_pMPRA1_K562_activ_elem_f, sep="\t", header=True, index=False)

pool1_pNoCMVMPRA1_HeLa_activs_per_elem.to_csv(pool1_pNoCMVMPRA1_HeLa_activ_elem_f, sep="\t", header=True, index=False)
pool1_pNoCMVMPRA1_HepG2_activs_per_elem.to_csv(pool1_pNoCMVMPRA1_HepG2_activ_elem_f, sep="\t", header=True, index=False)
pool1_pNoCMVMPRA1_K562_activs_per_elem.to_csv(pool1_pNoCMVMPRA1_K562_activ_elem_f, sep="\t", header=True, index=False)

pool2_pMPRA1_HepG2_activs_per_elem.to_csv(pool2_pMPRA1_HepG2_activ_elem_f, sep="\t", header=True, index=False)
pool2_pMPRA1_K562_activs_per_elem.to_csv(pool2_pMPRA1_K562_activ_elem_f, sep="\t", header=True, index=False)


# ### activities per barcode, column per replicate

# In[24]:


pool1_pMPRA1_HeLa_activs_per_barc = pool1_pMPRA1_HeLa_barc_quant.groupby(["barcode"])[pool1_pMPRA1_HeLa_reps].agg("median").reset_index()
pool1_pMPRA1_HepG2_activs_per_barc = pool1_pMPRA1_HepG2_barc_quant.groupby(["barcode"])[pool1_pMPRA1_HepG2_reps].agg("median").reset_index()
pool1_pMPRA1_K562_activs_per_barc = pool1_pMPRA1_K562_barc_quant.groupby(["barcode"])[pool1_pMPRA1_K562_reps].agg("median").reset_index()

pool1_pNoCMVMPRA1_HeLa_activs_per_barc = pool1_pNoCMVMPRA1_HeLa_barc_quant.groupby(["barcode"])[pool1_pNoCMVMPRA1_HeLa_reps].agg("median").reset_index()
pool1_pNoCMVMPRA1_HepG2_activs_per_barc = pool1_pNoCMVMPRA1_HepG2_barc_quant.groupby(["barcode"])[pool1_pNoCMVMPRA1_HepG2_reps].agg("median").reset_index()
pool1_pNoCMVMPRA1_K562_activs_per_barc = pool1_pNoCMVMPRA1_K562_barc_quant.groupby(["barcode"])[pool1_pNoCMVMPRA1_K562_reps].agg("median").reset_index()

pool2_pMPRA1_HepG2_activs_per_barc = pool2_pMPRA1_HepG2_barc_quant.groupby(["barcode"])[pool2_pMPRA1_HepG2_reps].agg("median").reset_index()
pool2_pMPRA1_K562_activs_per_barc = pool2_pMPRA1_K562_barc_quant.groupby(["barcode"])[pool2_pMPRA1_K562_reps].agg("median").reset_index()


# In[25]:


pool1_pMPRA1_HeLa_activs_per_barc.to_csv(pool1_pMPRA1_HeLa_activ_barc_f, sep="\t", header=True, index=False)
pool1_pMPRA1_HepG2_activs_per_barc.to_csv(pool1_pMPRA1_HepG2_activ_barc_f, sep="\t", header=True, index=False)
pool1_pMPRA1_K562_activs_per_barc.to_csv(pool1_pMPRA1_K562_activ_barc_f, sep="\t", header=True, index=False)

pool1_pNoCMVMPRA1_HeLa_activs_per_barc.to_csv(pool1_pNoCMVMPRA1_HeLa_activ_barc_f, sep="\t", header=True, index=False)
pool1_pNoCMVMPRA1_HepG2_activs_per_barc.to_csv(pool1_pNoCMVMPRA1_HepG2_activ_barc_f, sep="\t", header=True, index=False)
pool1_pNoCMVMPRA1_K562_activs_per_barc.to_csv(pool1_pNoCMVMPRA1_K562_activ_barc_f, sep="\t", header=True, index=False)

pool2_pMPRA1_HepG2_activs_per_barc.to_csv(pool2_pMPRA1_HepG2_activ_barc_f, sep="\t", header=True, index=False)
pool2_pMPRA1_K562_activs_per_barc.to_csv(pool2_pMPRA1_K562_activ_barc_f, sep="\t", header=True, index=False)


# ## 6. plot replicate heatmaps

# ### pool 1 (TSS pool)

# In[26]:


pool1_pMPRA1_HeLa_activs_per_elem.columns = ["HeLa_%s" % x if "rna" in x else x for x in pool1_pMPRA1_HeLa_activs_per_elem.columns]
pool1_pMPRA1_HepG2_activs_per_elem.columns = ["HepG2_%s" % x if "rna" in x else x for x in pool1_pMPRA1_HepG2_activs_per_elem.columns]
pool1_pMPRA1_K562_activs_per_elem.columns = ["K562_%s" % x if "rna" in x else x for x in pool1_pMPRA1_K562_activs_per_elem.columns]

pool1_pNoCMVMPRA1_HeLa_activs_per_elem.columns = ["HeLa_noCMV_%s" % x if "rna" in x else x for x in pool1_pNoCMVMPRA1_HeLa_activs_per_elem.columns]
pool1_pNoCMVMPRA1_HepG2_activs_per_elem.columns = ["HepG2_noCMV_%s" % x if "rna" in x else x for x in pool1_pNoCMVMPRA1_HepG2_activs_per_elem.columns]
pool1_pNoCMVMPRA1_K562_activs_per_elem.columns = ["K562_noCMV_%s" % x if "rna" in x else x for x in pool1_pNoCMVMPRA1_K562_activs_per_elem.columns]


# In[27]:


pool1 = pool1_pMPRA1_HeLa_activs_per_elem.merge(pool1_pMPRA1_HepG2_activs_per_elem, on=["unique_id", "element"], how="outer").merge(pool1_pMPRA1_K562_activs_per_elem, on=["unique_id", "element"], how="outer")
pool1_all = pool1_pMPRA1_HeLa_activs_per_elem.merge(pool1_pMPRA1_HepG2_activs_per_elem, on=["unique_id", "element"], how="outer").merge(pool1_pMPRA1_K562_activs_per_elem, on=["unique_id", "element"], how="outer").merge(pool1_pNoCMVMPRA1_HeLa_activs_per_elem, on=["unique_id", "element"], how="outer").merge(pool1_pNoCMVMPRA1_HepG2_activs_per_elem, on=["unique_id", "element"], how="outer").merge(pool1_pNoCMVMPRA1_K562_activs_per_elem, on=["unique_id", "element"], how="outer")
pool1_all.head()


# In[28]:


pool1_no_neg = pool1[(~pool1.unique_id.str.contains("RANDOM")) | (~pool1.unique_id.str.contains("SCRAMBLED"))]
pool1_no_neg = pool1_no_neg.drop("unique_id", axis=1)
pool1_no_neg_corr = pool1_no_neg.corr(method="spearman")

pool1_all_no_neg = pool1_all[(~pool1_all.unique_id.str.contains("RANDOM")) | (~pool1_all.unique_id.str.contains("SCRAMBLED"))]
pool1_all_no_neg = pool1_all_no_neg.drop("unique_id", axis=1)
pool1_all_no_neg_corr = pool1_all_no_neg.corr(method="spearman")


# In[29]:


pool1_all_no_neg_corr = pool1_all_no_neg.corr(method="pearson")
cmap = sns.cubehelix_palette(as_cmap=True)
cg = sns.clustermap(pool1_all_no_neg_corr, figsize=(7.2,7.2), cmap=cmap, annot=False)
_ = plt.setp(cg.ax_heatmap.yaxis.get_majorticklabels(), rotation=0)
plt.suptitle("pearson correlation of replicates\nquantile-normalized activities per element (no neg controls)")
cg.savefig("Fig_S3B.pdf", dpi="figure", transparent=True, bbox_inches="tight")


# ### pool 2 (deletion pool)

# In[30]:


pool2_pMPRA1_HepG2_activs_per_elem.columns = ["HepG2_%s" % x if "rna" in x else x for x in pool2_pMPRA1_HepG2_activs_per_elem.columns]
pool2_pMPRA1_K562_activs_per_elem.columns = ["K562_%s" % x if "rna" in x else x for x in pool2_pMPRA1_K562_activs_per_elem.columns]


# In[31]:


pool2 = pool2_pMPRA1_HepG2_activs_per_elem.merge(pool2_pMPRA1_K562_activs_per_elem, on=["unique_id", "element"], how="outer")
pool2.head()


# In[32]:


pool2_no_neg = pool2[(~pool2.unique_id.str.contains("RANDOM")) | (~pool2.unique_id.str.contains("SCRAMBLED"))]
pool2_no_neg = pool2_no_neg.drop("unique_id", axis=1)
pool2_no_neg_corr = pool2_no_neg.corr(method="spearman")


# In[33]:


pool2_no_neg_corr = pool2_no_neg.corr(method="pearson")
cg = sns.clustermap(pool2_no_neg_corr, figsize=(7.2, 7.2), annot=True, cmap=cmap, annot_kws={"fontsize":fontsize})
_ = plt.setp(cg.ax_heatmap.yaxis.get_majorticklabels(), rotation=0)
plt.suptitle("pearson correlation of replicates\nquantile-normalized activities per element (no neg controls)")


# ## 7. average pairwise spearman & pearson w/in each condition

# ### pool1 - HeLa min. promoter

# In[34]:


pool1_hela_cmv_p = pool1_all_no_neg[["HeLa_rna_1", "HeLa_rna_2", "HeLa_rna_3", "HeLa_rna_4"]].corr(method="pearson")
pool1_hela_cmv_s = pool1_all_no_neg[["HeLa_rna_1", "HeLa_rna_2", "HeLa_rna_3", "HeLa_rna_4"]].corr(method="spearman")

tmp = np.tril(pool1_hela_cmv_p, k=-1)
tmp[tmp == 0] = np.nan
pool1_hela_cmv_avg_p = np.nanmean(tmp)

tmp = np.tril(pool1_hela_cmv_s, k=-1)
tmp[tmp == 0] = np.nan
pool1_hela_cmv_avg_s = np.nanmean(tmp)

print("avg pearson: %s" % (pool1_hela_cmv_avg_p))
print("avg spearman: %s" % (pool1_hela_cmv_avg_s))


# ### pool1 - HepG2 min. promoter

# In[35]:


pool1_hepg2_cmv_p = pool1_all_no_neg[["HepG2_rna_3", "HepG2_rna_4", "HepG2_rna_5", "HepG2_rna_6", "HepG2_rna_7", "HepG2_rna_8", "HepG2_rna_9", "HepG2_rna_10", "HepG2_rna_11", "HepG2_rna_12", "HepG2_rna_13", "HepG2_rna_14"]].corr(method="pearson")
pool1_hepg2_cmv_s = pool1_all_no_neg[["HepG2_rna_3", "HepG2_rna_4", "HepG2_rna_5", "HepG2_rna_6", "HepG2_rna_7", "HepG2_rna_8", "HepG2_rna_9", "HepG2_rna_10", "HepG2_rna_11", "HepG2_rna_12", "HepG2_rna_13", "HepG2_rna_14"]].corr(method="spearman")

tmp = np.tril(pool1_hepg2_cmv_p, k=-1)
tmp[tmp == 0] = np.nan
pool1_hepg2_cmv_avg_p = np.nanmean(tmp)

tmp = np.tril(pool1_hepg2_cmv_s, k=-1)
tmp[tmp == 0] = np.nan
pool1_hepg2_cmv_avg_s = np.nanmean(tmp)

print("avg pearson: %s" % (pool1_hepg2_cmv_avg_p))
print("avg spearman: %s" % (pool1_hepg2_cmv_avg_s))


# ### pool1 - K562 min. promoter

# In[36]:


pool1_k562_cmv_p = pool1_all_no_neg[["K562_rna_1", "K562_rna_2", "K562_rna_3", "K562_rna_4"]].corr(method="pearson")
pool1_k562_cmv_s = pool1_all_no_neg[["K562_rna_1", "K562_rna_2", "K562_rna_3", "K562_rna_4"]].corr(method="spearman")

tmp = np.tril(pool1_k562_cmv_p, k=-1)
tmp[tmp == 0] = np.nan
pool1_k562_cmv_avg_p = np.nanmean(tmp)

tmp = np.tril(pool1_k562_cmv_s, k=-1)
tmp[tmp == 0] = np.nan
pool1_k562_cmv_avg_s = np.nanmean(tmp)

print("avg pearson: %s" % (pool1_k562_cmv_avg_p))
print("avg spearman: %s" % (pool1_k562_cmv_avg_s))


# ### pool1 - HeLa no min. promoter

# In[37]:


pool1_hela_no_cmv_p = pool1_all_no_neg[["HeLa_noCMV_rna_1", "HeLa_noCMV_rna_2", "HeLa_noCMV_rna_3", "HeLa_noCMV_rna_4"]].corr(method="pearson")
pool1_hela_no_cmv_s = pool1_all_no_neg[["HeLa_noCMV_rna_1", "HeLa_noCMV_rna_2", "HeLa_noCMV_rna_3", "HeLa_noCMV_rna_4"]].corr(method="spearman")

tmp = np.tril(pool1_hela_no_cmv_p, k=-1)
tmp[tmp == 0] = np.nan
pool1_hela_no_cmv_avg_p = np.nanmean(tmp)

tmp = np.tril(pool1_hela_no_cmv_s, k=-1)
tmp[tmp == 0] = np.nan
pool1_hela_no_cmv_avg_s = np.nanmean(tmp)

print("avg pearson: %s" % (pool1_hela_no_cmv_avg_p))
print("avg spearman: %s" % (pool1_hela_no_cmv_avg_s))


# ### pool1 - HepG2 no min. promoter

# In[38]:


pool1_hepg2_no_cmv_p = pool1_all_no_neg[["HepG2_noCMV_rna_3", "HepG2_noCMV_rna_4", "HepG2_noCMV_rna_5", "HepG2_noCMV_rna_6"]].corr(method="pearson")
pool1_hepg2_no_cmv_s = pool1_all_no_neg[["HepG2_noCMV_rna_3", "HepG2_noCMV_rna_4", "HepG2_noCMV_rna_5", "HepG2_noCMV_rna_6"]].corr(method="spearman")

tmp = np.tril(pool1_hepg2_no_cmv_p, k=-1)
tmp[tmp == 0] = np.nan
pool1_hepg2_no_cmv_avg_p = np.nanmean(tmp)

tmp = np.tril(pool1_hepg2_no_cmv_s, k=-1)
tmp[tmp == 0] = np.nan
pool1_hepg2_no_cmv_avg_s = np.nanmean(tmp)

print("avg pearson: %s" % (pool1_hepg2_no_cmv_avg_p))
print("avg spearman: %s" % (pool1_hepg2_no_cmv_avg_s))


# ### pool1 - K562 no min. promoter

# In[39]:


pool1_k562_no_cmv_p = pool1_all_no_neg[["K562_noCMV_rna_1", "K562_noCMV_rna_2", "K562_noCMV_rna_3", "K562_noCMV_rna_4"]].corr(method="pearson")
pool1_k562_no_cmv_s = pool1_all_no_neg[["K562_noCMV_rna_1", "K562_noCMV_rna_2", "K562_noCMV_rna_3", "K562_noCMV_rna_4"]].corr(method="spearman")

tmp = np.tril(pool1_k562_no_cmv_p, k=-1)
tmp[tmp == 0] = np.nan
pool1_k562_no_cmv_avg_p = np.nanmean(tmp)

tmp = np.tril(pool1_k562_no_cmv_s, k=-1)
tmp[tmp == 0] = np.nan
pool1_k562_no_cmv_avg_s = np.nanmean(tmp)

print("avg pearson: %s" % (pool1_k562_no_cmv_avg_p))
print("avg spearman: %s" % (pool1_k562_no_cmv_avg_s))


# ## pool1 average

# In[40]:


pool1_cmv_avg_p = np.mean([pool1_hela_cmv_avg_p, pool1_hepg2_cmv_avg_p, pool1_k562_cmv_avg_p])
pool1_no_cmv_avg_p = np.mean([pool1_hela_no_cmv_avg_p, pool1_hepg2_no_cmv_avg_p, pool1_k562_no_cmv_avg_p])
pool1_avg_p = np.mean([pool1_cmv_avg_p, pool1_no_cmv_avg_p])

pool1_cmv_avg_s = np.mean([pool1_hela_cmv_avg_s, pool1_hepg2_cmv_avg_s, pool1_k562_cmv_avg_s])
pool1_no_cmv_avg_s = np.mean([pool1_hela_no_cmv_avg_s, pool1_hepg2_no_cmv_avg_s, pool1_k562_no_cmv_avg_s])
pool1_avg_s = np.mean([pool1_cmv_avg_s, pool1_no_cmv_avg_s])

print("avg pool1 CMV pearson: %s" % (pool1_cmv_avg_p))
print("avg pool1 no CMV pearson: %s" % (pool1_no_cmv_avg_p))
print("avg all pool1 pearson: %s" % (pool1_avg_p))

print("")
print("avg pool1 CMV spearman: %s" % (pool1_cmv_avg_s))
print("avg pool1 no CMV spearman: %s" % (pool1_no_cmv_avg_s))
print("avg all pool1 spearman: %s" % (pool1_avg_s))


# ### pool2 - HepG2 (min. promoter)

# In[41]:


pool2_hepg2_cmv_p = pool2_no_neg[["HepG2_rna_3", "HepG2_rna_4", "HepG2_rna_5", "HepG2_rna_6", "HepG2_rna_7", "HepG2_rna_8", "HepG2_rna_9", "HepG2_rna_10"]].corr(method="pearson")
pool2_hepg2_cmv_s = pool2_no_neg[["HepG2_rna_3", "HepG2_rna_4", "HepG2_rna_5", "HepG2_rna_6", "HepG2_rna_7", "HepG2_rna_8", "HepG2_rna_9", "HepG2_rna_10"]].corr(method="spearman")

tmp = np.tril(pool2_hepg2_cmv_p, k=-1)
tmp[tmp == 0] = np.nan
pool2_hepg2_cmv_avg_p = np.nanmean(tmp)

tmp = np.tril(pool2_hepg2_cmv_s, k=-1)
tmp[tmp == 0] = np.nan
pool2_hepg2_cmv_avg_s = np.nanmean(tmp)

print("avg pearson: %s" % (pool2_hepg2_cmv_avg_p))
print("avg spearman: %s" % (pool2_hepg2_cmv_avg_s))


# ### pool2 - K562 (min. promoter)

# In[42]:


pool2_k562_cmv_p = pool2_no_neg[["K562_rna_1", "K562_rna_2", "K562_rna_3", "K562_rna_4"]].corr(method="pearson")
pool2_k562_cmv_s = pool2_no_neg[["K562_rna_1", "K562_rna_2", "K562_rna_3", "K562_rna_4"]].corr(method="spearman")

tmp = np.tril(pool2_k562_cmv_p, k=-1)
tmp[tmp == 0] = np.nan
pool2_k562_cmv_avg_p = np.nanmean(tmp)

tmp = np.tril(pool2_k562_cmv_s, k=-1)
tmp[tmp == 0] = np.nan
pool2_k562_cmv_avg_s = np.nanmean(tmp)

print("avg pearson: %s" % (pool2_k562_cmv_avg_p))
print("avg spearman: %s" % (pool2_k562_cmv_avg_s))


# ## pool2 average

# In[43]:


pool2_avg_p = np.mean([pool2_hepg2_cmv_avg_p, pool2_k562_cmv_avg_p])
pool2_avg_s = np.mean([pool2_hepg2_cmv_avg_s, pool2_k562_cmv_avg_s])

print("avg all pool2 pearson: %s" % (pool2_avg_p))
print("avg all pool2 spearman: %s" % (pool2_avg_s))
```
