## Supplementary material for "High-throughput functional analysis of IncRNA core promoters elucidatesrules governing tissue-specificity": 03__pvals.py

```
# coding: utf-8

# # 02__pvals
# # calculating significantly active/repressive tiles (compared to neg ctrls)
# 
# in this notebook, i calculate significantly active/repressive tiles by comparing them to random negative control sequences. this is done per replicate. the *barcode* activity values -- corresponding to a given reference element -- are compared to all negative control activity values using a two-sided wilcoxon test. i require a minimum of 10 barcodes to be present (Pool1 reference sequences have at least 15 barcodes, and Pool2 reference sequences have at least 80 barcodes) in order to calculate a p-value, otherwise i leave it NA. once the p-values are calculated per replicate, i then combine the p-values using stouffer's method. finally, i correct the p-values for multiple hypothesis testing using the Bonferroni family-wise error rate correction method. 
# 
# i define significantly active tiles as those that have a corrected p-value (q-value) of < 0.05, and that are at least a median foldchange of 0.5 above the negative control sequences in 75% of replicates. i define significantly repressive tiles as those that have a q-value of < 0.05, and that are at most a median fold change of -0.5 below the negative control sequences in 75% of replicates.
# 
# since HepG2 has more replicates than the other cell lines, in order to ensure that HepG2 results could be compared to the other cell lines at about the same power, i also downsampled the HepG2 replicates to get a "downsampled q-value". in that case, i randomly sampled 4 replicates, combined the p-values, corrected the p-values, and repeated the sampling process 100 times. i then considered the tile significant if it had a significant q-value in 75% of samples, with the same rules as above.
# 
# ------
# 
# no figures in this notebook

# In[1]:


import warnings
warnings.filterwarnings('ignore')

import pandas as pd
import numpy as np
import matplotlib as mpl
import matplotlib.pyplot as plt
import seaborn as sns
import sys
import time

from decimal import Decimal
from itertools import chain
from mpl_toolkits.axes_grid1 import make_axes_locatable
from scipy import stats
from statsmodels.sandbox.stats import multicomp

# import utils
sys.path.append("../../utils")
from plotting_utils import *
from misc_utils import *
from norm_utils import *

get_ipython().magic('matplotlib inline')


# In[2]:


sns.set(**PAPER_PRESET)
fontsize = PAPER_FONTSIZE


# In[3]:


np.random.seed(12345)


# ## functions

# In[4]:


def is_sig(row, col, thresh):
    if pd.isnull(row[col]):
        return "NA"
    else:
        if row[col] < thresh:
            return "sig"
        else:
            return "not sig"
        
def is_sig_bin(row, col, thresh):
    if pd.isnull(row[col]):
        return np.nan
    else:
        if row[col] < thresh:
            return 1
        else:
            return 0


# ## variables

# In[5]:


activs_dir = "../../data/02__activs"
out_dir = "../../data/03__pvals"
get_ipython().system('mkdir -p $out_dir')


# In[6]:


pool1_index_f = "../../data/00__index/tss_oligo_pool.index.txt"
pool2_index_f = "../../data/00__index/dels_oligo_pool.index.txt"


# In[7]:


pool1_pMPRA1_HeLa_activ_elem_f = "%s/POOL1__pMPRA1__HeLa__activities_per_element.txt" % activs_dir
pool1_pMPRA1_HeLa_activ_barc_f = "%s/POOL1__pMPRA1__HeLa__activities_per_barcode.txt" % activs_dir
pool1_pMPRA1_HeLa_pval_f = "%s/POOL1__pMPRA1__HeLa__pvals.txt" % out_dir

pool1_pMPRA1_HepG2_activ_elem_f = "%s/POOL1__pMPRA1__HepG2__activities_per_element.txt" % activs_dir
pool1_pMPRA1_HepG2_activ_barc_f = "%s/POOL1__pMPRA1__HepG2__activities_per_barcode.txt" % activs_dir
pool1_pMPRA1_HepG2_pval_f = "%s/POOL1__pMPRA1__HepG2__pvals.txt" % out_dir

pool1_pMPRA1_K562_activ_elem_f = "%s/POOL1__pMPRA1__K562__activities_per_element.txt" % activs_dir
pool1_pMPRA1_K562_activ_barc_f = "%s/POOL1__pMPRA1__K562__activities_per_barcode.txt" % activs_dir
pool1_pMPRA1_K562_pval_f = "%s/POOL1__pMPRA1__K562__pvals.txt" % out_dir

pool1_pNoCMVMPRA1_HeLa_activ_elem_f = "%s/POOL1__pNoCMVMPRA1__HeLa__activities_per_element.txt" % activs_dir
pool1_pNoCMVMPRA1_HeLa_activ_barc_f = "%s/POOL1__pNoCMVMPRA1__HeLa__activities_per_barcode.txt" % activs_dir
pool1_pNoCMVMPRA1_HeLa_pval_f = "%s/POOL1__pNoCMVMPRA1__HeLa__pvals.txt" % out_dir

pool1_pNoCMVMPRA1_HepG2_activ_elem_f = "%s/POOL1__pNoCMVMPRA1__HepG2__activities_per_element.txt" % activs_dir
pool1_pNoCMVMPRA1_HepG2_activ_barc_f = "%s/POOL1__pNoCMVMPRA1__HepG2__activities_per_barcode.txt" % activs_dir
pool1_pNoCMVMPRA1_HepG2_pval_f = "%s/POOL1__pNoCMVMPRA1__HepG2__pvals.txt" % out_dir

pool1_pNoCMVMPRA1_K562_activ_elem_f = "%s/POOL1__pNoCMVMPRA1__K562__activities_per_element.txt" % activs_dir
pool1_pNoCMVMPRA1_K562_activ_barc_f = "%s/POOL1__pNoCMVMPRA1__K562__activities_per_barcode.txt" % activs_dir
pool1_pNoCMVMPRA1_K562_pval_f = "%s/POOL1__pNoCMVMPRA1__K562__pvals.txt" % out_dir

pool2_pMPRA1_HepG2_activ_elem_f = "%s/POOL2__pMPRA1__HepG2__activities_per_element.txt" % activs_dir
pool2_pMPRA1_HepG2_activ_barc_f = "%s/POOL2__pMPRA1__HepG2__activities_per_barcode.txt" % activs_dir
pool2_pMPRA1_HepG2_pval_f = "%s/POOL2__pMPRA1__HepG2__pvals.txt" % out_dir

pool2_pMPRA1_K562_activ_elem_f = "%s/POOL2__pMPRA1__K562__activities_per_element.txt" % activs_dir
pool2_pMPRA1_K562_activ_barc_f = "%s/POOL2__pMPRA1__K562__activities_per_barcode.txt" % activs_dir
pool2_pMPRA1_K562_pval_f = "%s/POOL2__pMPRA1__K562__pvals.txt" % out_dir


# In[8]:


# number of times to downsample hepg2 replicates
n_samples = 100


# ## 1. import data

# In[9]:


pool1_index = pd.read_table(pool1_index_f, sep="\t")
pool2_index = pd.read_table(pool2_index_f, sep="\t")


# In[10]:


pool1_index_elem = pool1_index[["element", "oligo_type", "unique_id", "dupe_info", "SNP"]]
pool2_index_elem = pool2_index[["element", "oligo_type", "unique_id", "dupe_info", "SNP"]]

pool1_index_elem = pool1_index_elem.drop_duplicates()
pool2_index_elem = pool2_index_elem.drop_duplicates()


# In[11]:


pool1_pMPRA1_HeLa_activ_barc = pd.read_table(pool1_pMPRA1_HeLa_activ_barc_f)
pool1_pMPRA1_HepG2_activ_barc = pd.read_table(pool1_pMPRA1_HepG2_activ_barc_f)
pool1_pMPRA1_K562_activ_barc = pd.read_table(pool1_pMPRA1_K562_activ_barc_f)
pool1_pMPRA1_K562_activ_barc.head()


# In[12]:


pool1_pNoCMVMPRA1_HeLa_activ_barc = pd.read_table(pool1_pNoCMVMPRA1_HeLa_activ_barc_f)
pool1_pNoCMVMPRA1_HepG2_activ_barc = pd.read_table(pool1_pNoCMVMPRA1_HepG2_activ_barc_f)
pool1_pNoCMVMPRA1_K562_activ_barc = pd.read_table(pool1_pNoCMVMPRA1_K562_activ_barc_f)


# In[13]:


pool2_pMPRA1_HepG2_activ_barc = pd.read_table(pool2_pMPRA1_HepG2_activ_barc_f)
pool2_pMPRA1_K562_activ_barc = pd.read_table(pool2_pMPRA1_K562_activ_barc_f)


# In[14]:


pool1_pMPRA1_HeLa_activ_elem = pd.read_table(pool1_pMPRA1_HeLa_activ_elem_f)
pool1_pMPRA1_HepG2_activ_elem = pd.read_table(pool1_pMPRA1_HepG2_activ_elem_f)
pool1_pMPRA1_K562_activ_elem = pd.read_table(pool1_pMPRA1_K562_activ_elem_f)

pool1_pNoCMVMPRA1_HeLa_activ_elem = pd.read_table(pool1_pNoCMVMPRA1_HeLa_activ_elem_f)
pool1_pNoCMVMPRA1_HepG2_activ_elem = pd.read_table(pool1_pNoCMVMPRA1_HepG2_activ_elem_f)
pool1_pNoCMVMPRA1_K562_activ_elem = pd.read_table(pool1_pNoCMVMPRA1_K562_activ_elem_f)

pool2_pMPRA1_HepG2_activ_elem = pd.read_table(pool2_pMPRA1_HepG2_activ_elem_f)
pool2_pMPRA1_K562_activ_elem = pd.read_table(pool2_pMPRA1_K562_activ_elem_f)

pool1_pMPRA1_HeLa_activ_elem.head()


# ## 2. merge with index

# In[15]:


pool1_pMPRA1_HeLa_activ_barc = pool1_pMPRA1_HeLa_activ_barc.merge(pool1_index, left_on="barcode", right_on="barcode", how="left")
pool1_pMPRA1_HepG2_activ_barc = pool1_pMPRA1_HepG2_activ_barc.merge(pool1_index, left_on="barcode", right_on="barcode", how="left")
pool1_pMPRA1_K562_activ_barc = pool1_pMPRA1_K562_activ_barc.merge(pool1_index, left_on="barcode", right_on="barcode", how="left")

pool1_pNoCMVMPRA1_HeLa_activ_barc = pool1_pNoCMVMPRA1_HeLa_activ_barc.merge(pool1_index, left_on="barcode", right_on="barcode", how="left")
pool1_pNoCMVMPRA1_HepG2_activ_barc = pool1_pNoCMVMPRA1_HepG2_activ_barc.merge(pool1_index, left_on="barcode", right_on="barcode", how="left")
pool1_pNoCMVMPRA1_K562_activ_barc = pool1_pNoCMVMPRA1_K562_activ_barc.merge(pool1_index, left_on="barcode", right_on="barcode", how="left")

pool2_pMPRA1_HepG2_activ_barc = pool2_pMPRA1_HepG2_activ_barc.merge(pool2_index, left_on="barcode", right_on="barcode", how="left")
pool2_pMPRA1_K562_activ_barc = pool2_pMPRA1_K562_activ_barc.merge(pool2_index, left_on="barcode", right_on="barcode", how="left")


# ## 3. calculate p-values

# ### using RANDOM sequences as negative controls

# ### p values calculated per replicate

# In[16]:


pool1_pMPRA1_HeLa_activ_barc["better_type"] = pool1_pMPRA1_HeLa_activ_barc.apply(better_type, axis=1)
pool1_pMPRA1_HeLa_reps = [x for x in pool1_pMPRA1_HeLa_activ_barc.columns if "rna" in x]
pool1_pMPRA1_HeLa_us, pool1_pMPRA1_HeLa_pvals, pool1_pMPRA1_HeLa_fcs = element_p_val_per_rep_neg_controls(pool1_pMPRA1_HeLa_activ_barc, pool1_pMPRA1_HeLa_reps, 10, ["RANDOM"], ["WILDTYPE", "FLIPPED", "CONTROL", "SNP", "CONTROL_SNP"])


# In[17]:


pool1_pMPRA1_HepG2_activ_barc["better_type"] = pool1_pMPRA1_HepG2_activ_barc.apply(better_type, axis=1)
pool1_pMPRA1_HepG2_reps = [x for x in pool1_pMPRA1_HepG2_activ_barc.columns if "rna" in x]
pool1_pMPRA1_HepG2_us, pool1_pMPRA1_HepG2_pvals, pool1_pMPRA1_HepG2_fcs = element_p_val_per_rep_neg_controls(pool1_pMPRA1_HepG2_activ_barc, pool1_pMPRA1_HepG2_reps, 10, ["RANDOM"], ["WILDTYPE", "FLIPPED", "CONTROL", "SNP", "CONTROL_SNP"])


# In[18]:


pool1_pMPRA1_K562_activ_barc["better_type"] = pool1_pMPRA1_K562_activ_barc.apply(better_type, axis=1)
pool1_pMPRA1_K562_reps = [x for x in pool1_pMPRA1_K562_activ_barc.columns if "rna" in x]
pool1_pMPRA1_K562_us, pool1_pMPRA1_K562_pvals, pool1_pMPRA1_K562_fcs = element_p_val_per_rep_neg_controls(pool1_pMPRA1_K562_activ_barc, pool1_pMPRA1_K562_reps, 10, ["RANDOM"], ["WILDTYPE", "FLIPPED", "CONTROL", "SNP", "CONTROL_SNP"])


# In[19]:


pool1_pNoCMVMPRA1_HeLa_activ_barc["better_type"] = pool1_pNoCMVMPRA1_HeLa_activ_barc.apply(better_type, axis=1)
pool1_pNoCMVMPRA1_HeLa_reps = [x for x in pool1_pNoCMVMPRA1_HeLa_activ_barc.columns if "rna" in x]
pool1_pNoCMVMPRA1_HeLa_us, pool1_pNoCMVMPRA1_HeLa_pvals, pool1_pNoCMVMPRA1_HeLa_fcs = element_p_val_per_rep_neg_controls(pool1_pNoCMVMPRA1_HeLa_activ_barc, pool1_pNoCMVMPRA1_HeLa_reps, 10, ["RANDOM"], ["WILDTYPE", "FLIPPED", "CONTROL", "SNP", "CONTROL_SNP"])


# In[20]:


pool1_pNoCMVMPRA1_HepG2_activ_barc["better_type"] = pool1_pNoCMVMPRA1_HepG2_activ_barc.apply(better_type, axis=1)
pool1_pNoCMVMPRA1_HepG2_reps = [x for x in pool1_pNoCMVMPRA1_HepG2_activ_barc.columns if "rna" in x]
pool1_pNoCMVMPRA1_HepG2_us, pool1_pNoCMVMPRA1_HepG2_pvals, pool1_pNoCMVMPRA1_HepG2_fcs = element_p_val_per_rep_neg_controls(pool1_pNoCMVMPRA1_HepG2_activ_barc, pool1_pNoCMVMPRA1_HepG2_reps, 10, ["RANDOM"], ["WILDTYPE", "FLIPPED", "CONTROL", "SNP", "CONTROL_SNP"])


# In[21]:


pool1_pNoCMVMPRA1_K562_activ_barc["better_type"] = pool1_pNoCMVMPRA1_K562_activ_barc.apply(better_type, axis=1)
pool1_pNoCMVMPRA1_K562_reps = [x for x in pool1_pNoCMVMPRA1_K562_activ_barc.columns if "rna" in x]
pool1_pNoCMVMPRA1_K562_us, pool1_pNoCMVMPRA1_K562_pvals, pool1_pNoCMVMPRA1_K562_fcs = element_p_val_per_rep_neg_controls(pool1_pNoCMVMPRA1_K562_activ_barc, pool1_pNoCMVMPRA1_K562_reps, 10, ["RANDOM"], ["WILDTYPE", "FLIPPED", "CONTROL", "SNP", "CONTROL_SNP"])


# In[22]:


pool2_pMPRA1_HepG2_activ_barc["better_type"] = pool2_pMPRA1_HepG2_activ_barc.apply(better_type, axis=1)
pool2_pMPRA1_HepG2_reps = [x for x in pool2_pMPRA1_HepG2_activ_barc.columns if "rna" in x]
pool2_pMPRA1_HepG2_us, pool2_pMPRA1_HepG2_pvals, pool2_pMPRA1_HepG2_fcs = element_p_val_per_rep_neg_controls(pool2_pMPRA1_HepG2_activ_barc, pool2_pMPRA1_HepG2_reps, 10, ["RANDOM"], ["WILDTYPE", "FLIPPED", "CONTROL", "SNP", "CONTROL_SNP"])


# In[23]:


pool2_pMPRA1_K562_activ_barc["better_type"] = pool2_pMPRA1_K562_activ_barc.apply(better_type, axis=1)
pool2_pMPRA1_K562_reps = [x for x in pool2_pMPRA1_K562_activ_barc.columns if "rna" in x]
pool2_pMPRA1_K562_us, pool2_pMPRA1_K562_pvals, pool2_pMPRA1_K562_fcs = element_p_val_per_rep_neg_controls(pool2_pMPRA1_K562_activ_barc, pool2_pMPRA1_K562_reps, 10, ["RANDOM"], ["WILDTYPE", "FLIPPED", "CONTROL", "SNP", "CONTROL_SNP"])


# In[24]:


# change dicts into dfs
pool1_pMPRA1_HeLa_pvals_df = pd.DataFrame.from_dict(pool1_pMPRA1_HeLa_pvals, orient="index").reset_index()
pool1_pMPRA1_HeLa_fcs_df = pd.DataFrame.from_dict(pool1_pMPRA1_HeLa_fcs, orient="index").reset_index()

pool1_pMPRA1_HepG2_pvals_df = pd.DataFrame.from_dict(pool1_pMPRA1_HepG2_pvals, orient="index").reset_index()
pool1_pMPRA1_HepG2_fcs_df = pd.DataFrame.from_dict(pool1_pMPRA1_HepG2_fcs, orient="index").reset_index()

pool1_pMPRA1_K562_pvals_df = pd.DataFrame.from_dict(pool1_pMPRA1_K562_pvals, orient="index").reset_index()
pool1_pMPRA1_K562_fcs_df = pd.DataFrame.from_dict(pool1_pMPRA1_K562_fcs, orient="index").reset_index()

pool1_pNoCMVMPRA1_HeLa_pvals_df = pd.DataFrame.from_dict(pool1_pNoCMVMPRA1_HeLa_pvals, orient="index").reset_index()
pool1_pNoCMVMPRA1_HeLa_fcs_df = pd.DataFrame.from_dict(pool1_pNoCMVMPRA1_HeLa_fcs, orient="index").reset_index()

pool1_pNoCMVMPRA1_HepG2_pvals_df = pd.DataFrame.from_dict(pool1_pNoCMVMPRA1_HepG2_pvals, orient="index").reset_index()
pool1_pNoCMVMPRA1_HepG2_fcs_df = pd.DataFrame.from_dict(pool1_pNoCMVMPRA1_HepG2_fcs, orient="index").reset_index()

pool1_pNoCMVMPRA1_K562_pvals_df = pd.DataFrame.from_dict(pool1_pNoCMVMPRA1_K562_pvals, orient="index").reset_index()
pool1_pNoCMVMPRA1_K562_fcs_df = pd.DataFrame.from_dict(pool1_pNoCMVMPRA1_K562_fcs, orient="index").reset_index()

pool2_pMPRA1_HepG2_pvals_df = pd.DataFrame.from_dict(pool2_pMPRA1_HepG2_pvals, orient="index").reset_index()
pool2_pMPRA1_HepG2_fcs_df = pd.DataFrame.from_dict(pool2_pMPRA1_HepG2_fcs, orient="index").reset_index()

pool2_pMPRA1_K562_pvals_df = pd.DataFrame.from_dict(pool2_pMPRA1_K562_pvals, orient="index").reset_index()
pool2_pMPRA1_K562_fcs_df = pd.DataFrame.from_dict(pool2_pMPRA1_K562_fcs, orient="index").reset_index()


# In[25]:


# merge w/ barc data
pool1_pMPRA1_HeLa_activ_barc = pool1_pMPRA1_HeLa_activ_barc.merge(pool1_pMPRA1_HeLa_pvals_df, left_on="element", 
                                                                  right_on="index", how="left",
                                                                  suffixes=("", "_pval")).drop("index", axis=1)
pool1_pMPRA1_HeLa_activ_barc = pool1_pMPRA1_HeLa_activ_barc.merge(pool1_pMPRA1_HeLa_fcs_df, left_on="element", 
                                                                  right_on="index", how="left", 
                                                                  suffixes=("", "_log2fc")).drop("index", axis=1)


pool1_pMPRA1_HepG2_activ_barc = pool1_pMPRA1_HepG2_activ_barc.merge(pool1_pMPRA1_HepG2_pvals_df, left_on="element", 
                                                                   right_on="index", how="left",
                                                                   suffixes=("", "_pval")).drop("index", axis=1)
pool1_pMPRA1_HepG2_activ_barc = pool1_pMPRA1_HepG2_activ_barc.merge(pool1_pMPRA1_HepG2_fcs_df, left_on="element", 
                                                                    right_on="index", how="left", 
                                                                    suffixes=("", "_log2fc")).drop("index", axis=1)


pool1_pMPRA1_K562_activ_barc = pool1_pMPRA1_K562_activ_barc.merge(pool1_pMPRA1_K562_pvals_df, left_on="element", 
                                                                  right_on="index", how="left",
                                                                  suffixes=("", "_pval")).drop("index", axis=1)
pool1_pMPRA1_K562_activ_barc = pool1_pMPRA1_K562_activ_barc.merge(pool1_pMPRA1_K562_fcs_df, left_on="element", 
                                                                  right_on="index", how="left", 
                                                                  suffixes=("", "_log2fc")).drop("index", axis=1)


pool1_pNoCMVMPRA1_HeLa_activ_barc = pool1_pNoCMVMPRA1_HeLa_activ_barc.merge(pool1_pNoCMVMPRA1_HeLa_pvals_df, 
                                                                            left_on="element", 
                                                                            right_on="index", how="left",
                                                                            suffixes=("", "_pval")).drop("index", axis=1)
pool1_pNoCMVMPRA1_HeLa_activ_barc = pool1_pNoCMVMPRA1_HeLa_activ_barc.merge(pool1_pNoCMVMPRA1_HeLa_fcs_df, 
                                                                            left_on="element", 
                                                                            right_on="index", how="left", 
                                                                            suffixes=("", "_log2fc")).drop("index", axis=1)

pool1_pNoCMVMPRA1_HepG2_activ_barc = pool1_pNoCMVMPRA1_HepG2_activ_barc.merge(pool1_pNoCMVMPRA1_HepG2_pvals_df, 
                                                                              left_on="element", 
                                                                              right_on="index", how="left",
                                                                              suffixes=("", "_pval")).drop("index", axis=1)
pool1_pNoCMVMPRA1_HepG2_activ_barc = pool1_pNoCMVMPRA1_HepG2_activ_barc.merge(pool1_pNoCMVMPRA1_HepG2_fcs_df, 
                                                                              left_on="element", 
                                                                              right_on="index", how="left", 
                                                                              suffixes=("", "_log2fc")).drop("index", axis=1)


pool1_pNoCMVMPRA1_K562_activ_barc = pool1_pNoCMVMPRA1_K562_activ_barc.merge(pool1_pNoCMVMPRA1_K562_pvals_df, 
                                                                            left_on="element", 
                                                                            right_on="index", how="left",
                                                                            suffixes=("", "_pval")).drop("index", axis=1)
pool1_pNoCMVMPRA1_K562_activ_barc = pool1_pNoCMVMPRA1_K562_activ_barc.merge(pool1_pNoCMVMPRA1_K562_fcs_df, 
                                                                            left_on="element", 
                                                                            right_on="index", how="left", 
                                                                            suffixes=("", "_log2fc")).drop("index", axis=1)


pool2_pMPRA1_HepG2_activ_barc = pool2_pMPRA1_HepG2_activ_barc.merge(pool2_pMPRA1_HepG2_pvals_df, left_on="element", 
                                                                   right_on="index", how="left",
                                                                   suffixes=("", "_pval")).drop("index", axis=1)
pool2_pMPRA1_HepG2_activ_barc = pool2_pMPRA1_HepG2_activ_barc.merge(pool2_pMPRA1_HepG2_fcs_df, left_on="element", 
                                                                    right_on="index", how="left", 
                                                                    suffixes=("", "_log2fc")).drop("index", axis=1)


pool2_pMPRA1_K562_activ_barc = pool2_pMPRA1_K562_activ_barc.merge(pool2_pMPRA1_K562_pvals_df, left_on="element", 
                                                                  right_on="index", how="left",
                                                                  suffixes=("", "_pval")).drop("index", axis=1)
pool2_pMPRA1_K562_activ_barc = pool2_pMPRA1_K562_activ_barc.merge(pool2_pMPRA1_K562_fcs_df, left_on="element", 
                                                                  right_on="index", how="left", 
                                                                  suffixes=("", "_log2fc")).drop("index", axis=1)


# In[26]:


# extract columns
drop_cols = ["barcode", "full_oligo", "oligo_id"]

all_barc_dfs = [pool1_pMPRA1_HeLa_activ_barc, pool1_pMPRA1_HepG2_activ_barc, pool1_pMPRA1_K562_activ_barc,
                pool1_pNoCMVMPRA1_HeLa_activ_barc, pool1_pNoCMVMPRA1_HepG2_activ_barc, 
                pool1_pNoCMVMPRA1_K562_activ_barc, pool2_pMPRA1_HepG2_activ_barc, pool2_pMPRA1_K562_activ_barc]

all_elem_dfs = [pool1_pMPRA1_HeLa_activ_elem, pool1_pMPRA1_HepG2_activ_elem, pool1_pMPRA1_K562_activ_elem,
                pool1_pNoCMVMPRA1_HeLa_activ_elem, pool1_pNoCMVMPRA1_HepG2_activ_elem, 
                pool1_pNoCMVMPRA1_K562_activ_elem, pool2_pMPRA1_HepG2_activ_elem, pool2_pMPRA1_K562_activ_elem]

all_reps = [pool1_pMPRA1_HeLa_reps, pool1_pMPRA1_HepG2_reps, pool1_pMPRA1_K562_reps,
            pool1_pNoCMVMPRA1_HeLa_reps, pool1_pNoCMVMPRA1_HepG2_reps, pool1_pNoCMVMPRA1_K562_reps,
            pool2_pMPRA1_HepG2_reps, pool2_pMPRA1_K562_reps]

all_grp_dfs = []

for barc_df, elem_df, reps in zip(all_barc_dfs, all_elem_dfs, all_reps):
    samp_drop_cols = list(drop_cols)
    samp_drop_cols.extend(reps)
    
    df = elem_df.merge(barc_df.drop(samp_drop_cols, axis=1), on=["unique_id", "element"], how="left")
    df = df.drop_duplicates()
    all_grp_dfs.append(df)


# In[27]:


# correct p values
all_corr_dfs = []
for df in all_grp_dfs:
    pval_cols = [x for x in df.columns if "_pval" in x]
    for col in pval_cols:
        sub_df = df[~pd.isnull(df[col])][["unique_id", "element", col]]
        new_pvals = multicomp.multipletests(sub_df[col], method="bonferroni")[1]
        padj_col = "rna_%s_padj" % (col.split("_")[1])
        sub_df[padj_col] = new_pvals
        sub_df.drop(col, axis=1, inplace=True)
        df = df.merge(sub_df, on=["unique_id", "element"], how="left")
    all_corr_dfs.append(df)


# ## 4. use stouffer's method to combine p-values across replicates

# in this case, combine the *uncorrected* pvalues and *then adjust* using stouffer's method

# In[28]:


all_names = ["POOL1__pMPRA1__HeLa", "POOL1__pMPRA1__HepG2", "POOL1__pMPRA1__K562", "POOL1__pNoCMVMPRA1__HeLa",
             "POOL1__pNoCMVMPRA1__HepG2", "POOL1__pNoCMVMPRA1__K562", "POOL2__pMPRA1__HepG2", "POOL2__pMPRA1__K562"]


# In[29]:


all_comb_dfs = []
for name, df in zip(all_names, all_corr_dfs):
    cell_type = name.split("__")[2]
    print(cell_type)
    pval_cols = [x for x in df.columns if "_pval" in x]
    df["combined_pval"] = df.apply(combine_pvals, reps=pval_cols, axis=1)
    
    # correct combined pvals
    df_no_nan = df[~pd.isnull(df["combined_pval"])][["unique_id", "combined_pval"]]
    combined_padjs = multicomp.multipletests(df_no_nan["combined_pval"], method="bonferroni")[1]
    df_no_nan["combined_padj"] = combined_padjs
    df_no_nan.drop("combined_pval", axis=1, inplace=True)
    df = df.merge(df_no_nan, on="unique_id", how="left")
    
    l2fc_cols = [x for x in df.columns if "_log2fc" in x]
    df["combined_sig"] = df.apply(is_sig_combined, col="combined_padj", axis=1, thresh=0.05, l2fc_cols=l2fc_cols)
    df["combined_class"] = df.apply(is_active_or_repressive, sig_col="combined_sig", axis=1, 
                                    thresh=0.5, l2fc_cols=l2fc_cols)
    
    # if HepG2, where we have a lot of reps, downsample them 100x
    if cell_type == "HepG2":
        for n in range(n_samples):
            samp_pval_cols = list(np.random.choice(pval_cols, size=4))
            samp_reps = [x.split("_")[1] for x in samp_pval_cols]
            samp_l2fc_cols = []
            for x in samp_reps:
                samp_l2fc_cols.append("rna_%s_log2fc" % x)
            
            df["samp_%s_combined_pval" % n] = df.apply(combine_pvals, reps=samp_pval_cols, axis=1)
            
            # correct combined pvals
            df_no_nan = df[~pd.isnull(df["samp_%s_combined_pval" % n])][["unique_id", "samp_%s_combined_pval" % n]]
            combined_padjs = multicomp.multipletests(df_no_nan["samp_%s_combined_pval" % n], method="bonferroni")[1]
            df_no_nan["samp_%s_combined_padj" % n] = combined_padjs
            df = df.merge(df_no_nan, on="unique_id", how="left")
            df["samp_%s_combined_sig" % n] = df.apply(is_sig_combined, col="samp_%s_combined_padj" % n,
                                                      thresh=0.05, l2fc_cols=samp_l2fc_cols, axis=1)
            df["samp_%s_combined_class" % n] = df.apply(is_active_or_repressive, sig_col="samp_%s_combined_sig" % n, 
                                                        axis=1, thresh=0.5, l2fc_cols=samp_l2fc_cols)

    
        # consider it significant if sig in >75% of samples
        # take class that occurs max # of times
        samp_combined_cols = [x for x in df.columns if "samp" in x and "combined_sig" in x]
        samp_class_cols = [x for x in df.columns if "samp" in x and "combined_class" in x]
        
        df["downsamp_combined_sig"] = df.apply(downsamp_is_sig_combined, samp_combined_cols=samp_combined_cols, 
                                               n_samples=n_samples, axis=1)
        df["downsamp_combined_class"] = df.apply(downsamp_is_active_or_repressive, sig_col="downsamp_combined_sig",
                                                 samp_class_cols=samp_class_cols, axis=1)
        to_drop = [x for x in df.columns if x.startswith("samp_")]
        df.drop(to_drop, axis=1, inplace=True)
    
    
    tots = df.groupby(["better_type"])["element"].agg("count").reset_index()
    sigs = df[df["combined_sig"] == "sig"].groupby(["better_type"])["element"].agg("count").reset_index()
    perc = tots.merge(sigs, on="better_type", how="left", suffixes=("_tot", "_sig"))
    perc.fillna(0, inplace=True)
    perc["perc_sig"] = (perc["element_sig"]/perc["element_tot"])*100
    print(perc)
    
    
    if cell_type == "HepG2":
        sigs = df[df["downsamp_combined_sig"] == "sig"].groupby(["better_type"])["element"].agg("count").reset_index()
        perc = tots.merge(sigs, on="better_type", how="left", suffixes=("_tot", "_sig"))
        perc.fillna(0, inplace=True)
        perc["perc_sig"] = (perc["element_sig"]/perc["element_tot"])*100
        print(perc)
        
    all_comb_dfs.append(df)


# In[30]:


all_comb_dfs[1].head()


# In[31]:


all_comb_dfs[1].combined_sig.value_counts()


# In[32]:


all_comb_dfs[1].combined_class.value_counts()


# In[33]:


all_comb_dfs[1].downsamp_combined_sig.value_counts()


# In[34]:


all_comb_dfs[1].downsamp_combined_class.value_counts()


# ## 5. write files

# ### p-vals per element

# In[35]:


all_files = [pool1_pMPRA1_HeLa_pval_f, pool1_pMPRA1_HepG2_pval_f, pool1_pMPRA1_K562_pval_f,
             pool1_pNoCMVMPRA1_HeLa_pval_f, pool1_pNoCMVMPRA1_HepG2_pval_f, pool1_pNoCMVMPRA1_K562_pval_f,
             pool2_pMPRA1_HepG2_pval_f, pool2_pMPRA1_K562_pval_f]

final_dfs = []
for name, df, f in zip(all_names, all_comb_dfs, all_files):
    print(name)
    cols = ["unique_id", "element", "oligo_type"]
    pval_cols = [x for x in df.columns if ("_padj" in x or "_log2fc" in x) and "non_na" not in x]
    cols.extend(pval_cols)
    cols.extend(["combined_sig"])
    cols.extend(["combined_class"])
    if "HepG2" in name:
        downsamp_cols = [x for x in df.columns if "down" in x]
        cols.extend(downsamp_cols)
    
    sub_df = df[cols]
    sub_df.drop_duplicates(inplace=True)
    print(len(sub_df))
    #print(sub_df.head())
    sub_df.to_csv(f, sep="\t", index=False)
    final_dfs.append(sub_df)


# In[37]:


for name, df in zip(all_names, final_dfs):
    print(name)
    if "POOL1" in name:
        ids = pool1_index_elem["unique_id"]
    elif "POOL2" in name:
        ids = pool2_index_elem["unique_id"]
    missing = ids[~ids.isin(df["unique_id"])]
    print(len(missing))


# In[ ]:
```
