## Supplementary material for "High-throughput functional analysis of IncRNA core promoters elucidatesrules governing tissue-specificity": 04__activities.py

```
# coding: utf-8

# # 04__activities
# # analyzing activity levels per element (neg ctrls; between biotypes)
# 
# in this notebook, i perform analyses examining the activities of reference tiles in both pool1 and pool2. i compare reference sequences to negative controls, examine reference activities between biotypes, examine how MPRA activity compares to CAGE expression, and determine how many sequences are expressed across cell types.
# 
# ------
# 
# figures in this notebook:
# - **Fig 1C, Fig S4, Fig S12**: boxplots comparing reference sequences to negative controls & between biotypes
# - **Fig S6**: boxplots comparing biotypes when either randomly selecting TSSs or expression-matching TSSs
# - **Fig 1D**: KDE plot comparing CAGE cell type specificity and MPRA cell type specificity
# - **Fig S5**: KDE plot comparing mean CAGE expression and mean MPRA activity
# - **Fig 1E**: barplot showing % of reference sequences active in 1 or all 3 cell types

# In[1]:


import warnings
warnings.filterwarnings('ignore')

import pandas as pd
import numpy as np
import matplotlib as mpl
import matplotlib.pyplot as plt
import seaborn as sns
import sys
import time

from itertools import chain
from decimal import Decimal
from scipy import stats
from statsmodels.sandbox.stats import multicomp

from sklearn.preprocessing import StandardScaler
from sklearn.neighbors import NearestNeighbors

# import utils
sys.path.append("../../utils")
from plotting_utils import *
from misc_utils import *
from norm_utils import *

get_ipython().magic('matplotlib inline')


# In[2]:


sns.set(**PAPER_PRESET)
fontsize = PAPER_FONTSIZE


# ## variables

# In[3]:


activ_dir = "../../data/02__activs"
pval_dir = "../../data/03__pvals"
index_dir = "../../data/00__index"


# In[4]:


pool1_hela_barc_activ_f = "POOL1__pMPRA1__HeLa__activities_per_barcode.txt"
pool1_hepg2_barc_activ_f = "POOL1__pMPRA1__HepG2__activities_per_barcode.txt"
pool1_k562_barc_activ_f = "POOL1__pMPRA1__K562__activities_per_barcode.txt"

pool1_hela_elem_activ_f = "POOL1__pMPRA1__HeLa__activities_per_element.txt"
pool1_hepg2_elem_activ_f = "POOL1__pMPRA1__HepG2__activities_per_element.txt"
pool1_k562_elem_activ_f = "POOL1__pMPRA1__K562__activities_per_element.txt"

pool1_hela_pvals_f = "POOL1__pMPRA1__HeLa__pvals.txt"
pool1_hepg2_pvals_f = "POOL1__pMPRA1__HepG2__pvals.txt"
pool1_k562_pvals_f = "POOL1__pMPRA1__K562__pvals.txt"


# In[5]:


pool1_nocmv_hela_barc_activ_f = "POOL1__pNoCMVMPRA1__HeLa__activities_per_barcode.txt"
pool1_nocmv_hepg2_barc_activ_f = "POOL1__pNoCMVMPRA1__HepG2__activities_per_barcode.txt"
pool1_nocmv_k562_barc_activ_f = "POOL1__pNoCMVMPRA1__K562__activities_per_barcode.txt"

pool1_nocmv_hela_elem_activ_f = "POOL1__pNoCMVMPRA1__HeLa__activities_per_element.txt"
pool1_nocmv_hepg2_elem_activ_f = "POOL1__pNoCMVMPRA1__HepG2__activities_per_element.txt"
pool1_nocmv_k562_elem_activ_f = "POOL1__pNoCMVMPRA1__K562__activities_per_element.txt"

pool1_nocmv_hela_pvals_f = "POOL1__pNoCMVMPRA1__HeLa__pvals.txt"
pool1_nocmv_hepg2_pvals_f = "POOL1__pNoCMVMPRA1__HepG2__pvals.txt"
pool1_nocmv_k562_pvals_f = "POOL1__pNoCMVMPRA1__K562__pvals.txt"


# In[6]:


pool2_hepg2_barc_activ_f = "POOL2__pMPRA1__HepG2__activities_per_barcode.txt"
pool2_k562_barc_activ_f = "POOL2__pMPRA1__K562__activities_per_barcode.txt"

pool2_hepg2_elem_activ_f = "POOL2__pMPRA1__HepG2__activities_per_element.txt"
pool2_k562_elem_activ_f = "POOL2__pMPRA1__K562__activities_per_element.txt"

pool2_hepg2_pvals_f = "POOL2__pMPRA1__HepG2__pvals.txt"
pool2_k562_pvals_f = "POOL2__pMPRA1__K562__pvals.txt"


# In[7]:


pool1_index_f = "%s/tss_oligo_pool.index.txt" % index_dir
pool2_index_f = "%s/dels_oligo_pool.index.txt" % index_dir


# In[8]:


annot_f = "../../misc/00__tss_properties/mpra_id_to_biotype_map.txt"
id_map_f = "../../misc/00__tss_properties/mpra_tss_detailed_info.txt"
enh_id_map_f = "../../misc/00__tss_properties/enhancer_id_map.txt"
sel_map_f = "../../misc/00__tss_properties/mpra_tss_selection_info.txt"
cage_exp_f = "../../misc/01__cage/All_TSS_and_enh.CAGE_grouped_exp.tissue_sp.txt"
rna_seq_exp_f = "../../misc/01__cage/Expression.all.cells.txt"


# In[9]:


pool1_phylop_f = "../../data/00__index/pool1_tss.phylop46way.txt"


# ## 1. import data

# In[10]:


pool1_index = pd.read_table(pool1_index_f, sep="\t")
pool2_index = pd.read_table(pool2_index_f, sep="\t")


# In[11]:


pool1_index_elem = pool1_index[["element", "oligo_type", "unique_id", "dupe_info", "SNP"]]
pool2_index_elem = pool2_index[["element", "oligo_type", "unique_id", "dupe_info", "SNP"]]

pool1_index_elem = pool1_index_elem.drop_duplicates()
pool2_index_elem = pool2_index_elem.drop_duplicates()


# In[12]:


annot = pd.read_table(annot_f, sep="\t")
annot.head()


# In[13]:


id_map = pd.read_table(id_map_f, sep="\t")
id_map.head()


# In[14]:


sel_map = pd.read_table(sel_map_f, sep="\t")
sel_map.head()


# In[15]:


enh_id_map = pd.read_table(enh_id_map_f, sep="\t")
enh_id_map.head()


# In[16]:


cage_exp = pd.read_table(cage_exp_f, sep="\t")
cage_exp.head()


# In[17]:


rna_seq_exp = pd.read_table(rna_seq_exp_f, sep="\t")
rna_seq_exp.head()


# In[18]:


pool1_phylop = pd.read_table(pool1_phylop_f, sep="\t", header=None)
cols = ["chr", "start", "end", "unique_id", "score", "strand", "length"]
cols.extend(list(np.arange(-80, 34)))
pool1_phylop.columns = cols
pool1_phylop.head()


# ### pool 1

# In[19]:


pool1_hela_elem_norm = pd.read_table("%s/%s" % (activ_dir, pool1_hela_elem_activ_f), sep="\t")
pool1_hepg2_elem_norm = pd.read_table("%s/%s" % (activ_dir, pool1_hepg2_elem_activ_f), sep="\t")
pool1_k562_elem_norm = pd.read_table("%s/%s" % (activ_dir, pool1_k562_elem_activ_f), sep="\t")
pool1_hepg2_elem_norm.head()


# In[20]:


pool1_hela_reps = [x for x in pool1_hela_elem_norm.columns if "rna_" in x]
pool1_hepg2_reps = [x for x in pool1_hepg2_elem_norm.columns if "rna_" in x]
pool1_k562_reps = [x for x in pool1_k562_elem_norm.columns if "rna_" in x]
pool1_hepg2_reps


# In[21]:


pool1_hela_barc_norm = pd.read_table("%s/%s" % (activ_dir, pool1_hela_barc_activ_f), sep="\t")
pool1_hepg2_barc_norm = pd.read_table("%s/%s" % (activ_dir, pool1_hepg2_barc_activ_f), sep="\t")
pool1_k562_barc_norm = pd.read_table("%s/%s" % (activ_dir, pool1_k562_barc_activ_f), sep="\t")
pool1_hepg2_barc_norm.head()


# In[22]:


pool1_hela_pvals = pd.read_table("%s/%s" % (pval_dir, pool1_hela_pvals_f), sep="\t")
pool1_hepg2_pvals = pd.read_table("%s/%s" % (pval_dir, pool1_hepg2_pvals_f), sep="\t")
pool1_k562_pvals = pd.read_table("%s/%s" % (pval_dir, pool1_k562_pvals_f), sep="\t")
pool1_hepg2_pvals.head()


# ### pool 2

# In[23]:


pool2_hepg2_elem_norm = pd.read_table("%s/%s" % (activ_dir, pool2_hepg2_elem_activ_f), sep="\t")
pool2_k562_elem_norm = pd.read_table("%s/%s" % (activ_dir, pool2_k562_elem_activ_f), sep="\t")
pool2_hepg2_elem_norm.head()


# In[24]:


pool2_hepg2_reps = [x for x in pool2_hepg2_elem_norm.columns if "rna_" in x]
pool2_k562_reps = [x for x in pool2_k562_elem_norm.columns if "rna_" in x]
pool2_hepg2_reps


# In[25]:


pool2_hepg2_barc_norm = pd.read_table("%s/%s" % (activ_dir, pool2_hepg2_barc_activ_f), sep="\t")
pool2_k562_barc_norm = pd.read_table("%s/%s" % (activ_dir, pool2_k562_barc_activ_f), sep="\t")
pool2_hepg2_barc_norm.head()


# In[26]:


pool2_hepg2_pvals = pd.read_table("%s/%s" % (pval_dir, pool2_hepg2_pvals_f), sep="\t")
pool2_k562_pvals = pd.read_table("%s/%s" % (pval_dir, pool2_k562_pvals_f), sep="\t")
pool2_hepg2_pvals.head()


# ## 2. merge with index

# ### pool 1

# In[27]:


wt_index = pool1_index_elem.merge(annot, left_on="unique_id", right_on="seqID")
wt_index = wt_index[wt_index["oligo_type"].isin(["WILDTYPE", "WILDTYPE_BUT_HAS_SNP"])]
wt_index.sample(5)


# In[28]:


wt_index.PromType2.value_counts()


# In[29]:


pool1_hela_elem_norm = pool1_hela_elem_norm.merge(pool1_index_elem, on=["unique_id", "element"], how="left")
pool1_hepg2_elem_norm = pool1_hepg2_elem_norm.merge(pool1_index_elem, on=["unique_id", "element"], how="left")
pool1_k562_elem_norm = pool1_k562_elem_norm.merge(pool1_index_elem, on=["unique_id", "element"], how="left")


# In[30]:


pool1_hela_barc_norm = pool1_hela_barc_norm.merge(pool1_index, left_on="barcode", right_on="barcode", how="left")
pool1_hepg2_barc_norm = pool1_hepg2_barc_norm.merge(pool1_index, left_on="barcode", right_on="barcode", how="left")
pool1_k562_barc_norm = pool1_k562_barc_norm.merge(pool1_index, left_on="barcode", right_on="barcode", how="left")


# In[31]:


pool1_hela_elem_norm["better_type"] = pool1_hela_elem_norm.apply(better_type, axis=1)
pool1_hepg2_elem_norm["better_type"] = pool1_hepg2_elem_norm.apply(better_type, axis=1)
pool1_k562_elem_norm["better_type"] = pool1_k562_elem_norm.apply(better_type, axis=1)


# In[32]:


pool1_hela_elem_norm = pool1_hela_elem_norm.merge(pool1_hela_pvals.drop("oligo_type", axis=1), on=["unique_id", "element"], how="left")
pool1_hepg2_elem_norm = pool1_hepg2_elem_norm.merge(pool1_hepg2_pvals.drop("oligo_type", axis=1), on=["unique_id", "element"], how="left")
pool1_k562_elem_norm = pool1_k562_elem_norm.merge(pool1_k562_pvals.drop("oligo_type", axis=1), on=["unique_id", "element"], how="left")
pool1_hepg2_elem_norm.head()


# ### pool 2

# In[33]:


pool2_hepg2_elem_norm = pool2_hepg2_elem_norm.merge(pool2_index_elem, on=["unique_id", "element"], how="left")
pool2_k562_elem_norm = pool2_k562_elem_norm.merge(pool2_index_elem, on=["unique_id", "element"], how="left")


# In[34]:


pool2_hepg2_barc_norm = pool2_hepg2_barc_norm.merge(pool2_index, left_on="barcode", right_on="barcode", how="left")
pool2_k562_barc_norm = pool2_k562_barc_norm.merge(pool2_index, left_on="barcode", right_on="barcode", how="left")


# In[35]:


pool2_hepg2_elem_norm["better_type"] = pool2_hepg2_elem_norm.apply(better_type, axis=1)
pool2_k562_elem_norm["better_type"] = pool2_k562_elem_norm.apply(better_type, axis=1)


# In[36]:


pool2_hepg2_elem_norm = pool2_hepg2_elem_norm.merge(pool2_hepg2_pvals.drop("oligo_type", axis=1), on=["unique_id", "element"], how="left")
pool2_k562_elem_norm = pool2_k562_elem_norm.merge(pool2_k562_pvals.drop("oligo_type", axis=1), on=["unique_id", "element"], how="left")
pool2_hepg2_elem_norm.head()


# ## 3. count significantly active/inactive tiles

# ### pool 1

# In[37]:


pool1_hela_elem_norm["overall_mean"] = pool1_hela_elem_norm[pool1_hela_reps].mean(axis=1)
pool1_hepg2_elem_norm["overall_mean"] = pool1_hepg2_elem_norm[pool1_hepg2_reps].mean(axis=1)
pool1_k562_elem_norm["overall_mean"] = pool1_k562_elem_norm[pool1_k562_reps].mean(axis=1)

pool1_hela_elem_norm["overall_median"] = pool1_hela_elem_norm[pool1_hela_reps].median(axis=1)
pool1_hepg2_elem_norm["overall_median"] = pool1_hepg2_elem_norm[pool1_hepg2_reps].median(axis=1)
pool1_k562_elem_norm["overall_median"] = pool1_k562_elem_norm[pool1_k562_reps].median(axis=1)


# In[38]:


for cell, df in zip(["HeLa", "HepG2", "K562"], [pool1_hela_elem_norm, pool1_hepg2_elem_norm, pool1_k562_elem_norm]):
    print("%s: combined class" % cell)
    print(df.combined_class.value_counts())
    print("")
    if cell == "HepG2":
        print("%s: downsampled class" % cell)
        print(df.downsamp_combined_class.value_counts())
        print("")


# ### pool 2

# In[39]:


pool2_hepg2_elem_norm["overall_mean"] = pool2_hepg2_elem_norm[pool2_hepg2_reps].mean(axis=1)
pool2_k562_elem_norm["overall_mean"] = pool2_k562_elem_norm[pool2_k562_reps].mean(axis=1)

pool2_hepg2_elem_norm["overall_median"] = pool2_hepg2_elem_norm[pool2_hepg2_reps].median(axis=1)
pool2_k562_elem_norm["overall_median"] = pool2_k562_elem_norm[pool2_k562_reps].median(axis=1)


# In[40]:


for cell, df in zip(["HepG2", "K562"], [pool2_hepg2_elem_norm, pool2_k562_elem_norm]):
    print("%s: combined class" % cell)
    print(df.combined_class.value_counts())
    print("")
    if cell == "HepG2":
        print("%s: downsampled class" % cell)
        print(df.downsamp_combined_class.value_counts())
        print("")


# ## 4. boxplots: neg ctrls vs reference

# In[41]:


pool1_hepg2_df = pool1_hepg2_elem_norm.merge(annot, left_on="unique_id", right_on="seqID", how="left")
pool1_hela_df = pool1_hela_elem_norm.merge(annot, left_on="unique_id", right_on="seqID", how="left")
pool1_k562_df = pool1_k562_elem_norm.merge(annot, left_on="unique_id", right_on="seqID", how="left")
pool1_hepg2_df.head()


# In[42]:


pool1_hepg2_df["oligo_reg"] = pool1_hepg2_df.unique_id.str.split("__", expand=True)[2]
pool1_hela_df["oligo_reg"] = pool1_hela_df.unique_id.str.split("__", expand=True)[2]
pool1_k562_df["oligo_reg"] = pool1_k562_df.unique_id.str.split("__", expand=True)[2]
pool1_hepg2_df.head()


# In[43]:


def add_neg_ctrl_promtype(row):
    if row["better_type"] == "RANDOM":
        return "random"
    elif row["better_type"] == "SCRAMBLED":
        return "scrambled"
    elif row["better_type"] == "CONTROL":
        return "control"
    else:
        return row["PromType2"]


# In[44]:


pool1_hepg2_df["PromType2"] = pool1_hepg2_df.apply(add_neg_ctrl_promtype, axis=1)
pool1_hela_df["PromType2"] = pool1_hela_df.apply(add_neg_ctrl_promtype, axis=1)
pool1_k562_df["PromType2"] = pool1_k562_df.apply(add_neg_ctrl_promtype, axis=1)
pool1_hepg2_df.sample(10)


# ### pool 1

# In[45]:


order = ["RANDOM", "SCRAMBLED", "WILDTYPE"]
palette = {"RANDOM": "gray", "SCRAMBLED": "gray", "WILDTYPE": "black"}

f, axarr = plt.subplots(3, sharex=True, sharey=False, figsize=(1.78, 5))
neg_control_plot(pool1_hela_elem_norm, order, palette, fontsize, "HeLa", axarr[0], None, "HeLa MPRA activity", 
                 False, False, False, None)
neg_control_plot(pool1_hepg2_elem_norm, order, palette, fontsize, "HepG2", axarr[1], None, "HepG2 MPRA activity", 
                 False, False, False, None)
neg_control_plot(pool1_k562_elem_norm, order, palette, fontsize, "K562", axarr[2], None, "K562 MPRA activity", 
                 False, False, False, None)
plt.tight_layout()


# ### pool 2

# In[46]:


f, axarr = plt.subplots(2, sharex=True, sharey=False, figsize=(1.78, 3.2))
neg_control_plot(pool2_hepg2_elem_norm, order, palette, fontsize, "HepG2", axarr[0], None, "HepG2 MPRA activity", 
                 False, False, False, None)
neg_control_plot(pool2_k562_elem_norm, order, palette, fontsize, "K562", axarr[1], None, "K562 MPRA activity", 
                 False, False, False, None)
plt.tight_layout()
f.savefig("Fig_S12.pdf", bbox_inches="tight", dpi="figure")


# ## 5. boxplots: across TSS classes

# In[47]:


palette = {"random": "gray", "scrambled": "gray", "Enhancer": sns.color_palette("deep")[1], 
           "intergenic": sns.color_palette("deep")[2], "protein_coding": sns.color_palette("deep")[5], 
           "div_lnc": sns.color_palette("deep")[3], "div_pc": sns.color_palette("deep")[0]}


# In[48]:


# random
order = ["random", "Enhancer", "intergenic", "div_lnc", "protein_coding", "div_pc"]
labels = ["random", "eRNAs", "lincRNAs", "div. lncRNAs", "mRNAs", "div. mRNAs"]

f, axarr = plt.subplots(3, sharex=True, sharey=False, figsize=(5.3, 8))
promtype_plot(pool1_hela_df, order, palette, labels, fontsize, "HeLa", axarr[0], None, 
              "HeLa MPRA activity", False, False, False, None, True)
promtype_plot(pool1_hepg2_df, order, palette, labels, fontsize, "HepG2", axarr[1], None, 
              "HepG2 MPRA activity", False, False, False, None, True)
promtype_plot(pool1_k562_df, order, palette, labels, fontsize, "K562", axarr[2], None, 
              "K562 MPRA activity", False, False, False, None, True)
plt.tight_layout()
f.savefig("FigS4.pdf", bbox_inches="tight", dpi="figure")


# In[49]:


# random
order = ["random", "Enhancer", "intergenic", "div_lnc", "protein_coding", "div_pc"]
labels = ["random", "eRNAs", "lincRNAs", "div. lncRNAs", "mRNAs", "div. mRNAs"]

f, axarr = plt.subplots(3, sharex=True, sharey=False, figsize=(5.3, 8))
promtype_plot(pool1_hela_df, order, palette, labels, fontsize, "HeLa", axarr[0], None, 
              "HeLa MPRA activity", False, False, False, None, False)
promtype_plot(pool1_hepg2_df, order, palette, labels, fontsize, "HepG2", axarr[1], None, 
              "HepG2 MPRA activity", False, False, False, None, False)
promtype_plot(pool1_k562_df, order, palette, labels, fontsize, "K562", axarr[2], None, 
              "K562 MPRA activity", False, False, False, None, False)
plt.tight_layout()
f.savefig("Fig1C.pdf", bbox_inches="tight", dpi="figure")


# ## expression-match

# In[50]:


def fix_enh_cage_id(row):
    if row.PromType2 == "Enhancer":
        if not pd.isnull(row.enhancer_id_x):
            return row.enhancer_id_x
        elif not pd.isnull(row.enhancer_id_y):
            return row.enhancer_id_y
        else:
            return np.nan
    else:
        return row.cage_id


# In[51]:


sel_map["cage_id"] = sel_map["TSS_id"]
sel_map = sel_map.merge(enh_id_map[["TSS_id_Pos", "enhancer_id"]],
                        left_on="TSS_id", right_on="TSS_id_Pos", how="left")
sel_map = sel_map.merge(enh_id_map[["TSS_id_Neg", "enhancer_id"]],
                        left_on="TSS_id", right_on="TSS_id_Neg", how="left")
sel_map["cage_id"] = sel_map.apply(fix_enh_cage_id, axis=1)
sel_map.drop(["TSS_id_Pos", "enhancer_id_x", "TSS_id_Neg", "enhancer_id_y"], axis=1, inplace=True)
sel_map_expr = sel_map.merge(cage_exp, on="cage_id", how="left")
sel_map_expr.sample(5)


# In[52]:


sel_map_expr["log_av_exp"] = np.log10(sel_map_expr["av_exp"])
sel_map_expr.selected.unique()


# In[53]:


rand_sel_types = ["lncRNA", "eRNA.random", "mRNA.random", "mRNA.bidirec70-160"]


# In[54]:


rand_sel_ids = sel_map_expr[sel_map_expr["selected"].isin(rand_sel_types)]


# In[55]:


rand_sel_ids.PromType2.value_counts()


# In[56]:


def get_matching_pairs(df_1, df_2, col, scaler=True):

    x_1 = np.asarray(df_1[col])
    x_2 = np.asarray(df_2[col])
    x_1 = np.reshape(x_1, (len(df_1), 1))
    x_2 = np.reshape(x_2, (len(df_2), 1))

    if scaler == True:
        scaler = StandardScaler()
    if scaler:
        scaler.fit(x_2)
        x_2 = scaler.transform(x_2)
        x_1 = scaler.transform(x_1)
        
    nbrs = NearestNeighbors(n_neighbors=1).fit(x_2)
    distances, indices = nbrs.kneighbors(x_1)
    indices = indices.reshape(indices.shape[0])
    matched = df_2.ix[indices]
    return matched


# In[57]:


# match every biotype to the selected lncRNAs
promtypes = ["Enhancer", "protein_coding", "div_lnc", "div_pc"]
lncRNAs = sel_map_expr[(sel_map_expr["selected"] == "lncRNA") & 
                       (sel_map_expr["PromType2"] == "intergenic")].drop_duplicates()

print("total selected lncRNAs: %s" % len(lncRNAs))
print("min lncRNA expression: %s" % np.min(lncRNAs["av_exp"]))
print("max lncRNA expression: %s" % np.max(lncRNAs["av_exp"]))

all_matched = lncRNAs.copy()
for promtype in promtypes:
    sub_df = sel_map_expr[sel_map_expr["PromType2"] == promtype].drop_duplicates().reset_index()
    matched = get_matching_pairs(lncRNAs, sub_df, col="log_av_exp")
    print("%s (total=%s, matched=%s)" % (promtype, len(sub_df), len(matched)))
    all_matched = all_matched.append(matched)
all_matched = all_matched.drop_duplicates()


# In[58]:


all_matched.PromType2.value_counts()


# In[59]:


label_dict = {"Enhancer": "eRNAs", "intergenic": "lincRNAs", "div_lnc": "div. lncRNAs", "protein_coding": "mRNAs",
              "div_pc": "div. mRNAs"}
distplot_biotypes(rand_sel_ids, (3, 2.5), palette, label_dict, (0, 1.01), "log10(average CAGE expression)", False, None)


# In[60]:


distplot_biotypes(all_matched, (3, 2.5), palette, label_dict, (0, 1.01), "log10(average CAGE expression)", False, None)


# In[61]:


pool1_hela_rand = pool1_hela_df[pool1_hela_df["oligo_reg"].isin(rand_sel_ids["oligo_reg"])]
pool1_hepg2_rand = pool1_hepg2_df[pool1_hepg2_df["oligo_reg"].isin(rand_sel_ids["oligo_reg"])]
pool1_k562_rand = pool1_k562_df[pool1_k562_df["oligo_reg"].isin(rand_sel_ids["oligo_reg"])]


# In[63]:


order = ["Enhancer", "intergenic", "div_lnc", "protein_coding", "div_pc"]
labels = ["eRNAs", "lincRNAs", "div. lncRNAs", "mRNAs", "div. mRNAs"]

f, axarr = plt.subplots(3, sharex=True, sharey=False, figsize=(4, 8))
promtype_plot(pool1_hela_rand, order, palette, labels, fontsize, "HeLa", axarr[0], None, 
              "HeLa MPRA activity", False, False, False, None, False)
promtype_plot(pool1_hepg2_rand, order, palette, labels, fontsize, "HepG2", axarr[1], None, 
              "HepG2 MPRA activity", False, False, False, None, False)
promtype_plot(pool1_k562_rand, order, palette, labels, fontsize, "K562", axarr[2], None, 
              "K562 MPRA activity", False, False, False, None, False)
plt.tight_layout()
f.savefig("FigS6_1.pdf", bbox_inches="tight", dpi="figure")


# In[64]:


pool1_hela_exp = pool1_hela_df[pool1_hela_df["oligo_reg"].isin(all_matched["oligo_reg"])]
pool1_hepg2_exp = pool1_hepg2_df[pool1_hepg2_df["oligo_reg"].isin(all_matched["oligo_reg"])]
pool1_k562_exp = pool1_k562_df[pool1_k562_df["oligo_reg"].isin(all_matched["oligo_reg"])]


# In[65]:


f, axarr = plt.subplots(3, sharex=True, sharey=False, figsize=(4, 8))
promtype_plot(pool1_hela_exp, order, palette, labels, 
              fontsize, "HeLa", axarr[0], None, "HeLa MPRA activity", False, False, False, None, False)
promtype_plot(pool1_hepg2_exp, order, palette, labels, 
              fontsize, "HepG2", axarr[1], None, "HepG2 MPRA activity", False, False, False, None, False)
promtype_plot(pool1_k562_exp, order, palette, labels, 
              fontsize, "K562", axarr[2], None, "K562 MPRA activity", False, False, False, None, False)
plt.tight_layout()
f.savefig("FigS6_2.pdf", bbox_inches="tight", dpi="figure")


# the rest of the analysis only uses pool 1 (the TSS pool), as it looks at patterns in expression differences between TSS classes

# ## 6. barplots: find % of sequences active across cell types

# In[66]:


pool1_hela_df["cell"] = "HeLa"
pool1_hepg2_df["cell"] = "HepG2"
pool1_k562_df["cell"] = "K562"

all_df = pool1_hela_df[["unique_id", "better_type", "cell", "PromType2", "combined_class", "overall_mean"]].append(pool1_hepg2_df[["unique_id", "better_type", "cell", "PromType2", "combined_class", "overall_mean"]]).append(pool1_k562_df[["unique_id", "better_type", "cell", "PromType2", "combined_class", "overall_mean"]])


# In[67]:


df = all_df[all_df["better_type"] == "WILDTYPE"]
activ_grp = df.groupby("unique_id")["cell", "combined_class"].agg(lambda x: list(x)).reset_index()
activ_grp = activ_grp.merge(annot, left_on="unique_id", right_on="seqID", how="left").drop("seqID", axis=1)
activ_grp = activ_grp[(activ_grp["PromType2"].isin(TSS_CLASS_ORDER)) & 
                      ~(activ_grp["unique_id"].str.contains("SCRAMBLED"))]
activ_grp.sample(10)


# In[68]:


activ_grp["active_in_only_one"] = activ_grp.apply(active_in_only_one, axis=1)
activ_grp["active_in_only_two"] = activ_grp.apply(active_in_only_two, axis=1)
activ_grp["active_in_only_three"] = activ_grp.apply(active_in_only_three, axis=1)
activ_grp.sample(5)


# In[69]:


for PromType2 in TSS_CLASS_ORDER:
    df = activ_grp[activ_grp["PromType2"] == PromType2]
    active_in_1 = len(df[df["active_in_only_one"]])
    active_in_3 = len(df[df["active_in_only_three"]])
    print("%s | active in 1: %s, active in 3: %s" % (PromType2, active_in_1, active_in_3))


# In[70]:


activ_counts_1 = activ_grp.groupby(["PromType2", "active_in_only_one"])["unique_id"].agg("count").reset_index()
activ_pcts_1 = activ_counts_1.groupby("PromType2")["unique_id"].apply(lambda x: 100 * x / float(x.sum()))
activ_counts_1["percent"] = activ_pcts_1

activ_counts_2 = activ_grp.groupby(["PromType2", "active_in_only_two"])["unique_id"].agg("count").reset_index()
activ_pcts_2 = activ_counts_2.groupby("PromType2")["unique_id"].apply(lambda x: 100 * x / float(x.sum()))
activ_counts_2["percent"] = activ_pcts_2

activ_counts_3 = activ_grp.groupby(["PromType2", "active_in_only_three"])["unique_id"].agg("count").reset_index()
activ_pcts_3 = activ_counts_3.groupby("PromType2")["unique_id"].apply(lambda x: 100 * x / float(x.sum()))
activ_counts_3["percent"] = activ_pcts_3

activ_counts_1 = activ_counts_1[activ_counts_1["active_in_only_one"]]
activ_counts_2 = activ_counts_2[activ_counts_2["active_in_only_two"]]
activ_counts_3 = activ_counts_3[activ_counts_3["active_in_only_three"]]

activ_counts = activ_counts_1.merge(activ_counts_2, on="PromType2").merge(activ_counts_3, on="PromType2")
activ_counts.drop(["active_in_only_one", "unique_id_x", "active_in_only_two", "unique_id_y", 
                   "active_in_only_three", "unique_id"],
                  axis=1, inplace=True)
activ_counts.columns = ["PromType2", "active_in_only_one", "active_in_only_two", "active_in_only_three"]
activ_counts = pd.melt(activ_counts, id_vars="PromType2")
activ_counts.head()


# In[71]:


df = activ_counts[activ_counts["PromType2"] != "antisense"]
df["PromType2"] = pd.Categorical(df["PromType2"], TSS_CLASS_ORDER)
df.sort_values(by="PromType2")

plt.figure(figsize=(3.56, 2.3))
ax = sns.barplot(data=df, x="variable", y="value", hue="PromType2", ci=None, palette=TSS_CLASS_PALETTE)
ax.set_xticklabels(["active in 1", "active in 2", "active in 3"], rotation=30)

plt.legend(bbox_to_anchor=(1.35, 1))
plt.ylim((0, 50))
plt.ylabel("percent of sequences", size=fontsize)
plt.xlabel("")
plt.title("% of elements active in # of cell types")


# In[72]:


colors = []
for c in TSS_CLASS_ORDER:
    colors.append(TSS_CLASS_PALETTE[c])
colors


# In[73]:


# better plot showing tissue sp
df = activ_counts[activ_counts["PromType2"] != "antisense"]
df["PromType2"] = pd.Categorical(df["PromType2"], TSS_CLASS_ORDER)
df.sort_values(by="PromType2")

plt.figure(figsize=(3,2.3))
ax = sns.barplot(data=df[df["variable"]!="active_in_only_two"], x="PromType2", y="value", 
                 ci=None, hue="variable", linewidth=1.5)
ax.set_xticklabels(["eRNAs", "lincRNAs", "div. lncRNAs", "mRNAs", "div. mRNAs"], rotation=30)

colors = colors*2
for i, p in enumerate(ax.patches):
    if i < 5:
        p.set_facecolor(colors[i])
    else:
        p.set_facecolor("white")
        p.set_edgecolor(colors[i])
        p.set_alpha(1)
        p.set_hatch("///")

ax.legend().set_visible(False)
plt.ylim((0, 50))
plt.ylabel("% of sequences that are active\nin 1 and 3 cell types", fontsize=fontsize)
plt.xlabel("")
plt.savefig("Fig_1E.pdf", bbox_inches="tight", dpi="figure")


# ## 7. kdeplot: compare to CAGE

# In[74]:


hepg2_activ = pool1_hepg2_df[["unique_id", "element", "better_type", "overall_mean", "PromType2"]]
hela_activ = pool1_hela_df[["unique_id", "element", "better_type", "overall_mean"]]
k562_activ = pool1_k562_df[["unique_id", "element", "better_type", "overall_mean"]]

all_activ = hepg2_activ.merge(hela_activ, on=["unique_id", "element", "better_type"], how="left").merge(k562_activ, on=["unique_id", "element", "better_type"], how="left")
all_activ.columns = ["unique_id", "element", "better_type", "HepG2", "PromType2", "HeLa", "K562"]
all_activ = all_activ[["unique_id", "element", "better_type", "PromType2", "HepG2", "HeLa", "K562"]]
all_activ = all_activ[(all_activ["PromType2"].isin(TSS_CLASS_ORDER)) & 
                      ~(all_activ["unique_id"].str.contains("SCRAMBLED")) &
                      (all_activ["better_type"] == "WILDTYPE")]
all_activ.sample(5)


# In[75]:


all_activ["combined_class"] = ""
all_activ = all_activ.merge(pool1_hela_elem_norm[["unique_id", "element", "combined_class"]], on=["unique_id", "element"], how="left", suffixes=("", "_HeLa")).merge(pool1_hepg2_elem_norm[["unique_id", "element", "combined_class"]], on=["unique_id", "element"], how="left", suffixes=("", "_HepG2")).merge(pool1_k562_elem_norm[["unique_id", "element", "combined_class"]], on=["unique_id", "element"], how="left", suffixes=("", "_K562"))
all_activ.drop("combined_class", axis=1, inplace=True)
all_activ.head()


# In[76]:


all_activ["oligo_reg"] = all_activ.unique_id.str.split("__", expand=True)[2]
all_activ.sample(5)


# In[77]:


id_map = id_map[["oligo_reg", "gene_id", "K562_rep1", "K562_rep2", "K562_rep3", "HeLa_rep1", "HeLa_rep2", "HeLa_rep3", 
                 "HepG2_rep1", "HepG2_rep2", "HepG2_rep3"]]
all_activ = all_activ.merge(id_map, on="oligo_reg")
all_activ.sample(5)


# In[78]:


all_activ["K562_av"] = all_activ[["K562_rep1", "K562_rep2", "K562_rep3"]].mean(axis=1)
all_activ["HeLa_av"] = all_activ[["HeLa_rep1", "HeLa_rep2", "HeLa_rep3"]].mean(axis=1)
all_activ["HepG2_av"] = all_activ[["HepG2_rep1", "HepG2_rep2", "HepG2_rep3"]].mean(axis=1)

all_activ["K562_log_av"] = np.log10(all_activ["K562_av"]+1)
all_activ["HeLa_log_av"] = np.log10(all_activ["HeLa_av"]+1)
all_activ["HepG2_log_av"] = np.log10(all_activ["HepG2_av"]+1)


# In[79]:


all_activ = all_activ[(~all_activ["unique_id"].str.contains("SNP_INDIV")) & 
                      (~all_activ["unique_id"].str.contains("SNP_PLUS_HAPLO")) & 
                      (~all_activ["unique_id"].str.contains("FLIPPED"))]
all_activ.sample(5)


# In[80]:


# first scale mpra ranges to be positive
all_activ["hepg2_scaled"] = scale_range(all_activ["HepG2"], 0, 100)
all_activ["hela_scaled"] = scale_range(all_activ["HeLa"], 0, 100)
all_activ["k562_scaled"] = scale_range(all_activ["K562"], 0, 100)


# In[81]:


cage_ts = calculate_tissue_specificity(all_activ[["HepG2_log_av", "K562_log_av", "HeLa_log_av"]])
all_activ["cage_activ"] = all_activ[["HepG2_log_av", "K562_log_av", "HeLa_log_av"]].mean(axis=1)
all_activ["cage_ts"] = cage_ts

mpra_ts = calculate_tissue_specificity(all_activ[["hepg2_scaled", "k562_scaled", "hela_scaled"]])
all_activ["mpra_activ"] = all_activ[["HepG2", "K562", "HeLa"]].mean(axis=1)
all_activ["mpra_ts"] = mpra_ts
all_activ.head()


# In[82]:


cmap = sns.light_palette("darkslategray", as_cmap=True)


# In[83]:


no_nan = all_activ[(~pd.isnull(all_activ["mpra_ts"])) & (~pd.isnull(all_activ["cage_ts"]))]
g = sns.jointplot(data=no_nan, x="cage_ts", y="mpra_ts", kind="kde", shade_lowest=False, size=2.3, space=0,
                  stat_func=None, cmap=cmap, color="darkslategrey")
g.ax_joint.axhline(y=0.2, color="black", linewidth=1, linestyle="dashed")
g.ax_joint.axvline(x=0.5, color="black", linewidth=1, linestyle="dashed")
g.set_axis_labels("CAGE cell-type specificity", "MPRA cell-type specificity")
r, p = stats.spearmanr(no_nan["cage_ts"], no_nan["mpra_ts"])
g.ax_joint.annotate("r = {:.2f}\np = {:.2e}".format(r, Decimal(p)), xy=(.1, .75), xycoords=ax.transAxes, 
                    fontsize=5)
g.ax_joint.annotate("n = %s" % len(no_nan), xy=(.5, .8), xycoords=ax.transAxes, 
                    fontsize=5)
g.savefig("Fig_1D.pdf", bbox_inches="tight", dpi="figure")


# In[84]:


def cage_v_mpra_ts(row):
    if row["cage_ts"] > 0.5 and row["mpra_ts"] > 0.2:
        return "ts in both"
    elif row["cage_ts"] > 0.5 and row["mpra_ts"] <= 0.2:
        return "ts in cage, not mpra"
    elif row["cage_ts"] <= 0.5 and row["mpra_ts"] > 0.2:
        return "ts in mpra, not cage"
    else:
        return "not ts in both"
    
no_nan["ts_status"] = no_nan.apply(cage_v_mpra_ts, axis=1)
no_nan.ts_status.value_counts()


# In[85]:


tot = 692+402+310+236
upper_left = 310
upper_right = 402
lower_left = 692
lower_right = 236
print("upper left: %s" % (upper_left/tot))
print("upper right: %s" % (upper_right/tot))
print("lower left: %s" % (lower_left/tot))
print("lower right: %s" % (lower_right/tot))


# In[86]:


(692+402)/(692+402+310+236)


# In[97]:


no_nan = all_activ[(~pd.isnull(all_activ["mpra_activ"])) & (~pd.isnull(all_activ["cage_activ"]))]
g = sns.jointplot(data=no_nan, x="cage_activ", y="mpra_activ", kind="kde", shade_lowest=False, size=2.3, space=0,
                  stat_func=None, xlim=(-0.4, 0.9), ylim=(-3.5, 3), cmap=cmap, color="darkslategray")
g.set_axis_labels("mean log10(CAGE expression)", "mean MPRA activity")
r, p = stats.spearmanr(no_nan["cage_activ"], no_nan["mpra_activ"])
g.ax_joint.annotate("r = {:.2f}\np = {:.2e}".format(r, Decimal(p)), xy=(.06, .75), xycoords=ax.transAxes, 
                    fontsize=5)
g.ax_joint.annotate("n = %s" % len(no_nan), xy=(.48, .8), xycoords=ax.transAxes, 
                    fontsize=5)
g.savefig("Fig_S5.pdf", bbox_inches="tight", dpi="figure")


# ## 8. compare MPRA and CAGE to RNA-seq

# In[88]:


rna_seq_exp = rna_seq_exp[["gene_id", "HepG2", "HeLa-S3", "K562"]]
rna_seq_exp.columns = ["gene_id", "HepG2_rna_seq", "HeLa_rna_seq", "K562_rna_seq"]
rna_seq_exp["HepG2_rna_seq_log"] = np.log10(rna_seq_exp["HepG2_rna_seq"]+1)
rna_seq_exp["HeLa_rna_seq_log"] = np.log10(rna_seq_exp["HeLa_rna_seq"]+1)
rna_seq_exp["K562_rna_seq_log"] = np.log10(rna_seq_exp["K562_rna_seq"]+1)
all_activ_rna_seq = all_activ.merge(rna_seq_exp, on="gene_id")
all_activ_rna_seq.sample(5)


# In[89]:


all_activ_rna_seq.PromType2.value_counts()


# In[90]:


for cell in ["HepG2", "HeLa", "K562"]:
    mpra_col = cell
    seq_col = "%s_rna_seq_log" % cell
    
    no_nan = all_activ_rna_seq[(~pd.isnull(all_activ_rna_seq[mpra_col])) & (~pd.isnull(all_activ_rna_seq[seq_col]))]
    g = sns.jointplot(data=no_nan, x=mpra_col, y=seq_col, kind="kde", shade_lowest=False, size=2.3, space=0,
                      stat_func=None, cmap=cmap, color="darkslategray", ylim=(-0.5, 3))
    g.set_axis_labels("%s MPRA activity" % cell, "log10(%s RNA-seq expression)" % cell)
    r, p = stats.spearmanr(no_nan[mpra_col], no_nan[seq_col])
    g.ax_joint.annotate("r = {:.2f}\np = {:.2e}".format(r, Decimal(p)), xy=(.1, .75), xycoords=ax.transAxes, 
                        fontsize=5)
    
    # add n-value
    g.ax_joint.annotate("n = %s" % len(no_nan), xy=(.475, .8), xycoords=ax.transAxes, 
                        fontsize=5)
    
    plt.show()
    #g.savefig("%s_mpra_v_seq.pdf" % cell, bbox_inches="tight", dpi="figure")


# In[91]:


for cell in ["HepG2", "HeLa", "K562"]:
    cage_col = "%s_log_av" % cell
    seq_col = "%s_rna_seq_log" % cell
    
    no_nan = all_activ_rna_seq[(~pd.isnull(all_activ_rna_seq[cage_col])) & (~pd.isnull(all_activ_rna_seq[seq_col]))]
    g = sns.jointplot(data=no_nan, x=cage_col, y=seq_col, kind="kde", shade_lowest=False, size=2.3, space=0,
                      stat_func=None, cmap=cmap, color="darkslategray", ylim=(-0.5, 2.5), xlim=(-0.5, 2))
    g.set_axis_labels("log10(%s CAGE expression)" % cell, "log10(%s RNA-seq expression)" % cell)
    r, p = stats.spearmanr(no_nan[cage_col], no_nan[seq_col])
    g.ax_joint.annotate("r = {:.2f}\np = {:.2e}".format(r, Decimal(p)), xy=(.1, .75), xycoords=ax.transAxes, 
                        fontsize=5)
    
    # add n-value
    g.ax_joint.annotate("n = %s" % len(no_nan), xy=(.475, .8), xycoords=ax.transAxes, 
                        fontsize=5)
    
    plt.show()
    #g.savefig("%s_cage_v_seq.pdf" % cell, bbox_inches="tight", dpi="figure")


# ## 10. write final files

# In[92]:


# write file with tissue-specificities for later use
final = all_activ[["unique_id", "PromType2", "cage_activ", "cage_ts", "mpra_activ", "mpra_ts"]]
final.to_csv("../../data/02__activs/POOL1__pMPRA1__CAGE_vs_MPRA_activs.txt", sep="\t", index=False)


# In[93]:


# also write file with tss types
sel_map_expr.to_csv("../../misc/00__tss_properties/CAGE_expr_properties.txt", sep="\t", index=False)


# In[ ]:
```
