## Supplementary material for "High-throughput functional analysis of IncRNA core promoters elucidatesrules governing tissue-specificity": 05__divergent.py

```
# coding: utf-8

# # 05__divergent
# # analyzing divergent sequences: conservation and directionality preferences
# 
# in this notebook, i analyze two properties of divergent sequences: (1) their conservation (using phylop 46-way placental mammal alignments) and (2) their "directionality preference". by that i mean whether or not they are more likely to activate transcription in the sense direction vs. the antisense direction.
# 
# ------
# 
# no figures in this notebook

# In[1]:


import warnings
warnings.filterwarnings('ignore')

import pandas as pd
import numpy as np
import matplotlib as mpl
import matplotlib.pyplot as plt
import seaborn as sns
import sys
import time

from itertools import chain
from decimal import Decimal
from scipy import stats
from scipy import signal
from statsmodels.sandbox.stats import multicomp

# import utils
sys.path.append("../../utils")
from plotting_utils import *
from misc_utils import *
from norm_utils import *

get_ipython().magic('matplotlib inline')


# In[2]:


sns.set(**PAPER_PRESET)
fontsize = PAPER_FONTSIZE


# ## functions

# In[3]:


def padj_in_one(row):
    if not pd.isnull(row["hepg2_sig"]):
        if not pd.isnull(row["hepg2_sig"]):
            return True
    elif not pd.isnull(row["hela_sig"]):
        if not pd.isnull(row["hela_sig"]):
            return True
    elif not pd.isnull(row["k562_sig"]):
        if not pd.isnull(row["k562_sig"]):
            return True
    else:
        return False


# In[4]:


def sig_in_one(row):
    if row["hepg2_sig"] == "sig" or row["k562_sig"] == "sig" or row["hela_sig"] == "sig":
        return True
    else:
        return False


# ## variables

# In[5]:


biotypes = ["div_lnc", "div_pc", "enhancerMid", "intergenic", "protein_coding"]


# In[6]:


# note - this file will have to be changed if re-running outside of harvard odyssey as it's not in the git (large)
bw_f = "/n/rinn_data2/users/kaia/bigwig/all_chr.phyloP46way.placental.bw"


# In[7]:


index_dir = "../../data/00__index"
index_f = "%s/tss_oligo_pool.index.txt" % index_dir


# In[8]:


hepg2_activ_f = "../../data/02__activs/POOL1__pMPRA1__HepG2__activities_per_barcode.txt"
hela_activ_f = "../../data/02__activs/POOL1__pMPRA1__HeLa__activities_per_barcode.txt"
k562_activ_f = "../../data/02__activs/POOL1__pMPRA1__K562__activities_per_barcode.txt"


# In[9]:


annot_f = "../../misc/00__tss_properties/mpra_id_to_biotype_map.txt"


# # conservation

# ## 1. find phyloP scores for TSS +/- 1000

# In[10]:


in_path = "../../data/00__index/0__all_tss"
out_path = "tmp"
get_ipython().system('mkdir -p $out_path')
up = 1000
down = 1000

for b in biotypes:
    bed_f = "%s/TSS.start.1perGencGene.500bp.%s.bed" % (in_path, b)
    phylop_f = "%s/TSS.start.1perGencGene.500bp.%s.%sup%sdown.phylop.txt" % (out_path, b, up, down)
    get_ipython().system('bwtool matrix $up:$down $bed_f $bw_f $phylop_f')


# ## 2. read files and find column avg

# In[11]:


data = pd.DataFrame()

up = 1000
down = 1000

for b in biotypes:
    phylop_f = "%s/TSS.start.1perGencGene.500bp.%s.%sup%sdown.phylop.txt" % (out_path, b, up, down)
    tmp = pd.read_table(phylop_f, sep="\t", header=None)
    avg = list(tmp.mean(axis=0))
    ste = list(tmp.std(axis=0)/(np.sqrt(len(tmp))))
    n = len(tmp)
    tmp = pd.DataFrame(data={"mean": avg, "ste": ste})
    tmp["type"] = b
    tmp["num"] = n
    tmp = tmp.reset_index()
    tmp.columns = ["idx", "mean", "ste", "type", "num"]
    tmp["ntd"] = list(range(-1*up, down))
    data = data.append(tmp)

data["y1"] = data["mean"] - data["ste"]
data["y2"] = data["mean"] + data["ste"]
data.sample(5)


# ## 3. plot

# In[12]:


palette = {"intergenic": sns.color_palette()[2], "enhancerMid": sns.color_palette()[1],
           "div_pc": sns.color_palette()[0], "protein_coding": sns.color_palette()[5], 
           "div_lnc": sns.color_palette()[3]}


# In[13]:


fig = plt.figure(figsize=(7,2))

lower = -400
upper = 400

for b in biotypes:
    df = data[(data["type"] == b) & (data["ntd"] >= lower) & (data["ntd"] < upper)]
    x = signal.savgol_filter(df["mean"], 15, 1)
    plt.fill_between(df["ntd"], df["y1"], df["y2"], color=palette[b], alpha=0.5)
    plt.plot(df["ntd"], x, color=palette[b], linewidth=3, label="%s (%s)" % (b, df["num"].iloc[0]))
plt.xlim((lower, upper))
plt.axvline(x=-75, color="black", linestyle="dashed", linewidth=1)
plt.axvline(x=25, color="black", linestyle="dashed", linewidth=1)
plt.legend(ncol=1, loc=1, bbox_to_anchor=(1.25, 1))
plt.xlabel("nucleotide (0=TSS)")
plt.ylabel("phylop 100-way")


# In[14]:


order = ["enhancerMid", "intergenic", "div_lnc", "protein_coding", "div_pc"]


# In[15]:


data["log_mean"] = np.log(data["mean"]+1)
data.head()


# In[16]:


plt.figure(figsize=(2.5, 2.5))
ax = sns.boxplot(data=data, x="type", y="log_mean", order=order, palette=palette,
                 flierprops=dict(marker='o', markersize=5))
ax.set_xticklabels(["eRNAs", "lincRNAs", "div. lncRNAs", "mRNAs", "div. mRNAs"], rotation=30)
mimic_r_boxplot(ax)
ax.set_ylabel("log(mean(phyloP 100-way score))")
ax.set_xlabel("")


# # directionality preference

# ## 1. import data

# In[17]:


index = pd.read_table(index_f, sep="\t")


# In[18]:


index_elem = index[["element", "oligo_type", "unique_id", "dupe_info", "SNP", "seq_name"]]
index_elem = index_elem.drop_duplicates()


# In[19]:


hepg2_activ = pd.read_table(hepg2_activ_f, sep="\t")
hela_activ = pd.read_table(hela_activ_f, sep="\t")
k562_activ = pd.read_table(k562_activ_f, sep="\t")


# In[20]:


hepg2_reps = [x for x in hepg2_activ.columns if "rna" in x]
hela_reps = [x for x in hela_activ.columns if "rna" in x]
k562_reps = [x for x in k562_activ.columns if "rna" in x]


# In[21]:


annot = pd.read_table(annot_f, sep="\t")
annot.head()


# ## 2. find mean activ per barcode and merge info

# In[22]:


hepg2_cols = ["barcode"] + ["hepg2_%s" % x for x in hepg2_reps]
k562_cols = ["barcode"] + ["k562_%s" % x for x in k562_reps]
hela_cols = ["barcode"] + ["hela_%s" % x for x in hela_reps]
hela_cols


# In[23]:


hepg2_activ.columns = hepg2_cols
k562_activ.columns = k562_cols
hela_activ.columns = hela_cols
hela_activ.head()


# In[24]:


data = index.merge(hepg2_activ[hepg2_cols], on="barcode").merge(hela_activ[hela_cols], on="barcode").merge(k562_activ[k562_cols], on="barcode")
data.sample(5)


# ## 3. make map of flipped to not flipped

# In[25]:


index_elem["cage_id"] = index_elem.apply(get_cage_id, axis=1)
index_elem.sample(5)


# In[26]:


flip_map = {}
for cage_id in list(index_elem["cage_id"].unique()):
    if cage_id == "none":
        continue
    unique_seqs = index_elem[index_elem["cage_id"] == cage_id].seq_name.unique()
    if len(unique_seqs) == 1:
        continue
    else:
        flip_map[unique_seqs[0]] = unique_seqs[1]


# ## 4. calculate differences b/w flipped and not flipped
# require at least 10 barcodes in each

# In[27]:


hepg2_results = {}
hela_results = {}
k562_results = {}

for sense in flip_map.keys():
    antisense = flip_map[sense]
    
    for results_dict, reps in zip([hepg2_results, k562_results, hela_results], [hepg2_cols, k562_cols, hela_cols]):
        rep_results = {"antisense": antisense}
        for rep in reps:
            if rep == "barcode":
                continue

            # extract sense/antisense barcode values for each cell type
            sense_vals = data[(data["seq_name"] == sense) & 
                              (data["oligo_type"].isin(["WILDTYPE", "WILDTYPE_BUT_HAS_SNP"]))][rep]
            antisense_vals = data[(data["seq_name"] == antisense) & 
                                  (data["oligo_type"].isin(["FLIPPED"]))][rep]
            
            # get non-nan values
            sense_vals = np.asarray(sense_vals)
            antisense_vals = np.asarray(antisense_vals)
            sense_nonan_vals = sense_vals[~np.isnan(sense_vals)]
            antisense_nonan_vals = antisense_vals[~np.isnan(antisense_vals)]

            # find "log2fc" for antisense/sense (as diff b/w medians)
            sense_median = np.nanmedian(sense_nonan_vals)
            antisense_median = np.nanmedian(antisense_nonan_vals)
            l2fc = antisense_median - sense_median

            # perform wilcoxon rank sum test
            if len(sense_nonan_vals) >= 10 and len(antisense_nonan_vals) >= 10:
                u, pval = stats.mannwhitneyu(sense_nonan_vals, antisense_nonan_vals, alternative="two-sided", use_continuity=False)
            else:
                u, pval = "NA__not_enough_barcodes", "NA__not_enough_barcodes"
                
            tmp = {"%s_l2fc" % rep: l2fc, "%s_pval" % rep: pval}
            rep_results.update(tmp)
        # add results to dicts
        results_dict[sense] = rep_results


# In[28]:


hepg2_results = pd.DataFrame.from_dict(data=hepg2_results, orient="index").reset_index()
hela_results = pd.DataFrame.from_dict(data=hela_results, orient="index").reset_index()
k562_results = pd.DataFrame.from_dict(data=k562_results, orient="index").reset_index()

hepg2_results.head()


# ## 5. combine and correct p-vals

# In[29]:


hepg2_pvals = [x for x in hepg2_results.columns if "_pval" in x and "combined" not in x]
hela_pvals = [x for x in hela_results.columns if "_pval" in x and "combined" not in x]
k562_pvals = [x for x in k562_results.columns if "_pval" in x and "combined" not in x]

hepg2_results["hepg2_combined_pval"] = hepg2_results.apply(combine_pvals, reps=hepg2_pvals, axis=1)
hela_results["hela_combined_pval"] = hela_results.apply(combine_pvals, reps=hela_pvals, axis=1)
k562_results["k562_combined_pval"] = k562_results.apply(combine_pvals, reps=k562_pvals, axis=1)
k562_results.head()


# In[30]:


hepg2_reps = [x for x in hepg2_activ.columns if "rna" in x]
hela_reps = [x for x in hela_activ.columns if "rna" in x]
k562_reps = [x for x in k562_activ.columns if "rna" in x]


# In[31]:


all_dfs = []
for df, reps, cell in zip([hepg2_results, hela_results, k562_results], [hepg2_reps, hela_reps, k562_reps], ["hepg2", "hela", "k562"]):
    reps.extend(["%s_combined" % cell])
    print(cell)
    for rep in reps:
        print(rep)
        col = "%s_pval" % rep
        sub_df = df[~(df[col].astype(str).str.contains("NA")) & ~(pd.isnull(df[col]))][["index", "antisense", col]]

        new_pvals = multicomp.multipletests(sub_df[col], method="bonferroni")[1]
        sub_df["%s_padj" % (rep)] = new_pvals
        sub_df.drop(col, axis=1, inplace=True)

        df = df.merge(sub_df, on=["index", "antisense"], how="left")
    all_dfs.append(df)


# In[32]:


for cell, df in zip(["hepg2", "hela", "k562"], all_dfs):
    print(cell)
    l2fc_cols = [x for x in df.columns if "_l2fc" in x and "combined" not in x]
    print(l2fc_cols)
    comb_col = "%s_combined_padj" % (cell)
    print(comb_col)
    df["%s_sig" % cell] = df.apply(is_sig_combined, col=comb_col, thresh=0.05, l2fc_cols=l2fc_cols, axis=1)


# ## 6. put it all in 1 dataframe

# In[33]:


all_results = all_dfs[0].merge(all_dfs[1], on=["index", "antisense"], how="outer").merge(all_dfs[2], on=["index", "antisense"], how="outer")
all_results.head()


# In[34]:


all_results["padj_in_one"] = all_results.apply(padj_in_one, axis=1)
all_results.sample(5)


# ## 7. find out how many are significantly different in at least 1 cell type

# In[35]:


annot["sense"] = annot["seqID"].str.split("__", expand=True)[1] + "__" + annot["seqID"].str.split("__", expand=True)[2]
annot_wt = annot[~(annot["seqID"].str.contains("SNP_INDIV")) & ~(annot["seqID"].str.contains("HAPLO")) & ~(annot["seqID"].str.contains("SCRAMBLED"))]
annot_wt.head()


# In[36]:


all_results.head()


# In[37]:


sig_results = all_results[all_results["padj_in_one"]]
sig_results = sig_results.merge(annot_wt, left_on="index", right_on="sense", how="left")
sig_results.head()


# In[38]:


sig_results["sig_in_any"] = sig_results.apply(sig_in_one, axis=1)
sig_results.head()


# In[39]:


sig_results.sig_in_any.value_counts()


# ## 8. plot

# In[40]:


plt.figure(figsize=(3, 2))

ax = sns.countplot(data=sig_results, x="PromType2", color="lightgray", order=TSS_CLASS_ORDER)
sns.countplot(data=sig_results[sig_results["sig_in_any"]], x="PromType2", palette=TSS_CLASS_PALETTE, 
              order=TSS_CLASS_ORDER, ax=ax)
ax.set_xticklabels(["eRNAs", "lincRNAs", "div. lncRNAs", "mRNAs", "div. mRNAs"], rotation=30)

counts = []
tots = []
for i, p in enumerate(ax.patches):
    if i <= 4 :
        tots.append(p.get_height())
    else:
        counts.append(p.get_height())
    text = int(p.get_height())
    ax.annotate(text, (p.get_x() + p.get_width()/2., p.get_height()), 
                ha="center", va="bottom", size=fontsize)

for i in range(len(counts)):
    percent = (float(counts[i])/tots[i])*100
    text = "%.1f%%" % percent
    ax.annotate(text, (i, 5), ha="center", va="bottom", size=fontsize)

plt.xlabel("")
plt.ylim((0, 130))
plt.title("count of seqs that are sig. diff. (padj < 0.05)\nin either HepG2, HeLa, or K562")


# ## 9. remove tmp files

# In[41]:


get_ipython().system('rm tmp/*.txt')
get_ipython().system('rmdir tmp')


# In[ ]:
```
