## Supplementary material for "High-throughput functional analysis of IncRNA core promoters elucidatesrules governing tissue-specificity": 06__tf_tissue_sp.py

```
# coding: utf-8

# # 06__tf_tissue_sp
# # calculating tissue specificity of TFs (across HepG2, HeLa, and K562)
# 
# in this notebook, i calculate the tissue specificity of TFs across the 3 cell types in our MPRAs using ENCODE RNA-seq data. then, i correlate motif coverage (# bp covered and maximum coverage) as well as average TF specificity with MPRA activities and specificities.
# 
# note: the FIMO mappings and coverage calculations were done separately (see methods)
# 
# ------
# 
# figures in this notebook:
# - **Fig 2C**: KDE plot of correlations of MPRA activity & specificity with each of the 3 metrics

# In[1]:


import warnings
warnings.filterwarnings('ignore')

import pandas as pd
import numpy as np
import matplotlib as mpl
import matplotlib.pyplot as plt
import seaborn as sns
import sys
import time

from itertools import chain
from decimal import Decimal
from scipy import stats
from scipy import signal
from statsmodels.sandbox.stats import multicomp

# import utils
sys.path.append("../../utils")
from plotting_utils import *
from misc_utils import *
from norm_utils import *

get_ipython().magic('matplotlib inline')


# In[2]:


sns.set(**PAPER_PRESET)
fontsize = PAPER_FONTSIZE


# ## functions

# In[3]:


def fix_small_decimal(row):
    if row.max_cov < 1:
        return 0
    else:
        return row.max_cov


# In[4]:


def get_cage_id(row):
    if "Enhancer" in row.unique_id:
        return row.unique_id.split("__")[1]
    else:
        return row.unique_id.split("__")[2]


# ## variables

# In[5]:


index_dir = "../../data/00__index"
index_f = "%s/tss_oligo_pool.index.txt" % index_dir


# In[6]:


hepg2_activ_f = "../../data/02__activs/POOL1__pMPRA1__HepG2__activities_per_element.txt"
hela_activ_f = "../../data/02__activs/POOL1__pMPRA1__HeLa__activities_per_element.txt"
k562_activ_f = "../../data/02__activs/POOL1__pMPRA1__K562__activities_per_element.txt"


# In[7]:


fimo_f = "../../misc/03__fimo/00__fimo_outputs/all_fimo_map.new_deduped.txt.gz"
fimo_chip_f = "../../misc/03__fimo/00__fimo_outputs/all_fimo_map.new_chip_intersected.new_deduped.txt.gz"
pool1_fimo_f = "../../misc/03__fimo/00__fimo_outputs/pool1_fimo_map.new_deduped.txt"
pool1_fimo_chip_f = "../../misc/03__fimo/00__fimo_outputs/pool1_fimo_map.new_chip_intersected.new_deduped.txt"
pool1_fimo_no_ets_f = "../../misc/03__fimo/00__fimo_outputs/pool1_fimo_map.no_ETS_motifs.new_deduped.txt"
pool1_fimo_no_ets_chip_f = "../../misc/03__fimo/00__fimo_outputs/pool1_fimo_map.new_chip_intersected.no_ETS_motifs.new_deduped.txt"


# In[8]:


fimo_bp_cov_f = "../../data/04__coverage/all_fimo_map.new_deduped.bp_covered.txt"
fimo_max_cov_f = "../../data/04__coverage/all_fimo_map.new_deduped.max_coverage.txt"

fimo_chip_bp_cov_f = "../../data/04__coverage/all_fimo_map.new_chip_intersected.new_deduped.bp_covered.txt"
fimo_chip_max_cov_f = "../../data/04__coverage/all_fimo_map.new_chip_intersected.new_deduped.max_coverage.txt"

fimo_clust_bp_cov_f = "../../data/04__coverage/all_fimo_map.bulyk_clusters.new_deduped.bp_covered.txt"
fimo_clust_max_cov_f = "../../data/04__coverage/all_fimo_map.bulyk_clusters.new_deduped.max_coverage.txt"

fimo_mosbat_bp_cov_f = "../../data/04__coverage/all_fimo_map.mosbat_clusters.bp_covered.txt"
fimo_mosbat_max_cov_f = "../../data/04__coverage/all_fimo_map.mosbat_clusters.max_coverage.txt"


# In[9]:


fimo_no_ets_bp_cov_f = "../../data/04__coverage/all_fimo_map.no_ETS_motifs.new_deduped.bp_covered.txt"
fimo_no_ets_max_cov_f = "../../data/04__coverage/all_fimo_map.no_ETS_motifs.new_deduped.max_coverage.txt"

fimo_no_ets_chip_bp_cov_f = "../../data/04__coverage/all_fimo_map.new_chip_intersected.no_ETS_motifs.new_deduped.bp_covered.txt"
fimo_no_ets_chip_max_cov_f = "../../data/04__coverage/all_fimo_map.new_chip_intersected.no_ETS_motifs.new_deduped.max_coverage.txt"


# In[10]:


pool1_fimo_bp_cov_f = "../../data/04__coverage/pool1_fimo_map.new_deduped.bp_covered.txt"
pool1_fimo_max_cov_f = "../../data/04__coverage/pool1_fimo_map.new_deduped.max_coverage.txt"

pool1_fimo_chip_bp_cov_f = "../../data/04__coverage/pool1_fimo_map.new_chip_intersected.new_deduped.bp_covered.txt"
pool1_fimo_chip_max_cov_f = "../../data/04__coverage/pool1_fimo_map.new_chip_intersected.new_deduped.max_coverage.txt"


# In[11]:


pool1_fimo_no_ets_bp_cov_f = "../../data/04__coverage/pool1_fimo_map.no_ETS_motifs.new_deduped.bp_covered.txt"
pool1_fimo_no_ets_max_cov_f = "../../data/04__coverage/pool1_fimo_map.no_ETS_motifs.new_deduped.max_coverage.txt"

pool1_fimo_no_ets_chip_bp_cov_f = "../../data/04__coverage/pool1_fimo_map.new_chip_intersected.no_ETS_motifs.new_deduped.bp_covered.txt"
pool1_fimo_no_ets_chip_max_cov_f = "../../data/04__coverage/pool1_fimo_map.new_chip_intersected.no_ETS_motifs.new_deduped.max_coverage.txt"


# In[12]:


tf_ts_f = "../../data/04__coverage/TF_tissue_specificities.from_CAGE.txt"


# In[13]:


cage_v_mpra_f = "../../data/02__activs/POOL1__pMPRA1__CAGE_vs_MPRA_activs.txt"


# In[14]:


tss_cage_map_f = "../../misc/00__tss_properties/mpra_tss_detailed_info.txt"
enh_cage_map_f = "../../misc/00__tss_properties/enhancer_id_map.txt"


# ## 1. import data

# In[15]:


fimo = pd.read_table(fimo_f, sep="\t", header=None, compression="gzip")
fimo.columns = ["motif_chr", "motif_start", "motif_end", "unique_id", "score", "strand", "chr", "start", "end", 
                "motif", "motif_score", "motif_strand"]
fimo.head()


# In[16]:


fimo_chip = pd.read_table(fimo_chip_f, sep="\t", header=None, compression="gzip")
fimo_chip.columns = ["motif_chr", "motif_start", "motif_end", "unique_id", "score", "strand", "chr", "start", "end", 
                     "motif", "motif_score", "motif_strand"]


# In[17]:


pool1_fimo = pd.read_table(pool1_fimo_f, sep="\t", header=None)
pool1_fimo.columns = ["motif_chr", "motif_start", "motif_end", "unique_id", "score", "strand", "chr", "start", "end", 
                      "motif", "motif_score", "motif_strand"]


# In[18]:


pool1_fimo_chip = pd.read_table(pool1_fimo_chip_f, sep="\t", header=None)
pool1_fimo_chip.columns = ["motif_chr", "motif_start", "motif_end", "unique_id", "score", "strand", "chr", "start", 
                           "end", "motif", "motif_score", "motif_strand"]


# In[19]:


pool1_fimo_no_ets = pd.read_table(pool1_fimo_no_ets_f, sep="\t", header=None)
pool1_fimo_no_ets.columns = ["motif_chr", "motif_start", "motif_end", "unique_id", "score", "strand", "chr", "start", 
                             "end", "motif", "motif_score", "motif_strand"]


# In[20]:


pool1_fimo_no_ets_chip = pd.read_table(pool1_fimo_no_ets_chip_f, sep="\t", header=None)
pool1_fimo_no_ets_chip.columns = ["motif_chr", "motif_start", "motif_end", "unique_id", "score", "strand", "chr", 
                                  "start", "end", "motif", "motif_score", "motif_strand"]


# In[21]:


index = pd.read_table(index_f, sep="\t")
index_elem = index[["element", "oligo_type", "unique_id", "dupe_info", "SNP", "seq_name"]]
index_elem = index_elem.drop_duplicates()


# In[22]:


hepg2_activ = pd.read_table(hepg2_activ_f, sep="\t")
hela_activ = pd.read_table(hela_activ_f, sep="\t")
k562_activ = pd.read_table(k562_activ_f, sep="\t")


# In[23]:


hepg2_reps = [x for x in hepg2_activ.columns if "rna" in x]
hela_reps = [x for x in hela_activ.columns if "rna" in x]
k562_reps = [x for x in k562_activ.columns if "rna" in x]


# In[24]:


fimo_bp_cov = pd.read_table(fimo_bp_cov_f, sep="\t", header=None)
fimo_bp_cov.columns = ["chr", "start", "end", "unique_id", "score", "strand", "n_motifs", "n_bp_cov", "seq_len", 
                       "frac_bp_cov"]

fimo_max_cov = pd.read_table(fimo_max_cov_f, sep="\t", header=None)
fimo_max_cov.columns = ["unique_id", "max_cov"]

fimo_cov = fimo_bp_cov.merge(fimo_max_cov, on="unique_id")
print(len(fimo_cov))
fimo_cov.head()


# In[25]:


fimo_chip_bp_cov = pd.read_table(fimo_chip_bp_cov_f, sep="\t", header=None)
fimo_chip_bp_cov.columns = ["chr", "start", "end", "unique_id", "score", "strand", "n_motifs", "n_bp_cov", "seq_len", 
                            "frac_bp_cov"]

fimo_chip_max_cov = pd.read_table(fimo_chip_max_cov_f, sep="\t", header=None)
fimo_chip_max_cov.columns = ["unique_id", "max_cov"]

fimo_chip_cov = fimo_chip_bp_cov.merge(fimo_chip_max_cov, on="unique_id")
print(len(fimo_chip_cov))


# In[26]:


fimo_clust_bp_cov = pd.read_table(fimo_clust_bp_cov_f, sep="\t", header=None)
fimo_clust_bp_cov.columns = ["chr", "start", "end", "unique_id", "score", "strand", "n_motifs", "n_bp_cov", "seq_len", 
                             "frac_bp_cov"]

fimo_clust_max_cov = pd.read_table(fimo_clust_max_cov_f, sep="\t", header=None)
fimo_clust_max_cov.columns = ["unique_id", "max_cov"]

fimo_clust_cov = fimo_clust_bp_cov.merge(fimo_clust_max_cov, on="unique_id")
print(len(fimo_clust_cov))


# In[27]:


fimo_mosbat_bp_cov = pd.read_table(fimo_mosbat_bp_cov_f, sep="\t", header=None)
fimo_mosbat_bp_cov.columns = ["chr", "start", "end", "unique_id", "score", "strand", "n_motifs", "n_bp_cov", "seq_len", 
                             "frac_bp_cov"]

fimo_mosbat_max_cov = pd.read_table(fimo_mosbat_max_cov_f, sep="\t", header=None)
fimo_mosbat_max_cov.columns = ["unique_id", "max_cov"]

fimo_mosbat_cov = fimo_mosbat_bp_cov.merge(fimo_mosbat_max_cov, on="unique_id")
print(len(fimo_mosbat_cov))


# In[28]:


pool1_fimo_bp_cov = pd.read_table(pool1_fimo_bp_cov_f, sep="\t", header=None)
pool1_fimo_bp_cov.columns = ["chr", "start", "end", "unique_id", "score", "strand", "n_motifs", "n_bp_cov", "seq_len", 
                             "frac_bp_cov"]

pool1_fimo_max_cov = pd.read_table(pool1_fimo_max_cov_f, sep="\t", header=None)
pool1_fimo_max_cov.columns = ["unique_id", "max_cov"]

pool1_fimo_cov = pool1_fimo_bp_cov.merge(pool1_fimo_max_cov, on="unique_id")
print(len(pool1_fimo_cov))


# In[29]:


pool1_fimo_chip_bp_cov = pd.read_table(pool1_fimo_chip_bp_cov_f, sep="\t", header=None)
pool1_fimo_chip_bp_cov.columns = ["chr", "start", "end", "unique_id", "score", "strand", "n_motifs", "n_bp_cov", 
                                  "seq_len", "frac_bp_cov"]

pool1_fimo_chip_max_cov = pd.read_table(pool1_fimo_chip_max_cov_f, sep="\t", header=None)
pool1_fimo_chip_max_cov.columns = ["unique_id", "max_cov"]

pool1_fimo_chip_cov = pool1_fimo_chip_bp_cov.merge(pool1_fimo_chip_max_cov, on="unique_id")
print(len(pool1_fimo_chip_cov))


# In[30]:


pool1_fimo_no_ets_bp_cov = pd.read_table(pool1_fimo_no_ets_bp_cov_f, sep="\t", header=None)
pool1_fimo_no_ets_bp_cov.columns = ["chr", "start", "end", "unique_id", "score", "strand", "n_motifs", "n_bp_cov", 
                                    "seq_len", "frac_bp_cov"]

pool1_fimo_no_ets_max_cov = pd.read_table(pool1_fimo_no_ets_max_cov_f, sep="\t", header=None)
pool1_fimo_no_ets_max_cov.columns = ["unique_id", "max_cov"]

pool1_fimo_no_ets_cov = pool1_fimo_no_ets_bp_cov.merge(pool1_fimo_no_ets_max_cov, on="unique_id")
print(len(pool1_fimo_no_ets_cov))


# In[31]:


pool1_fimo_no_ets_chip_bp_cov = pd.read_table(pool1_fimo_no_ets_chip_bp_cov_f, sep="\t", header=None)
pool1_fimo_no_ets_chip_bp_cov.columns = ["chr", "start", "end", "unique_id", "score", "strand", "n_motifs", 
                                         "n_bp_cov", "seq_len", "frac_bp_cov"]

pool1_fimo_no_ets_chip_max_cov = pd.read_table(pool1_fimo_no_ets_chip_max_cov_f, sep="\t", header=None)
pool1_fimo_no_ets_chip_max_cov.columns = ["unique_id", "max_cov"]

pool1_fimo_no_ets_chip_cov = pool1_fimo_no_ets_chip_bp_cov.merge(pool1_fimo_no_ets_chip_max_cov, on="unique_id")
print(len(pool1_fimo_no_ets_chip_cov))


# In[32]:


all_cov_dfs = {"fimo": fimo_cov, "fimo_chip": fimo_chip_cov, "fimo_clust": fimo_clust_cov,  
               "pool1_fimo": pool1_fimo_cov, "pool1_fimo_chip": pool1_fimo_chip_cov, 
               "pool1_fimo_no_ets": pool1_fimo_no_ets_cov, 
               "pool1_fimo_no_ets_chip": pool1_fimo_no_ets_chip_cov,
               "fimo_mosbat": fimo_mosbat_cov}

all_motif_dfs = {"fimo": fimo, "fimo_chip": fimo_chip, "pool1_fimo": pool1_fimo, "pool1_fimo_chip": pool1_fimo_chip,
                 "pool1_fimo_no_ets": pool1_fimo_no_ets, "pool1_fimo_no_ets_chip": pool1_fimo_no_ets_chip}


# In[33]:


for key in all_cov_dfs.keys():
    df = all_cov_dfs[key]
    df["max_cov"] = df.apply(fix_small_decimal, axis=1)
    df["log_n_motifs"] = np.log(df["n_motifs"]+1)
    df["log_bp_cov"] = np.log(df["n_bp_cov"]+1)
    df["log_max_cov"] = np.log(df["max_cov"]+1)
    df["cage_id"] = df.apply(get_cage_id, axis=1)

fimo_chip_cov.head()


# In[34]:


for key in all_motif_dfs.keys():
    df = all_motif_dfs[key]
    df["cage_id"] = df.apply(get_cage_id, axis=1)
    
fimo.sample(5)


# In[35]:


tf_ts = pd.read_table(tf_ts_f, sep="\t")
tf_ts.head()


# In[36]:


cage_v_mpra = pd.read_table(cage_v_mpra_f, sep="\t")
cage_v_mpra["oligo_reg"] = cage_v_mpra["unique_id"].str.split("__", expand=True)[2]
cage_v_mpra.head()


# In[37]:


tss_cage_map = pd.read_table(tss_cage_map_f, sep="\t")
tss_cage_map.head()


# In[38]:


enh_cage_map = pd.read_table(enh_cage_map_f, sep="\t")
enh_cage_map.head()


# ## 2. for pool1: join coverage and motif files with MPRA expr/spec files

# In[39]:


# since enhancers have 2 TSS_ids, need to join these separately
tmp = cage_v_mpra.merge(tss_cage_map[["oligo_reg", "TSS_id"]], on="oligo_reg", how="left")
tmp_enh_pos = tmp[(tmp["unique_id"].str.contains("Enhancer")) & (tmp["TSS_id"].str[-1] == "+")]
tmp_enh_neg = tmp[(tmp["unique_id"].str.contains("Enhancer")) & (tmp["TSS_id"].str[-1] == "-")]

tmp_enh_pos = tmp_enh_pos.merge(enh_cage_map[["TSS_id_Pos", "enhancer_id"]], left_on="TSS_id", right_on="TSS_id_Pos",
                                how="left")
tmp_enh_neg = tmp_enh_neg.merge(enh_cage_map[["TSS_id_Neg", "enhancer_id"]], left_on="TSS_id", right_on="TSS_id_Neg",
                                how="left")
tmp_enh_pos = tmp_enh_pos.drop("TSS_id_Pos", axis=1)
tmp_enh_neg = tmp_enh_neg.drop("TSS_id_Neg", axis=1)
tmp_enh = tmp_enh_pos.append(tmp_enh_neg)
tmp_enh["TSS_id"] = tmp_enh["enhancer_id"]
tmp_enh.drop("enhancer_id", axis=1, inplace=True)
tmp_enh.sample(5)


# In[40]:


tmp_no_enh = tmp[~tmp["unique_id"].str.contains("Enhancer")]
cage_v_mpra = tmp_no_enh.append(tmp_enh)
cage_v_mpra.sample(5)


# In[42]:


cage_v_mpra.PromType2.value_counts()


# In[43]:


pool1_fimo_cov = pool1_fimo_cov.merge(cage_v_mpra, on="unique_id")
pool1_fimo = pool1_fimo.merge(cage_v_mpra, on="unique_id")
pool1_fimo_cov["cage_id"] = pool1_fimo_cov["TSS_id"]
pool1_fimo["cage_id"] = pool1_fimo["TSS_id"]
all_cov_dfs["pool1_fimo"] = pool1_fimo_cov
all_motif_dfs["pool1_fimo"] = pool1_fimo
print(len(pool1_fimo_cov))
pool1_fimo_cov.sample(5)


# In[44]:


pool1_fimo_chip_cov = pool1_fimo_chip_cov.merge(cage_v_mpra, on="unique_id")

# for old chip files:
#pool1_fimo_chip = pool1_fimo_chip.drop("unique_id", axis=1).merge(cage_v_mpra, left_on="cage_id", right_on="TSS_id")

# for new chip files:
pool1_fimo_chip = pool1_fimo_chip.merge(cage_v_mpra, on="unique_id")

pool1_fimo_chip_cov["cage_id"] = pool1_fimo_chip_cov["TSS_id"]
pool1_fimo_chip["cage_id"] = pool1_fimo_chip["TSS_id"]
all_cov_dfs["pool1_fimo_chip"] = pool1_fimo_chip_cov
all_motif_dfs["pool1_fimo_chip"] = pool1_fimo_chip
print(len(pool1_fimo_chip_cov))


# In[45]:


pool1_fimo_no_ets_cov = pool1_fimo_no_ets_cov.merge(cage_v_mpra, on="unique_id")
pool1_fimo_no_ets = pool1_fimo_no_ets.merge(cage_v_mpra, on="unique_id")
pool1_fimo_no_ets_cov["cage_id"] = pool1_fimo_no_ets_cov["TSS_id"]
pool1_fimo_no_ets["cage_id"] = pool1_fimo_no_ets["TSS_id"]
all_cov_dfs["pool1_fimo_no_ets"] = pool1_fimo_no_ets_cov
all_motif_dfs["pool1_fimo_no_ets"] = pool1_fimo_no_ets
print(len(pool1_fimo_no_ets_cov))


# In[46]:


pool1_fimo_no_ets_chip_cov = pool1_fimo_no_ets_chip_cov.merge(cage_v_mpra, on="unique_id")

# for old chip files:
# pool1_fimo_no_ets_chip = pool1_fimo_no_ets_chip.drop("unique_id", axis=1).merge(cage_v_mpra, left_on="cage_id", 
#                                                                                 right_on="TSS_id")

# for new chip files:
pool1_fimo_no_ets_chip = pool1_fimo_no_ets_chip.merge(cage_v_mpra, on="unique_id")

pool1_fimo_no_ets_chip_cov["cage_id"] = pool1_fimo_no_ets_chip_cov["TSS_id"]
pool1_fimo_no_ets_chip["cage_id"] = pool1_fimo_no_ets_chip["TSS_id"]
all_cov_dfs["pool1_fimo_no_ets_chip"] = pool1_fimo_no_ets_chip_cov
all_motif_dfs["pool1_fimo_no_ets_chip"] = pool1_fimo_no_ets_chip
print(len(pool1_fimo_no_ets_chip_cov))


# In[47]:


pool1_fimo_no_ets_chip.head()


# ## 3. find avg TF/motif specificity per tile

# In[48]:


all_spec_dfs = {}
for key in all_motif_dfs.keys():
    print(key)
    df = all_motif_dfs[key]
    df["motif"] = df["motif"].str.upper()
    df = df.merge(tf_ts, left_on="motif", right_on="tf", how="left")
    df_nonan = df[~pd.isnull(df["tissue_sp_3"])]
    df_deduped = df_nonan.drop_duplicates(subset=["motif", "unique_id"])
    avg_sp = df_deduped.groupby(["cage_id"])["tissue_sp_3"].agg("mean").reset_index()
    avg_sp.columns = ["cage_id", "avg_tf_tissue_sp"]
    avg_sp["log_avg_tf_tissue_sp"] = np.log(avg_sp["avg_tf_tissue_sp"]+1)
    all_spec_dfs[key] = avg_sp
avg_sp.sample(5)


# ## 4. merge and write coverage files

# In[49]:


file_prefixes = {"fimo": "all_fimo_map", "fimo_chip": "all_fimo_map.chip_intersected", 
                 "fimo_clust": "all_fimo_map.bulyk_clusters", 
                 "pool1_fimo": "pool1_fimo_map", "pool1_fimo_chip": "pool1_fimo_map.chip_intersected", 
                 "pool1_fimo_no_ets": "pool1_fimo_map.no_ETS_motifs", 
                 "pool1_fimo_no_ets_chip": "pool1_fimo_map.chip_intersected.no_ETS_motifs",
                 "fimo_mosbat": "all_fimo_map.mosbat_clusters"}

for key in all_cov_dfs.keys():
    print(key)
    name = file_prefixes[key]
    cov_df = all_cov_dfs[key]
    if key in all_spec_dfs.keys():
        spec_df = all_spec_dfs[key]
        merge_df = cov_df.merge(spec_df, on="cage_id", how="left")
        merge_df = merge_df[["unique_id", "cage_id", "n_motifs", "n_bp_cov", "max_cov", "avg_tf_tissue_sp", 
                             "log_n_motifs", "log_bp_cov", "log_max_cov", "log_avg_tf_tissue_sp"]].drop_duplicates()
    else:
        merge_df = cov_df[["unique_id", "cage_id", "n_motifs", "n_bp_cov", "max_cov", 
                                 "log_n_motifs", "log_bp_cov", "log_max_cov"]].drop_duplicates()
    
    merge_df.to_csv("../../data/04__coverage/%s.all_cov.new.txt" % name, sep="\t", index=False)
    
    if "pool1" in name:
        merge_df = merge_df.merge(cage_v_mpra[["unique_id", "mpra_activ", "mpra_ts"]], on="unique_id")
    all_cov_dfs[key] = merge_df


# ## 5. plot correlations w/ MPRA data

# ## tissue specificity

# #### fimo only

# In[50]:


df = all_cov_dfs["pool1_fimo"]
df.head()


# In[51]:


#cmap = sns.light_palette("#8da0cb", as_cmap=True)
cmap = sns.light_palette("darkslategray", as_cmap=True)


# In[52]:


no_nan = df[~pd.isnull(df["log_avg_tf_tissue_sp"]) &
            ~pd.isnull(df["mpra_activ"])]
len(no_nan)


# In[53]:


fig = plt.figure(figsize=(1.2, 1.2))
ax = sns.kdeplot(no_nan["log_avg_tf_tissue_sp"], no_nan["mpra_activ"], cmap=cmap, 
                 shade=True, shade_lowest=False)
ax.set_ylabel("mean MPRA activity")
ax.set_xlabel("log(mean TF tissue specificity)")

r, p = stats.spearmanr(no_nan["log_avg_tf_tissue_sp"], no_nan["mpra_activ"])
print("r: %s, spearman p: %s" % (r, p))
ax.annotate("r = {:.2f}".format(r), xy=(.05, .9), xycoords=ax.transAxes, fontsize=fontsize)

# add n-value
ax.annotate("n = %s" % len(no_nan), ha="right", xy=(.96, .9), xycoords=ax.transAxes, 
            fontsize=fontsize)


# In[54]:


no_nan = df[~pd.isnull(df["log_avg_tf_tissue_sp"]) &
            ~pd.isnull(df["mpra_ts"])]
len(no_nan)


# In[55]:


fig = plt.figure(figsize=(1.2, 1.2))
ax = sns.kdeplot(no_nan["log_avg_tf_tissue_sp"], no_nan["mpra_ts"], cmap=cmap, 
                 shade=True, shade_lowest=False)
ax.set_ylabel("tissue specificity in MPRA")
ax.set_xlabel("log(mean TF tissue specificity)")

r, p = stats.spearmanr(no_nan["log_avg_tf_tissue_sp"], no_nan["mpra_ts"])
print("r: %s, spearman p: %s" % (r, p))
ax.annotate("r = {:.2f}".format(r), xy=(.05, .9), xycoords=ax.transAxes, fontsize=fontsize)

# add n-value
ax.annotate("n = %s" % len(no_nan), ha="right", xy=(.96, .9), xycoords=ax.transAxes, 
            fontsize=fontsize)


# #### fimo intersected w/ chip

# In[56]:


df = all_cov_dfs["pool1_fimo_chip"]


# In[57]:


no_nan = df[~pd.isnull(df["log_avg_tf_tissue_sp"]) &
            ~pd.isnull(df["mpra_activ"])]
len(no_nan)


# In[58]:


fig = plt.figure(figsize=(1.2, 1.2))
ax = sns.kdeplot(no_nan["log_avg_tf_tissue_sp"], no_nan["mpra_activ"], cmap=cmap, 
                 shade=True, shade_lowest=False)
ax.set_ylabel("mean MPRA activity")
ax.set_xlabel("log(mean TF tissue specificity)")

r, p = stats.spearmanr(no_nan["log_avg_tf_tissue_sp"], no_nan["mpra_activ"])
print("r: %s, spearman p: %s" % (r, p))
ax.annotate("r = {:.2f}".format(r), xy=(.05, .9), xycoords=ax.transAxes, fontsize=fontsize)

# add n-value
ax.annotate("n = %s" % len(no_nan), ha="right", xy=(.96, .9), xycoords=ax.transAxes, 
            fontsize=fontsize)
fig.savefig("Fig_2C_3.pdf", bbox_inches="tight", dpi="figure")


# In[59]:


no_nan = df[~pd.isnull(df["log_avg_tf_tissue_sp"]) &
            ~pd.isnull(df["mpra_ts"])]
len(no_nan)


# In[60]:


fig = plt.figure(figsize=(1.2, 1.2))
ax = sns.kdeplot(no_nan["log_avg_tf_tissue_sp"], no_nan["mpra_ts"], cmap=cmap, 
                 shade=True, shade_lowest=False)
ax.set_ylabel("tissue specificity in MPRA")
ax.set_xlabel("log(mean TF tissue specificity)")

r, p = stats.spearmanr(no_nan["log_avg_tf_tissue_sp"], no_nan["mpra_ts"])
print("r: %s, spearman p: %s" % (r, p))
ax.annotate("r = {:.2f}".format(r), xy=(.05, .9), xycoords=ax.transAxes, fontsize=fontsize)

# add n-value
ax.annotate("n = %s" % len(no_nan), ha="right", xy=(.96, .9), xycoords=ax.transAxes, 
            fontsize=fontsize)
fig.savefig("Fig_2C_6.pdf", bbox_inches="tight", dpi="figure")


# ## number of bp covered

# #### fimo only

# In[61]:


df = all_cov_dfs["pool1_fimo"]


# In[62]:


no_nan = df[~pd.isnull(df["log_bp_cov"]) &
            ~pd.isnull(df["mpra_activ"])]

# for these, only look at those with >1 motif
no_nan = no_nan[no_nan["n_motifs"] > 0]
len(no_nan)


# In[63]:


fig = plt.figure(figsize=(1.2, 1.2))
ax = sns.kdeplot(no_nan["log_bp_cov"], no_nan["mpra_activ"], cmap="Blues", 
                 shade=True, shade_lowest=False)
ax.set_ylabel("mean MPRA activity")
ax.set_xlabel("log(number of bp covered by motif)")
ax.set_xlim((3,5))

r, p = stats.spearmanr(no_nan["log_bp_cov"], no_nan["mpra_activ"])
print("r: %s, spearman p: %s" % (r, p))
ax.annotate("r = {:.2f}".format(r), xy=(.05, .9), xycoords=ax.transAxes, fontsize=fontsize)

# add n-value
ax.annotate("n = %s" % len(no_nan), ha="right", xy=(.96, .9), xycoords=ax.transAxes, 
            fontsize=fontsize)


# In[64]:


no_nan = df[~pd.isnull(df["log_bp_cov"]) &
            ~pd.isnull(df["mpra_ts"])]

# for these, only look at those with >1 motif
no_nan = no_nan[no_nan["n_motifs"] > 0]
len(no_nan)


# In[65]:


fig = plt.figure(figsize=(1.2, 1.2))
ax = sns.kdeplot(no_nan["log_bp_cov"], no_nan["mpra_ts"], cmap="Blues", 
                 shade=True, shade_lowest=False)
ax.set_ylabel("tissue specificity in MPRA")
ax.set_xlabel("log(number of bp covered by motif)")
ax.set_xlim((3, 5))

r, p = stats.spearmanr(no_nan["log_bp_cov"], no_nan["mpra_ts"])
print("r: %s, spearman p: %s" % (r, p))
ax.annotate("r = {:.2f}".format(r), xy=(.05, .9), xycoords=ax.transAxes, fontsize=fontsize)

# add n-value
ax.annotate("n = %s" % len(no_nan), ha="right", xy=(.96, .9), xycoords=ax.transAxes, 
            fontsize=fontsize)


# #### fimo intersected w/ chip

# In[66]:


df = all_cov_dfs["pool1_fimo_chip"]


# In[67]:


no_nan = df[~pd.isnull(df["log_bp_cov"]) &
            ~pd.isnull(df["mpra_activ"])]

# for these, only look at those with >1 motif
no_nan = no_nan[no_nan["n_motifs"] > 0]
len(no_nan)


# In[68]:


fig = plt.figure(figsize=(1.2, 1.2))
no_0 = no_nan[no_nan["n_bp_cov"] > 0]
all_0 = no_nan[no_nan["n_bp_cov"] == 0]
ax = sns.kdeplot(no_0["log_bp_cov"], no_0["mpra_activ"], cmap="Blues", 
                 shade=True, shade_lowest=False)
ax.set_ylabel("mean MPRA activity")
ax.set_xlabel("log(number of bp covered by motif)")

r, p = stats.spearmanr(no_nan["log_bp_cov"], no_nan["mpra_activ"])
print("r: %s, spearman p: %s" % (r, p))
ax.annotate("r = {:.2f}".format(r), xy=(.05, .9), xycoords=ax.transAxes, fontsize=fontsize)

# add n-value
ax.annotate("n = %s" % len(no_nan), ha="right", xy=(.96, .9), xycoords=ax.transAxes, 
            fontsize=fontsize)

fig.savefig("Fig_2C_1.pdf", bbox_inches="tight", dpi="figure")


# In[69]:


no_nan = df[~pd.isnull(df["log_bp_cov"]) &
            ~pd.isnull(df["mpra_ts"])]

# for these, only look at those with >1 motif
no_nan = no_nan[no_nan["n_motifs"] > 0]
len(no_nan)


# In[70]:


fig = plt.figure(figsize=(1.2, 1.2))
ax = sns.kdeplot(no_nan["log_bp_cov"], no_nan["mpra_ts"], cmap="Blues", 
                 shade=True, shade_lowest=False)
ax.set_ylabel("tissue specificity in MPRA")
ax.set_xlabel("log(number of bp covered by motif)")

r, p = stats.spearmanr(no_nan["log_bp_cov"], no_nan["mpra_ts"])
print("r: %s, spearman p: %s" % (r, p))
ax.annotate("r = {:.2f}".format(r), xy=(.05, .9), xycoords=ax.transAxes, fontsize=fontsize)

# add n-value
ax.annotate("n = %s" % len(no_nan), ha="right", xy=(.96, .9), xycoords=ax.transAxes, 
            fontsize=fontsize)

fig.savefig("Fig_2C_4.pdf", bbox_inches="tight", dpi="figure")


# ## max overlapping coverage

# #### fimo only

# In[71]:


df = all_cov_dfs["pool1_fimo"]


# In[72]:


cmap = sns.light_palette("firebrick", as_cmap=True)


# In[73]:


no_nan = df[~pd.isnull(df["log_max_cov"]) &
            ~pd.isnull(df["mpra_activ"])]

# for these, only look at those with >1 motif
no_nan = no_nan[no_nan["n_motifs"] > 0]
len(no_nan)


# In[74]:


fig = plt.figure(figsize=(1.2, 1.2))
ax = sns.kdeplot(no_nan["log_max_cov"], no_nan["mpra_activ"], cmap=cmap, 
                 shade=True, shade_lowest=False)
ax.set_ylabel("mean MPRA activity")
ax.set_xlabel("log(max overlapping motifs)")

r, p = stats.spearmanr(no_nan["log_max_cov"], no_nan["mpra_activ"])
print("r: %s, spearman p: %s" % (r, p))
ax.annotate("r = {:.2f}".format(r), xy=(.05, .9), xycoords=ax.transAxes, fontsize=fontsize)

# add n-value
ax.annotate("n = %s" % len(no_nan), ha="right", xy=(.96, .9), xycoords=ax.transAxes, 
            fontsize=fontsize)


# In[75]:


no_nan = df[~pd.isnull(df["log_max_cov"]) &
            ~pd.isnull(df["mpra_ts"])]

# for these, only look at those with >1 motif
no_nan = no_nan[no_nan["n_motifs"] > 0]
len(no_nan)


# In[76]:


fig = plt.figure(figsize=(1.2, 1.2))
ax = sns.kdeplot(no_nan["log_max_cov"], no_nan["mpra_ts"], cmap=cmap, 
                 shade=True, shade_lowest=False)
ax.set_ylabel("tissue specificity in MPRA")
ax.set_xlabel("log(max overlapping motifs)")

r, p = stats.spearmanr(no_nan["log_max_cov"], no_nan["mpra_ts"])
print("r: %s, spearman p: %s" % (r, p))
ax.annotate("r = {:.2f}".format(r), xy=(.05, .9), xycoords=ax.transAxes, fontsize=fontsize)

# add n-value
ax.annotate("n = %s" % len(no_nan), ha="right", xy=(.96, .9), xycoords=ax.transAxes, 
            fontsize=fontsize)


# #### fimo intersected w/ chip

# In[77]:


df = all_cov_dfs["pool1_fimo_chip"]


# In[78]:


no_nan = df[~pd.isnull(df["log_max_cov"]) &
            ~pd.isnull(df["mpra_activ"])]

# for these, only look at those with >1 motif
no_nan = no_nan[no_nan["n_motifs"] > 0]
len(no_nan)


# In[79]:


fig = plt.figure(figsize=(1.2, 1.2))
ax = sns.kdeplot(no_nan["log_max_cov"], no_nan["mpra_activ"], cmap=cmap, 
                 shade=True, shade_lowest=False)
ax.set_ylabel("mean MPRA activity")
ax.set_xlabel("log(max overlapping motifs)")

r, p = stats.spearmanr(no_nan["log_max_cov"], no_nan["mpra_activ"])
print("r: %s, spearman p: %s" % (r, p))
ax.annotate("r = {:.2f}".format(r), xy=(.05, .9), xycoords=ax.transAxes, fontsize=fontsize)

# add n-value
ax.annotate("n = %s" % len(no_nan), ha="right", xy=(.96, .9), xycoords=ax.transAxes, 
            fontsize=fontsize)

fig.savefig("Fig_2C_2.pdf", bbox_inches="tight", dpi="figure")


# In[80]:


no_nan = df[~pd.isnull(df["log_max_cov"]) &
            ~pd.isnull(df["mpra_ts"])]

# for these, only look at those with >1 motif
no_nan = no_nan[no_nan["n_motifs"] > 0]
len(no_nan)


# In[81]:


fig = plt.figure(figsize=(1.2, 1.2))
ax = sns.kdeplot(no_nan["log_max_cov"], no_nan["mpra_ts"], cmap=cmap, 
                 shade=True, shade_lowest=False)
ax.set_ylabel("tissue specificity in MPRA")
ax.set_xlabel("log(max overlapping motifs)")

r, p = stats.spearmanr(no_nan["log_max_cov"], no_nan["mpra_ts"])
print("r: %s, spearman p: %s" % (r, p))
ax.annotate("r = {:.2f}".format(r), xy=(.05, .9), xycoords=ax.transAxes, fontsize=fontsize)

# add n-value
ax.annotate("n = %s" % len(no_nan), ha="right", xy=(.96, .9), xycoords=ax.transAxes, 
            fontsize=fontsize)

fig.savefig("Fig_2C_5.pdf", bbox_inches="tight", dpi="figure")
```
