## Supplementary material for "High-throughput functional analysis of IncRNA core promoters elucidatesrules governing tissue-specificity": 00__parse_pwm_files.py

```
# coding: utf-8

# # notebook to parse PWM files and write them to MEME format

# In[1]:


from os import walk


# In[2]:


file_dir = "../../misc/02__bulyk_clusters/00__original_files/glossary-pwm"


# In[3]:


out_dir = "../../misc/02__bulyk_clusters/01__meme_file"
get_ipython().system('mkdir -p $out_dir')


# In[4]:


files = []
for (dirpath, dirnames, filenames) in walk(file_dir):
    files.extend(filenames)
    break


# In[5]:


print(len(files))
files[0:5]


# In[6]:


file_contents = {}
for filename in files:
    path = "%s/%s" % (file_dir, filename)
    tf = filename.split(".")[0]
    with open(path) as f:
        content = f.readlines()
    content = [x.strip() for x in content]
    file_contents[tf] = content
len(file_contents)


# In[8]:


out_file = "glossary-pwm.txt"
with open("%s/%s" % (out_dir, out_file), "w") as f:
    f.write("MEME version 4\n\n")
    f.write("ALPHABET= ACGT\n\n")
    f.write("strands: + -\n\n")
    f.write("Background letter frequences (from uniform background):\n")
    f.write("A 0.25000 C 0.25000 G 0.25000 T 0.25000\n\n")
    for tf in file_contents:
        f.write("MOTIF %s\n\n" % tf)
        f.write("letter-probability matrix: alength= 4 w= %s nsites= 20 E= 0\n" % (len(file_contents[tf])-1))
        for i, line in enumerate(file_contents[tf]):
            if i == 0:
                continue
            else:
                split_line = line.split("\t")
                new_line = " %s\t%s\t%s\t%s\n" % (split_line[1], split_line[2], split_line[3], split_line[4])
                f.write(new_line)
        f.write("\n")
f.close()


# In[ ]:
```
