## Supplementary material for "High-throughput functional analysis of IncRNA core promoters elucidatesrules governing tissue-specificity": 07__motif_coverage.py

```
# coding: utf-8

# # 07__motif_coverage
# # analyzing how motif coverage correlates with MPRA properties and biotypes; clustering similar motifs
# 
# in this notebook, i look at how the coverage metrics (# bp covered and max coverage of motifs; done separately, see methods) look within biotypes. i look at either the motifs alone, the motifs *after* limiting to only those motifs which have been validated by a corresponding chip peak, and non-redundant 8mer motifs from Mariani et al. i also cluster the motifs using MoSBAT (done separately using their webtool) and re-calculate the metrics. finally, i examine the conservation of overlapping motifs using phylop 46-way placental mammal scores.
# 
# ------
# 
# figures in this notebook:
# - **Fig 2D and 2F**: cumulative density plots of # bp covered and max motif coverage across biotypes
# - **Fig 2E and 2G**: cumulative density plots of # bp covered and max motif coverage within lincRNAs and mRNAs, separated by specificity profiles
# - **Fig S9**: heatmap of MoSBAT clustered motifs, and more cumulative density plots (after using the two clustering methods)
# - **Fig 2H**: conservation of overlapping motifs via phylop46way scores

# In[1]:


import warnings
warnings.filterwarnings('ignore')

import pandas as pd
import numpy as np
import matplotlib as mpl
import matplotlib.pyplot as plt
import seaborn as sns
import sys
import time

from itertools import chain
from decimal import Decimal
from random import shuffle
from scipy import stats
from scipy import signal
from scipy.spatial import distance
from scipy.cluster import hierarchy
from statsmodels.sandbox.stats import multicomp

# import utils
sys.path.append("../../utils")
from plotting_utils import *
from misc_utils import *
from norm_utils import *

get_ipython().magic('matplotlib inline')


# In[2]:


sns.set(**PAPER_PRESET)
fontsize = PAPER_FONTSIZE


# ## variables

# In[3]:


mosbat_file = "../../misc/02__mosbat/results.from_mosbat.txt"


# In[30]:


all_tss_f = "../../data/00__index/0__all_tss/All.TSS.114bp.uniq.new.bed"
cage_expr_f = "../../misc/01__cage/All_TSS_and_enh.CAGE_grouped_exp.tissue_sp.txt"


# In[31]:


fimo_cov_f = "../../data/04__coverage/all_fimo_map.all_cov.new.txt"
fimo_chip_cov_f = "../../data/04__coverage/all_fimo_map.chip_intersected.all_cov.new.txt"
fimo_clust_cov_f = "../../data/04__coverage/all_fimo_map.bulyk_clusters.all_cov.new.txt"
fimo_mosbat_cov_f = "../../data/04__coverage/all_fimo_map.mosbat_clusters.all_cov.new.txt"

pool1_fimo_cov_f = "../../data/04__coverage/pool1_fimo_map.all_cov.new.txt"
pool1_fimo_chip_cov_f = "../../data/04__coverage/all_fimo_map.chip_intersectedall_cov.new.txt"


# In[6]:


fimo_phylop_f = "../../misc/03__fimo/03__phylop_meta_plot/all_fimo_map.phylop46way.txt"
fimo_chip_phylop_f = "../../misc/03__fimo/03__phylop_meta_plot/all_fimo_map.chip_intersected.phylop46way.txt"
fimo_clust_phylop_f = "../../misc/03__fimo/03__phylop_meta_plot/all_fimo_map.bulyk_clusters.phylop46way.txt"


# In[7]:


all_phylop_f = "../../data/00__index/0__all_tss/All.TSS.114bp.uniq.phylop46way.txt"


# In[8]:


dnase_f = "../../misc/05__dnase/All.TSS.114bp.uniq.count_DNase_accessible_samples.txt"


# note: the reason why some IDs are not in the expression file is these are FANTOM CAT IDs that for some reason are not present in the FANTOM5 robust set. so, we exclude these.

# ## 1. import data

# In[9]:


corr = pd.read_table(mosbat_file, sep="\t")


# In[10]:


all_phylop = pd.read_table(all_phylop_f, sep="\t")
all_phylop.columns = ["chr", "start", "end", "unique_id", "score", "strand", "size", "num_data", "min", "max",
                      "mean", "median"]
all_phylop.head()


# In[11]:


fimo_phylop = pd.read_table(fimo_phylop_f, sep="\t", header=None)
cols = ["chr", "start", "end", "motif", "n_ov", "tss_dist"]
cols.extend(np.arange(-150, 150, step=1))
fimo_phylop.columns = cols


# In[12]:


fimo_chip_phylop = pd.read_table(fimo_chip_phylop_f, sep="\t", header=None)
fimo_chip_phylop.columns = cols


# In[13]:


fimo_clust_phylop = pd.read_table(fimo_clust_phylop_f, sep="\t", header=None)
fimo_clust_phylop.columns = cols


# In[14]:


dnase = pd.read_table(dnase_f, sep="\t", header=None)
dnase.columns = ["unique_id", "n_accessible"]
dnase["PromType2"] = dnase.unique_id.str.split("__", expand=True)[0]


# In[15]:


fimo_cov = pd.read_table(fimo_cov_f, sep="\t")
fimo_chip_cov = pd.read_table(fimo_chip_cov_f, sep="\t")
fimo_clust_cov = pd.read_table(fimo_clust_cov_f, sep="\t")
fimo_mosbat_cov = pd.read_table(fimo_mosbat_cov_f, sep="\t")
fimo_chip_cov.sample(5)


# In[16]:


print(len(fimo_cov))
print(len(fimo_chip_cov))
print(len(fimo_clust_cov))
print(len(fimo_mosbat_cov))


# In[18]:


fimo_cov["PromType2"] = fimo_cov["unique_id"].str.split("__", expand=True)[0]
fimo_chip_cov["PromType2"] = fimo_chip_cov["unique_id"].str.split("__", expand=True)[0]
fimo_clust_cov["PromType2"] = fimo_clust_cov["unique_id"].str.split("__", expand=True)[0]
fimo_mosbat_cov["PromType2"] = fimo_mosbat_cov["unique_id"].str.split("__", expand=True)[0]
fimo_cov.sample(5)


# In[19]:


# filter to those that have at least 1 motif so distributions are not 0-skewed
fimo_cov = fimo_cov[fimo_cov["n_motifs"] > 0]
print(len(fimo_cov))

fimo_chip_cov = fimo_chip_cov[fimo_chip_cov["n_motifs"] > 0]
print(len(fimo_chip_cov))

fimo_clust_cov = fimo_clust_cov[fimo_clust_cov["n_motifs"] > 0]
print(len(fimo_clust_cov))

fimo_mosbat_cov = fimo_mosbat_cov[fimo_mosbat_cov["n_motifs"] > 0]
print(len(fimo_mosbat_cov))


# In[20]:


cage_expr = pd.read_table(cage_expr_f, sep="\t")
cage_expr.head()


# In[21]:


# inner merging with this file ensures that we are only looking at robust TSSs and robust enhancers
fimo_cov = fimo_cov.merge(cage_expr, on="cage_id", how="left")
fimo_cov.head()


# In[22]:


fimo_chip_cov = fimo_chip_cov.merge(cage_expr, on="cage_id", how="left")
fimo_chip_cov.sample(5)


# In[23]:


fimo_clust_cov = fimo_clust_cov.merge(cage_expr, on="cage_id", how="left")
fimo_clust_cov.head()


# In[24]:


fimo_mosbat_cov = fimo_mosbat_cov.merge(cage_expr, on="cage_id", how="left")
fimo_mosbat_cov.head()


# In[25]:


chip_cov_exp = fimo_chip_cov[~pd.isnull(fimo_chip_cov["av_exp"])]
motif_cov_exp = fimo_cov[~pd.isnull(fimo_cov["av_exp"])]
cluster_cov_exp = fimo_clust_cov[~pd.isnull(fimo_clust_cov["av_exp"])]
mosbat_cov_exp = fimo_mosbat_cov[~pd.isnull(fimo_mosbat_cov["av_exp"])]


# In[26]:


motif_cov_exp.PromType2.value_counts()


# In[27]:


chip_cov_exp.PromType2.value_counts()


# In[28]:


cluster_cov_exp.PromType2.value_counts()


# In[32]:


all_tss = pd.read_table(all_tss_f, sep="\t", header=None)
all_tss.columns = ["chr", "start", "end", "name", "score", "strand"]
all_tss["PromType2"] = all_tss["name"].str.split("__", expand=True)[0]
all_tss.PromType2.value_counts()


# ## 2. plot # bp covered & max cov across biotypes

# ### all motifs

# In[33]:


enh_vals = fimo_cov[fimo_cov["PromType2"] == "Enhancer"]["log_bp_cov"]
linc_vals = fimo_cov[fimo_cov["PromType2"] == "intergenic"]["log_bp_cov"]
dlnc_vals = fimo_cov[fimo_cov["PromType2"] == "div_lnc"]["log_bp_cov"]
pc_vals = fimo_cov[fimo_cov["PromType2"] == "protein_coding"]["log_bp_cov"]
dpc_vals = fimo_cov[fimo_cov["PromType2"] == "div_pc"]["log_bp_cov"]

fig = plt.figure(figsize=(2.25, 1.6))
ax = sns.kdeplot(data=enh_vals, cumulative=True, bw=0.1, color=TSS_CLASS_PALETTE["Enhancer"], 
                 label="eRNAs (n=%s)" % len(enh_vals))
sns.kdeplot(data=linc_vals, cumulative=True, bw=0.1, color=TSS_CLASS_PALETTE["intergenic"], 
            label="lincRNAs (n=%s)" % len(linc_vals), ax=ax)
sns.kdeplot(data=dlnc_vals, cumulative=True, bw=0.1, color=TSS_CLASS_PALETTE["div_lnc"], 
            label="div. lncRNAs (n=%s)" % len(dlnc_vals), ax=ax)
sns.kdeplot(data=pc_vals, cumulative=True, bw=0.1, color=TSS_CLASS_PALETTE["protein_coding"], 
            label="mRNAs (n=%s)" % len(pc_vals), ax=ax)
sns.kdeplot(data=dpc_vals, cumulative=True, bw=0.1, color=TSS_CLASS_PALETTE["div_pc"], 
            label="div. mRNAs (n=%s)" % len(dpc_vals), ax=ax)
ax.set_xlabel("log(# of bp covered)")
ax.set_ylabel("cumulative density")
ax.set_xlim((2, 5))
ax.set_ylim((0, 1.05))
plt.legend(handlelength=1)


# In[34]:


# for each group, split into tissue-sp v dynamic v ubiquitous
idxs = list(range(0, 5))
promtypes = ["Enhancer", "intergenic", "protein_coding"]
names = ["eRNAs", "lincRNAs", "mRNAs"]
df = motif_cov_exp
col = "log_bp_cov"
xlabel = "log(# of bp covered)"
xlim = (2, 5)

f, axarr = plt.subplots(nrows=1, ncols=3, sharex=False, sharey=True, figsize=(5.5, 1.6))

for i, promtype, name in zip(idxs, promtypes, names):
    print("i: %s, promtype: %s, name: %s" % (i, promtype, name))
    ax = axarr[i]
    
    promtype_vals = df[df["PromType2"] == promtype]
    ts_vals = promtype_vals[promtype_vals["tss_type"] == "tissue-specific"][col]
    ub_vals = promtype_vals[promtype_vals["tss_type"] == "ubiquitous"][col]
    dy_vals = promtype_vals[promtype_vals["tss_type"] == "dynamic"][col]
    
    ub_ts_u, ub_ts_pval = stats.mannwhitneyu(ub_vals, ts_vals, alternative="two-sided", use_continuity=False)
    print("ub/ts pval: %s" % ub_ts_pval)
    
    ub_dy_u, ub_dy_pval = stats.mannwhitneyu(ub_vals, dy_vals, alternative="two-sided", use_continuity=False)
    print("ub/dy pval: %s" % ub_dy_pval)
    
    ts_dy_u, ts_dy_pval = stats.mannwhitneyu(ts_vals, dy_vals, alternative="two-sided", use_continuity=False)
    print("ts/dy pval: %s" % ts_dy_pval)
    
    sns.kdeplot(data=ts_vals, cumulative=True, bw=0.1, color=TSS_CLASS_PALETTE[promtype], linestyle="dashed",
                label="tissue-specific\n(n=%s)" % len(ts_vals), ax=ax)
    sns.kdeplot(data=ub_vals, cumulative=True, bw=0.1, color=TSS_CLASS_PALETTE[promtype], linestyle="solid",
                label="ubiquitous\n(n=%s)" % len(ub_vals), ax=ax)
    sns.kdeplot(data=dy_vals, cumulative=True, bw=0.1, color=TSS_CLASS_PALETTE[promtype], linestyle="dotted",
                label="dynamic\n(n=%s)" % len(dy_vals), ax=ax)
    
    ax.set_title("%s (%s total)" % (name, len(promtype_vals)))
    ax.set_xlabel(xlabel)
    ax.legend(loc="upper left", handlelength=2)
    ax.set_xlim(xlim)
    
    if i == 0:
        ax.set_ylabel("cumulative density")

plt.subplots_adjust(wspace=0.1)
plt.ylim((0, 1.05))


# In[35]:


enh_vals = fimo_cov[fimo_cov["PromType2"] == "Enhancer"]["log_max_cov"]
linc_vals = fimo_cov[fimo_cov["PromType2"] == "intergenic"]["log_max_cov"]
dlnc_vals = fimo_cov[fimo_cov["PromType2"] == "div_lnc"]["log_max_cov"]
pc_vals = fimo_cov[fimo_cov["PromType2"] == "protein_coding"]["log_max_cov"]
dpc_vals = fimo_cov[fimo_cov["PromType2"] == "div_pc"]["log_max_cov"]

fig = plt.figure(figsize=(2.35, 1.6))
ax = sns.kdeplot(data=enh_vals, cumulative=True, bw=0.1, color=TSS_CLASS_PALETTE["Enhancer"], 
                 label="eRNAs\n(n=%s)" % len(enh_vals))
sns.kdeplot(data=linc_vals, cumulative=True, bw=0.1, color=TSS_CLASS_PALETTE["intergenic"], 
            label="lincRNAs\n(n=%s)" % len(linc_vals), ax=ax)
sns.kdeplot(data=dlnc_vals, cumulative=True, bw=0.1, color=TSS_CLASS_PALETTE["div_lnc"], 
            label="div. lncRNAs\n(n=%s)" % len(dlnc_vals), ax=ax)
sns.kdeplot(data=pc_vals, cumulative=True, bw=0.1, color=TSS_CLASS_PALETTE["protein_coding"], 
            label="mRNAs\n(n=%s)" % len(pc_vals), ax=ax)
sns.kdeplot(data=dpc_vals, cumulative=True, bw=0.1, color=TSS_CLASS_PALETTE["div_pc"], 
            label="div. mRNAs\n(n=%s)" % len(dpc_vals), ax=ax)
ax.set_xlabel("log(max coverage)")
ax.set_ylabel("cumulative density")
ax.set_ylim((0, 1.05))
ax.legend(loc="bottom right", handlelength=1)
#fig.savefig("max_cov.all_biotypes.for_poster.pdf", dpi="figure", bbox_inches="tight")


# In[36]:


# for each group, split into tissue-sp v dynamic v ubiquitous
idxs = list(range(0, 5))
promtypes = ["Enhancer", "intergenic", "protein_coding"]
names = ["eRNAs", "lincRNAs", "mRNAs"]
df = motif_cov_exp
col = "log_max_cov"
xlabel = "log(max coverage)"
xlim = (0, 5.5)

f, axarr = plt.subplots(nrows=1, ncols=3, sharex=False, sharey=True, figsize=(5.5, 1.6))

for i, promtype, name in zip(idxs, promtypes, names):
    print("i: %s, promtype: %s, name: %s" % (i, promtype, name))
    ax = axarr[i]
    
    promtype_vals = df[df["PromType2"] == promtype]
    ts_vals = promtype_vals[promtype_vals["tss_type"] == "tissue-specific"][col]
    ub_vals = promtype_vals[promtype_vals["tss_type"] == "ubiquitous"][col]
    dy_vals = promtype_vals[promtype_vals["tss_type"] == "dynamic"][col]
    
    ub_ts_u, ub_ts_pval = stats.mannwhitneyu(ub_vals, ts_vals, alternative="two-sided", use_continuity=False)
    print("ub/ts pval: %s" % ub_ts_pval)
    
    ub_dy_u, ub_dy_pval = stats.mannwhitneyu(ub_vals, dy_vals, alternative="two-sided", use_continuity=False)
    print("ub/dy pval: %s" % ub_dy_pval)
    
    ts_dy_u, ts_dy_pval = stats.mannwhitneyu(ts_vals, dy_vals, alternative="two-sided", use_continuity=False)
    print("ts/dy pval: %s" % ts_dy_pval)
    
    sns.kdeplot(data=ts_vals, cumulative=True, bw=0.1, color=TSS_CLASS_PALETTE[promtype], linestyle="dashed",
                label="tissue-sp.\n(n=%s)" % len(ts_vals), ax=ax)
    sns.kdeplot(data=ub_vals, cumulative=True, bw=0.1, color=TSS_CLASS_PALETTE[promtype], linestyle="solid",
                label="ubiq.\n(n=%s)" % len(ub_vals), ax=ax)
    sns.kdeplot(data=dy_vals, cumulative=True, bw=0.1, color=TSS_CLASS_PALETTE[promtype], linestyle="dotted",
                label="dynamic\n(n=%s)" % len(dy_vals), ax=ax)
    
    ax.set_title("%s (%s total)" % (name, len(promtype_vals)))
    ax.set_xlabel(xlabel)
    ax.legend(loc="bottom right", handlelength=2)
    ax.set_xlim(xlim)
    ax.set_ylim((0, 1.05))
    
    if i == 0:
        ax.set_ylabel("cumulative density")


# ### ChIP-validated motifs

# In[37]:


enh_vals = fimo_chip_cov[fimo_chip_cov["PromType2"] == "Enhancer"]["log_bp_cov"]
linc_vals = fimo_chip_cov[fimo_chip_cov["PromType2"] == "intergenic"]["log_bp_cov"]
dlnc_vals = fimo_chip_cov[fimo_chip_cov["PromType2"] == "div_lnc"]["log_bp_cov"]
pc_vals = fimo_chip_cov[fimo_chip_cov["PromType2"] == "protein_coding"]["log_bp_cov"]
dpc_vals = fimo_chip_cov[fimo_chip_cov["PromType2"] == "div_pc"]["log_bp_cov"]

fig = plt.figure(figsize=(2.25, 1.6))
ax = sns.kdeplot(data=enh_vals, cumulative=True, bw=0.1, color=TSS_CLASS_PALETTE["Enhancer"], 
                 label="eRNAs\n(n=%s)" % len(enh_vals))
sns.kdeplot(data=linc_vals, cumulative=True, bw=0.1, color=TSS_CLASS_PALETTE["intergenic"], 
            label="lincRNAs\n(n=%s)" % len(linc_vals), ax=ax)
sns.kdeplot(data=dlnc_vals, cumulative=True, bw=0.1, color=TSS_CLASS_PALETTE["div_lnc"], 
            label="div. lncRNAs\n(n=%s)" % len(dlnc_vals), ax=ax)
sns.kdeplot(data=pc_vals, cumulative=True, bw=0.1, color=TSS_CLASS_PALETTE["protein_coding"], 
            label="mRNAs\n(n=%s)" % len(pc_vals), ax=ax)
sns.kdeplot(data=dpc_vals, cumulative=True, bw=0.1, color=TSS_CLASS_PALETTE["div_pc"], 
            label="div. mRNAs\n(n=%s)" % len(dpc_vals), ax=ax)
ax.set_xlabel("log(# of bp covered)")
ax.set_ylabel("cumulative density")
ax.set_ylim((0, 1.05))
plt.legend(handlelength=1)
fig.savefig("Fig_2D.pdf", bbox_inches="tight", dpi="figure")


# In[38]:


# for each group, split into tissue-sp v dynamic v ubiquitous
idxs = list(range(0, 5))
promtypes = ["Enhancer", "intergenic", "protein_coding"]
names = ["eRNAs", "lincRNAs", "mRNAs"]
df = chip_cov_exp
col = "log_bp_cov"
xlabel = "log(# of bp covered)"
xlim = (0, 5)

f, axarr = plt.subplots(nrows=1, ncols=3, sharex=False, sharey=True, figsize=(5.5, 1.6))

for i, promtype, name in zip(idxs, promtypes, names):
    print("i: %s, promtype: %s, name: %s" % (i, promtype, name))
    ax = axarr[i]
    
    promtype_vals = df[df["PromType2"] == promtype]
    ts_vals = promtype_vals[promtype_vals["tss_type"] == "tissue-specific"][col]
    ub_vals = promtype_vals[promtype_vals["tss_type"] == "ubiquitous"][col]
    dy_vals = promtype_vals[promtype_vals["tss_type"] == "dynamic"][col]
    
    ub_ts_u, ub_ts_pval = stats.mannwhitneyu(ub_vals, ts_vals, alternative="two-sided", use_continuity=False)
    print("ub/ts pval: %s" % ub_ts_pval)
    
    ub_dy_u, ub_dy_pval = stats.mannwhitneyu(ub_vals, dy_vals, alternative="two-sided", use_continuity=False)
    print("ub/dy pval: %s" % ub_dy_pval)
    
    ts_dy_u, ts_dy_pval = stats.mannwhitneyu(ts_vals, dy_vals, alternative="two-sided", use_continuity=False)
    print("ts/dy pval: %s" % ts_dy_pval)
    
    sns.kdeplot(data=ts_vals, cumulative=True, bw=0.1, color=TSS_CLASS_PALETTE[promtype], linestyle="dashed",
                label="tissue-sp.\n(n=%s)" % len(ts_vals), ax=ax)
    sns.kdeplot(data=ub_vals, cumulative=True, bw=0.1, color=TSS_CLASS_PALETTE[promtype], linestyle="solid",
                label="ubiq.\n(n=%s)" % len(ub_vals), ax=ax)
    sns.kdeplot(data=dy_vals, cumulative=True, bw=0.1, color=TSS_CLASS_PALETTE[promtype], linestyle="dotted",
                label="dynamic\n(n=%s)" % len(dy_vals), ax=ax)
    
    ax.set_title("%s (%s total)" % (name, len(promtype_vals)))
    ax.set_xlabel(xlabel)
    ax.legend(loc="bottom right", handlelength=2)
    ax.set_xlim(xlim)
    ax.set_ylim((0, 1.05))
    
    if i == 0:
        ax.set_ylabel("cumulative density")
        
plt.subplots_adjust(wspace=0.1)
f.savefig("Fig_2E.pdf", bbox_inches="tight", dpi="figure")


# In[41]:


enh_vals = fimo_chip_cov[fimo_chip_cov["PromType2"] == "Enhancer"]["log_max_cov"]
linc_vals = fimo_chip_cov[fimo_chip_cov["PromType2"] == "intergenic"]["log_max_cov"]
dlnc_vals = fimo_chip_cov[fimo_chip_cov["PromType2"] == "div_lnc"]["log_max_cov"]
pc_vals = fimo_chip_cov[fimo_chip_cov["PromType2"] == "protein_coding"]["log_max_cov"]
dpc_vals = fimo_chip_cov[fimo_chip_cov["PromType2"] == "div_pc"]["log_max_cov"]

fig = plt.figure(figsize=(2.25, 1.6))
ax = sns.kdeplot(data=enh_vals, cumulative=True, bw=0.1, color=TSS_CLASS_PALETTE["Enhancer"], 
                 label="eRNAs (n=%s)" % len(enh_vals))
sns.kdeplot(data=linc_vals, cumulative=True, bw=0.1, color=TSS_CLASS_PALETTE["intergenic"], 
            label="lincRNAs (n=%s)" % len(linc_vals), ax=ax)
sns.kdeplot(data=dlnc_vals, cumulative=True, bw=0.1, color=TSS_CLASS_PALETTE["div_lnc"], 
            label="div. lncRNAs (n=%s)" % len(dlnc_vals), ax=ax)
sns.kdeplot(data=pc_vals, cumulative=True, bw=0.1, color=TSS_CLASS_PALETTE["protein_coding"], 
            label="mRNAs (n=%s)" % len(pc_vals), ax=ax)
sns.kdeplot(data=dpc_vals, cumulative=True, bw=0.1, color=TSS_CLASS_PALETTE["div_pc"], 
            label="div. mRNAs (n=%s)" % len(dpc_vals), ax=ax)
ax.set_xlabel("log(max coverage)")
ax.set_ylabel("cumulative density")
ax.set_ylim((0, 1.05))
ax.set_xlim((0.5, 3.45))
plt.legend(handlelength=1)

fig.savefig("Fig_2F.pdf", bbox_inches="tight", dpi="figure")


# In[46]:


# for each group, split into tissue-sp v dynamic v ubiquitous
idxs = list(range(0, 5))
promtypes = ["Enhancer", "intergenic", "protein_coding"]
names = ["eRNAs", "lincRNAs", "mRNAs"]
df = chip_cov_exp
col = "log_max_cov"
xlabel = "log(max coverage)"
xlim = (-0.95, 3.1)

f, axarr = plt.subplots(nrows=1, ncols=3, sharex=False, sharey=True, figsize=(5.5, 1.6))

for i, promtype, name in zip(idxs, promtypes, names):
    print("i: %s, promtype: %s, name: %s" % (i, promtype, name))
    ax = axarr[i]
    
    promtype_vals = df[df["PromType2"] == promtype]
    ts_vals = promtype_vals[promtype_vals["tss_type"] == "tissue-specific"][col]
    ub_vals = promtype_vals[promtype_vals["tss_type"] == "ubiquitous"][col]
    dy_vals = promtype_vals[promtype_vals["tss_type"] == "dynamic"][col]
    
    ub_ts_u, ub_ts_pval = stats.mannwhitneyu(ub_vals, ts_vals, alternative="two-sided", use_continuity=False)
    print("ub/ts pval: %s" % ub_ts_pval)
    
    ub_dy_u, ub_dy_pval = stats.mannwhitneyu(ub_vals, dy_vals, alternative="two-sided", use_continuity=False)
    print("ub/dy pval: %s" % ub_dy_pval)
    
    ts_dy_u, ts_dy_pval = stats.mannwhitneyu(ts_vals, dy_vals, alternative="two-sided", use_continuity=False)
    print("ts/dy pval: %s" % ts_dy_pval)
    
    sns.kdeplot(data=ts_vals, cumulative=True, bw=0.1, color=TSS_CLASS_PALETTE[promtype], linestyle="dashed",
                label="tissue-sp.\n(n=%s)" % len(ts_vals), ax=ax)
    sns.kdeplot(data=ub_vals, cumulative=True, bw=0.1, color=TSS_CLASS_PALETTE[promtype], linestyle="solid",
                label="ubiq.\n(n=%s)" % len(ub_vals), ax=ax)
    sns.kdeplot(data=dy_vals, cumulative=True, bw=0.1, color=TSS_CLASS_PALETTE[promtype], linestyle="dotted",
                label="dynamic\n(n=%s)" % len(dy_vals), ax=ax)
    
    ax.set_title("%s (%s total)" % (name, len(promtype_vals)))
    ax.set_xlabel(xlabel)
    ax.legend(loc="upper left", handlelength=1.8)
    ax.set_ylim((0, 1.05))
    ax.set_xlim(xlim)
    
    if i == 0:
        ax.set_ylabel("cumulative density")
plt.subplots_adjust(wspace=0.1)
f.savefig("Fig_2G.pdf", bbox_inches="tight", dpi="figure")


# ### clustered motifs

# In[47]:


enh_vals = fimo_clust_cov[fimo_clust_cov["PromType2"] == "Enhancer"]["log_bp_cov"]
linc_vals = fimo_clust_cov[fimo_clust_cov["PromType2"] == "intergenic"]["log_bp_cov"]
dlnc_vals = fimo_clust_cov[fimo_clust_cov["PromType2"] == "div_lnc"]["log_bp_cov"]
pc_vals = fimo_clust_cov[fimo_clust_cov["PromType2"] == "protein_coding"]["log_bp_cov"]
dpc_vals = fimo_clust_cov[fimo_clust_cov["PromType2"] == "div_pc"]["log_bp_cov"]

fig = plt.figure(figsize=(2.25, 1.6))
ax = sns.kdeplot(data=enh_vals, cumulative=True, bw=0.1, color=TSS_CLASS_PALETTE["Enhancer"], 
                 label="eRNAs\n(n=%s)" % len(enh_vals))
sns.kdeplot(data=linc_vals, cumulative=True, bw=0.1, color=TSS_CLASS_PALETTE["intergenic"], 
            label="lincRNAs\n(n=%s)" % len(linc_vals), ax=ax)
sns.kdeplot(data=dlnc_vals, cumulative=True, bw=0.1, color=TSS_CLASS_PALETTE["div_lnc"], 
            label="div. lncRNAs\n(n=%s)" % len(dlnc_vals), ax=ax)
sns.kdeplot(data=pc_vals, cumulative=True, bw=0.1, color=TSS_CLASS_PALETTE["protein_coding"], 
            label="mRNAs\n(n=%s)" % len(pc_vals), ax=ax)
sns.kdeplot(data=dpc_vals, cumulative=True, bw=0.1, color=TSS_CLASS_PALETTE["div_pc"], 
            label="div. mRNAs\n(n=%s)" % len(dpc_vals), ax=ax)
ax.set_xlabel("log(# of bp covered by non-redundant 8mer motifs)")
ax.set_ylabel("cumulative density")
ax.set_ylim((0, 1.05))
plt.legend(handlelength=1)
fig.savefig("Fig_S9C_1.pdf", bbox_inches="tight", dpi="figure")


# In[48]:


# for each group, split into tissue-sp v dynamic v ubiquitous
idxs = list(range(0, 5))
promtypes = ["Enhancer", "intergenic", "protein_coding"]
names = ["eRNAs", "lincRNAs", "mRNAs"]
df = cluster_cov_exp
col = "log_bp_cov"
xlabel = "log(# of bp covered by non-redundant 8mer motifs)"
xlim = (0, 5)

f, axarr = plt.subplots(nrows=1, ncols=3, sharex=False, sharey=True, figsize=(5.5, 1.6))

for i, promtype, name in zip(idxs, promtypes, names):
    print("i: %s, promtype: %s, name: %s" % (i, promtype, name))
    ax = axarr[i]
    
    promtype_vals = df[df["PromType2"] == promtype]
    ts_vals = promtype_vals[promtype_vals["tss_type"] == "tissue-specific"][col]
    ub_vals = promtype_vals[promtype_vals["tss_type"] == "ubiquitous"][col]
    dy_vals = promtype_vals[promtype_vals["tss_type"] == "dynamic"][col]
    
    ub_ts_u, ub_ts_pval = stats.mannwhitneyu(ub_vals, ts_vals, alternative="two-sided", use_continuity=False)
    print("ub/ts pval: %s" % ub_ts_pval)
    
    ub_dy_u, ub_dy_pval = stats.mannwhitneyu(ub_vals, dy_vals, alternative="two-sided", use_continuity=False)
    print("ub/dy pval: %s" % ub_dy_pval)
    
    ts_dy_u, ts_dy_pval = stats.mannwhitneyu(ts_vals, dy_vals, alternative="two-sided", use_continuity=False)
    print("ts/dy pval: %s" % ts_dy_pval)
    
    sns.kdeplot(data=ts_vals, cumulative=True, bw=0.1, color=TSS_CLASS_PALETTE[promtype], linestyle="dashed",
                label="tissue-sp.\n(n=%s)" % len(ts_vals), ax=ax)
    sns.kdeplot(data=ub_vals, cumulative=True, bw=0.1, color=TSS_CLASS_PALETTE[promtype], linestyle="solid",
                label="ubiq.\n(n=%s)" % len(ub_vals), ax=ax)
    sns.kdeplot(data=dy_vals, cumulative=True, bw=0.1, color=TSS_CLASS_PALETTE[promtype], linestyle="dotted",
                label="dynamic\n(n=%s)" % len(dy_vals), ax=ax)
    
    ax.set_title("%s (%s total)" % (name, len(promtype_vals)))
    ax.set_xlabel(xlabel)
    ax.legend(loc="bottom right", handlelength=2)
    ax.set_xlim(xlim)
    ax.set_ylim((0, 1.05))
    
    if i == 0:
        ax.set_ylabel("cumulative density")
plt.subplots_adjust(wspace=0.1)


# In[50]:


enh_vals = fimo_clust_cov[fimo_clust_cov["PromType2"] == "Enhancer"]["log_max_cov"]
linc_vals = fimo_clust_cov[fimo_clust_cov["PromType2"] == "intergenic"]["log_max_cov"]
dlnc_vals = fimo_clust_cov[fimo_clust_cov["PromType2"] == "div_lnc"]["log_max_cov"]
pc_vals = fimo_clust_cov[fimo_clust_cov["PromType2"] == "protein_coding"]["log_max_cov"]
dpc_vals = fimo_clust_cov[fimo_clust_cov["PromType2"] == "div_pc"]["log_max_cov"]

fig = plt.figure(figsize=(2.25, 1.6))
ax = sns.kdeplot(data=enh_vals, cumulative=True, bw=0.1, color=TSS_CLASS_PALETTE["Enhancer"], 
                 label="eRNAs (n=%s)" % len(enh_vals))
sns.kdeplot(data=linc_vals, cumulative=True, bw=0.1, color=TSS_CLASS_PALETTE["intergenic"], 
            label="lincRNAs (n=%s)" % len(linc_vals), ax=ax)
sns.kdeplot(data=dlnc_vals, cumulative=True, bw=0.1, color=TSS_CLASS_PALETTE["div_lnc"], 
            label="div. lncRNAs (n=%s)" % len(dlnc_vals), ax=ax)
sns.kdeplot(data=pc_vals, cumulative=True, bw=0.1, color=TSS_CLASS_PALETTE["protein_coding"], 
            label="mRNAs (n=%s)" % len(pc_vals), ax=ax)
sns.kdeplot(data=dpc_vals, cumulative=True, bw=0.1, color=TSS_CLASS_PALETTE["div_pc"], 
            label="div. mRNAs (n=%s)" % len(dpc_vals), ax=ax)
ax.set_xlabel("log(max coverage of non-redundant 8mer motifs)")
ax.set_ylabel("cumulative density")
ax.set_ylim((0, 1.05))
plt.legend(handlelength=0.8)
fig.savefig("Fig_S9C_2.pdf", bbox_inches="tight", dpi="figure")


# In[51]:


# for each group, split into tissue-sp v dynamic v ubiquitous
idxs = list(range(0, 5))
promtypes = ["Enhancer", "intergenic", "protein_coding"]
names = ["eRNAs", "lincRNAs", "mRNAs"]
df = cluster_cov_exp
col = "log_max_cov"
xlabel = "log(max coverage of non-redundant 8mer motifs)"
xlim = (-0.5, 3)

f, axarr = plt.subplots(nrows=1, ncols=3, sharex=False, sharey=True, figsize=(5.5, 1.6))

for i, promtype, name in zip(idxs, promtypes, names):
    print("i: %s, promtype: %s, name: %s" % (i, promtype, name))
    ax = axarr[i]
    
    promtype_vals = df[df["PromType2"] == promtype]
    ts_vals = promtype_vals[promtype_vals["tss_type"] == "tissue-specific"][col]
    ub_vals = promtype_vals[promtype_vals["tss_type"] == "ubiquitous"][col]
    dy_vals = promtype_vals[promtype_vals["tss_type"] == "dynamic"][col]
    
    ub_ts_u, ub_ts_pval = stats.mannwhitneyu(ub_vals, ts_vals, alternative="two-sided", use_continuity=False)
    print("ub/ts pval: %s" % ub_ts_pval)
    
    ub_dy_u, ub_dy_pval = stats.mannwhitneyu(ub_vals, dy_vals, alternative="two-sided", use_continuity=False)
    print("ub/dy pval: %s" % ub_dy_pval)
    
    ts_dy_u, ts_dy_pval = stats.mannwhitneyu(ts_vals, dy_vals, alternative="two-sided", use_continuity=False)
    print("ts/dy pval: %s" % ts_dy_pval)
    
    sns.kdeplot(data=ts_vals, cumulative=True, bw=0.1, color=TSS_CLASS_PALETTE[promtype], linestyle="dashed",
                label="tissue-sp.\n(n=%s)" % len(ts_vals), ax=ax)
    sns.kdeplot(data=ub_vals, cumulative=True, bw=0.1, color=TSS_CLASS_PALETTE[promtype], linestyle="solid",
                label="ubiq.\n(n=%s)" % len(ub_vals), ax=ax)
    sns.kdeplot(data=dy_vals, cumulative=True, bw=0.1, color=TSS_CLASS_PALETTE[promtype], linestyle="dotted",
                label="dynamic\n(n=%s)" % len(dy_vals), ax=ax)
    
    ax.set_title("%s (%s total)" % (name, len(promtype_vals)))
    ax.set_xlabel(xlabel)
    ax.legend(loc="bottom right", handlelength=2)
    ax.set_xlim(xlim)
    ax.set_ylim((0,1.05))
    
    if i == 0:
        ax.set_ylabel("cumulative density")
plt.subplots_adjust(wspace=0.1)


# ## 2. cluster the motifs using MoSBAT output

# In[52]:


corr.set_index(corr["Motif"], inplace=True)
corr.drop("Motif", axis=1, inplace=True)
corr.head()


# In[53]:


row_linkage = hierarchy.linkage(distance.pdist(corr, 'correlation'), method="average")
col_linkage = hierarchy.linkage(distance.pdist(corr.T, 'correlation'), method="average")


# In[54]:


dists = plot_dendrogram(row_linkage, 0.4, "correlation")


# In[55]:


clusters = hierarchy.fcluster(row_linkage, 0.1, criterion="distance")


# In[56]:


print("n clusters: %s" % np.max(clusters))


# In[57]:


cluster_map = pd.DataFrame.from_dict(dict(zip(list(corr.index), clusters)), orient="index")
cluster_map.columns = ["cluster"]
cluster_map.head()


# In[58]:


cluster_map.loc["KLF5"]


# ## 3. plot clustered motif heatmap

# In[59]:


colors = sns.husl_palette(np.max(clusters), s=0.75)
shuffle(colors)
lut = dict(zip(range(np.min(clusters), np.max(clusters)+1), colors))
row_colors = cluster_map["cluster"].map(lut)


# In[60]:


cmap = sns.cubehelix_palette(8, start=.5, light=1, dark=0.25, hue=0.9, rot=-0.75, as_cmap=True)


# In[61]:


cg = sns.clustermap(corr, method="average", row_linkage=row_linkage, robust=True,
                    col_linkage=col_linkage, cmap=cmap, figsize=(5, 5), row_colors=row_colors,
                    linewidths=0, rasterized=True)
cg.savefig("Fig_S9A.pdf", bbox_inches="tight", dpi="figure")


# In[62]:


cluster_map.to_csv("../../misc/02__mosbat/cluster_map.txt", sep="\t", index=True)


# ## 4. re-plot # bp covered and max coverage per biotype *after* clustering
# note that i sent the cluster results to marta, who re-ran her coverage scripts using them, and i re-upload them in this notebook (so in real life there is a break between the above part and the following part of this notebook)

# In[63]:


fimo_mosbat_cov.head()


# In[67]:


enh_vals = fimo_mosbat_cov[fimo_mosbat_cov["PromType2"] == "Enhancer"]["log_bp_cov"]
linc_vals = fimo_mosbat_cov[fimo_mosbat_cov["PromType2"] == "intergenic"]["log_bp_cov"]
dlnc_vals = fimo_mosbat_cov[fimo_mosbat_cov["PromType2"] == "div_lnc"]["log_bp_cov"]
pc_vals = fimo_mosbat_cov[fimo_mosbat_cov["PromType2"] == "protein_coding"]["log_bp_cov"]
dpc_vals = fimo_mosbat_cov[fimo_mosbat_cov["PromType2"] == "div_pc"]["log_bp_cov"]

fig = plt.figure(figsize=(2.25, 1.6))
ax = sns.kdeplot(data=enh_vals, cumulative=True, color=TSS_CLASS_PALETTE["Enhancer"], 
                 label="eRNAs (n=%s)" % len(enh_vals))
sns.kdeplot(data=linc_vals, cumulative=True, color=TSS_CLASS_PALETTE["intergenic"], 
            label="lincRNAs (n=%s)" % len(linc_vals), ax=ax)
sns.kdeplot(data=dlnc_vals, cumulative=True, color=TSS_CLASS_PALETTE["div_lnc"], 
            label="div. lncRNAs (n=%s)" % len(dlnc_vals), ax=ax)
sns.kdeplot(data=pc_vals, cumulative=True, color=TSS_CLASS_PALETTE["protein_coding"], 
            label="mRNAs (n=%s)" % len(pc_vals), ax=ax)
sns.kdeplot(data=dpc_vals, cumulative=True, color=TSS_CLASS_PALETTE["div_pc"], 
            label="div. mRNAs (n=%s)" % len(dpc_vals), ax=ax)
ax.set_xlabel("log(# of bp covered, deduped by motif cluster)")
ax.set_ylabel("cumulative density")
ax.set_ylim((0, 1.05))
ax.set_xlim((2.5, 5))
plt.legend(handlelength=1)
fig.savefig("Fig_S9B_1.pdf", bbox_inches="tight", dpi="figure")


# In[69]:


# for each group, split into tissue-sp v dynamic v ubiquitous
idxs = list(range(0, 5))
promtypes = ["Enhancer", "intergenic", "protein_coding"]
names = ["eRNAs", "lincRNAs", "mRNAs"]
df = mosbat_cov_exp
col = "log_bp_cov"
xlabel = "log(# of bp covered, deduped by motif cluster)"
xlim = (2.5, 5)

f, axarr = plt.subplots(nrows=1, ncols=3, sharex=False, sharey=True, figsize=(5.5, 1.6))

for i, promtype, name in zip(idxs, promtypes, names):
    print("i: %s, promtype: %s, name: %s" % (i, promtype, name))
    ax = axarr[i]
    
    promtype_vals = df[df["PromType2"] == promtype]
    ts_vals = promtype_vals[promtype_vals["tss_type"] == "tissue-specific"][col]
    ub_vals = promtype_vals[promtype_vals["tss_type"] == "ubiquitous"][col]
    dy_vals = promtype_vals[promtype_vals["tss_type"] == "dynamic"][col]
    
    sns.kdeplot(data=ts_vals, cumulative=True, color=TSS_CLASS_PALETTE[promtype], linestyle="dashed",
                label="tissue-sp.\n(n=%s)" % len(ts_vals), ax=ax)
    sns.kdeplot(data=ub_vals, cumulative=True, color=TSS_CLASS_PALETTE[promtype], linestyle="solid",
                label="ubiq.\n(n=%s)" % len(ub_vals), ax=ax)
    sns.kdeplot(data=dy_vals, cumulative=True, color=TSS_CLASS_PALETTE[promtype], linestyle="dotted",
                label="dynamic\n(n=%s)" % len(dy_vals), ax=ax)
    
    ax.set_title("%s (%s total)" % (name, len(promtype_vals)))
    ax.set_xlabel(xlabel)
    ax.legend(loc="bottom right", handlelength=2)
    ax.set_xlim(xlim)
    ax.set_ylim((0,1.05))
    
    if i == 0:
        ax.set_ylabel("cumulative density")
plt.subplots_adjust(wspace=0.1)


# In[75]:


enh_vals = fimo_mosbat_cov[fimo_mosbat_cov["PromType2"] == "Enhancer"]["log_max_cov"]
linc_vals = fimo_mosbat_cov[fimo_mosbat_cov["PromType2"] == "intergenic"]["log_max_cov"]
dlnc_vals = fimo_mosbat_cov[fimo_mosbat_cov["PromType2"] == "div_lnc"]["log_max_cov"]
pc_vals = fimo_mosbat_cov[fimo_mosbat_cov["PromType2"] == "protein_coding"]["log_max_cov"]
dpc_vals = fimo_mosbat_cov[fimo_mosbat_cov["PromType2"] == "div_pc"]["log_max_cov"]

fig = plt.figure(figsize=(2.25, 1.6))
ax = sns.kdeplot(data=enh_vals, cumulative=True, color=TSS_CLASS_PALETTE["Enhancer"], 
                 label="eRNAs (n=%s)" % len(enh_vals))
sns.kdeplot(data=linc_vals, cumulative=True, color=TSS_CLASS_PALETTE["intergenic"], 
            label="lincRNAs (n=%s)" % len(linc_vals), ax=ax)
sns.kdeplot(data=dlnc_vals, cumulative=True, color=TSS_CLASS_PALETTE["div_lnc"], 
            label="div. lncRNAs (n=%s)" % len(dlnc_vals), ax=ax)
sns.kdeplot(data=pc_vals, cumulative=True, color=TSS_CLASS_PALETTE["protein_coding"], 
            label="mRNAs (n=%s)" % len(pc_vals), ax=ax)
sns.kdeplot(data=dpc_vals, cumulative=True, color=TSS_CLASS_PALETTE["div_pc"], 
            label="div. mRNAs (n=%s)" % len(dpc_vals), ax=ax)
ax.set_xlabel("log(max coverage, deduped by motif cluster)")
ax.set_ylabel("cumulative density")
plt.xlim((-1.35, 3.75))
plt.ylim((0,1.05))
plt.legend(handlelength=0.8)
fig.savefig("Fig_S9B_2.pdf", bbox_inches="tight", dpi="figure")


# In[76]:


# for each group, split into tissue-sp v dynamic v ubiquitous
idxs = list(range(0, 5))
promtypes = ["Enhancer", "intergenic", "protein_coding"]
names = ["eRNAs", "lincRNAs", "mRNAs"]
df = mosbat_cov_exp
col = "log_max_cov"
xlabel = "log(max coverage, deduped by motif cluster)"
xlim = (-1, 4)

f, axarr = plt.subplots(nrows=1, ncols=3, sharex=False, sharey=True, figsize=(5.5, 1.6))

for i, promtype, name in zip(idxs, promtypes, names):
    print("i: %s, promtype: %s, name: %s" % (i, promtype, name))
    ax = axarr[i]
    
    promtype_vals = df[df["PromType2"] == promtype]
    ts_vals = promtype_vals[promtype_vals["tss_type"] == "tissue-specific"][col]
    ub_vals = promtype_vals[promtype_vals["tss_type"] == "ubiquitous"][col]
    dy_vals = promtype_vals[promtype_vals["tss_type"] == "dynamic"][col]
    
    sns.kdeplot(data=ts_vals, cumulative=True, color=TSS_CLASS_PALETTE[promtype], linestyle="dashed",
                label="tissue-sp.\n(n=%s)" % len(ts_vals), ax=ax)
    sns.kdeplot(data=ub_vals, cumulative=True, color=TSS_CLASS_PALETTE[promtype], linestyle="solid",
                label="ubiq.\n(n=%s)" % len(ub_vals), ax=ax)
    sns.kdeplot(data=dy_vals, cumulative=True, color=TSS_CLASS_PALETTE[promtype], linestyle="dotted",
                label="dynamic\n(n=%s)" % len(dy_vals), ax=ax)
    
    ax.set_title("%s (%s total)" % (name, len(promtype_vals)))
    ax.set_xlabel(xlabel)
    ax.legend(loc="bottom right", handlelength=2)
    ax.set_xlim(xlim)
    ax.set_ylim((0, 1.05))
    
    if i == 0:
        ax.set_ylabel("cumulative density")
plt.subplots_adjust(wspace=0.1)


# ## 5. look at conservation of nucleotides vs. motif coverage

# ### fimo only

# In[77]:


res_dict = {}
df = fimo_phylop
nuc_cols = list(np.arange(-150, 150, step=1))

prev_max = 0
for max_motifs in [1, 10, 30, 86]:
    sub = df[(df["n_ov"] > prev_max) & (df["n_ov"] <= max_motifs)]
    n_motifs = len(sub)
    nums = np.asarray(sub[nuc_cols])
    
    avg = np.nanmean(nums, axis=0)
    std = np.nanstd(nums, axis=0)
    
    y1 = avg - std
    y2 = avg + std
    
    res_dict[max_motifs] = {"n_motifs": n_motifs, "avg": avg, "y1": y1, "y2": y2}
    prev_max = max_motifs


# In[78]:


fig = plt.figure(figsize=(5.75,1.6))
palette = {1: sns.cubehelix_palette(4, start=.75, rot=-.75)[0], 10: sns.cubehelix_palette(4, start=.75, rot=-.75)[1], 
           30: sns.cubehelix_palette(4, start=.75, rot=-.75)[2], 86: sns.cubehelix_palette(4, start=.75, rot=-.75)[3]}
labels = ["1 motif", "2-10 motifs", "11-30 motifs", "31+ motifs"]

for n, label in zip(res_dict.keys(), labels):
    res = res_dict[n]
    n_motifs = res["n_motifs"]
    avg = res["avg"]
    y1 = res["y1"]
    y2 = res["y2"]
    # x = signal.savgol_filter(df["mean"], 15, 1)
    # plt.fill_between(nuc_cols, y1, y2, color=palette[n], alpha=0.5)
    plt.plot(nuc_cols, avg, color=palette[n], linewidth=2, label="%s (n=%s)" % (label, n_motifs))
# plt.xlim((lower, upper))
# plt.axvline(x=-75, color="black", linestyle="dashed", linewidth=1)
# plt.axvline(x=25, color="black", linestyle="dashed", linewidth=1)
plt.legend(ncol=1, loc=1, handlelength=1)
plt.xlabel("nucleotide (0 = middle of motif)")
plt.ylabel("phylop 46-way")


# ### fimo + chip

# In[79]:


fimo_chip_phylop["n_ov"].max()


# In[80]:


res_dict = {}
df = fimo_chip_phylop
nuc_cols = list(np.arange(-75, 75, step=1))

prev_max = 0
for max_motifs in [1, 4, 8, 13]:
    sub = df[(df["n_ov"] > prev_max) & (df["n_ov"] <= max_motifs)]
    n_motifs = len(sub)
    nums = np.asarray(sub[nuc_cols])
    
    avg = np.nanmean(nums, axis=0)
    std = np.nanstd(nums, axis=0)
    
    y1 = avg - std
    y2 = avg + std
    
    res_dict[max_motifs] = {"n_motifs": n_motifs, "avg": avg, "y1": y1, "y2": y2}
    prev_max = max_motifs


# In[81]:


fig = plt.figure(figsize=(5.75,1.6))
palette = {1: sns.cubehelix_palette(4, start=.75, rot=-.75)[0], 4: sns.cubehelix_palette(4, start=.75, rot=-.75)[1], 
           8: sns.cubehelix_palette(4, start=.75, rot=-.75)[2], 13: sns.cubehelix_palette(4, start=.75, rot=-.75)[3]}
labels = ["1 motif", "2-4 motifs", "5-8 motifs", "9+ motifs"]

for n, label in zip(res_dict.keys(), labels):
    res = res_dict[n]
    n_motifs = res["n_motifs"]
    avg = res["avg"]
    y1 = res["y1"]
    y2 = res["y2"]
    # x = signal.savgol_filter(df["mean"], 15, 1)
    # plt.fill_between(nuc_cols, y1, y2, color=palette[n], alpha=0.5)
    plt.plot(nuc_cols, avg, color=palette[n], linewidth=2, label="%s (n=%s)" % (label, n_motifs))
# plt.xlim((lower, upper))
# plt.axvline(x=-75, color="black", linestyle="dashed", linewidth=1)
# plt.axvline(x=25, color="black", linestyle="dashed", linewidth=1)
plt.legend(ncol=1, loc=1, handlelength=1)
plt.xlabel("nucleotide (0 = middle of motif)")
plt.ylabel("phylop 46-way")
fig.savefig("Fig_2H.pdf", bbox_inches="tight", dpi="figure")


# ### clusters

# In[82]:


fimo_clust_phylop["n_ov"].max()


# In[83]:


res_dict = {}
df = fimo_clust_phylop
nuc_cols = list(np.arange(-75, 75, step=1))

prev_max = 0
for max_motifs in [1, 3, 6, 13]:
    sub = df[(df["n_ov"] > prev_max) & (df["n_ov"] <= max_motifs)]
    n_motifs = len(sub)
    nums = np.asarray(sub[nuc_cols])
    
    avg = np.nanmean(nums, axis=0)
    std = np.nanstd(nums, axis=0)
    
    y1 = avg - std
    y2 = avg + std
    
    res_dict[max_motifs] = {"n_motifs": n_motifs, "avg": avg, "y1": y1, "y2": y2}
    prev_max = max_motifs


# In[84]:


fig = plt.figure(figsize=(5.75,1.6))
palette = {1: sns.cubehelix_palette(4, start=.75, rot=-.75)[0], 3: sns.cubehelix_palette(4, start=.75, rot=-.75)[1], 
           6: sns.cubehelix_palette(4, start=.75, rot=-.75)[2], 13: sns.cubehelix_palette(4, start=.75, rot=-.75)[3]}
labels = ["1 motif", "2-3 motifs", "4-6 motifs", "7+ motifs"]

for n, label in zip(res_dict.keys(), labels):
    res = res_dict[n]
    n_motifs = res["n_motifs"]
    avg = res["avg"]
    y1 = res["y1"]
    y2 = res["y2"]
    # x = signal.savgol_filter(df["mean"], 15, 1)
    # plt.fill_between(nuc_cols, y1, y2, color=palette[n], alpha=0.5)
    plt.plot(nuc_cols, avg, color=palette[n], linewidth=2, label="%s (n=%s)" % (label, n_motifs))
# plt.xlim((lower, upper))
# plt.axvline(x=-75, color="black", linestyle="dashed", linewidth=1)
# plt.axvline(x=25, color="black", linestyle="dashed", linewidth=1)
plt.legend(ncol=1, loc=2, handlelength=1)
plt.xlabel("nucleotide (0 = middle of motif)")
plt.ylabel("phylop 46-way")


# ## 6. look at motif overlap vs. dnase accessibility

# In[85]:


dnase_merged = dnase.merge(motif_cov_exp, on=["unique_id", "PromType2"])
dnase_merged["log_n_accessible"] = np.log10(dnase_merged["n_accessible"]+1)
dnase_merged.head()


# In[86]:


fig = plt.figure(figsize=(1.2, 1.2))
ax = sns.kdeplot(dnase_merged["log_max_cov"], dnase_merged["log_n_accessible"], 
                 cmap=sns.light_palette("firebrick", as_cmap=True), 
                 shade=True, shade_lowest=False, bw=0.13)
ax.set_ylabel("log(count of DNase\naccessible samples)")
ax.set_xlabel("log(max overlapping motifs)")
ax.set_ylim((0.5, 2.5))

r, p = stats.spearmanr(dnase_merged["log_max_cov"], dnase_merged["log_n_accessible"])
print("r: %s, spearman p: %s" % (r, p))
ax.annotate("r = {:.2f}".format(r), xy=(.05, .9), xycoords=ax.transAxes, fontsize=fontsize)

# add n-value
ax.annotate("n = %s" % len(dnase_merged), ha="right", xy=(.96, .9), xycoords=ax.transAxes, 
            fontsize=fontsize)
#fig.savefig("max_cov.v.dnase.pdf", bbox_inches="tight", dpi="figure")


# In[87]:


fig = plt.figure(figsize=(1.2, 1.2))
ax = sns.kdeplot(dnase_merged["log_bp_cov"], dnase_merged["log_n_accessible"], 
                 cmap="Blues", 
                 shade=True, shade_lowest=False, bw=0.13)
ax.set_ylabel("log(count of DNase\naccessible samples)")
ax.set_xlabel("log(number of bp covered by motif)")
ax.set_ylim((0.5, 2.5))
ax.set_xlim((3, 5.5))

r, p = stats.spearmanr(dnase_merged["log_bp_cov"], dnase_merged["log_n_accessible"])
print("r: %s, spearman p: %s" % (r, p))
ax.annotate("r = {:.2f}".format(r), xy=(.05, .9), xycoords=ax.transAxes, fontsize=fontsize)

# add n-value
ax.annotate("n = %s" % len(dnase_merged), ha="right", xy=(.96, .9), xycoords=ax.transAxes, 
            fontsize=fontsize)
#fig.savefig("bp_cov.v.dnase.pdf", bbox_inches="tight", dpi="figure")


# In[ ]:
```
