## Supplementary material for "High-throughput functional analysis of IncRNA core promoters elucidatesrules governing tissue-specificity": 08__deletions.py

```
# coding: utf-8

# # 08__deletions
# # analyzing single-nucleotide deletions MPRA and preparing input for MIND
# 
# in this notebook, i do two things: (1) i calculate the effect size of deletions (log2 foldchange between the deletion activity and the reference activity) from the deletion MPRA and (2) i compare those effect sizes to the number of TF motifs that are computationally predicted by FIMO to be gained or lost by the deletion. note that these FIMO predictions are done separately. 
# 
# ------
# 
# figures in this notebook:
# - **Fig 3B, S13A**: scatter plots showing the comparison between deletion effect sizes and motif disruptions
# - **Fig 3C, S13B**: specific examples of deletion profiles across a sequence (HOTAIR and DLEU1) and how they compare to motif disruption profiles

# In[1]:


import warnings
warnings.filterwarnings('ignore')

import pandas as pd
import numpy as np
import matplotlib as mpl
import matplotlib.pyplot as plt
import seaborn as sns
import sys
import time

from itertools import chain
from decimal import Decimal
from matplotlib import gridspec
from random import shuffle
from scipy import stats
from scipy import signal
from scipy.spatial import distance
from scipy.cluster import hierarchy
from scipy.stats import spearmanr
from scipy.stats import pearsonr
from statsmodels.sandbox.stats import multicomp

# import utils
sys.path.append("../../utils")
from plotting_utils import *
from misc_utils import *
from norm_utils import *
from del_utils import *

get_ipython().magic('matplotlib inline')


# In[2]:


sns.set(**PAPER_PRESET)
fontsize = PAPER_FONTSIZE


# ## variables

# In[3]:


activ_dir = "../../data/02__activs"
pval_dir = "../../data/03__pvals"
index_dir = "../../data/00__index"


# In[4]:


pool2_hepg2_barc_activ_f = "%s/POOL2__pMPRA1__HepG2__activities_per_barcode.txt" % (activ_dir)
pool2_k562_barc_activ_f = "%s/POOL2__pMPRA1__K562__activities_per_barcode.txt" % (activ_dir)

pool2_hepg2_elem_activ_f = "%s/POOL2__pMPRA1__HepG2__activities_per_element.txt" % (activ_dir)
pool2_k562_elem_activ_f = "%s/POOL2__pMPRA1__K562__activities_per_element.txt" % (activ_dir)

pool2_hepg2_pvals_f = "%s/POOL2__pMPRA1__HepG2__pvals.txt" % (pval_dir)
pool2_k562_pvals_f = "%s/POOL2__pMPRA1__K562__pvals.txt" % (pval_dir)


# In[5]:


pool2_index_f = "%s/dels_oligo_pool.index.txt" % (index_dir)
pool2_index_del_f = "%s/dels_oligo_pool.index.with_deletion_info.txt" % (index_dir)


# In[6]:


annot_f = "%s/tssid_all_biotypes.txt" % (index_dir)
tfbs_f = "../../data/04__coverage/pool2_n_motifs_map.txt"


# ## 1. import data

# In[7]:


hepg2_elem_res = pd.read_table(pool2_hepg2_elem_activ_f, sep="\t")
k562_elem_res = pd.read_table(pool2_k562_elem_activ_f, sep="\t")
hepg2_elem_res.head()


# In[8]:


hepg2_barc_res = pd.read_table(pool2_hepg2_barc_activ_f, sep="\t")
k562_barc_res = pd.read_table(pool2_k562_barc_activ_f, sep="\t")
hepg2_barc_res.head()


# In[9]:


hepg2_wt_pvals = pd.read_table(pool2_hepg2_pvals_f, sep="\t")
k562_wt_pvals = pd.read_table(pool2_k562_pvals_f, sep="\t")
hepg2_wt_pvals.head()


# In[10]:


index = pd.read_table(pool2_index_f, sep="\t")
index.head()


# In[11]:


tfbs = pd.read_table(tfbs_f, sep="\t", header=None)
tfbs.columns = ["unique_id", "n_tfs"]
tfbs.head()


# ## 2. add required deletion information to index

# In[12]:


# first create a dict of del seq : wt seq map
index_dels = index[index["oligo_type"].str.contains("DELETION")][["oligo_type", "tile_name", "element"]].drop_duplicates()
print("mapping %s unique deletion sequences" % (len(index_dels)))
index_wt = index[index["oligo_type"] == "WILDTYPE"][["oligo_type", "tile_name", "element"]].drop_duplicates()
index_wt_w_snp = index[index["oligo_type"] == "WILDTYPE_BUT_HAS_SNP"][["oligo_type", "tile_name", "element"]].drop_duplicates()
index_fl = index[index["oligo_type"] == "FLIPPED"][["oligo_type", "tile_name", "element"]].drop_duplicates()
dels_dict = {}
for i, row in index_dels.iterrows():
    if "WILDTYPE_BUT_HAS_SNP" in row.oligo_type:
        seq = index_wt_w_snp[index_wt_w_snp["tile_name"] == row.tile_name]["element"].iloc[0]
    elif "WILDTYPE" in row.oligo_type:
        seq = index_wt[index_wt["tile_name"] == row.tile_name]["element"].iloc[0]
    elif "FLIPPED" in row.oligo_type:
        seq = index_fl[index_fl["tile_name"] == row.tile_name]["element"].iloc[0]
    dels_dict[row.element] = seq


# In[13]:


# add deletion number to index
print("finding deletion numbers...")
index["del_num"] = index.apply(get_del_num, axis=1)


# In[14]:


print("finding deletion bases...")
index["del_base"] = index.apply(get_del_base, seq_map=dels_dict, axis=1)


# In[15]:


index_elem = index.drop(["RE_count_1", "RE_count_2", "RE_count_3", "barcode", "full_oligo"], axis=1)
index_elem = index_elem.drop_duplicates(subset=["oligo_type", "seq_name", "dupe_info", "element"])
index_elem["dupe_info"] = index_elem.apply(fix_dupe_info, axis=1)
index_elem.head()


# ## 3. merge with index

# In[16]:


hepg2_barc = index.merge(hepg2_barc_res, left_on="barcode", right_on="barcode", how="left")
k562_barc = index.merge(k562_barc_res, left_on="barcode", right_on="barcode", how="left")
hepg2_barc.head()


# In[17]:


hepg2 = index_elem.merge(hepg2_elem_res, on=["unique_id", "element"], how="left")
k562 = index_elem.merge(k562_elem_res, on=["unique_id", "element"], how="left")
hepg2.head()


# In[18]:


# add overall mean column and standard dev column
hepg2_reps = [col for col in hepg2.columns if "rna" in col]
hepg2["overall_mean"] = hepg2[hepg2_reps].mean(axis=1)
hepg2["lfcSE"] = hepg2[hepg2_reps].std(axis=1)/np.sqrt(len(hepg2_reps))

hepg2_barc["overall_mean"] = hepg2_barc[hepg2_reps].mean(axis=1)
hepg2_barc["lfcSE"] = hepg2_barc[hepg2_reps].std(axis=1)/np.sqrt(len(hepg2_reps))


# In[19]:


# add overall mean column and standard dev column
k562_reps = [col for col in k562.columns if "rna" in col]
k562["overall_mean"] = k562[k562_reps].mean(axis=1)
k562["lfcSE"] = k562[k562_reps].std(axis=1)/np.sqrt(len(k562_reps))

k562_barc["overall_mean"] = k562_barc[k562_reps].mean(axis=1)
k562_barc["lfcSE"] = k562_barc[k562_reps].std(axis=1)/np.sqrt(len(k562_reps))


# ## 4. calculate p-value for deletions

# In[20]:


hepg2_barcode_value_dict = get_barcode_value_map(hepg2, hepg2_barc, hepg2_reps)


# In[21]:


k562_barcode_value_dict = get_barcode_value_map(k562, k562_barc, k562_reps)


# In[22]:


hepg2_pvals, hepg2_l2fcs = calculate_p_value(hepg2_barcode_value_dict)


# In[23]:


k562_pvals, k562_l2fcs = calculate_p_value(k562_barcode_value_dict)


# In[24]:


# combine and adjust pvals
hepg2_all_pvals = combine_and_adjust_pvals(hepg2_pvals, hepg2_l2fcs, 0.05, hepg2_reps)
hepg2_all_pvals.sample(5)


# In[25]:


# combine and adjust pvals
k562_all_pvals = combine_and_adjust_pvals(k562_pvals, k562_l2fcs, 0.05, k562_reps)
k562_all_pvals.sample(5)


# In[26]:


# add info back to original dfs
hepg2 = hepg2.merge(hepg2_all_pvals, left_on="element", right_on="index", how="left")
hepg2.head()


# In[27]:


# add info back to original dfs
k562 = k562.merge(k562_all_pvals, left_on="element", right_on="index", how="left")
k562.head()


# ## 5. wrangle deletion data into format we need

# In[28]:


unique_names = index_elem[index_elem.oligo_type.str.contains("DELETION")]["tile_name"].unique()
[x for x in unique_names if "Enhancer" in x]


# In[29]:


# first flatten the dataframe so dupes at seq level are dealt with
hepg2 = tidy_split(hepg2, "dupe_info", sep=",", keep=False)
k562 = tidy_split(k562, "dupe_info", sep=",", keep=False)
hepg2.head()


# In[30]:


# then fix deletion numbers of those that have new rows due to a dupe
hepg2["del_num_fixed"] = hepg2.apply(fix_del_num, axis=1)
k562["del_num_fixed"] = k562.apply(fix_del_num, axis=1)
hepg2.head()


# In[31]:


hepg2_dels = wrangle_deletion_data(hepg2, unique_names, hepg2_wt_pvals)
hepg2_dels["Enhancer.noflip.NA__chr11:65264635-65265953__chr11:65265021..65265229,-,2.1"].head()


# In[32]:


k562_dels = wrangle_deletion_data(k562, unique_names, k562_wt_pvals)
k562_dels["Enhancer.noflip.NA__chr11:65264635-65265953__chr11:65265021..65265229,-,2.1"].head()


# In[33]:


k562_dels["Antisense.noflip.NA__p1@MEG3__chr14:101292281..101292489,+,17.1"].head()


# ## 6. write files

# In[34]:


# hepg2
filenames = []
clean_names = []
hepg2_dels_clean = {}
path = "../../data/05__deletions/HepG2"
for key in hepg2_dels:
    df = hepg2_dels[key]
    filename, clean_name = fix_names(key, "HepG2", hepg2, NAME_DICT, LOC_DICT)
    filenames.append(filename)
    clean_names.append(clean_name)
    filename = "%s/%s" % (path, filename)
    df.to_csv(filename, sep="\t", index=False)
    hepg2_dels_clean[clean_name] = df
name_map = pd.DataFrame({"filename": filenames, "seq_name": clean_names})
name_map.to_csv("../../data/05__deletions/HepG2_filename_map.txt", sep="\t", index=False)


# In[35]:


# k562
filenames = []
clean_names = []
k562_dels_clean = {}
path = "../../data/05__deletions/K562"
for key in k562_dels:
    df = k562_dels[key]
    filename, clean_name = fix_names(key, "K562", k562, NAME_DICT, LOC_DICT)
    filenames.append(filename)
    clean_names.append(clean_name)
    filename = "%s/%s" % (path, filename)
    df.to_csv(filename, sep="\t", index=False)
    k562_dels_clean[clean_name] = df
name_map = pd.DataFrame({"filename": filenames, "seq_name": clean_names})
name_map.to_csv("../../data/05__deletions/K562_filename_map.txt", sep="\t", index=False)


# ## 7. compare deletion profiles to predicted motif profiles (seq by seq)

# In[36]:


def fix_names(row, name_dict, loc_dict):
    old_name = row["unique_id"]
    chrom = old_name.split("__")[3].split(":")[0]
    start = int(old_name.split("__")[3].split(":")[1].split("..")[0])
    end = int(old_name.split("__")[3].split(":")[1].split("..")[1].split(",")[0])
    strand = old_name.split("__")[3].split(",")[1]
    locs = "%s:%s-%s" % (chrom, start, end)
    if strand == "+":
        text_strand = "plus"
    else:
        text_strand = "minus"
    tile_num = int(old_name.split("__")[4].split(".")[1])
    
    name = old_name.split("__")[2]
    coords = old_name.split("__")[3].split(",")[0]
    try:
        gene = name.split(",")[0].split("@")[1]
        prom = name.split(",")[0].split("@")[0]
    except:
        gene = "X"
        prom = "pX"
    
    if gene not in name_dict.keys() and coords not in loc_dict.keys():
        name = "%s__%s__tile%s" % (gene, prom, tile_num)
    elif gene in name_dict.keys():
        name = "%s__%s__tile%s" % (name_dict[gene], prom, tile_num)
    elif coords in loc_dict.keys():
        name = "%s__%s__tile%s" % (loc_dict[coords], prom, tile_num)
    
    clean_name = "%s__%s" % (name, text_strand)
    return clean_name


# In[37]:


tfbs["del_id"] = tfbs.apply(fix_names, name_dict=NAME_DICT, loc_dict=LOC_DICT, axis=1)
tfbs["delpos"] = tfbs.apply(get_del_num, axis=1)
tfbs["del_num"] = tfbs.apply(get_del_num, axis=1)
tfbs.sample(5)


# In[38]:


tfbs_wt = tfbs[(~tfbs["unique_id"].str.contains("DELETION")) & 
               (~tfbs["unique_id"].str.contains("SNP_INDIV")) & 
               (~tfbs["unique_id"].str.contains("HAPLO"))]
tfbs_wt.sample(5)


# In[39]:


index_del_info = index_elem[["unique_id", "dupe_info", "del_num"]]
index_tfbs = index_del_info.merge(tfbs, on=["unique_id", "del_num"])
index_tfbs.head()


# In[40]:


index_tfbs = tidy_split(index_tfbs, "dupe_info", sep=",", keep=False)
index_tfbs.head()


# In[41]:


index_tfbs["delpos"] = index_tfbs.apply(fix_del_num, axis=1)
index_tfbs.head()


# In[42]:


hepg2_dels_fixed = {}
k562_dels_fixed = {}
for dels_dict, fixed_dict in zip([hepg2_dels_clean, k562_dels_clean], [hepg2_dels_fixed, k562_dels_fixed]):
    for name in dels_dict:
        df = dels_dict[name]
        df["del_id"] = name
        df = df.merge(index_tfbs, on=["del_id", "delpos"], how="left")
        fixed_dict[name] = df
hepg2_dels_fixed["ZFAS1__p1__tile2__plus"].head()


# In[43]:


hepg2_all_dels_tfs = pd.DataFrame()
k562_all_dels_tfs = pd.DataFrame()
for dels_fixed, dels_df, cell in zip([hepg2_dels_fixed, k562_dels_fixed], [hepg2_all_dels_tfs, k562_all_dels_tfs],
                                    ["HepG2", "K562"]):
    print(cell)
    for name in dels_fixed:
        df = dels_fixed[name].sort_values(by="delpos")
        seq = list(df["seq"])
        scores = list(df["mean.log2FC"])
        yerrs = list(df["se"])

        # get wt motif # for reference
        del_id = df.del_id.iloc[0]
        wt_tfs = tfbs_wt[tfbs_wt["del_id"] == del_id]["n_tfs"].iloc[0]
        df["delta_tfs"] = df["n_tfs"] - wt_tfs
        motif_vals = list(df["delta_tfs"])
        dels_df = dels_df.append(df)

        # plot
        if name == "HOTAIR__p1__tile2__minus" and cell == "HepG2":
            plot_peaks_and_tfbs((5.6, 2.5), 94, name, cell, scores, yerrs, 
                                motif_vals, seq, "Fig_3C", True)
        if name == "DLEU1__p1__tile1__plus" and cell == "HepG2":
            plot_peaks_and_tfbs((5.6, 2.5), 94, name, cell, scores, yerrs, 
                                motif_vals, seq, "Fig_S13B", True)
#         if name == "ZFAS1__p1__tile2__plus" and cell == "HepG2":
#             plot_peaks_and_tfbs((5.6, 2.5), 94, name, cell, scores, yerrs, 
#                                 motif_vals, seq, "ZFAS_tile2.for_talk.pdf", True)
            
#         if name == "ZFAS1__p1__tile2__plus" and cell == "HepG2":
#             plot_peaks_and_tfbs((5.6, 1.5), 94, name, cell, scores, yerrs, 
#                                 motif_vals, seq, "ZFAS_tile2.for_poster.pdf", True)
        
    if cell == "HepG2":
        hepg2_all_dels_tfs = dels_df.copy()
    else:
        k562_all_dels_tfs = dels_df.copy()


# ## 8. compare effect sizes w/ number of motifs gained or lost (for all seqs)

# In[44]:


# hepg2
hepg2_all_dels_tfs["delta_tfs_abs"] = np.abs(hepg2_all_dels_tfs["delta_tfs"])
hepg2_all_dels_tfs["del_abs"] = np.abs(hepg2_all_dels_tfs["mean.log2FC"])
hepg2_all_dels_tfs["delta_tfs_log"] = np.log(hepg2_all_dels_tfs["delta_tfs"]+1)
no_nans = hepg2_all_dels_tfs[~pd.isnull(hepg2_all_dels_tfs["mean.log2FC"])]
sig_only = hepg2_all_dels_tfs[hepg2_all_dels_tfs["sig"] == "sig"]
len(sig_only)


# In[45]:


sig_only.head()


# In[47]:


g = sns.jointplot(data=sig_only, x="delta_tfs", y="mean.log2FC", kind="reg", space=0, size=2.5, 
                  stat_func=spearmanr, 
                  marginal_kws={"hist": False}, color="darkgrey", scatter_kws={"s": 25})
print("HepG2")
g.set_axis_labels(r"$\Delta$ motifs (del-ref)", "deletion effect size")

# add n-value
g.ax_joint.annotate("n = %s" % len(sig_only), ha="right", xy=(.95, .05), xycoords=g.ax_joint.transAxes, 
                    fontsize=fontsize)

g.savefig("Fig_3B.pdf", bbox_inches="tight", dpi="figure")


# In[48]:


# k562
k562_all_dels_tfs["delta_tfs_abs"] = np.abs(k562_all_dels_tfs["delta_tfs"])
k562_all_dels_tfs["del_abs"] = np.abs(k562_all_dels_tfs["mean.log2FC"])
k562_all_dels_tfs["delta_tfs_log"] = np.log(k562_all_dels_tfs["delta_tfs"]+1)
no_nans = k562_all_dels_tfs[~pd.isnull(k562_all_dels_tfs["mean.log2FC"])]
sig_only = k562_all_dels_tfs[k562_all_dels_tfs["sig"] == "sig"]
len(sig_only)


# In[49]:


g = sns.jointplot(data=sig_only, x="delta_tfs", y="mean.log2FC", kind="reg", space=0, size=2.5, 
                  stat_func=spearmanr, 
                  marginal_kws={"hist": False}, color="darkgrey", scatter_kws={"s": 25})
print("K562")
g.set_axis_labels(r"$\Delta$ motifs (del-ref)", "deletion effect size")

# add n-value
g.ax_joint.annotate("n = %s" % len(sig_only), ha="right", xy=(.95, .05), xycoords=g.ax_joint.transAxes, 
                    fontsize=fontsize)

g.savefig("Fig_S13A.pdf", bbox_inches="tight", dpi="figure")


# In[ ]:
```
