## Supplementary material for "High-throughput functional analysis of IncRNA core promoters elucidatesrules governing tissue-specificity": 09__tfbs_identification.py

```
# coding: utf-8

# # 09__tbs_identification
# # finding functional TFBSs based on deletion data
# 
# in this notebook, i find "peaks" in the deletion data, defined as any stretch of >= 5 nucleotides with effect sizes of <= -1.5 * the average standard deviation of the deletion effect sizes in that tile. (scaling by the standard deviation ensures that extremely noisy tiles will have higher thresholds for peak calling). then, i intersect the mapped FIMO motifs with these peaks, and consider "functional" motifs to be those that overlap a peak by at least 1 nucleotide. i also limit the motifs to only those that are expressed in the given cell type. i make a heatmap of all of the predicted "functional" TF motifs across all of the lncRNAs we examined, and i examine patterns between tile specificity and the number of functional motifs in the tile.
# 
# ------
# 
# figures in this notebook:
# - **Fig 3E**: barplot of deletion effect sizes and sequence logo plotted proportionally to the loss scores of lncRNA DLEU1 promoter (DLEU1_HepG2). 
# - **Fig 3D, S14**: heatmap showing all of the functional TFBSs
# - **Fig S15**: boxplot showing number of functional motifs in TSSs that are active in only one cell type vs. both

# In[1]:


import warnings
warnings.filterwarnings('ignore')

import pandas as pd
import numpy as np
import matplotlib as mpl
import matplotlib.pyplot as plt
import seaborn as sns
import sys
import time

from os import walk
from scipy.stats import spearmanr

# import utils
sys.path.append("../../utils")
from plotting_utils import *
from misc_utils import *
from norm_utils import *
from del_utils import *

get_ipython().magic('matplotlib inline')


# In[2]:


sns.set(**PAPER_PRESET)
fontsize = PAPER_FONTSIZE


# ## variables

# In[3]:


del_dir = "../../data/05__deletions"
out_dir = "../../data/06__tfbs_results"


# In[4]:


# file w/ tfs and their expression
tf_expr_f = "../../data/04__coverage/TF_tissue_specificities.from_CAGE.txt"


# In[5]:


fimo_f = "../../misc/03__fimo/00__fimo_outputs/pool2_fimo_map.orig.txt"


# In[6]:


index_f = "../../data/00__index/dels_oligo_pool.index.txt"


# In[7]:


peak_signal_cutoff = 1.5
peak_length_cutoff = 5
del_buffer = 11
seq_len = 94


# ## 1. import deletion data and tf expr file

# In[8]:


# hepg2
hepg2_files = []
for (dirpath, dirnames, filenames) in walk("%s/HepG2" % del_dir):
    hepg2_files.extend(filenames)
    break


# In[9]:


# k562
k562_files = []
for (dirpath, dirnames, filenames) in walk("%s/K562" % del_dir):
    k562_files.extend(filenames)
    break


# In[10]:


hepg2_data = {}
k562_data = {}
for files, data, cell in zip([hepg2_files, k562_files], [hepg2_data, k562_data], ["HepG2", "K562"]):
    data_dir = "%s/%s" % (del_dir, cell)
    for f in files:
        df = pd.read_table("%s/%s" % (data_dir, f))
        data[f] = df


# In[11]:


# import tf expr data
tf_expr = pd.read_table(tf_expr_f, sep="\t")
tf_expr.head()


# In[12]:


fimo = pd.read_table(fimo_f, sep="\t")
fimo.columns = ["motif", "unique_id", "start", "stop", "strand", "score", "p-value", "q-value", "seq"]
fimo.head()


# In[13]:


# filter to tfs that are expr in the 2 cell lines
hepg2_tfs = tf_expr[tf_expr["HepG2_exp"] > 0.1]["tf"]
k562_tfs = tf_expr[tf_expr["K562_exp"] > 0.1]["tf"]


# In[14]:


index = pd.read_table(index_f, sep="\t")
index_elem = index[["unique_id", "tile_name", "chr", "tile_start", "tile_end", "strand", "tile_number"]].drop_duplicates()
index_elem.head()


# ## 2. filter dfs to only those that are sig active (wt_class) and make loss score

# In[15]:


def loss_score(row):
    if row["mean.log2FC"] < 0:
        return -row["mean.log2FC"]
    else:
        return 0


# In[16]:


hepg2_data_filt = {}
k562_data_filt = {}
for data, data_filt in zip([hepg2_data, k562_data], [hepg2_data_filt, k562_data_filt]):
    for key in data:
        df = data[key]
        if df.wt_class.iloc[0] == "sig active":
            df["loss_score"] = df.apply(loss_score, axis=1)
            data_filt[key] = df


# ## 3. find "peaks" in the deletion data

# In[17]:


def find_peaks(df, peak_signal_cutoff, peak_length_cutoff):
    peak_status = []
    effect_sizes = list(df["mean.log2FC"])
    avg_sd = df["sd"].mean()
    count = 0
    in_peak = False
    overall_count = 1
    scaled_peak_signal_cutoff = avg_sd * peak_signal_cutoff
    print("sd: %s, scaled cutoff: %s" % (avg_sd, scaled_peak_signal_cutoff))
    for x in effect_sizes:
        if x <= -scaled_peak_signal_cutoff:
            count += 1
            if count >= peak_length_cutoff:
                if in_peak == False:
                    # replace the last peak_length_cutoff elements with "peak"
                    tmp = peak_status[:(-peak_length_cutoff-1)]
                    diff = len(peak_status) - len(tmp)
                    peak_status = list(tmp)
                    peak_status.extend(["peak"] * (diff+1))
                    in_peak = True
                else:
                    peak_status.append("peak")
            else:
                peak_status.append("no peak")
        else:
            in_peak = False
            count = 0
            peak_status.append("no peak")
        overall_count += 1
    df["peak"] = peak_status
    return df


# In[18]:


hepg2_data_peaks = {}
k562_data_peaks = {}

for data_filt, data_peaks, cell in zip([hepg2_data_filt, k562_data_filt], [hepg2_data_peaks, k562_data_peaks],
                                       ["HepG2", "K562"]):
    for key in data_filt:
        name = key.split(".")[0]
        strand = key.split(".")[2]
        gene_name = "%s__%s" % (name, strand)
        print(gene_name)
        df = data_filt[key]
        df = find_peaks(df, peak_signal_cutoff, peak_length_cutoff)
        data_peaks[gene_name] = df
        
        # write file
        cell_dir = "%s/%s/0__peaks" % (out_dir, cell)
        get_ipython().system('mkdir -p $cell_dir')
        df.to_csv("%s/%s.tfbs_peaks.txt" % (cell_dir, gene_name), sep="\t", index=False)


# ## 4. intersect FIMO motifs w/ peaks

# In[19]:


fimo["fixed_name"] = fimo.apply(fix_fimo_names, name_dict=NAME_DICT, loc_dict=LOC_DICT, axis=1)
fimo.head()


# In[20]:


def getOverlap(a, b):
    return max(a[0], b[0]) - min(a[1], b[1])


# In[21]:


hepg2_motif_peaks = {}
k562_motif_peaks = {}

for data_peaks, data_motifs, cell in zip([hepg2_data_peaks, k562_data_peaks], [hepg2_motif_peaks, k562_motif_peaks],
                                         ["HepG2", "K562"]):
    print("")
    print(cell)
    for key in data_peaks:
        print(key)
        df = data_peaks[key]

        # put del_df bps in terms of 1-94
        df["delpos_fixed"] = list(range(1, 95))

        fimo_sub = fimo[(fimo["fixed_name"] == key) & (~fimo["unique_id"].str.contains("DELETION"))
                        & (~fimo["unique_id"].str.contains("SNP_INDIV"))]
        fimo_sub = fimo_sub.sort_values(by="start")
        #print(fimo_sub)
        scores = list(df["mean.log2FC"])
        yerrs = list(df["se"])
        scaled_scores = list(df["loss_score"])
        bases = list(df["seq"])
        motif_positions = list(zip(list(fimo_sub["start"]), list(fimo_sub["stop"]), list(fimo_sub["motif"])))
        print(list(fimo_sub["start"]))
        print(list(fimo_sub["stop"]))

        # get peak positions in tuples
        prev_p = "no peak"
        starts = []
        ends = []
        for i, p in zip(list(df["delpos_fixed"]), list(df["peak"])):
            if p == "peak" and prev_p == "no peak":
                starts.append(i-1)
            elif p == "no peak" and prev_p == "peak":
                ends.append(i-1)
            elif i == 94 and prev_p == "peak":
                ends.append(i)
            prev_p = p
        widths = list(zip(starts, ends))

        motif_positions_neg = [(x-del_buffer, y-del_buffer, m) for x, y, m in motif_positions]
        motif_positions_filt = [(x, y, m) for x, y, m in motif_positions_neg if x > 0 and y < seq_len]
        motif_positions_fixed = [(x, y) for x, y, m in motif_positions_filt]
        motif_names = [m for x, y, m in motif_positions_filt]


        if key == "MEG3__p1__tile2__plus":
            plot_peaks_and_fimo((5.6, 2), seq_len, key, widths, scores, yerrs, scaled_scores, bases, 
                                motif_positions_fixed, motif_names, "MEG3_%s.pdf" % cell, ".", True)
        elif key == "DLEU1__p1__tile2__plus":
            plot_peaks_and_fimo((5.6, 2), seq_len, key, widths, scores, yerrs, scaled_scores, bases, 
                                motif_positions_fixed, motif_names, "DLEU1_%s.pdf" % cell, ".", True)
#             plot_peaks_and_fimo((5.6, 2), seq_len, key, [], scores, yerrs, scaled_scores, bases, 
#                                 motif_positions_fixed, motif_names, "DLEU1_%s.for_talk.pdf" % cell, ".", True)
#         elif key == "ZFAS1__p1__tile2__plus":
#             plot_peaks_and_fimo((5.6, 2), seq_len, key, [], scores, yerrs, scaled_scores, bases, 
#                                 motif_positions_fixed, motif_names, "ZFAS1_%s.for_talk.pdf" % cell, ".", True)
        else:
            plot_peaks_and_fimo((5.6, 2), seq_len, key, widths, scores, yerrs, scaled_scores, bases, 
                                motif_positions_fixed, motif_names, None, None, False)

        motifs = []
        starts = []
        ends = []
        in_peaks = []
        for start, end, motif in motif_positions_filt:
            motifs.append(motif)
            starts.append(start)
            ends.append(end)
            added = False
            for w in widths:
                overlap = getOverlap(w, [start, end])
                if overlap < 0:
                    in_peaks.append("in peak")
                    added = True
                    break
            if not added:
                in_peaks.append("no peak")
        data_motifs[key] = {"motif": motifs, "start": starts, "end": ends, "peak_overlap": in_peaks}
        print({"motif": motifs, "start": starts, "end": ends, "peak_overlap": in_peaks})


# In[22]:


hepg2_motif_dfs = {}
k562_motif_dfs = {}

for data_motifs, dfs, cell in zip([hepg2_motif_peaks, k562_motif_peaks], [hepg2_motif_dfs, k562_motif_dfs], 
                                  ["HepG2", "K562"]):
    for key in data_motifs:
        data = data_motifs[key]
        df = pd.DataFrame.from_dict(data)
        df = df[["motif", "start", "end", "peak_overlap"]]
        df = df.drop_duplicates()
        dfs[key] = df
        
        # write file
        cell_dir = "%s/%s/1__motifs" % (out_dir, cell)
        get_ipython().system('mkdir -p $cell_dir')
        df = df.sort_values(by="start", ascending=True)
        df.to_csv("%s/%s.tfbs_peaks.txt" % (cell_dir, key), sep="\t", index=False)

hepg2_motif_dfs["GAS5__p1__tile2__minus"].head()


# In[23]:


# find total # of tested motifs found to be in peaks
for motif_dfs, cell in zip([hepg2_motif_dfs, k562_motif_dfs], ["HepG2", "K562"]):
    print(cell)
    tot_motifs = 0
    tot_func_motifs = 0
    for key in motif_dfs:
        df = motif_dfs[key]
        tot_motifs += len(df)
        tot_func_motifs += len(df[df["peak_overlap"] == "in peak"])
    print("tot motifs: %s, tot func: %s, perc: %s" % (tot_motifs, tot_func_motifs, tot_func_motifs/tot_motifs))


# ## 5. limit to TFs expressed in each cell line & in peaks

# In[24]:


hepg2_motif_dfs_filt = {}
k562_motif_dfs_filt = {}

for motif_dfs, motif_dfs_filt, tfs in zip([hepg2_motif_dfs, k562_motif_dfs], 
                                          [hepg2_motif_dfs_filt, k562_motif_dfs_filt],
                                          [hepg2_tfs, k562_tfs]):
    for key in motif_dfs:
        df = motif_dfs[key]
        sub = df[(df["motif"].isin(tfs)) & (df["peak_overlap"] == "in peak")]
        motif_dfs_filt[key] = sub

hepg2_motif_dfs_filt["GAS5__p1__tile2__minus"].head()


# ## 6. make heatmap with TFs mapped in each sequence
# note: use only HepG2 since there are more seqs expressed in HepG2 and use results filtered by TFs expr in HepG2

# In[25]:


# first, put all gene data in dictionary of list of dataframes
hepg2_gene_data = {}
k562_gene_data = {}

for tile_data, gene_data in zip([hepg2_motif_dfs_filt, k562_motif_dfs_filt], [hepg2_gene_data, k562_gene_data]):
    for key in tile_data:
        data = tile_data[key]

        if "enhancer" not in key:
            gene_name = key.split("_")[0]
            prom_name = key.split("_")[1]
            tile_name = key.split("_")[2]
            strand_name = key.split("_")[3]
        else:
            gene_name = key.split("_")[0] + "_" + key.split("_")[1]
            prom_name = key.split("_")[2]
            tile_name = key.split("_")[3]
            strand_name = key.split("_")[4]

        data["gene_name"] = gene_name
        data["prom_name"] = prom_name
        data["tile_name"] = tile_name
        data["strand_name"] = strand_name

        if "LINC" in gene_name:
            if "00467" not in gene_name:
                if "enhancer" in gene_name:
                    gene_name = "enhancer_LINC-PINT"
                else:
                    gene_name = "LINC-PINT"
        if gene_name not in gene_data:
            gene_data[gene_name] = [data]
        else:
            current_gene_data = gene_data[gene_name]
            current_gene_data.append(data)

list(hepg2_gene_data.keys())[0:5]


# In[26]:


hepg2_sig_data = {}
k562_sig_data = {}

hepg2_all_motifs = []
k562_all_motifs = []

for sig_data, all_motifs, gene_data in zip([hepg2_sig_data, k562_sig_data], [hepg2_all_motifs, k562_all_motifs],
                                           [hepg2_gene_data, k562_gene_data]):
    for gene in gene_data:
        dfs = gene_data[gene]
        gene_motifs = []
        for df in dfs:
            gene_motifs.extend(list(df["motif"]))
        gene_motifs = list(set(gene_motifs))
        all_motifs.extend(gene_motifs)
        if len(gene_motifs) == 0:
            continue
        sig_data[gene] = gene_motifs


# In[27]:


hepg2_all_motifs = list(set(hepg2_all_motifs))
len(hepg2_all_motifs)


# In[28]:


hepg2_motif_idx_dict = {k:v for k, v in zip(hepg2_all_motifs, list(range(0, len(hepg2_all_motifs))))}


# In[29]:


hepg2_motif_array = np.zeros((len(hepg2_sig_data), len(hepg2_all_motifs)))
for i, gene in enumerate(hepg2_sig_data):
    motif_data = hepg2_sig_data[gene]
    motif_idxs = [hepg2_motif_idx_dict[motif] for motif in motif_data]
    #print(gene)
    for j in motif_idxs:
        hepg2_motif_array[i, j] = 1

hepg2_mo_df = pd.DataFrame(hepg2_motif_array, index=list(hepg2_sig_data.keys()), columns=hepg2_all_motifs)
hepg2_mo_df.head()


# In[30]:


cmap = sns.light_palette("firebrick", reverse=False, as_cmap=True)


# In[31]:


cg = sns.clustermap(hepg2_mo_df, annot=False, cmap=cmap, figsize=(2.25, 3))
cg.savefig("Fig_3D.pdf", bbox_inches="tight", dpi="figure")


# In[32]:


cg = sns.clustermap(hepg2_mo_df.T, annot=False, cmap=cmap, figsize=(5, 12))
cg.savefig("Fig_S14.pdf", bbox_inches="tight", dpi="figure")


# ## 7. plot number of motifs found in seqs expressed in only one cell type vs. two

# In[33]:


expr_in_hepg2_not_k562 = [x for x in hepg2_gene_data.keys() if x not in k562_gene_data.keys()]
expr_in_both = [x for x in hepg2_gene_data.keys() if x in k562_gene_data.keys()]
expr_in_hepg2_not_k562


# In[34]:


hepg2_sig_data.keys()


# In[35]:


results_dict = {}
for gene in expr_in_hepg2_not_k562:
    try:
        sig_motifs = hepg2_sig_data[gene]
        n_sig_motifs = len(sig_motifs)
    except:
        n_sig_motifs = 0
    results_dict[gene] = (n_sig_motifs, "on in hepg2, not k562")
    
for gene in expr_in_both:
    try:
        hepg2_sig_motifs = hepg2_sig_data[gene]
    except:
        hepg2_sig_motifs = []
    try:
        k562_sig_motifs = k562_sig_data[gene]
    except:
        k562_sig_motifs = []
    hepg2_sig_motifs.extend(k562_sig_motifs)
    all_motifs = list(set(hepg2_sig_motifs))
    n_sig_motifs = len(all_motifs)
    results_dict[gene] = (n_sig_motifs, "on in both")

results_df = pd.DataFrame.from_dict(results_dict, orient="index").reset_index()
results_df.columns = ["gene", "n_sig_motifs", "type"]
results_df.head()


# In[36]:


results_df.type.value_counts()


# In[37]:


fig = plt.figure(figsize=(2.5, 2))
ax = sns.boxplot(data=results_df, x="type", y="n_sig_motifs", flierprops = dict(marker='o', markersize=5))
ax.set_xticklabels(["active in one cell type", "active in both cell types"], rotation=30)
mimic_r_boxplot(ax)
plt.xlabel("")
plt.ylabel("# of significant motifs")
plt.ylim((-8, 33))


# calc p-vals b/w dists
one_dist = np.asarray(results_df[results_df["type"] == "on in hepg2, not k562"]["n_sig_motifs"])
both_dist = np.asarray(results_df[results_df["type"] == "on in both"]["n_sig_motifs"])

one_dist = one_dist[~np.isnan(one_dist)]
both_dist = both_dist[~np.isnan(both_dist)]

u, pval = stats.mannwhitneyu(one_dist, both_dist, alternative="less", use_continuity=False)

# statistical annotation
annotate_pval(ax, 0.2, 0.8, 28, 0, 0, pval, fontsize, False, None, None)
ax.text(0, -6, len(one_dist), horizontalalignment='center', color=sns.color_palette()[0])
ax.text(1, -6, len(both_dist), horizontalalignment='center', color=sns.color_palette()[1])

fig.savefig("Fig_S15.pdf", dpi="figure", bbox_inches="tight")


# In[38]:


pval


# ## 8. plot correlation b/w number of motifs found and ref tile activity

# In[39]:


hepg2_dict = {}
k562_dict = {}

for del_dict, motif_dict, d in zip([hepg2_data_peaks, k562_data_peaks], 
                                   [hepg2_motif_dfs, k562_motif_dfs], 
                                   [hepg2_dict, k562_dict]):
    for key in del_dict:
        df = del_dict[key]
        wt_activ = df["wt_activ"].iloc[0]
        
        # find num sig motifs
        motifs = motif_dict[key]
        n_tot_sig = len(list(set(list(motifs["motif"]))))
            
        d[key] = [wt_activ, n_tot_sig]


# In[40]:


hepg2_activ = pd.DataFrame.from_dict(hepg2_dict, orient="index").reset_index()
hepg2_activ.columns = ["seq_name", "activ", "n_sig"]

k562_activ = pd.DataFrame.from_dict(k562_dict, orient="index").reset_index()
k562_activ.columns = ["seq_name", "activ", "n_sig"]


# In[41]:


hepg2_activ.head()
```
