## Supplementary material for "High-throughput functional analysis of IncRNA core promoters elucidatesrules governing tissue-specificity": 10__snps.py

```
# coding: utf-8

# # 10__snps
# # finding regulatory SNPs in MPRA data
# 
# in this notebook, i call regulatory SNPs by comparing the barcode activities for a reference tile to the barcode activities for the alternative tile. i do this per replicate and then combine the p-values and adjust them (see supplemental methods).
# 
# ------
# 
# figures in this notebook:
# - **Fig S19**: volcano plots in HepG2 & K562
# - **Fig S17**: control SNP swarm plots in HepG2 & K562
# - **Fig S22**: GWAS SNP swarm plots in HepG2 & K562
# - **Fig 4B and S18**: scatter plots between number of motifs predicted to be disrupted by FIMO and SNP effect sizes in HepG2 and K562
# - **Fig 4C**: scatter plot showing effect size differences between HepG2 & K562
# - **Fig S20B**: barplot showing significant SNPs found when down-sampling replicates in HepG2
# - **Fig S21**: boxplot showing effect sizes across biotypes

# In[1]:


import warnings
warnings.filterwarnings('ignore')

import numpy as np
import matplotlib as mpl
import matplotlib.pyplot as plt
import pandas as pd
import seaborn as sns
import sys
import time

from decimal import Decimal
from scipy import stats
from scipy.stats import spearmanr
from statsmodels.sandbox.stats import multicomp

# import utils
sys.path.append("../../utils")
from plotting_utils import *
from misc_utils import *
from norm_utils import *
from snp_utils import *

get_ipython().magic('matplotlib inline')


# In[2]:


sns.set(**PAPER_PRESET)
fontsize = PAPER_FONTSIZE
np.random.seed(SEED)


# ## variables

# In[3]:


index_dir = "../../data/00__index"
activ_dir = "../../data/02__activs"
pval_dir = "../../data/03__pvals"


# In[4]:


pool1_index_f = "%s/tss_oligo_pool.index.txt" % index_dir
pool2_index_f = "%s/dels_oligo_pool.index.txt" % index_dir


# In[5]:


hepg2_pool1_activ_f = "%s/POOL1__pMPRA1__HepG2__activities_per_barcode.txt" % activ_dir
k562_pool1_activ_f = "%s/POOL1__pMPRA1__K562__activities_per_barcode.txt" % activ_dir
hepg2_pool2_activ_f = "%s/POOL2__pMPRA1__HepG2__activities_per_barcode.txt" % activ_dir
k562_pool2_activ_f = "%s/POOL2__pMPRA1__K562__activities_per_barcode.txt" % activ_dir


# In[6]:


hepg2_pool1_pvals_f = "%s/POOL1__pMPRA1__HepG2__pvals.txt" % pval_dir
k562_pool1_pvals_f = "%s/POOL1__pMPRA1__K562__pvals.txt" % pval_dir
hepg2_pool2_pvals_f = "%s/POOL2__pMPRA1__HepG2__pvals.txt" % pval_dir
k562_pool2_pvals_f = "%s/POOL2__pMPRA1__K562__pvals.txt" % pval_dir


# In[7]:


# number of times to downsample hepg2
n_samples = 100


# In[8]:


fimo_f = "../../data/04__coverage/pool1_n_motifs_map.txt"


# In[9]:


annot_f = "../../misc/00__tss_properties/mpra_id_to_biotype_map.txt"


# ## 1. import data

# In[10]:


pool1_index = pd.read_table(pool1_index_f, sep="\t")
pool2_index = pd.read_table(pool2_index_f, sep="\t")


# In[11]:


hepg2_pool1_activ = pd.read_table(hepg2_pool1_activ_f, sep="\t")
k562_pool1_activ = pd.read_table(k562_pool1_activ_f, sep="\t")
hepg2_pool2_activ = pd.read_table(hepg2_pool2_activ_f, sep="\t")
k562_pool2_activ = pd.read_table(k562_pool2_activ_f, sep="\t")


# In[12]:


hepg2_pool1_reps = [x for x in hepg2_pool1_activ.columns if x != "barcode"]
k562_pool1_reps = [x for x in k562_pool1_activ.columns if x != "barcode"]
hepg2_pool2_reps = [x for x in hepg2_pool2_activ.columns if x != "barcode"]
k562_pool2_reps = [x for x in k562_pool2_activ.columns if x != "barcode"]

hepg2_pool1_activ["rep_mean"] = np.nanmean(hepg2_pool1_activ[hepg2_pool1_reps], axis=1)
k562_pool1_activ["rep_mean"] = np.nanmean(k562_pool1_activ[k562_pool1_reps], axis=1)
hepg2_pool2_activ["rep_mean"] = np.nanmean(hepg2_pool2_activ[hepg2_pool2_reps], axis=1)
k562_pool2_activ["rep_mean"] = np.nanmean(k562_pool2_activ[k562_pool2_reps], axis=1)

hepg2_pool1_activ.head()


# In[13]:


pool1_index_elem = pool1_index[["oligo_type", "dupe_info", "seq_name", "tile_name", "chr", "locus_start", "locus_end",
                                "strand", "element", "unique_id", "SNP"]].drop_duplicates()
pool2_index_elem = pool2_index[["oligo_type", "dupe_info", "seq_name", "tile_name", "chr", "locus_start", "locus_end",
                                "strand", "element", "unique_id", "SNP"]].drop_duplicates()


# In[14]:


hepg2_pool1_pvals = pd.read_table(hepg2_pool1_pvals_f, sep="\t")
k562_pool1_pvals = pd.read_table(k562_pool1_pvals_f, sep="\t")
hepg2_pool2_pvals = pd.read_table(hepg2_pool2_pvals_f, sep="\t")
k562_pool2_pvals = pd.read_table(k562_pool2_pvals_f, sep="\t")


# In[15]:


fimo = pd.read_table(fimo_f, sep="\t", header=None)
fimo.columns = ["seqID", "n_motifs"]
fimo.head()


# In[16]:


annot = pd.read_table(annot_f, sep="\t")
annot.head()


# ## 2. merge w/ index

# In[17]:


hepg2_pool1_data = pool1_index.merge(hepg2_pool1_activ, on="barcode", how="outer")
k562_pool1_data = pool1_index.merge(k562_pool1_activ, on="barcode", how="outer")
hepg2_pool2_data = pool2_index.merge(hepg2_pool2_activ, on="barcode", how="outer")
k562_pool2_data = pool2_index.merge(k562_pool2_activ, on="barcode", how="outer")
hepg2_pool1_data.head()


# ## 3. extract SNP pairs

# create map of wt unique_id : snp unique_ids

# In[18]:


hepg2_pool1_wt_w_snp_seqs = hepg2_pool1_data[hepg2_pool1_data["oligo_type"].isin(["WILDTYPE_BUT_HAS_SNP", 
                                                                                  "CONTROL_BUT_HAS_SNP"])]
k562_pool1_wt_w_snp_seqs = k562_pool1_data[k562_pool1_data["oligo_type"].isin(["WILDTYPE_BUT_HAS_SNP", 
                                                                               "CONTROL_BUT_HAS_SNP"])]
hepg2_pool2_wt_w_snp_seqs = hepg2_pool2_data[hepg2_pool2_data["oligo_type"].isin(["WILDTYPE_BUT_HAS_SNP", 
                                                                                  "CONTROL_BUT_HAS_SNP"])]
k562_pool2_wt_w_snp_seqs = k562_pool2_data[k562_pool2_data["oligo_type"].isin(["WILDTYPE_BUT_HAS_SNP", 
                                                                               "CONTROL_BUT_HAS_SNP"])]


# In[19]:


hepg2_pool1_wt_w_snp_seqs = hepg2_pool1_wt_w_snp_seqs[["oligo_type", "dupe_info", "seq_name", "tile_name", "chr", 
                                                       "locus_start", "locus_end", "strand", "element", "unique_id", 
                                                       "SNP"]].drop_duplicates()
k562_pool1_wt_w_snp_seqs = k562_pool1_wt_w_snp_seqs[["oligo_type", "dupe_info", "seq_name", "tile_name", "chr", 
                                                     "locus_start", "locus_end", "strand", "element", "unique_id", 
                                                     "SNP"]].drop_duplicates()
hepg2_pool2_wt_w_snp_seqs = hepg2_pool2_wt_w_snp_seqs[["oligo_type", "dupe_info", "seq_name", "tile_name", "chr", 
                                                       "locus_start", "locus_end", "strand", "element", "unique_id", 
                                                       "SNP"]].drop_duplicates()
k562_pool2_wt_w_snp_seqs = k562_pool2_wt_w_snp_seqs[["oligo_type", "dupe_info", "seq_name", "tile_name", "chr", 
                                                     "locus_start", "locus_end", "strand", "element", "unique_id", 
                                                     "SNP"]].drop_duplicates()


# In[20]:


hepg2_pool1_wt_df = hepg2_pool1_wt_w_snp_seqs.copy()
k562_pool1_wt_df = k562_pool1_wt_w_snp_seqs.copy()
hepg2_pool2_wt_df = hepg2_pool2_wt_w_snp_seqs.copy()
k562_pool2_wt_df = k562_pool2_wt_w_snp_seqs.copy()


# In[21]:


print("mapping HepG2 pool1 SNPs")
hepg2_pool1_snp_map = map_snps(hepg2_pool1_wt_w_snp_seqs, pool1_index_elem)
print("mapping K562 pool1 SNPs")
k562_pool1_snp_map = map_snps(k562_pool1_wt_w_snp_seqs, pool1_index_elem)
print("mapping HepG2 pool2 SNPs")
hepg2_pool2_snp_map = map_snps(hepg2_pool2_wt_w_snp_seqs, pool2_index_elem)
print("mapping K562 pool2 SNPs")
k562_pool2_snp_map = map_snps(k562_pool2_wt_w_snp_seqs, pool2_index_elem)


# ## 4. calculate p-values (wilcox test using mean across replicates, 1 for each barcode)

# In[22]:


hepg2_pool1_log2fc_cols = [x for x in hepg2_pool1_pvals.columns if "_log2fc" in x]
k562_pool1_log2fc_cols = [x for x in k562_pool1_pvals.columns if "_log2fc" in x]
hepg2_pool2_log2fc_cols = [x for x in hepg2_pool2_pvals.columns if "_log2fc" in x]
k562_pool2_log2fc_cols = [x for x in k562_pool2_pvals.columns if "_log2fc" in x]


# In[23]:


hepg2_pool1_pvals["mean_log2fc"] = hepg2_pool1_pvals[hepg2_pool1_log2fc_cols].mean(axis=1)
k562_pool1_pvals["mean_log2fc"] = k562_pool1_pvals[k562_pool1_log2fc_cols].mean(axis=1)
hepg2_pool2_pvals["mean_log2fc"] = hepg2_pool2_pvals[hepg2_pool2_log2fc_cols].mean(axis=1)
k562_pool2_pvals["mean_log2fc"] = k562_pool2_pvals[k562_pool2_log2fc_cols].mean(axis=1)


# note if HepG2 -- to compare to other cell types, downsample to 4 reps. do this 100x and take minimum.

# ### active

# In[24]:


min_barcodes = 20
activ_alpha = 0.5
active_l2fc_thresh = 0.5
repr_l2fc_thresh = -0.5


# In[25]:


k562_pool1_active_snp_data = get_snp_results(k562_pool1_reps, k562_pool1_snp_map, k562_pool1_data, k562_pool1_pvals,
                                             min_barcodes, activ_alpha, active_l2fc_thresh, repr_l2fc_thresh, 
                                             "active", "POOL1", 0)


# In[26]:


k562_pool2_active_snp_data = get_snp_results(k562_pool2_reps, k562_pool2_snp_map, k562_pool2_data, k562_pool2_pvals,
                                             min_barcodes, activ_alpha, active_l2fc_thresh, repr_l2fc_thresh, 
                                             "active", "POOL2", 0)


# In[27]:


hepg2_pool1_active_snp_data = get_snp_results(hepg2_pool1_reps, hepg2_pool1_snp_map, hepg2_pool1_data, hepg2_pool1_pvals,
                                              min_barcodes, activ_alpha, active_l2fc_thresh, repr_l2fc_thresh, 
                                              "active", "POOL1", n_samples)


# In[28]:


hepg2_pool2_active_snp_data = get_snp_results(hepg2_pool2_reps, hepg2_pool2_snp_map, hepg2_pool2_data, hepg2_pool2_pvals,
                                              min_barcodes, activ_alpha, active_l2fc_thresh, repr_l2fc_thresh, 
                                              "active", "POOL2", 0)


# ### repressive

# In[29]:


k562_pool1_repressive_snp_data = get_snp_results(k562_pool1_reps, k562_pool1_snp_map, k562_pool1_data, k562_pool1_pvals,
                                                min_barcodes, activ_alpha, active_l2fc_thresh, repr_l2fc_thresh, 
                                                "repressive", "POOL1", 0)


# In[30]:


k562_pool2_repressive_snp_data = get_snp_results(k562_pool2_reps, k562_pool2_snp_map, k562_pool2_data, k562_pool2_pvals,
                                                 min_barcodes, activ_alpha, active_l2fc_thresh, repr_l2fc_thresh, 
                                                 "repressive", "POOL2", 0)


# In[31]:


hepg2_pool1_repressive_snp_data = get_snp_results(hepg2_pool1_reps, hepg2_pool1_snp_map, hepg2_pool1_data, hepg2_pool1_pvals,
                                                  min_barcodes, activ_alpha, active_l2fc_thresh, repr_l2fc_thresh, 
                                                  "repressive", "POOL1", n_samples)


# In[32]:


hepg2_pool2_repressive_snp_data = get_snp_results(hepg2_pool2_reps, hepg2_pool2_snp_map, hepg2_pool2_data, hepg2_pool2_pvals,
                                                  min_barcodes, activ_alpha, active_l2fc_thresh, repr_l2fc_thresh, 
                                                  "repressive", "POOL2", 0)


# ### combine and find sig SNPs

# In[33]:


hepg2_pool1_log2fc_cols = [x for x in hepg2_pool1_active_snp_data.columns if "_l2fc" in x and "combined" not in x]
k562_pool1_log2fc_cols = [x for x in k562_pool1_active_snp_data.columns if "_l2fc" in x and "combined" not in x]
hepg2_pool2_log2fc_cols = [x for x in hepg2_pool2_active_snp_data.columns if "_l2fc" in x and "combined" not in x]
k562_pool2_log2fc_cols = [x for x in k562_pool2_active_snp_data.columns if "_l2fc" in x and "combined" not in x]


# In[34]:


k562_pool1_log2fc_cols


# In[35]:


# filter - require the same direction in at least 75% of non-na replicates
def sig_status(row, col, thresh, l2fc_cols):
    if "NA" in str(row[col]) or pd.isnull(row[col]):
        return "NA__too_many_rep_NAs"
    elif row[col] < thresh:
        l2fcs = list(row[l2fc_cols])
        neg = [x for x in l2fcs if x < 0]
        pos = [x for x in l2fcs if x > 0]
        perc_neg = len(neg)/float(len(neg)+len(pos))
        perc_pos = len(pos)/float(len(neg)+len(pos))
        if perc_neg > 0.75 or perc_pos > 0.75:
            return "sig"
        else:
            return "not sig"
    else:
        return "not sig"


# In[36]:


nums = [0, 1, 2, 3]
active_dfs = [k562_pool1_active_snp_data, k562_pool2_active_snp_data, hepg2_pool1_active_snp_data, hepg2_pool2_active_snp_data]
repressive_dfs = [k562_pool1_repressive_snp_data, k562_pool2_repressive_snp_data, hepg2_pool1_repressive_snp_data, hepg2_pool2_repressive_snp_data]
all_reps = [k562_pool1_reps, k562_pool2_reps, hepg2_pool1_reps, hepg2_pool2_reps]
all_rep_l2fcs = [k562_pool1_log2fc_cols, k562_pool2_log2fc_cols, hepg2_pool1_log2fc_cols, hepg2_pool2_log2fc_cols]

for i, active_df, repressive_df, reps, rep_l2fcs in zip(nums, active_dfs, repressive_dfs, all_reps, all_rep_l2fcs):
    print(i)
    active_df["combined_sig"] = active_df.apply(sig_status, col="combined_padj", thresh=0.05, 
                                                l2fc_cols=rep_l2fcs, axis=1)
    repressive_df["combined_sig"] = repressive_df.apply(sig_status, col="combined_padj", thresh=0.05, 
                                                        l2fc_cols=rep_l2fcs, axis=1)

    for rep in reps:
        active_df["%s_sig" % rep] = active_df.apply(sig_status, col="%s_padj" % rep, thresh=0.05, 
                                                    l2fc_cols=rep_l2fcs, axis=1)
        repressive_df["%s_sig" % rep] = repressive_df.apply(sig_status, col="%s_padj" % rep, thresh=0.05, 
                                                            l2fc_cols=rep_l2fcs, axis=1)


# In[37]:


hepg2_pool1_active_snp_data.combined_sig.value_counts()


# In[38]:


hepg2_pool1_active_snp_data.downsamp_sig.value_counts()


# In[39]:


hepg2_pool1_repressive_snp_data.combined_sig.value_counts()


# ## 5. make summary plot for snps (pool1 active only)

# In[40]:


hepg2_pool1_active_snp_data["type"] = hepg2_pool1_active_snp_data.apply(snp_type, col="combined_sig", axis=1)
k562_pool1_active_snp_data["type"] = k562_pool1_active_snp_data.apply(snp_type, col="combined_sig", axis=1)
hepg2_pool2_active_snp_data["type"] = hepg2_pool2_active_snp_data.apply(snp_type, col="combined_sig", axis=1)
k562_pool2_active_snp_data["type"] = k562_pool2_active_snp_data.apply(snp_type, col="combined_sig", axis=1)

hepg2_pool1_active_snp_data["downsamp_type"] = hepg2_pool1_active_snp_data.apply(snp_type, col="combined_sig", axis=1)


# In[41]:


palette = {"not sig": "gray", "sig": sns.color_palette()[2]}


# In[42]:


for active_snp_data, cell in zip([hepg2_pool1_active_snp_data, k562_pool1_active_snp_data], ["HepG2", "K562"]):
    print(cell)
    snp_data = active_snp_data[(~active_snp_data["combined_sig"].str.contains("NA")) &
                           (~active_snp_data["unique_id"].str.contains("CONTROL")) &
                           (~active_snp_data["unique_id"].str.contains("HAPLO"))].copy()
    snp_data["combined_neg_log_pval"] = -np.log10(snp_data["combined_pval"].astype(float))
    print(snp_data.combined_sig.value_counts())
    g = sns.lmplot(data=snp_data, x="combined_l2fc", y="combined_neg_log_pval", hue="combined_sig", fit_reg=False,
                   palette=palette, size=2.2, scatter_kws={"s": 25})
    g.set_axis_labels("SNP log2(alt/ref)", "-log10(p-value)")
    plt.show()
    plt.close()
    g.savefig("Fig_S19_%s.pdf" % cell, dpi="figure", bbox_inches="tight")


# ## 6. make control snp plots (in pool 1)

# In[43]:


hepg2_pool1_active_snp_data = hepg2_pool1_active_snp_data.merge(pool1_index_elem[["unique_id", "SNP"]], 
                                                                on="unique_id", how="left")
k562_pool1_active_snp_data = k562_pool1_active_snp_data.merge(pool1_index_elem[["unique_id", "SNP"]], 
                                                              on="unique_id", how="left")
print(len(hepg2_pool1_active_snp_data))
print(len(k562_pool1_active_snp_data))

hepg2_pool1_repressive_snp_data = hepg2_pool1_repressive_snp_data.merge(pool1_index_elem[["unique_id", "SNP"]], 
                                                                        on="unique_id", how="left")
k562_pool1_repressive_snp_data = k562_pool1_repressive_snp_data.merge(pool1_index_elem[["unique_id", "SNP"]], 
                                                                        on="unique_id", how="left")
print(len(hepg2_pool1_repressive_snp_data))
print(len(k562_pool1_repressive_snp_data))


# In[44]:


hepg2_pool1_active_sig_data = hepg2_pool1_active_snp_data[hepg2_pool1_active_snp_data["combined_sig"] == "sig"]
hepg2_pool1_repressive_sig_data = hepg2_pool1_repressive_snp_data[hepg2_pool1_repressive_snp_data["combined_sig"] == "sig"]

k562_pool1_active_sig_data = k562_pool1_active_snp_data[k562_pool1_active_snp_data["combined_sig"] == "sig"]
k562_pool1_repressive_sig_data = k562_pool1_repressive_snp_data[k562_pool1_repressive_snp_data["combined_sig"] == "sig"]


# In[45]:


hepg2_pool1_snp_data_all = hepg2_pool1_active_snp_data.append(hepg2_pool1_repressive_snp_data)
hepg2_pool1_ctrl_data_all = hepg2_pool1_snp_data_all[hepg2_pool1_snp_data_all["wt_id"].str.contains("CONTROL")]
hepg2_pool1_ctrl_data_all_grp = hepg2_pool1_ctrl_data_all.groupby(["unique_id", "wt_id", "SNP"])["combined_sig"].apply(list).reset_index()

k562_pool1_snp_data_all = k562_pool1_active_snp_data.append(k562_pool1_repressive_snp_data)
k562_pool1_ctrl_data_all = k562_pool1_snp_data_all[k562_pool1_snp_data_all["wt_id"].str.contains("CONTROL")]
k562_pool1_ctrl_data_all_grp = k562_pool1_ctrl_data_all.groupby(["unique_id", "wt_id", "SNP"])["combined_sig"].apply(list).reset_index()


# In[46]:


hepg2_pool1_ctrl_snps = hepg2_pool1_active_snp_data[hepg2_pool1_active_snp_data["unique_id"].str.contains("CONTROL")]
k562_pool1_ctrl_snps = k562_pool1_active_snp_data[k562_pool1_active_snp_data["unique_id"].str.contains("CONTROL")]

print(len(hepg2_pool1_ctrl_snps))
print(len(k562_pool1_ctrl_snps))


# In[47]:


hepg2_pool1_ctrl_snps_filt = hepg2_pool1_ctrl_snps[~hepg2_pool1_ctrl_snps["combined_sig"].str.contains("NA")]
k562_pool1_ctrl_snps_filt = k562_pool1_ctrl_snps[~k562_pool1_ctrl_snps["combined_sig"].str.contains("NA")]


# In[48]:


hepg2_pool1_data["wt_or_snp"] = hepg2_pool1_data.apply(wt_or_snp, axis=1)
k562_pool1_data["wt_or_snp"] = k562_pool1_data.apply(wt_or_snp, axis=1)


# In[49]:


print("HepG2")
paired_swarmplots_w_pval(7, 4, (7.2, 10), hepg2_pool1_ctrl_snps_filt, hepg2_pool1_data, fontsize, ".", "Fig_S17_1", 
                         True)


# In[50]:


print("K562")
paired_swarmplots_w_pval(7, 4, (7.2, 10), k562_pool1_ctrl_snps_filt, k562_pool1_data, fontsize, ".", "Fig_S17_2", 
                         True)


# ## 7. make GWAS SNP plots

# In[51]:


hepg2_pool1_gwas = hepg2_pool1_active_sig_data[hepg2_pool1_active_sig_data["SNP"].isin(["rs3101018", "rs3785098", "rs4456788"])]
k562_pool1_gwas = k562_pool1_active_sig_data[k562_pool1_active_sig_data["SNP"].isin(["rs3101018", "rs3785098", "rs4456788"])]


# In[52]:


print("HepG2")
paired_swarmplots_w_pval(1, 3, (4.5, 2), hepg2_pool1_gwas, hepg2_pool1_data, fontsize, ".", "Fig_S22_1_tall", True)


# In[53]:


paired_swarmplots_w_pval(1, 3, (4.5, 1.5), hepg2_pool1_gwas, hepg2_pool1_data, fontsize, ".", "Fig_S22_1_short", True)


# In[54]:


hepg2_pool1_gwas.head()


# In[55]:


print("K562")
paired_swarmplots_w_pval(1, 3, (4.5, 2), k562_pool1_gwas, k562_pool1_data, fontsize, ".", "Fig_S22_2_tall", True)


# In[56]:


paired_swarmplots_w_pval(1, 3, (4.5, 1.5), k562_pool1_gwas, k562_pool1_data, fontsize, ".", "Fig_S22_2_short", True)


# ## 8. write files

# In[57]:


out_dir = "../../data/07__snps"


# In[58]:


hepg2_pool2_active_snp_data = hepg2_pool2_active_snp_data.merge(pool2_index_elem[["unique_id", "SNP"]], 
                                                                on="unique_id", how="left")
k562_pool2_active_snp_data = k562_pool2_active_snp_data.merge(pool2_index_elem[["unique_id", "SNP"]], 
                                                              on="unique_id", how="left")

hepg2_pool2_repressive_snp_data = hepg2_pool2_repressive_snp_data.merge(pool2_index_elem[["unique_id", "SNP"]], 
                                                                        on="unique_id", how="left")
k562_pool2_repressive_snp_data = k562_pool2_repressive_snp_data.merge(pool2_index_elem[["unique_id", "SNP"]], 
                                                                      on="unique_id", how="left")


# In[59]:


active_dfs = [k562_pool1_active_snp_data, k562_pool2_active_snp_data, hepg2_pool1_active_snp_data, hepg2_pool2_active_snp_data]
repressive_dfs = [k562_pool1_repressive_snp_data, k562_pool2_repressive_snp_data, hepg2_pool1_repressive_snp_data, hepg2_pool2_repressive_snp_data]
all_reps = [k562_pool1_reps, k562_pool2_reps, hepg2_pool1_reps, hepg2_pool2_reps]
cells = ["K562", "K562", "HepG2", "HepG2"]
pools = ["POOL1", "POOL2", "POOL1", "POOL2"]


# In[60]:


for active_df, repressive_df, reps, cell, pool in zip(active_dfs, repressive_dfs, all_reps, cells, pools):
    print("%s %s" % (cell, pool))
    col_order = ["unique_id", "wt_id", "SNP"]
    wt_meds = [x + "_wt_med" for x in reps]
    wt_meds.extend(["combined_wt_med"])
    snp_meds = [x + "_snp_med" for x in reps]
    snp_meds.extend(["combined_snp_med"])
    l2fcs = [x + "_l2fc" for x in reps]
    l2fcs.extend(["combined_l2fc"])
    pvals = [x + "_pval" for x in reps]
    pvals.extend(["combined_pval"])
    padjs = [x + "_padj" for x in reps]
    padjs.extend(["combined_padj"])
    sigs = [x + "_sig" for x in reps]
    sigs.extend(["combined_sig"])
    if cell == "HepG2" and pool == "POOL1":
        sigs.extend(["downsamp_sig"])
    
    col_order.extend(wt_meds)
    col_order.extend(snp_meds)
    col_order.extend(l2fcs)
    col_order.extend(pvals)
    col_order.extend(padjs)
    col_order.extend(sigs)
    
    active_df = active_df[col_order]
    repressive_df = repressive_df[col_order]
    
    active_f = "%s/%s__%s_active_snp_results.txt" % (out_dir, cell, pool)
    repressive_f = "%s/%s__%s_repressive_snp_results.txt" % (out_dir, cell, pool)
    
    active_df.to_csv(active_f, sep="\t", index=False)
    repressive_df.to_csv(repressive_f, sep="\t", index=False)


# ## 9. create nicer table for supplement

# In[61]:


hepg2_supp = hepg2_pool1_active_snp_data[["SNP", "unique_id", "combined_wt_med", "combined_snp_med", "combined_l2fc",
                                          "combined_padj", "combined_sig", "downsamp_sig"]]
hepg2_supp.columns = ["SNP", "unique_id", "HepG2_ref_activ", "HepG2_alt_activ", "HepG2_effect_size", "HepG2_padj", 
                      "HepG2_sig_status", "HepG2_downsampled_sig_status"]
hepg2_supp.replace("NA__too_many_rep_NAs", np.nan, inplace=True)
print(len(hepg2_supp))
hepg2_supp.sample(5)


# In[62]:


k562_supp = k562_pool1_active_snp_data[["SNP", "unique_id", "combined_wt_med", "combined_snp_med", "combined_l2fc",
                                        "combined_padj", "combined_sig"]]
k562_supp.columns = ["SNP", "unique_id", "K562_ref_activ", "K562_alt_activ", "K562_effect_size", "K562_padj", 
                     "K562_sig_status"]
k562_supp.replace("NA__too_many_rep_NAs", np.nan, inplace=True)
print(len(k562_supp))
k562_supp.sample(5)


# In[63]:


supp_table_s5 = hepg2_supp.merge(k562_supp, on=["SNP", "unique_id"]).drop_duplicates()
print(len(supp_table_s5))
supp_table_s5.sample(5)


# In[64]:


supp_table_s5.to_csv("%s/Supplemental_Table_S5.txt" % out_dir, sep="\t", index=False)


# ## 10. correlate SNPs with fimo motif predictions

# In[65]:


def reverse_snp_map(snp_map):
    rev_map = {}
    for key in snp_map:
        vals = snp_map[key][0]
        for val in vals:
            if val not in rev_map:
                rev_map[val] = key
            else:
                print("dupe val: %s" % val)
    return rev_map


# In[66]:


hepg2_pool1_rev_map = reverse_snp_map(hepg2_pool1_snp_map)
k562_pool1_rev_map = reverse_snp_map(k562_pool1_snp_map)


# In[67]:


fimo.set_index("seqID", inplace=True)
fimo.head()


# In[68]:


fimo_dict = fimo.to_dict(orient="index")


# In[69]:


def get_snp_tfbs_delta(row, rev_map, fimo_map):
    try:
        n_wt_tfs = fimo_map[rev_map[row["unique_id"]]]["n_motifs"]
        n_snp_tfs = fimo_map[row["unique_id"]]["n_motifs"]
    except:
        n_wt_tfs = np.nan
        n_snp_tfs = np.nan
    delta_tfs = n_snp_tfs - n_wt_tfs
    return delta_tfs


# In[70]:


supp_table_s5["delta_tfs"] = supp_table_s5.apply(get_snp_tfbs_delta, rev_map=hepg2_pool1_rev_map, 
                                                 fimo_map=fimo_dict, axis=1)
supp_table_s5.sample(5)


# In[71]:


df = supp_table_s5[supp_table_s5["HepG2_sig_status"] == "sig"]
g = sns.jointplot(data=df, x="delta_tfs", y="HepG2_effect_size", kind="reg", space=0, size=2.5, stat_func=spearmanr, 
                  marginal_kws={"hist": False}, color="darkgrey", scatter_kws={"s": 25})
g.set_axis_labels(r"$\Delta$ motifs (alt-ref)", "SNP effect size")

# add n-value
g.ax_joint.annotate("n = %s" % len(df), ha="right", xy=(.95, .05), xycoords=g.ax_joint.transAxes, 
                    fontsize=fontsize)

g.savefig("Fig_4B.pdf", dpi="figure", bbox_inches="tight")


# In[72]:


df = supp_table_s5[supp_table_s5["K562_sig_status"] == "sig"]
g = sns.jointplot(data=df, x="delta_tfs", y="K562_effect_size", kind="reg", space=0, size=2.5, stat_func=spearmanr, 
                  marginal_kws={"hist": False}, color="darkgrey", scatter_kws={"s": 25})
g.set_axis_labels(r"$\Delta$ motifs (alt-ref)", "SNP effect size")

# add n-value
g.ax_joint.annotate("n = %s" % len(df), ha="right", xy=(.95, .05), xycoords=g.ax_joint.transAxes, 
                    fontsize=fontsize)

g.savefig("Fig_S18.pdf", dpi="figure", bbox_inches="tight")


# ## 11. correlate SNPs across cell types

# In[73]:


def both_type(row):
    if row["HepG2_downsampled_sig_status"] == "sig" and row["K562_sig_status"] == "sig":
        return "sig in both"
    elif row["HepG2_downsampled_sig_status"] == "sig" and row["K562_sig_status"] == "not sig":
        return "sig in HepG2"
    elif row["HepG2_downsampled_sig_status"] == "not sig" and row["K562_sig_status"] == "sig":
        return "sig in K562"
    else:
        return "not sig in both"
    
supp_table_s5["sig_type"] = supp_table_s5.apply(both_type, axis=1)
supp_table_s5.sample(5)


# In[74]:


both_no_null = supp_table_s5[~pd.isnull(supp_table_s5["HepG2_sig_status"]) & ~pd.isnull(supp_table_s5["K562_sig_status"])]
palette = {"not sig in both": "lightgrey", "sig in both": "dimgrey", 
           "sig in HepG2": "salmon", "sig in K562": "firebrick"}
g = sns.lmplot(data=both_no_null, x="HepG2_effect_size", y="K562_effect_size", fit_reg=False, hue="sig_type",
               palette=palette, size=2.2, scatter_kws={"s": 20})
g.set_axis_labels("HepG2 log2(alt/ref)", "K562 log2(alt/ref)")
g.savefig("Fig_4C.pdf", dpi="figure", bbox_inches="tight")


# In[75]:


both_no_null.sig_type.value_counts()


# ## 12. compare results using all HepG2 reps vs. subset

# In[76]:


hepg2_all_sig = len(supp_table_s5[supp_table_s5["HepG2_sig_status"] == "sig"])
hepg2_down_sig = len(supp_table_s5[supp_table_s5["HepG2_downsampled_sig_status"] == "sig"])


# In[77]:


rep_nums = {"HepG2 (Sampled Replicates: 4)": [4, hepg2_down_sig], "HepG2 (All Replicates: 12)": [12, hepg2_all_sig]}
rep_nums = pd.DataFrame.from_dict(rep_nums, orient="index").reset_index()
rep_nums.columns = ["name", "reps", "snps"]
rep_nums.head()


# In[78]:


fig = plt.figure(figsize=(2.2, 2.2))
ax = sns.barplot(data=rep_nums, x="name", y="snps", color="lightgray")
ax.set_xlabel("")
ax.set_ylabel("# significant SNPs")
ax.set_xticklabels(["HepG2 (Sampled Replicates: 4)", "HepG2 (All Replicates: 12)"], rotation=30)
fig.savefig("Fig_S20B.pdf", dpi="figure", bbox_inches="tight")


# ## 13. compare effect sizes across biotypes

# In[79]:


supp_table_s5 = supp_table_s5.merge(annot, left_on="unique_id", right_on="seqID")
supp_table_s5.head()


# In[80]:


supp_table_s5["HepG2_abs_effect_size"] = np.abs(supp_table_s5["HepG2_effect_size"])
supp_table_s5["K562_abs_effect_size"] = np.abs(supp_table_s5["K562_effect_size"])


# In[81]:


fig = plt.figure(figsize=(3.5, 2.5))
ax = sns.boxplot(data=supp_table_s5, x="PromType2", y="HepG2_abs_effect_size", 
                 flierprops = dict(marker='o', markersize=5), order=TSS_CLASS_ORDER, palette=TSS_CLASS_PALETTE)
ax.set_xticklabels(["eRNAs", "lincRNAs", "div. lncRNAs", "mRNAs", "div. mRNAs"], rotation=30)
mimic_r_boxplot(ax)
ax.set_ylim((-0.35, 4))
plt.xlabel("")
plt.ylabel("absolute value(HepG2 effect size)")

# add n-value
ax.annotate(len(supp_table_s5[supp_table_s5["PromType2"] == "Enhancer"]), ha="center", va="top",
            xy=(0, -0.1), fontsize=fontsize, color=TSS_CLASS_PALETTE["Enhancer"])
ax.annotate(len(supp_table_s5[supp_table_s5["PromType2"] == "intergenic"]), ha="center", va="top",
            xy=(1, -0.1), fontsize=fontsize, color=TSS_CLASS_PALETTE["intergenic"])
ax.annotate(len(supp_table_s5[supp_table_s5["PromType2"] == "div_lnc"]), ha="center", va="top",
            xy=(2, -0.1), fontsize=fontsize, color=TSS_CLASS_PALETTE["div_lnc"])
ax.annotate(len(supp_table_s5[supp_table_s5["PromType2"] == "protein_coding"]), ha="center", va="top",
            xy=(3, -0.1), fontsize=fontsize, color=TSS_CLASS_PALETTE["protein_coding"])
ax.annotate(len(supp_table_s5[supp_table_s5["PromType2"] == "div_pc"]), ha="center", va="top",
            xy=(4, -0.1), fontsize=fontsize, color=TSS_CLASS_PALETTE["div_pc"])

fig.savefig("Fig_S21_1.pdf", dpi="figure", bbox_inches="tight")


# In[82]:


fig = plt.figure(figsize=(3.5, 2.5))
ax = sns.boxplot(data=supp_table_s5, x="PromType2", y="K562_abs_effect_size", 
                 flierprops = dict(marker='o', markersize=5), order=TSS_CLASS_ORDER, palette=TSS_CLASS_PALETTE)
ax.set_xticklabels(["eRNAs", "lincRNAs", "div. lncRNAs", "mRNAs", "div. mRNAs"], rotation=30)
mimic_r_boxplot(ax)
ax.set_ylim((-0.45, 5.25))
plt.xlabel("")
plt.ylabel("absolute value(K562 effect size)")

# add n-value
ax.annotate(len(supp_table_s5[supp_table_s5["PromType2"] == "Enhancer"]), ha="center", va="top",
            xy=(0, -0.1), fontsize=fontsize, color=TSS_CLASS_PALETTE["Enhancer"])
ax.annotate(len(supp_table_s5[supp_table_s5["PromType2"] == "intergenic"]), ha="center", va="top",
            xy=(1, -0.1), fontsize=fontsize, color=TSS_CLASS_PALETTE["intergenic"])
ax.annotate(len(supp_table_s5[supp_table_s5["PromType2"] == "div_lnc"]), ha="center", va="top",
            xy=(2, -0.1), fontsize=fontsize, color=TSS_CLASS_PALETTE["div_lnc"])
ax.annotate(len(supp_table_s5[supp_table_s5["PromType2"] == "protein_coding"]), ha="center", va="top",
            xy=(3, -0.1), fontsize=fontsize, color=TSS_CLASS_PALETTE["protein_coding"])
ax.annotate(len(supp_table_s5[supp_table_s5["PromType2"] == "div_pc"]), ha="center", va="top",
            xy=(4, -0.1), fontsize=fontsize, color=TSS_CLASS_PALETTE["div_pc"])

fig.savefig("Fig_S21_2.pdf", dpi="figure", bbox_inches="tight")


# ## 14. make GWAS supplemental table

# In[83]:


tss_snps = pd.read_table("../../misc/04__gwas/tss.snp.ragger.output.txt", sep="\t")
enh_snps = pd.read_table("../../misc/04__gwas/enh.snp.ragger.output.txt", sep="\t")
all_snps = tss_snps.append(enh_snps)


# In[84]:


n_snps_ld = all_snps.groupby(["SNP1 Name"])["Population"].agg("count").reset_index()
n_snps_ld.columns = ["SNP1 Name", "n_snps_in_ld"]
all_snps = all_snps.merge(n_snps_ld, on="SNP1 Name", how="left")
all_snps.head()


# In[85]:


gwas = pd.read_table("../../misc/04__gwas/gwas.catalog.h19.ucsc.txt", sep="\t")


# In[86]:


tmp_snps = all_snps[["SNP1 Name", "SNP2 Name", "R-squared", "D'", "Distance", "n_snps_in_ld"]].drop_duplicates()
tmp_gwas = gwas[["name", "pubMedID", "trait"]].drop_duplicates()
tmp = tmp_snps.merge(tmp_gwas, left_on="SNP1 Name", right_on="name", how="outer")
tmp.head()


# In[87]:


tmp_mpra = supp_table_s5[["SNP", "unique_id", "HepG2_effect_size", "HepG2_sig_status", "K562_effect_size", "K562_sig_status", "delta_tfs", "sig_type"]].drop_duplicates()
supp_table_s6 = tmp_mpra.merge(tmp, left_on="SNP", right_on="SNP2 Name")
supp_table_s6.head()


# In[88]:


supp_table_s6 = supp_table_s6[(supp_table_s6["HepG2_sig_status"] == "sig") | (supp_table_s6["K562_sig_status"] == "sig")]


# In[89]:


supp_table_s6 = supp_table_s6[["unique_id", "SNP", "HepG2_effect_size", "K562_effect_size", "HepG2_sig_status",
                               "K562_sig_status", "SNP1 Name", "R-squared", "D'", "Distance", "n_snps_in_ld", 
                               "pubMedID", "trait"]].drop_duplicates()
supp_table_s6.columns = ["unique_id", "tested_SNP", "HepG2_effect_size", "K562_effect_size", "HepG2_sig_status",
                         "K562_sig_status", "tagged_SNP", "r_squared", "d_prime", "distance_between_SNPs", 
                         "n_snps_in_ld", "pubmed_id", "trait"]
supp_table_s6.head()


# In[90]:


len(supp_table_s6)


# In[91]:


len(supp_table_s6[(supp_table_s6["HepG2_sig_status"] == "sig") & (supp_table_s6["K562_sig_status"] == "sig")])


# In[92]:


supp_table_s6.HepG2_sig_status.value_counts()


# In[93]:


supp_table_s6.K562_sig_status.value_counts()


# In[94]:


supp_table_s6.to_csv("../../data/07__snps/Supplemental_Table_S6.txt", sep="\t", index=False)


# In[ ]:
```
