## Supplementary material for "High-throughput functional analysis of IncRNA core promoters elucidatesrules governing tissue-specificity": 11__snp_del_comparison.py

```
# coding: utf-8

# # 11__snp_del_comparison
# # comparing SNP effect sizes to deletion effect sizes
# 
# in this notebook, i compare SNP effect sizes (log2(alt/ref)) to deletion effect sizes (log2(del/ref)) and also plot deletions and SNPs side-by-side across sequences
# 
# ------
# 
# figures in this notebook:
# - **Fig S16A**: scatter plot b/w SNP and del effect sizes in HepG2 and K562
# - **Fig S16B**: bar plot of del effect sizes with SNP plot below for MEG3

# In[42]:


import warnings
warnings.filterwarnings('ignore')

import numpy as np
import matplotlib as mpl
import matplotlib.pyplot as plt
import pandas as pd
import seaborn as sns
import sys
import time

from decimal import Decimal
from os import walk
from scipy import stats
from scipy.stats import spearmanr
from statsmodels.sandbox.stats import multicomp

# import utils
sys.path.append("../../utils")
from plotting_utils import *
from misc_utils import *
from norm_utils import *
from snp_utils import *
from del_utils import *

get_ipython().magic('matplotlib inline')


# In[43]:


sns.set(**PAPER_PRESET)
fontsize = PAPER_FONTSIZE
np.random.seed(SEED)


# ## variables

# In[44]:


snp_dir = "../../data/07__snps"
del_dir = "../../data/06__tfbs_results"


# In[45]:


hepg2_del_dir = "%s/HepG2/0__peaks" % del_dir
k562_del_dir = "%s/K562/0__peaks" % del_dir


# In[46]:


hepg2_snp_file = "%s/HepG2__POOL2_active_snp_results.txt" % snp_dir
k562_snp_file = "%s/K562__POOL2_active_snp_results.txt" % snp_dir


# In[47]:


index_file = "../../data/00__index/dels_oligo_pool.index.txt"


# ## 1. import data

# In[48]:


hepg2_snps = pd.read_table(hepg2_snp_file, sep="\t")
k562_snps = pd.read_table(k562_snp_file, sep="\t")
hepg2_snps.head()


# In[49]:


index = pd.read_table(index_file, sep="\t")
index.head()


# In[50]:


hepg2_files = []
for (dirpath, dirnames, filenames) in walk(hepg2_del_dir):
    hepg2_files.extend(filenames)
    break


# In[51]:


k562_files = []
for (dirpath, dirnames, filenames) in walk(k562_del_dir):
    k562_files.extend(filenames)
    break


# In[52]:


k562_files[0:5]


# In[53]:


hepg2_dels = {}
k562_dels = {}
for files, f_dir, dels in zip([hepg2_files, k562_files], [hepg2_del_dir, k562_del_dir], [hepg2_dels, k562_dels]):
    for f in files:
        name = f.split(".")[0]
        df = pd.read_table("%s/%s" % (f_dir, f), sep="\t")
        dels[name] = df


# In[54]:


hepg2_dels["ZFAS1__p1__tile2__plus"].head()


# ## 2. parse names in snps & dels so they match

# In[55]:


hepg2_snps["wt_id_new"] = hepg2_snps.apply(fix_snp_names, name_dict=NAME_DICT, loc_dict=LOC_DICT, axis=1)
k562_snps["wt_id_new"] = k562_snps.apply(fix_snp_names, name_dict=NAME_DICT, loc_dict=LOC_DICT, axis=1)
hepg2_snps.sample(5)


# ## 3. find bp num of SNP per tile

# In[56]:


hepg2_wt_seqs = index[index["unique_id"].isin(hepg2_snps["wt_id"])]
hepg2_wt_seqs = hepg2_wt_seqs[["unique_id", "element"]].drop_duplicates()
hepg2_wt_seqs_dict = {k:v for k,v in zip(list(hepg2_wt_seqs["unique_id"]), list(hepg2_wt_seqs["element"]))}
len(hepg2_wt_seqs_dict)


# In[57]:


k562_wt_seqs = index[index["unique_id"].isin(k562_snps["wt_id"])]
k562_wt_seqs = k562_wt_seqs[["unique_id", "element"]].drop_duplicates()
k562_wt_seqs_dict = {k:v for k,v in zip(list(k562_wt_seqs["unique_id"]), list(k562_wt_seqs["element"]))}
len(k562_wt_seqs_dict)


# In[58]:


# same in both, just make map w/ hepg2
wt_seqs_dict = hepg2_wt_seqs_dict.copy()
snp_seqs = index[index["unique_id"].isin(hepg2_snps["unique_id"])]
snp_seqs = snp_seqs[["unique_id", "element"]].drop_duplicates()
snp_seqs_dict = {k:v for k,v in zip(list(snp_seqs["unique_id"]), list(snp_seqs["element"]))}
len(snp_seqs_dict)


# In[59]:


snps_grp = hepg2_snps.groupby("wt_id")["unique_id"].agg("count").reset_index()
snps_grp.sort_values(by="unique_id", ascending=False).head()


# In[60]:


hepg2_snps["snp_pos"] = hepg2_snps.apply(get_snp_pos, wt_seqs_dict=wt_seqs_dict, snp_seqs_dict=snp_seqs_dict, max_snps_per_tile=11,
                                         axis=1)
k562_snps["snp_pos"] = k562_snps.apply(get_snp_pos, wt_seqs_dict=wt_seqs_dict, snp_seqs_dict=snp_seqs_dict, max_snps_per_tile=11,
                                       axis=1)
hepg2_snps.sample(5)


# In[61]:


hepg2_snps.snp_pos.min()


# In[62]:


hepg2_snps.snp_pos.max()


# ## 4. find SNPs that are significant
# for now only look at SNPs that are significantly *down* from active tiles

# In[63]:


hepg2_indiv_snps = hepg2_snps[~hepg2_snps["unique_id"].str.contains("HAPLO")]
hepg2_sig_snps = hepg2_indiv_snps[hepg2_indiv_snps["combined_sig"] == "sig"]
hepg2_not_sig_snps = hepg2_indiv_snps[(hepg2_indiv_snps["combined_sig"] == "not sig")]

k562_indiv_snps = k562_snps[~k562_snps["unique_id"].str.contains("HAPLO")]
k562_sig_snps = k562_indiv_snps[k562_indiv_snps["combined_sig"] == "sig"]
k562_not_sig_snps = k562_indiv_snps[(k562_indiv_snps["combined_sig"] == "not sig")]


# In[64]:


len(hepg2_indiv_snps)


# In[65]:


len(hepg2_sig_snps)


# In[66]:


len(hepg2_not_sig_snps)


# ## 5. overlap peak data

# In[67]:


def get_overlap(row, del_dfs):
    snp_loc = row["snp_pos"]
    try:
        df = del_dfs[row["wt_id_new"]]
    except KeyError:
        return "seq not tested"
    snp_loc_in_df_coords = snp_loc+11
    del_peak_locs = list(df[df["peak"] == "peak"]["delpos"])
    if snp_loc_in_df_coords in del_peak_locs:
        return "peak overlap"
    else:
        return "no peak overlap"


# In[68]:


hepg2_sig_snps["overlap"] = hepg2_sig_snps.apply(get_overlap, del_dfs=hepg2_dels, axis=1)
k562_sig_snps["overlap"] = k562_sig_snps.apply(get_overlap, del_dfs=k562_dels, axis=1)
hepg2_sig_snps.sample(5)


# In[69]:


hepg2_not_sig_snps["overlap"] = hepg2_not_sig_snps.apply(get_overlap, del_dfs=hepg2_dels, axis=1)
k562_not_sig_snps["overlap"] = k562_not_sig_snps.apply(get_overlap, del_dfs=k562_dels, axis=1)
hepg2_not_sig_snps.sample(5)


# In[70]:


for cell, sig_snps in zip(["HepG2", "K562"], [hepg2_sig_snps, k562_sig_snps]):
    print(cell)
    print(sig_snps.overlap.value_counts())
    print("")


# In[71]:


print("significantly down-reg snps only")
for cell, sig_snps in zip(["HepG2", "K562"], [hepg2_sig_snps, k562_sig_snps]):
    print(cell)
    sig_down_snps = sig_snps[sig_snps["combined_l2fc"] < 0]
    print(sig_down_snps.overlap.value_counts())
    print("")


# In[72]:


print("not significant SNPs")
for cell, not_sig_snps in zip(["HepG2", "K562"], [hepg2_not_sig_snps, k562_not_sig_snps]):
    print(cell)
    print(not_sig_snps.overlap.value_counts())
    print("")


# ## 6. which SNPs are reg but not in peaks?

# In[73]:


hepg2_sig_down_snps = hepg2_sig_snps[hepg2_sig_snps["combined_l2fc"] < 0]
k562_sig_down_snps = k562_sig_snps[k562_sig_snps["combined_l2fc"] < 0]
hepg2_sig_down_snps[hepg2_sig_down_snps["overlap"] == "no peak overlap"]


# ## 7. institute l2fc foldchange on snps

# In[74]:


hepg2_sig_down_snps_filt = hepg2_sig_down_snps[hepg2_sig_down_snps["combined_l2fc"] <= -1]
k562_sig_down_snps_filt = k562_sig_down_snps[k562_sig_down_snps["combined_l2fc"] <= -1]
print(len(hepg2_sig_down_snps_filt))
print(len(k562_sig_down_snps_filt))


# In[75]:


hepg2_sig_down_snps_filt.overlap.value_counts()


# In[76]:


k562_sig_down_snps_filt.overlap.value_counts()


# ## 8. plot correlation b/w deletions & snps

# In[77]:


all_hepg2_snp_ids = []
all_k562_snp_ids = []
all_hepg2_del_vals = []
all_hepg2_snp_vals = []
all_k562_del_vals = []
all_k562_snp_vals = []

for all_snp_ids, snps, dels, all_del_vals, all_snp_vals in zip([all_hepg2_snp_ids, all_k562_snp_ids], 
                                                               [hepg2_snps, k562_snps], [hepg2_dels, k562_dels], 
                                                               [all_hepg2_del_vals, all_k562_del_vals], 
                                                               [all_hepg2_snp_vals, all_k562_snp_vals]):
    for seq in list(snps.wt_id_new):
        if seq in dels:
            del_df = dels[seq]

            # put del_df bps in terms of 1-94
            del_df["delpos_fixed"] = list(range(1, 95))

            snp_df = snps[(snps["wt_id_new"] == seq)]
            merged = snp_df[["wt_id_new", "SNP", "snp_pos", 
                             "combined_l2fc", "combined_sig"]].merge(del_df, left_on="snp_pos", 
                                                                     right_on="delpos_fixed")
            snp_ids = list(merged["SNP"])
            del_vals = list(merged["mean.log2FC"])
            snp_vals = list(merged["combined_l2fc"])
            all_snp_ids.extend(snp_ids)
            all_del_vals.extend(del_vals)
            all_snp_vals.extend(snp_vals)

        else:
            continue


# In[78]:


hepg2_snp_del_df = pd.DataFrame()
hepg2_snp_del_df["SNP"] = all_hepg2_snp_ids
hepg2_snp_del_df["del_val"] = all_hepg2_del_vals
hepg2_snp_del_df["snp_val"] = all_hepg2_snp_vals
hepg2_snp_del_df.drop_duplicates(inplace=True)
hepg2_snp_del_nonan = hepg2_snp_del_df[~(pd.isnull(hepg2_snp_del_df["del_val"])) 
                                       & ~(pd.isnull(hepg2_snp_del_df["snp_val"]))]
hepg2_snp_del_nonan.sort_values(by="snp_val").head()


# In[79]:


k562_snp_del_df = pd.DataFrame()
k562_snp_del_df["SNP"] = all_k562_snp_ids
k562_snp_del_df["del_val"] = all_k562_del_vals
k562_snp_del_df["snp_val"] = all_k562_snp_vals
k562_snp_del_df.drop_duplicates(inplace=True)
k562_snp_del_nonan = k562_snp_del_df[~(pd.isnull(k562_snp_del_df["del_val"])) 
                                     & ~(pd.isnull(k562_snp_del_df["snp_val"]))]
k562_snp_del_nonan.sort_values(by="snp_val").head()


# In[80]:


g = sns.jointplot(data=hepg2_snp_del_nonan, x="snp_val", y="del_val", kind="reg", space=0, size=2.2, 
                  stat_func=spearmanr, 
                  marginal_kws={"hist": False}, color="darkgrey", scatter_kws={"s": 25})
g.set_axis_labels("SNP effect size", "deletion effect size")

# add n-value
g.ax_joint.annotate("n = %s" % len(hepg2_snp_del_nonan), ha="right", xy=(.95, .05), xycoords=g.ax_joint.transAxes, 
                    fontsize=fontsize)

g.savefig("Fig_S16A_HepG2.pdf", dpi="figure", bbox_inches="tight")


# In[81]:


g = sns.jointplot(data=k562_snp_del_nonan, x="snp_val", y="del_val", kind="reg", space=0, size=2.2, 
                  stat_func=spearmanr, 
                  marginal_kws={"hist": False}, color="darkgrey", scatter_kws={"s": 25})
g.set_axis_labels("SNP effect size", "deletion effect size")

# add n-value
g.ax_joint.annotate("n = %s" % len(k562_snp_del_nonan), ha="right", xy=(.95, .05), xycoords=g.ax_joint.transAxes, 
                    fontsize=fontsize)

g.savefig("Fig_S16A_K562.pdf", dpi="figure", bbox_inches="tight")


# ## 9. plot overlap of SNPs & deletions

# In[82]:


seq_len = 94

for cell, snps, dels in zip(["HepG2", "K562"], [hepg2_snps, k562_snps], [hepg2_dels, k562_dels]):
    print(cell)
    seqs_w_snps = snps["wt_id_new"].unique()
    
    for seq in seqs_w_snps:
        print(seq)
        try:
            del_df = dels[seq]
        except KeyError:
            print("deletions not tested")
            print("")
            continue

        # put del_df bps in terms of 1-94
        del_df["delpos_fixed"] = list(range(1, 95))

        # print snp ids
        snp_info = snps[(snps["wt_id_new"] == seq)][["SNP", "snp_pos"]]

        # merge dfs
        snp_df = snps[(snps["wt_id_new"] == seq)][["wt_id_new", "combined_l2fc", "combined_sig", "snp_pos"]]
        merged = del_df.merge(snp_df, left_on="delpos_fixed", right_on="snp_pos", how="left")

        # get everything we need to plot
        prev_p = "no peak"
        starts = []
        ends = []
        for i, p in zip(merged["delpos_fixed"], merged["peak"]):
            if p == "peak" and prev_p == "no peak":
                starts.append(i-1)
            elif p == "no peak" and prev_p == "peak":
                ends.append(i-1)
            elif i == 94 and prev_p == "peak":
                ends.append(i)
            prev_p = p
        widths = list(zip(starts, ends))

        scores = list(merged["mean.log2FC"])
        bases = list(merged["seq"])
        yerrs = list(merged["se"])
        scaled_scores = list(merged["loss_score"])

        snp_vals = list(merged["combined_l2fc"].fillna(0))
        snp_sigs = list(merged["combined_sig"].fillna("NA"))

        if "MEG3__p1__tile2__plus" in seq:
            print(snp_info)
            plot_peaks_and_snps((5.6, 2), seq_len, seq, widths, scores, yerrs, scaled_scores, 
                                snp_vals, snp_sigs, bases, "Fig_S16B_%s.pdf" % cell, ".", True)

#         else:
#             plot_peaks_and_snps((5.6, 2), seq_len, seq, widths, scores, yerrs, scaled_scores, 
#                                 snp_vals, snp_sigs, bases, None, None, False)


# In[ ]:
```
