## Supplementary material for "High-throughput functional analysis of IncRNA core promoters elucidatesrules governing tissue-specificity": 12__haplotypes.py

```
# coding: utf-8

# # 12__haplotypes
# # determining whether SNPs act additively vs epistatically
# 
# in this notebook, i determine whether SNPs interact together additively or epistatically. whenever multiple SNPs show up together in a tile (i.e., a "haplotype"), i determine the "expected" (additive) effect size of the haplotype by adding all of the effect sizes of the individual SNPs together. i then bootstrap this expected effect size 1000 times and call those falling outside of the 90% confidence interval as being epistatic.
# 
# ------
# 
# figures in this notebook:
# - **Fig 4D**: swarm plots of 2 representative haplotypes -- one additive and one not

# In[1]:


import warnings
warnings.filterwarnings('ignore')

import itertools
import numpy as np
import matplotlib as mpl
import matplotlib.pyplot as plt
import pandas as pd
import seaborn as sns
import sys
import time

from decimal import Decimal
from os import walk
from scipy import stats
from scipy.stats import spearmanr
from statsmodels.sandbox.stats import multicomp

# import utils
sys.path.append("../../utils")
from plotting_utils import *
from misc_utils import *
from norm_utils import *
from snp_utils import *
from del_utils import *

get_ipython().magic('matplotlib inline')


# In[2]:


sns.set(**PAPER_PRESET)
fontsize = PAPER_FONTSIZE
np.random.seed(SEED)


# ## variables

# In[3]:


hepg2_snp_data_f = "../../data/07__snps/HepG2__POOL1_active_snp_results.txt"
k562_snp_data_f = "../../data/07__snps/K562__POOL1_active_snp_results.txt"


# In[4]:


hepg2_activity_data_f = "../../data/02__activs/POOL1__pMPRA1__HepG2__activities_per_barcode.txt"
k562_activity_data_f = "../../data/02__activs/POOL1__pMPRA1__K562__activities_per_barcode.txt"


# In[5]:


index_f = "../../data/00__index/tss_oligo_pool.index.txt"


# ## 1. import data

# In[6]:


hepg2_snp_data = pd.read_table(hepg2_snp_data_f, sep="\t")
k562_snp_data = pd.read_table(k562_snp_data_f, sep="\t")
hepg2_snp_data.head()


# In[7]:


hepg2_activ = pd.read_table(hepg2_activity_data_f, sep="\t")
k562_activ = pd.read_table(k562_activity_data_f, sep="\t")
hepg2_activ.head()


# In[8]:


hepg2_reps = [x for x in hepg2_activ.columns if x != "barcode"]
k562_reps = [x for x in k562_activ.columns if x != "barcode"]

hepg2_activ["rep_mean"] = np.nanmean(hepg2_activ[hepg2_reps], axis=1)
k562_activ["rep_mean"] = np.nanmean(k562_activ[k562_reps], axis=1)

hepg2_activ.head()


# In[9]:


index = pd.read_table(index_f, sep="\t")
index.head()


# In[10]:


index_elem = index[["oligo_type", "dupe_info", "seq_name", "tile_name", "chr", "locus_start", "locus_end",
                    "strand", "element", "unique_id", "SNP"]].drop_duplicates()
index_elem.head()


# In[11]:


hepg2_data = index.merge(hepg2_activ, on="barcode", how="outer")
k562_data = index.merge(k562_activ, on="barcode", how="outer")
k562_data.head()


# ## 2. find haplotypes

# In[12]:


hepg2_haplo_ids = hepg2_snp_data[hepg2_snp_data["unique_id"].str.contains("WILDTYPE_SNP_PLUS_HAPLO")]["unique_id"]
hepg2_haplo_wts = hepg2_snp_data[hepg2_snp_data["unique_id"].str.contains("WILDTYPE_SNP_PLUS_HAPLO")]["wt_id"]

k562_haplo_ids = k562_snp_data[k562_snp_data["unique_id"].str.contains("WILDTYPE_SNP_PLUS_HAPLO")]["unique_id"]
k562_haplo_wts = k562_snp_data[k562_snp_data["unique_id"].str.contains("WILDTYPE_SNP_PLUS_HAPLO")]["wt_id"]


# In[13]:


hepg2_reps.extend(["combined"])
k562_reps.extend(["combined"])
k562_reps


# In[14]:


hepg2_all_haplo_sig_ids = []
hepg2_all_haplo_sig_wts = []

k562_all_haplo_sig_ids = []
k562_all_haplo_sig_wts = []

for snp_data, reps, all_haplo_sig_ids, all_haplo_sig_wts in zip([hepg2_snp_data, k562_snp_data], 
                                                                [hepg2_reps, k562_reps], 
                                                                [hepg2_all_haplo_sig_ids, k562_all_haplo_sig_ids],
                                                                [hepg2_all_haplo_sig_wts, k562_all_haplo_sig_wts]):
    for rep in reps:
        sig_col = "%s_sig" % rep
        haplo_sig_ids = snp_data[(snp_data["unique_id"].str.contains("WILDTYPE_SNP_PLUS_HAPLO")) & 
                                 (snp_data[sig_col]=="sig")]["unique_id"]
        haplo_wt_ids = snp_data[snp_data["unique_id"].str.contains("WILDTYPE_SNP_PLUS_HAPLO") & 
                                (snp_data[sig_col]=="sig")]["wt_id"]
        all_haplo_sig_ids.append(haplo_sig_ids)
        all_haplo_sig_wts.append(haplo_wt_ids)


# In[15]:


len(k562_all_haplo_sig_ids[-1])


# In[16]:


len(k562_all_haplo_sig_wts[-1])


# ## 3. bootstrap the effect size of haplotypes

# In[17]:


def n_char(row):
    return len(row["SNP"])
hepg2_snp_data["SNP_chars"] = hepg2_snp_data.apply(n_char, axis=1)
k562_snp_data["SNP_chars"] = k562_snp_data.apply(n_char, axis=1)
hepg2_snp_data.sample(5)


# In[18]:


def fixed_names(row):
    if "," in row["SNP"]:
        return "haplotype"
    else:
        return row["SNP"]
hepg2_snp_data["SNP_fixed"] = hepg2_snp_data.apply(fixed_names, axis=1)
k562_snp_data["SNP_fixed"] = k562_snp_data.apply(fixed_names, axis=1)
hepg2_snp_data.sample(5)


# In[19]:


hepg2_wt_seqs = index[index["unique_id"].isin(hepg2_snp_data["wt_id"])]
hepg2_wt_seqs = hepg2_wt_seqs[["unique_id", "element"]].drop_duplicates()
hepg2_wt_seqs_dict = {k:v for k,v in zip(list(hepg2_wt_seqs["unique_id"]), list(hepg2_wt_seqs["element"]))}
len(hepg2_wt_seqs_dict)


# In[20]:


k562_wt_seqs = index[index["unique_id"].isin(k562_snp_data["wt_id"])]
k562_wt_seqs = k562_wt_seqs[["unique_id", "element"]].drop_duplicates()
k562_wt_seqs_dict = {k:v for k,v in zip(list(k562_wt_seqs["unique_id"]), list(k562_wt_seqs["element"]))}
len(k562_wt_seqs_dict)


# In[21]:


hepg2_snp_seqs = index[index["unique_id"].isin(hepg2_snp_data["unique_id"])]
hepg2_snp_seqs = hepg2_snp_seqs[["unique_id", "element"]].drop_duplicates()
hepg2_snp_seqs_dict = {k:v for k,v in zip(list(hepg2_snp_seqs["unique_id"]), list(hepg2_snp_seqs["element"]))}
len(hepg2_snp_seqs_dict)


# In[22]:


k562_snp_seqs = index[index["unique_id"].isin(k562_snp_data["unique_id"])]
k562_snp_seqs = k562_snp_seqs[["unique_id", "element"]].drop_duplicates()
k562_snp_seqs_dict = {k:v for k,v in zip(list(k562_snp_seqs["unique_id"]), list(k562_snp_seqs["element"]))}
len(k562_snp_seqs_dict)


# In[23]:


def get_snp_pos(row, wt_seqs_dict, snp_seqs_dict, max_snps_per_tile):
    wt_id = row["wt_id"]
    snp_id = row["unique_id"]
    wt_seq = wt_seqs_dict[wt_id]
    snp_seq = snp_seqs_dict[snp_id]
    try:
        pos = [i for i in range(len(wt_seq)) if wt_seq[i] != snp_seq[i]][0]
    except:
        pos = [i for i in range(len(snp_seq)) if wt_seq[i] != snp_seq[i]][0]
    pos = pos-10
    return pos


# In[24]:


hepg2_snp_data["snp_pos"] = hepg2_snp_data.apply(get_snp_pos, wt_seqs_dict=hepg2_wt_seqs_dict, 
                                                 snp_seqs_dict=hepg2_snp_seqs_dict, max_snps_per_tile=11, axis=1)
k562_snp_data["snp_pos"] = k562_snp_data.apply(get_snp_pos, wt_seqs_dict=k562_wt_seqs_dict, 
                                               snp_seqs_dict=k562_snp_seqs_dict, max_snps_per_tile=11, axis=1)
k562_snp_data.sample(5)


# In[25]:


def get_num(row, num_dict):
    return num_dict[row["SNP_fixed"]]


# In[26]:


def plot_and_fit_bootstrap(df_w_data, col, name, n_sim, ci_percent, plotname, save):
    median_width = 0.3
    # df no haplo
    df_no_hap = df_w_data[~df_w_data["SNP_fixed"].isin(["haplotype"])]

    # find med. per SNP
    med_per_snp = df_no_hap.groupby(["SNP_fixed"])[col].agg("median").reset_index()
    max_snp = med_per_snp[col].max()
    min_snp = med_per_snp[col].min()
    wt_val = med_per_snp[med_per_snp["SNP_fixed"] == "wildtype"][col].iloc[0]

    med_per_snp = med_per_snp[med_per_snp["SNP_fixed"] != "wildtype"]
    order = ["wildtype"]
    if max_snp > wt_val:
        # order ascending w/ wt first
        SNP_order = med_per_snp.sort_values(by=col, ascending=True)["SNP_fixed"]
    else:
        # order descending w/ wt first
        SNP_order = med_per_snp.sort_values(by=col, ascending=False)["SNP_fixed"]
    order.extend(SNP_order)
    order.extend(["haplotype"])

    # fix snp names
    num_dict = dict(zip(order, list(range(0, len(order)))))
    df_no_hap["SNP_num"] = df_no_hap.apply(get_num, axis=1, num_dict=num_dict)
    df_w_data["SNP_num"] = df_w_data.apply(get_num, axis=1, num_dict=num_dict)
    num_order = list(range(0,len(order)))
    
    # find actual additive effect from data
    wt_median = np.nanmedian(df_w_data[df_w_data["SNP_num"] == 0][col])
    actual_effect = 0
    for s in range(1, len(num_order)-1):
        snp_median = np.nanmedian(df_w_data[df_w_data["SNP_num"] == s][col])
        actual_effect += snp_median - wt_median
    #print("wt med: %s | real effect: %s" % (wt_median, actual_effect))
    
    # bootstrap this additive effect
    effects = np.zeros((1, n_sim))
    for n in range(n_sim):
        effect = 0
        
        # find wt bootstrap med
        vals = np.array(df_w_data[df_w_data["SNP_num"] == 0][col])
        n_vals = len(vals)
        vals_boot = np.random.choice(vals, size=n_vals, replace=True)
        wt_boot_median = np.nanmedian(vals_boot)
        
        # find snp effect on wt
        for s in range(1, len(num_order)-1):
            vals = np.array(df_w_data[df_w_data["SNP_num"] == s][col])
            n_vals = len(vals)
            vals_boot = np.random.choice(vals, size=n_vals, replace=True)
            vals_boot_median = np.nanmedian(vals_boot)
            effect += vals_boot_median - wt_boot_median
        effects[0, n] = effect
    
    diffs = effects - actual_effect
    diffs_sort = np.sort(diffs)   
    
    # find bounds of confidence interval
    lower_percentile = (1 - ci_percent)/2.
    upper_percentile = 1 - lower_percentile
    #print("lower perc: %s | upper perc: %s" % (lower_percentile, upper_percentile))
    
    lower_idx = int(round(lower_percentile * n_sim) - 1)
    upper_idx = int(round(upper_percentile * n_sim) - 1)
    #print("lower idx: %s | upper idx: %s" % (lower_idx, upper_idx))
    
    lower_bound = actual_effect - diffs_sort[0, upper_idx]
    upper_bound = actual_effect - diffs_sort[0, lower_idx]
    #print("lower bound: %s | upper bound: %s" % (lower_bound, upper_bound))
    
    # translate that into an actual activity value
    additive_hap = actual_effect + wt_median
    additive_hap_low = lower_bound + wt_median
    additive_hap_high = upper_bound + wt_median
    #print("haplotype effect: [%s, %s] median: %s" % (additive_hap_low, additive_hap_high, additive_hap))
    
    # record difference b/w actual effect & additive effect
    additiveness = np.abs(actual_effect - additive_hap)
    
    # see if hap median falls w/in bounds
    hap_median = np.nanmedian(df_w_data[df_w_data["SNP_fixed"] == "haplotype"][col])
    if hap_median > additive_hap_low and hap_median < additive_hap_high:
        additive = "additive"
    elif hap_median < additive_hap_low:
        additive = "sub-additive"
    elif hap_median > additive_hap_high:
        additive = "super-additive"

    fig = plt.figure(figsize=(len(df_w_data.SNP.unique())*0.4+1,1))

    # swarm plot - no hap
    ax = sns.swarmplot(x="SNP_num", y=col, data=df_w_data[df_w_data["SNP_fixed"]=="wildtype"], 
                       color="lightgrey", order=num_order, zorder=1)
    sns.swarmplot(x="SNP_num", y=col, data=df_no_hap[df_no_hap["SNP_fixed"]!="wildtype"], 
                  color="darkgrey", 
                  order=num_order, zorder=1, ax=ax)
    ax.set_xlim((-0.5, len(order)))
    
    # swarm plot - hap
    sns.swarmplot(x="SNP_num", y=col, data=df_w_data[df_w_data["SNP_fixed"]=="haplotype"], 
                  color=sns.color_palette()[2], order=num_order, ax=ax, zorder=5)
    
    # median bars
    for tick, text in zip(ax.get_xticks(), ax.get_xticklabels()):
        snp = int(text.get_text())

        # calculate the median value for all replicates of either X or Y
        median_val = df_w_data[df_w_data["SNP_num"]==snp][col].median()

        # plot horizontal lines across the column, centered on the tick
        ax.plot([tick-median_width/2, tick+median_width/2], [median_val, median_val],
                lw=1, color='k', zorder=10)
    
    # fill between - confidence band
    hap_num = num_order[-1]
    x = [hap_num-0.5, hap_num+0.5]
    y1 = [additive_hap_low, additive_hap_low]
    y2 = [additive_hap_high, additive_hap_high]
    plt.fill_between(x, y1, y2, color=sns.color_palette()[2], alpha=0.2, zorder=1)
    
    # line delineating wt and hap
    plt.axvline(x=0.5, linestyle="dashed", color="black", linewidth=1)
    plt.axvline(x=hap_num-0.5, linestyle="dashed", color="black", linewidth=1)
    
    # line delineating bottom and top CIs
    plt.plot(x, y1, color="black", linewidth=1, zorder=1)
    plt.plot(x, y2, color="black", linewidth=1, zorder=1)
    
    # other plot aesthetics
    plt.xlabel("")
    if col != "rep_mean":
        plt.ylabel("MPRA activity")
    else:
        plt.ylabel("MPRA activity")
    title = "%s\n%s\n[additive = %s]" % (name, col, additive)
    plt.title(title)
    labels = ["reference"]
    labels.extend([x for x in order if x != "wildtype"])
    #print(labels)
    
    # find min and max for plot
    plot_min = np.min([df_w_data[col].min(), additive_hap_low])
    plot_max = np.max([df_w_data[col].max(), additive_hap_high])
    
    plt.ylim((plot_min-0.5, plot_max+0.5))
    ax.set_xticklabels(labels, rotation=30)
    if save:
        plt.show()
        fig.savefig("%s.pdf" % (plotname), dpi="figure", bbox_inches="tight")
    else:
        plt.close()
    return additive, additiveness


# In[27]:


len(all_haplo_sig_wts)


# In[28]:


hepg2_all_additive_results = {}
k562_all_additive_results = {}

hepg2_combined_haplo_sig_ids = hepg2_all_haplo_sig_ids[-1]
hepg2_combined_haplo_sig_wts = hepg2_all_haplo_sig_wts[-1]

k562_combined_haplo_sig_ids = k562_all_haplo_sig_ids[-1]
k562_combined_haplo_sig_wts = k562_all_haplo_sig_wts[-1]

zipped = zip([hepg2_combined_haplo_sig_ids, k562_combined_haplo_sig_ids], 
             [hepg2_combined_haplo_sig_wts, k562_combined_haplo_sig_wts],
             [hepg2_all_additive_results, k562_all_additive_results],
             [hepg2_data, k562_data], ["HepG2", "K562"])

for haplo_sig_ids, haplo_sig_wts, all_additive_results, data, cell in zipped:
    print(cell)
    col = "rep_mean"
    for i, wt_id in enumerate(haplo_sig_wts):
        seq_type = wt_id.split("__")[1].split(".")[0]
        flip_type = wt_id.split("__")[1].split(".")[1]
        loc = wt_id.split("__")[2].split(",")[0]
        chrom = loc.split(":")[0]
        start = loc.split(":")[1].split("..")[0]
        end = loc.split(":")[1].split("..")[1]
        strand = wt_id.split("__")[2].split(",")[1]
        name = "%s__%s__%s_%s_%s_%s" % (seq_type, flip_type, chrom, start, end, strand)

        # df finagling
        df = snp_data[snp_data["wt_id"] == wt_id][["unique_id", "SNP", "SNP_chars", "SNP_fixed", "snp_pos"]]
        snps = df[["SNP", "SNP_fixed", "SNP_chars"]].sort_values(by="SNP_chars")["SNP"]
        df_w_data = df.merge(data[["unique_id", "barcode", col]], on="unique_id")
        df_w_data = df_w_data.append(data[data["unique_id"] == wt_id][["unique_id", "barcode", col]])
        df_w_data["SNP_fixed"] = df_w_data["SNP_fixed"].fillna("wildtype")

        if len(df_w_data.SNP.unique()) <= 3:
            # more than 1 SNP does not exist
            continue

        # find min dist between snps
        pos = list(set(list(df["snp_pos"])))
        combos = list(itertools.combinations(pos, 2))
        dists = [np.abs(x[0]-x[1]) for x in combos]
        min_dist = np.min(dists)

        if name in ["Enhancer__noflip__chr13_21050140_21050254_+", "mrna__noflip__chr3_156878514_156878628_-"]:
            save = True
            plotname = "Fig_4D_%s_%s" % (name.split("__")[0], cell)
        else:
            save = False
            plotname = None
        additive, additiveness = plot_and_fit_bootstrap(df_w_data, col, name, 1000, 0.9, plotname, save)
        all_additive_results[name] = (additive, additiveness, min_dist)


# ## 3. find % of additive and non additive haplotypes

# In[29]:


hepg2_results = pd.DataFrame.from_dict(data=hepg2_all_additive_results, orient="index").reset_index()
hepg2_results.columns = ["name", "status", "additiveness", "min_dist"]
hepg2_results.head()


# In[30]:


k562_results = pd.DataFrame.from_dict(data=k562_all_additive_results, orient="index").reset_index()
k562_results.columns = ["name", "status", "additiveness", "min_dist"]
k562_results.head()


# In[31]:


hepg2_results.status.value_counts()


# In[32]:


k562_results.status.value_counts()


# In[ ]:
```
