## Supplementary material for "High-throughput functional analysis of IncRNA core promoters elucidatesrules governing tissue-specificity": README.txt

# some mRNAs were misclassified as not being divergent when they should have been, to fix this i ran:
bedtools closest -S -d -t first -a All.TSS.uniq.mRNAs.bed -b All.TSS.uniq.no_enh.bed | awk '{{OFS="\t"} if ($4 != $10) {print}}' | awk '{{OFS="\t"} {split($4, a,"__")} if ($13 < 1000) {print $4, "div_pc__" a[2] "__" a[3]} else {print $4, $4}}' > All.TSS.uniq.new_div_pc_IDs.txt

where the no_enh file contains all TSSs that do not correspond to enhancers
same thing for intergenic to get new divergent ones

then:
sort -k4,4 old/All.TSS.114bp.uniq.bed | join -a 1 -1 4 -2 1 - All.TSS.uniq.new_div_IDs.txt | awk '{{OFS="\t"} {split($1, a, "__")} if (a[1] == "protein_coding" || a[1] == "intergenic") {print $2, $3, $4, $7, $5, $6} else {print $2, $3, $4, $1, $5, $6}}' | sort | uniq > All.TSS.114bp.uniq.new.bed

to get the new file w/ new IDs
