## Supplementary material for "High-throughput functional analysis of IncRNA core promoters elucidatesrules governing tissue-specificity": README.txt

used bedtools to create files w/ # of motifs, # bp covered, and max coverage as follows:

!!! note: need to use bedtools v. 2.25 -- groupby is broken in 2.26
load modules:
module load centos6
module load bedtools2/2.25.0-fasrc01


for num motifs and num bp covered:
awk '{{OFS="\t"} {print $11, $12, $7, $8, $9, $10}}' ../../misc/03__fimo/00__fimo_outputs/pool1_fimo_map.new_deduped.txt | sort | uniq -f 2 | awk '{{OFS="\t"} {print $3, $4, $5, $6, $1, $2}}' | coverageBed -a ../../data/00__index/tss_oligo_pool.no_random.bed -b stdin > pool1_fimo_map.new_deduped.bp_covered.txt

for max coverage -- FINALLY FIXED:
bedtools intersect -wo -a ../00__index/tss_oligo_pool.no_random.single_bp.bed -b ../../misc/03__fimo/00__fimo_outputs/pool1_fimo_map.new_chip_intersected.new_deduped.txt | awk '{{OFS="\t"} {print $10, $12, $17, $18, $1, $2, $3, $4, $5, $6, $13, $14, $15, $16}}' | sort | uniq -f 4 | awk '{{OFS="\t"} {print $5, $6, $7, $8, $9, $10, $11, $12, $13, $14}}' | sort | uniq | bedtools groupby -g 1,2,3,4 -c 10 -o count_distinct -i stdin | awk '{{OFS="\t"} {print $5, $4}}' | sort -k2,2 -k1,1nr | uniq -f 1 | awk '{{OFS="\t"} {print $2, $1}}' > pool1_fimo_map.new_chip_intersected.new_deduped.max_coverage.txt

note that max coverage requires a bed file of every nucleotide in each 114p bp region (made using coverageBed -d -a promoters.bed -b promoters.bed!)
note that we need a bed file of the promoters used to find motifs and the motifs themselves in 12 column format (from 00__fimo_outputs)
also note that these files *de-dupe* by +/- (so if a motif shows up in exact same place with both + and -, only count once)


*** for the mosbat cluster output ***
bp covered:
coverageBed -a ../../data/00__index/0__all_tss/All.TSS.114bp.uniq.bed -b all_fimo_map.mosbat_cluster_info.merged_with_distinct_count.txt > all_fimo_map.mosbat_clusters.bp_covered.txt

max cov:
intersectBed -wo -a ../../data/00__index/0__all_tss/All.TSS.114bp.uniq.bed -b all_fimo_map.mosbat_cluster_info.merged_with_distinct_count.txt | bedtools groupby -g 4 -c 10 -o max -i stdin > all_fimo_map.mosbat_clusters.max_coverage.txt

where the merged_with_distinct_count file was merged based on the cluster number assigned from the heatmap
